## supporting information for "Negative and positive interspecific interactions involving jellyfish polyps in marine sessile communities"

#### S1 Contrast matrix for compositional manova

We used the following contrast matrix in the ilr transformation for final compositions:

$$\mathbf{V} = \begin{array}{r} \\ \textit{Aurelia aurita} \\ \text{Bare panel} \\ \textit{Botrylloides spp.} \\ \textit{Bugula spp.} \\ \textit{Molgula tubifera} \end{array} \begin{array}{c} \begin{array}{cccc} \mathbf{v}_1 & \mathbf{v}_2 & \mathbf{v}_3 & \mathbf{v}_4 \end{array} \\ \begin{pmatrix} 1/\sqrt{2} & -\sqrt{3/10} & 0 & 0 \\ -1/\sqrt{2} & -\sqrt{3/10} & 0 & 0 \\ 0 & \sqrt{2/15} & 1/\sqrt{6} & 1/\sqrt{2} \\ 0 & \sqrt{2/15} & -2/\sqrt{6} & 0 \\ 0 & \sqrt{2/15} & 1/\sqrt{6} & -1/\sqrt{2} \end{pmatrix} \end{array}. \tag{S1}$$

The columns of $\mathbf{V}$ represent the following logcontrasts: $\mathbf{v}_1$ between *A. aurita* and bare panel; $\mathbf{v}_2$ between the geometric mean of potential competitors and the geometric mean of *A. aurita* and bare panel; $\mathbf{v}_3$ between the geometric mean of ascidians and bryozoans; and $\mathbf{v}_4$ between colonial and solitary ascidians. This is a convenient basis because by setting ilr coordinates 3 and 4 to zero we can obtain the orthogonal projection of a subcomposition onto a subspace that preserves variation between *A. aurita*, bare panel and the geometric mean of potential competitors, but ignores variation among potential competitors (Pawlowsky-Glahn et al., 2015, p. 51). Ternary diagrams in such subspaces are the preferred way to represent marginal relationships among compositions with more than three parts, because this approach is consistent with the Aitchison geometry (van den Boogaart and Tolosana-Delgado, 2013, section 4.2.1). In addition, the choice of $\mathbf{v}_2$ as a logcontrast between geometric means results in a lower-dimensional representation in which differences have a population-dynamic interpretation. For two subcompositions $\mathbf{c}^{(1)}, \mathbf{c}^{(2)}$, a difference in the coefficient of $\mathbf{v}_1$ is proportional to $\log(c_1^{(2)}/c_1^{(1)}) - \log(c_2^{(2)}/c_2^{(1)})$, which has the form of a difference in proportional growth rates between *A. aurita* and bare panel over one unit of time. Similarly, a difference in the coefficient of $\mathbf{v}_2$ is proportional to

$$\frac{1}{3}\left(\log\frac{c_3^{(2)}}{c_3^{(1)}} + \log\frac{c_4^{(2)}}{c_4^{(1)}} + \log\frac{c_5^{(2)}}{c_5^{(1)}}\right) - \frac{1}{2}\left(\log\frac{c_1^{(2)}}{c_1^{(1)}} + \log\frac{c_2^{(2)}}{c_2^{(1)}}\right),$$

which has the form of a difference in mean proportional growth rates between potential competitors and *A. aurita* and bare panel over a unit of time. Thus, differences in such coordinates measure the amount of unequal proportional growth needed to transform one composition into another (Chong and Spencer, 2018). Under the alternative approach of dimension reduction using amalgamations such as $(c_1, c_2, c_3 + c_4 + c_5)$, in which the third part is the sum of relative abundances of potential competitors, there is no such population-dynamic interpretation.

### S2 Coding of treatment effects in compositional manova

For reference, we give the compositional manova model described in the main text again here:

$$\begin{aligned} \mathbf{y}_{jkl} &\sim \text{multinomial}(n_{jkl}, \boldsymbol{\rho}_{jkl}), \\ \boldsymbol{\rho}_{jkl} &= \text{ilr}^{-1}\left(\boldsymbol{\mu} + \boldsymbol{\alpha}_j + \boldsymbol{\beta}_k + \boldsymbol{\gamma}_{jk} + \boldsymbol{\delta}_l + \boldsymbol{\varepsilon}_{jkl}\right), \\ \boldsymbol{\delta}_l &\sim N(\mathbf{0}, \mathbf{Z}), \\ \boldsymbol{\varepsilon}_{jkl} &\sim N(\mathbf{0}, \boldsymbol{\Sigma}). \end{aligned} \tag{S2}$$

We coded treatment effects in the compositional manova using orthogonal contrasts. We represented treatment and depth effects by the matrices $\mathbf{F}, \mathbf{G}$ respectively:

$$\mathbf{F} = \begin{matrix} O \\ C \\ A \end{matrix} \begin{pmatrix} 1 & -1/2 \\ 0 & 1 \\ -1 & -1/2 \end{pmatrix},$$

$$\mathbf{G} = \begin{matrix} 1\,\text{m} \\ 3\,\text{m} \end{matrix} \begin{pmatrix} -1 \\ 1 \end{pmatrix}.$$

The first column of $\mathbf{F}$ represents the difference between the $O$ and $A$ treatments, while the second column represents the difference between the $C$ treatment and the other treatments. The single column of $\mathbf{G}$ represents the difference between depths. We constructed interaction contrasts $\mathbf{H}$ using the Kronecker product (denoted by $\otimes$) of $\mathbf{F}$ and $\mathbf{G}$ (Venables, 2018):

$$\mathbf{H} = \mathbf{F} \otimes \mathbf{G} = \begin{matrix} O, 1\,\mathrm{m} \\ O, 3\,\mathrm{m} \\ C, 1\,\mathrm{m} \\ C, 3\,\mathrm{m} \\ A, 1\,\mathrm{m} \\ A, 3\,\mathrm{m} \end{matrix} \begin{pmatrix} -1 & 1/2 \\ 1 & -1/2 \\ 0 & -1 \\ 0 & 1 \\ 1 & 1/2 \\ -1 & -1/2 \end{pmatrix},$$

in which the first column represents the difference in depth effects between the $O$ and $A$ treatments, and the second column represents the difference in depth effects between treatment $C$ and the other treatments. Then the effects of depth $j$, treatment $k$ and their interaction can be written as

$$\begin{aligned} \boldsymbol{\alpha}_j &= \mathbf{g}_j \boldsymbol{\alpha}, \\ \boldsymbol{\beta}_k &= \mathbf{f}_k \begin{pmatrix} \boldsymbol{\beta}_1 & \boldsymbol{\beta}_2 \end{pmatrix}, \\ \boldsymbol{\gamma}_{jk} &= \mathbf{h}_{jk} \begin{pmatrix} \boldsymbol{\gamma}_1 & \boldsymbol{\gamma}_2 \end{pmatrix}, \end{aligned}$$

where $\mathbf{g}_j$, $\mathbf{f}_k$ and $\mathbf{h}_{jk}$ are the row of $\mathbf{G}$ corresponding to depth $j$, the row of $\mathbf{F}$ corresponding to treatment $j$ and the row of $\mathbf{H}$ corresponding to the combination of depth $j$, treatment $k$, respectively, and $\boldsymbol{\alpha}, \boldsymbol{\beta}_1, \boldsymbol{\beta}_2, \boldsymbol{\gamma}_1, \boldsymbol{\gamma}_2$ are parameter vectors.

### S3 Priors for compositional manova

For the depth effect $\boldsymbol{\alpha}$, we used prior information from Chong and Spencer (2018). In that study, a Bayesian compositional linear model was used to model the relationship between community composition and depth, for communities on the dock wall of our study site. We took the posterior distribution of predicted differences in composition between communities at 3 m and 1 m and selected the subcomposition approximately corresponding to those taxa abundant enough to analyze at the end of our experiment. The Chong and Spencer (2018) model was based on amalgamated taxa. We therefore used the colonial ascidians taxon to represent *Botrylloides spp.* and the solitary ascidians taxon to represent *Molgula tubifera.* The remaining components (*Bugula spp.*, *Aurelia aurita* and bare wall/bare panel) had obvious matches between our data and Chong and Spencer (2018). We transformed to logratio coordinates using the contrast matrix $\mathbf{V}$ given in Equation S1. Pairwise scatter plots and marginal kernel density plots of a sample drawn from the posterior distribution of this difference suggested that a multivariate normal would be a reasonable approximation (Figure S1, green). We chose a multivariate normal prior centred on the posterior mean vector from Chong and Spencer (2018). However, because the experiment was done in a different year and on a different substrate, and the taxa in Chong and Spencer (2018) were amalgamated, we made our prior more diffuse and with a weaker correlation structure (Figure S1, orange):

$$\boldsymbol{\alpha} \sim N(\boldsymbol{\mu}_\alpha, \mathrm{diag}(2\mathbf{s})\mathbf{R}\,\mathrm{diag}(2\mathbf{s})),$$

where $\boldsymbol{\mu}_\alpha = (2.8, -2.6, -0.1, 0.1)^T$ is the sample posterior mean vector, $\mathbf{s}$ is the vector of sample posterior standard deviations, and $\mathbf{R}$ is a correlation matrix whose off-diagonal elements are all set to half the sample mean posterior correlation among all pairs of components, giving the prior covariance matrix

$$\boldsymbol{\Sigma}_\alpha = \mathrm{diag}(2\mathbf{s})\mathbf{R}\,\mathrm{diag}(2\mathbf{s})) = \begin{pmatrix} 12.7 & 3.2\times10^{-4} & 5.5\times10^{-5} & 8.5\times10^{-5} \\ 3.2\times10^{-4} & 7.1 & 4.1\times10^{-5} & 6.4\times10^{-5} \\ 5.5\times10^{-5} & 4.1\times10^{-5} & 0.21 & 1.1\times10^{-5} \\ 8.5\times10^{-5} & 6.4\times10^{-5} & 1.1\times10^{-5} & 0.5 \end{pmatrix}$$

This is close to a diagonal covariance matrix. Although it is a reasonable description of most aspects of the posterior distribution from Chong and Spencer (2018), it does not capture the strong correlation between $\alpha_1$ and $\alpha_2$. This is probably an advantage: we are not confident that data from an observational study in a different year are sufficient to support very strong prior information.

For the covariance matrices $\boldsymbol{\Sigma}$ and $\mathbf{Z}$ of the panel effects $\boldsymbol{\varepsilon}$ and block effects $\boldsymbol{\delta}$ respectively, we used generic weakly-informative priors, with independent half-Cauchy$(0, 2.5)$ priors on the standard deviations and an LKJ prior (Lewandowski et al., 2009) with scale parameter 2 on the correlation matrix. These were the same as the priors on variation among dock wall images in Chong and Spencer (2018). We did not use posterior estimates of the corresponding effect for dock wall images from Chong and Spencer (2018) because our panels were smaller than the dock wall images, and in past experiments (Maxatova, 2016; Edney, 2017; Sharpe, 2020), there appeared to be more variation among panels than adjacent areas of dock wall. We used the same prior for block effects because previous experiments (Maxatova, 2016; Edney, 2017; Presser, 2019; Sharpe, 2020) suggested that among-block differences could plausibly be as large as among-panel differences.

For the overall mean $\boldsymbol{\mu}$ in ilr coordinates, we chose independent univariate $N(0, 4)$ priors on each coordinate (where the second parameter is the variance). Among past experiments (Maxatova, 2016; Edney, 2017; Presser, 2019; Sharpe, 2020), typical composition was quite variable from year to year, with some taxa almost absent in some years and abundant in others. Centring priors on zero means that the most likely composition is equal relative abundance of each component. We chose the variance 4 by experimenting with simulated data, so that after back-transformation from ilr coordinates, compositions having almost none of a component or being almost filled by it were possible (with 80% of the probability for any component between relative abundances of about 0.004 and 0.6).

For the treatment effects $\boldsymbol{\beta}_1, \boldsymbol{\beta}_2$ and treatment by depth interactions $\boldsymbol{\gamma}_1, \boldsymbol{\gamma}_2$ we used independent univariate $N(0, 1)$ priors on each ilr coordinate. Based on the fact that we were aiming to remove half of the target taxon, changes at least as large as doubling or halving a single component should be highly plausible. We calculated the squared Aitchison distance (Aitchison, 1992) between the zero vector in the simplex and a vector in which a single component is doubled or halved. This is equal to the squared Euclidean distance from the origin in ilr coordinates (Egozcue et al., 2003). The probability of a random vector lying at least this squared distance $d^2$ from the origin, for a 4-dimensional multivariate normal distribution with mean vector $\mathbf{0}$ and covariance matrix $c\mathbf{I}_4$, is given by $P(\chi^2_4 \geq \frac{d^2}{c})$ (e.g. Härdle and Simar, 2019, Theorem 4.7). This probability is large for $c = 1$ (Figure S2, green line), and in fact, larger changes in a single component are also likely (Figure S2, orange, purple and pink lines). This prior is therefore sufficiently broad to cover likely scraping treatment effects. In the absence of detailed knowledge about the likely size of interactions between scraping treatments and depth, we used the same prior for this interaction.

### S4 Fitting, checking and calibration for compositional manova

We fitted the compositional manova using `cmdstan 2.23.0` (Carpenter et al., 2017). We ran 4 chains for 2500 warmup and 2500 sampling iterations. This took approximately 90 s on a 64-bit Ubuntu 20.04 system with 4 Intel Xeon 3.2 GHz cores and 16 GiB RAM. Effective sample size was at least 1242 (median 7561) and the potential scale reduction statistic was no larger than 1.003 (median 1.000), for all parameters (these values are for the full model: similar values were obtained for the simplified model reported in the results section). Inspection of trace plots did not reveal any evidence of failure to converge. Processing and visualization of samples from the posterior was done in R version 4.1.0 (R Core Team, 2020), with the packages `rstan 2.21.2` (Stan Development Team, 2020) and `compositions 2.0-2` (van den Boogaart et al., 2020).

We checked the assumption of multivariate normal distributions on the hierarchical block effects $\boldsymbol{\delta}$ and panel effects $\boldsymbol{\varepsilon}$ using $QQ$ plots of squared Mahalanobis distance from the origin against quantiles of the $\chi^2(4)$ distribution, for samples of size 1000 from the posterior. No large deviations from multivariate normality were apparent.

We carried out two graphical posterior predictive checks of the plausibility of the model: plots of sample proportions of each taxon from posterior predictive simulations against sample proportions from the real data, and plots of density curves for sample versions of the logit difference statistics described below (calculated using sample proportions with add-one pseudocounts, rather than by re-fitting models), for real data and posterior predictive simulations. Neither of these revealed strong departures of posterior predictive simulated data from the observed data.

We checked the performance of the sampling algorithm for the full model using simulation-based calibration (Talts et al., 2018), implemented in the `sbc()` function in `rstan 2.21.2`. We conditioned 500 times on draws from the prior predictive distribution, ran Stan on each draw for 2500 warmup and 2500 sampling iterations as above (taking approximately 32 h on a 64-bit Ubuntu 20.04 system with 4 Intel Xeon 3.2 GHz cores and 16 GiB RAM), thinned using the default value of 3, and inspected histograms of rank statistics for true parameters within the posterior samples. No large deviations from uniformity were apparent in the histograms of rank statistics, consistent with absence of bias in sampling from the posterior.

### S5 Model comparison for compositional manova

We compared versions of the compositional manova using leave-one-cluster-out cross-validation. The loss function was the Bayesian leave-one-cluster-out estimate of out-of-sample prediction error (expected log predictive density) $\mathrm{elpd}_{\mathrm{loco}}$, where a cluster corresponds to a block of panels:

$$\mathrm{elpd}_{\mathrm{loco}} = \sum_{l=1}^{L} \log f(\mathbf{y}_l|\mathbf{y}_{-l}).$$

Here, $L$ is the number of blocks and $f(\mathbf{y}_l|\mathbf{y}_{-l})$ is the posterior density of the $l$th block $\mathbf{y}_l$, given the data set $\mathbf{y}_{-l}$ in which block $l$ is excluded (Vehtari et al., 2017). The leave-one-cluster-out posterior density is

$$f(\mathbf{y}_l|\mathbf{y}_{-l}) = \int f(\mathbf{y}_l|\boldsymbol{\theta}) f(\boldsymbol{\theta}|\mathbf{y}_{-l}) \, \mathrm{d}\boldsymbol{\theta}, \tag{S3}$$

where $\boldsymbol{\theta}$ is the parameter vector $\{\boldsymbol{\mu}, \boldsymbol{\alpha}, \boldsymbol{\beta}_1, \boldsymbol{\beta}_2, \boldsymbol{\gamma}_1, \boldsymbol{\gamma}_2, \boldsymbol{\Sigma}, \mathbf{Z}\}$, $f(\mathbf{y}_l|\boldsymbol{\theta})$ is the density of the $l$th block given parameter vector $\boldsymbol{\theta}$, and $f(\boldsymbol{\theta}|\mathbf{y}_{-l})$ is the posterior density of $\boldsymbol{\theta}$ estimated from all blocks other than $l$. To estimate $f(\mathbf{y}_l|\boldsymbol{\theta})$, we need to integrate over the unknown block and panel effects for each new panel in a new block (with one panel from each of two depths and three removal treatments):

$$f(\mathbf{y}_l|\boldsymbol{\theta}) = \prod_{j=1}^{2} \prod_{k=1}^{3} \int \int f(\boldsymbol{\delta}|\boldsymbol{\theta}) f(\boldsymbol{\varepsilon}|\boldsymbol{\theta}) f(\mathbf{y}_{jkl}|\boldsymbol{\theta}, \boldsymbol{\delta}, \boldsymbol{\varepsilon}) \, \mathrm{d}\boldsymbol{\varepsilon} \, \mathrm{d}\boldsymbol{\delta}, \tag{S4}$$

where $f(\boldsymbol{\delta}|\boldsymbol{\theta})$ and $f(\boldsymbol{\varepsilon}|\boldsymbol{\theta})$ are the multivariate normal densities of block and panel effects, with mean vector $\mathbf{0}$ and covariance matrices $\mathbf{Z}$ and $\boldsymbol{\Sigma}$ respectively, and $f(\mathbf{y}_{jkl}|\boldsymbol{\theta}, \boldsymbol{\delta}, \boldsymbol{\varepsilon})$ is the multinomial probability for the counts from the panel at depth $j$, removal treatment $k$ in block $l$ (Equation S2). We estimated the integral in Equation S3 using samples of size 1000 from the posterior density of $f(\boldsymbol{\theta}|\mathbf{y}_{-l})$, obtained using Stan as described in section S4, on data sets with each block left out in turn. For each draw from each of these posterior densities, we estimated the integrals in Equation S4 by Monte Carlo integration in R, using samples of size 1000 from $f(\boldsymbol{\delta}|\boldsymbol{\theta})$ and $f(\boldsymbol{\varepsilon}|\boldsymbol{\theta})$. We compared models using the `loo_compare()` function in the R package `loo` version 2.4.1 (Vehtari et al., 2020). Computation of $\mathrm{elpd}_{\mathrm{loco}}$ for all the models considered took approximately 44 h on a 64-bit Ubuntu 20.04 system with 4 Intel Xeon 3.2 GHz cores and 16 GiB RAM.

### S6 Visualization of compositional manova results

In the compositional manova, let $\boldsymbol{\phi}'_{jk} = \boldsymbol{\alpha}'_j \oplus \boldsymbol{\beta}'_k \oplus \boldsymbol{\gamma}'_{jk} \in \mathbb{S}^4$ denote a treatment effect (the effect of the combination of depth $j$ and removal treatment $k$). Let $\mathbf{e}_1 = (1,0,0,0)^T, \ldots, \mathbf{e}_4 = (0,0,0,1)^T$ be the standard basis vectors in $\mathbb{R}^4$. Then a basis for $\mathbb{S}^4$ is given by the vectors $\mathbf{e}'_i = \mathrm{ilr}^{-1}\,\mathbf{e}_i, i = 1, \ldots, 4$. The orthogonal projection of a treatment effect onto a subset of these basis vectors with indices $S$ is given by

$$\bigoplus_{i \in S} \langle \boldsymbol{\phi}'_{jk}, \mathbf{e}'_i \rangle_a \odot \mathbf{e}'_i,$$

where $\bigoplus_{i \in S}$ denotes repeated perturbation, $\langle \cdot, \cdot \rangle_a$ denotes the Aitchison inner product (Egozcue et al., 2003) and $\odot$ represents the powering operator (Aitchison, 1986, p. 120). With $S = \{1, 2\}$, we obtain the projection of treatment effects onto the 2-simplex with parts representing *A. aurita*, bare panel and gm (potential competitors), where gm () denotes the geometric mean. With $S = \{3, 4\}$, we obtain treatment effects on the subcomposition of potential competitors, with parts *Botrylloides spp.*, *Bugula spp.* and *Molgula tubifera.*

We represented each of these projections of treatment effects in a ternary plot. We calculated 95 % highest posterior density credible regions for the projections of treatment effects using the algorithm in Hyndman (1996), implemented in the `hdr.2d()` function in R package `hdrcde 3.4` (Hyndman, 2018). We plotted the corresponding observations in the form $\hat{\boldsymbol{\rho}}_{ijkl} \ominus \hat{\boldsymbol{\mu}}'$, where $\hat{\boldsymbol{\rho}}_{ijkl}$ denotes the sample estimate of the relative abundance vector in panel $i$ from depth $j$, treatment $k$, block $l$ (with zeros replaced by $1/2$, which is half the detection limit), $\hat{\boldsymbol{\mu}}'$ denotes the estimated posterior mean overall metric centre from the manova, and $\ominus$ is the compositional difference operator (Egozcue et al., 2003).

We visualized the panel and block covariance matrices ($\boldsymbol{\Sigma}$ and $\mathbf{Z}$ respectively) by constructing ellipses of unit Mahalanobis distance around the origin in the first two and last two ilr coordinates. We constructed 100 such ellipses using draws from the posterior distributions of the corresponding submatrices of $\boldsymbol{\Sigma}$ and $\mathbf{Z}$, back-transformed and plotted them in ternary plots.

### S7 Measuring treatment effects

We assessed the effects of potential competitors on *A. aurita* using differences in logit (*A. aurita*) between potential competitor removal ($O$) and control ($C$) treatments. Similarly, we assessed the effects of *A. aurita* on potential competitors using differences in logit (potential competitors) between *A. aurita* removal ($A$) and control ($C$) treatments.

Differences in treatment metric centres from the manova are not directly relevant to our hypotheses, because we expect the metric centre to be affected by a removal treatment, even if the non-target taxon does not respond to this removal. In contrast, logit differences take the value zero if the non-target taxon does not respond. For example, consider the map $f_O$ representing removal of half the potential competitors:

$$\begin{aligned} &f_O : \mathbb{S}^4 \to \mathbb{S}^4, \\ &(c_1, c_2, c_3, c_4, c_5) \mapsto \left( c_1, c_2 + \frac{1}{2}(c_3 + c_4 + c_5), \frac{1}{2}c_3, \frac{1}{2}c_4, \frac{1}{2}c_5 \right). \end{aligned} \tag{S5}$$

The removal treatment has no direct effect on $\text{logit}(c_1) = \log(c_1/(1 - c_1))$, because we are simply turning some of the space occupied by potential competitors into bare panel, without changing $1 - c_1$. The same applies to repeated application of this treatment. Let $\text{logit}(c_1)_O$ and $\text{logit}(c_1)_C$ denote the values of $\text{logit}(c_1)$ in the $O$ and $C$ treatments respectively, at the end of the experiment. Under the null hypothesis that potential competitor removal does not affect *A. aurita*,

$$\text{logit}(c_1)_O - \text{logit}(c_1)_C = 0.$$

However, if potential competitors have a negative effect on *A. aurita*, we expect

$$\text{logit}(c_1)_O - \text{logit}(c_1)_C > 0.$$

Similarly, if *A. aurita* has a negative effect on potential competitors, we expect

$$\text{logit}(c_3 + c_4 + c_5)_A - \text{logit}(c_3 + c_4 + c_5)_C > 0.$$

We therefore calculated the posterior distributions of the logit difference statistics $\text{logit}(c_1)_O - \text{logit}(c_1)_C$ and $\text{logit}(c_3 + c_4 + c_5)_A - \text{logit}(c_3 + c_4 + c_5)_C$ for each treatment combination at each depth. We plotted these distributions using kernel density estimates (obtained using the `density()` function in R with default parameters). We calculated 95 % highest posterior density credible intervals using the `hdr()` function in R package `hdrcde 3.4` (Hyndman, 2018).

Removal treatments such as repeated application of $f_O$ (Equation S5) are settling processes (Pawlowsky-Glahn et al., 2015, section 9.3), which are not linear in the Aitchison geometry, even under the null response. The logit difference statistics which we use to measure the response, and models of community dynamics such as Equations S9 below, which determine whether responses are non-null, are also nonlinear in the Aitchison geometry. Nevertheless, the two-way manova model with interaction (Equation 2) can describe any pattern of variation in metric centres among treatments. Thus, our manova model can represent the outcome of removal treatments and resulting responses in community dynamics, but not the underlying mechanism.

### S8 Model for community dynamics including polyp growth on potential competitors

In the main text, we do not include polyp growth on potential competitors, because this was very rarely observed in the experimental data (although it is common in some years). Here, we describe and analyze the basic properties of a model with a third state variable representing polyps on potential competitors (the model we specified before collecting data), because it may be useful for future work. We show that in this model, it is possible for potential competitors to have either a positive or a negative effect on the proportional population growth rate of polyps growing on potential competitors, depending on parameter values.

To the model in the main text we add a third state variable, the density $y_2$ of *A. aurita* polyps on potential competitors, per unit surface area of substrate (numbers $\text{L}^{-2}$). We separate polyp density into those on substrate and those on potential competitors because polyps are able to settle on some organisms such as the ascidian

*Ascidiella aspersa* and the bivalve *Mytilus edulis*. In this model, increases in the proportion of substrate filled by potential competitors could potentially have either a negative or a positive net effect on polyps, depending on the amount of new surface area created and the ability of polyps to settle on this new area.

Our updated model is

$$\frac{\mathrm{d}x}{\mathrm{d}t} = a_0 (1 - x - \delta y_1) + a_1 x (1 - x - \delta y_1) + a_2 x, \tag{S6}$$

$$\frac{\mathrm{d}y_1}{\mathrm{d}t} = b_0 (1 - x - \delta y_1) + b_1 y_1 (1 - x - \delta y_1) + b_2 y_1 + b_3 y_2 (1 - x - \delta y_1), \tag{S7}$$

$$\frac{\mathrm{d}y_2}{\mathrm{d}t} = c_0 (\psi x - \delta y_2) + c_1 y_2 (\psi x - \delta y_2) + a_2 \psi x y_2 + c_2 y_2 + c_3 y_1 (\psi x - \delta y_2). \tag{S8}$$

The processes included in this model are sketched in Figure S3.

The dynamics of polyps on substrate (Equation S7) are the same as those of the basic model in the main text, apart from an additional term representing the contribution of budding by polyps on potential competitors to polyps on substrate, with proportional rate given by the positive parameter $b_3$ ($\mathrm{T}^{-1}$). The dynamics of polyps on potential competitors (Equation S8) have the same general form as Equation S7, apart from two key differences. First, the surface area unoccupied by polyps on competitors per unit substrate surface area is given by $\psi x - \delta y_2$, where the positive parameter $\psi$ (dimensionless) is the surface area available for occupation by polyps per unit substrate area occupied by potential competitors. The parameter $\psi$ would be equal to 1 for a perfectly flat potential competitor, but will generally be greater than 1. For example, a hemispherical potential competitor would have $\psi = 2$, because such an organism with radius $r$ would occupy $\pi r^2$ units of substrate area, but create $2\pi r^2$ units of surface area for settlement. Second, when a competitor dies, we assume that it falls from the substrate, taking any polyps on its surface with it. The term $a_2 \psi x y_2$ represents the resulting rate of decrease of polyp numbers on potential competitors per unit substrate area. Note that in order to keep track of total polyp numbers, $y_2$ must be per unit substrate area, not per unit competitor surface area. This is because the loss of a competitor organism together with its associated polyps does not reduce the number of polyps per unit competitor surface area. The other parameters are the proportional rate of settlement of polyps on potential competitors ($c_0$, positive, numbers $\mathrm{L}^{-2}\mathrm{T}^{-1}$), the proportional rate of increase of polyp number on potential competitors by budding of polyps on potential competitors ($c_1$, positive, $\mathrm{T}^{-1}$), the proportional death rate of polyps on potential competitors ($c_2$, negative, $\mathrm{T}^{-1}$), and the proportional rate of increase of polyps on potential competitors due to budding from polyps on substrate ($c_3$, positive, $\mathrm{T}^{-1}$).

We determine the signs of the elements of the community matrix for this model below (section S10).

### S9 Dimensionless form and community matrix for basic dynamic model

Writing the basic model in an equivalent dimensionless form will reduce the number of parameters, and will also make it a more natural description of point-count data from experiments. The proportion of substrate filled by potential competitors, $x$, is already dimensionless. Let $y_1^* = \delta y_1$ be the proportion of substrate filled by polyps on the substrate. Let $t^* = t\delta b_0$ be a dimensionless time variable, scaled by the proportional rate at which polyps on substrate fill area by settlement. Substituting these dimensionless variables into Equations 3 and 4, we obtain the dimensionless system

$$\begin{aligned}\frac{\mathrm{d}x}{\mathrm{d}t^*} &= \Pi_1(1-x-y_1^*) + \Pi_2 x(1-x-y_1^*) + \Pi_3 x,\\ \frac{\mathrm{d}y_1^*}{\mathrm{d}t^*} &= (1-x-y_1^*) + \Pi_4 y_1^*(1-x-y_1^*) + \Pi_5 y_1^*,\end{aligned} \tag{S9}$$

with parameters $\Pi_1 = a_0/(\delta b_0)$ (positive, ratio of settlement rate by potential competitors to the proportional rate at which polyps on substrate fill area by settlement), $\Pi_2 = a_1/(\delta b_0)$ (positive, ratio of growth rate of potential competitors to the proportional rate at which polyps on substrate fill area by settlement), $\Pi_3 = a_2/(\delta b_0)$ (negative, ratio of death rate of potential competitors to the proportional rate at which polyps on substrate fill area by settlement), $\Pi_4 = b_1/(\delta b_0)$ (positive, ratio of polyp budding rate onto substrate by polyps on substrate to the proportional rate at which polyps on substrate fill area by settlement), $\Pi_5 = b_2/(\delta b_0)$ (negative, ratio of polyp death rate on substrate to the proportional rate at which polyps on substrate fill area by settlement).

We measure interaction strengths using the community matrix of partial derivatives of proportional rates of change with respect to the dimensionless state variables. This is an appropriate choice of interaction strength measurement for our experiment, because it does not require the assumption of equilibrium (Laska and Wootton, 1998). We include effects on settlement, because we want to measure the overall effects on proportional rates of change of relative abundances. However, if we wanted a measure of habitat quality alone, it would be more appropriate to exclude effects on settlement (Drake and Richards, 2018).

Let $f(x, y_1^*) = \dfrac{1}{x}\dfrac{\mathrm{d}x}{\mathrm{d}t^*}, g(x, y_1^*) = \dfrac{1}{y_1^*}\dfrac{\mathrm{d}y_1^*}{\mathrm{d}t^*}$ be the proportional rates of change of the dimensionless variables $x, y_1^*$. Then the community matrix $\mathbf{C}$ is

$$\begin{aligned}\mathbf{C} &= \begin{pmatrix} \dfrac{\partial f}{\partial x} & \dfrac{\partial f}{\partial y_1^*} \\ \dfrac{\partial g}{\partial x} & \dfrac{\partial g}{\partial y_1^*} \end{pmatrix}\\ &= \begin{pmatrix} -\Pi_1(1-y_1^*)/x^2 - \Pi_2 & -\Pi_1/x - \Pi_2 \\ -1/y_1^* - \Pi_4 & -(1-x)/(y_1^*)^2 - \Pi_4 \end{pmatrix}.\end{aligned}$$

Now because $0 \leq x \leq 1$, $0 \leq y_1^* \leq 1$, $0 \leq x + y_1^* \leq 1$, and $\Pi_3$ is the only negative parameter appearing in the community matrix, the signs of the elements in the matrix are

$$\begin{pmatrix} - & - \\ - & - \end{pmatrix}. \tag{S10}$$

Thus, each group of organisms in the model has overall negative intra-group density dependence (diagonal elements), and potential competitors and polyps on substrate have negative effects on each other (elements

(1, 2) and (2, 1)).

#### S10 Dimensionless form and community matrix for model including polyp growth on potential competitors

We take the same approach to nondimensionalizing the model with polyp growth on potential competitors (section S8) as for the model in the main text. Let $y_2^* = \delta y_2$ be the area filled by polyps on potential competitors per unit substrate area: this may be greater than 1, because $\psi \geq 1$. Substituting this and the other dimensionless variables from the main text into Equations S6, S7 and S8, we obtain the dimensionless system

$$\begin{aligned}
\frac{\mathrm{d}x}{\mathrm{d}t^*} &= \Pi_1(1 - x - y_1^*) + \Pi_2 x(1 - x - y_1^*) + \Pi_3 x, \\
\frac{\mathrm{d}y_1^*}{\mathrm{d}t^*} &= (1 - x - y_1^*) + \Pi_4 y_1^*(1 - x - y_1^*) + \Pi_5 y_1^* + \Pi_6 y_2^*(1 - x - y_1^*), \\
\frac{\mathrm{d}y_2^*}{\mathrm{d}t^*} &= \Pi_7(\psi x - y_2^*) + \Pi_8 y_2^*(\psi x - y_2^*) + \Pi_3 \psi x y_2^* + \Pi_9 y_2^* + \Pi_{10} y_1^*(\psi x - y_2^*),
\end{aligned} \tag{S11}$$

with parameters (in additionl to those in the main text) $\Pi_6 = b_3/(\delta b_0)$ (positive, ratio of polyp budding rate onto substrate by polyps on potential competitors to the proportional rate at which polyps on substrate fill area by settlement), $\Pi_7 = c_0/b_0$ (positive, ratio of settlement rate of polyps on potential competitors to settlement rate of polyps on substrate), $\Pi_8 = c_1/(\delta b_0)$ (positive, ratio of polyp budding rate onto substrate by polyps on potential competitors to the proportional rate at which polyps on substrate fill area by settlement), $\Pi_9 = c_2/(\delta b_0)$ (negative, ratio of polyp death rate on potential competitors to the proportional rate at which polyps on substrate fill area by settlement) and $\Pi_{10} = c_3/(\delta b_0)$ (positive, ratio of polyp budding rate onto potential competitors by polyps on substrate to the proportional rate at which polyps on substrate fill area by settlement).

We use the community matrix to measure interaction strengths, as for the model in the main text, additionally defining $h(x, y_1^*, y_2^*) = \dfrac{1}{y_2^*}\dfrac{\mathrm{d}y_2^*}{\mathrm{d}t^*}$ to be the proportional rate of change of the dimensionless variable $y_2^*$. Then the community matrix is

$$\begin{aligned}
\mathbf{C} &= \begin{pmatrix} \frac{\partial f}{\partial x} & \frac{\partial f}{\partial y_1^*} & \frac{\partial f}{\partial y_2^*} \\ \frac{\partial g}{\partial x} & \frac{\partial g}{\partial y_1^*} & \frac{\partial g}{\partial y_2^*} \\ \frac{\partial h}{\partial x} & \frac{\partial h}{\partial y_1^*} & \frac{\partial h}{\partial y_2^*} \end{pmatrix} \\
&= \begin{pmatrix} -\Pi_1(1 - y_1^*)/x^2 - \Pi_2 & -\Pi_1/x - \Pi_2 & 0 \\ -(1 + \Pi_6 y_2^*)/y_1^* - \Pi_4 & -(1 - x)(1 + \Pi_6 y_2^*)/(y_1^*)^2 - \Pi_4 & \Pi_6(1 - x - y_1^*)/y_1^* \\ \psi((\Pi_7 + \Pi_{10} y_1^*)/y_2^* + \Pi_3 + \Pi_8) & \Pi_{10}(\psi x - y_2^*)/y_2^* & -\psi x(\Pi_7 + \Pi_{10} y_1^*)/(y_2^*)^2 - \Pi_8 \end{pmatrix}.
\end{aligned}$$

As in the main text, $0 \leq x \leq 1$, $0 \leq y_1^* \leq 1$, $0 \leq y_2^*$, $0 \leq x + y_1^* \leq 1$, $0 \leq \psi x - y_2^* \leq 1$, $\psi \geq 1$, and $\Pi_3$ is the only negative parameter, so the signs of the elements in the matrix are

$$\begin{pmatrix} - & - & 0 \\ - & - & + \\ ? & + & - \end{pmatrix}.$$

In addition to the effects discussed in the main text, polyps on substrate and polyps on potential competitors have positive effects on each other (elements $(2, 3)$ and $(3, 2)$), and polyps on potential competitors have no direct effect on potential competitors (element $(1, 3)$). The sign of the effect of potential competitors on polyps growing on potential competitors (element $(3, 1)$) is unknown until parameter values are known. However, the sign does not depend on $\psi$, the surface area available for occupation by polyps per unit substrate area occupied by potential competitors, although the magnitude does. Furthermore, the sign is more likely to be negative if $y_2^*$, the area filled by polyps on potential competitors per unit substrate area, is large, the ratio $\Pi_7$ of settlement rates of polyps on potential competitors to substrate is small, the ratio $\Pi_{10}$ of polyp budding rate onto potential competitors by polyps on substrate the the proportional rate at which polyps on substrate fill area by settlement is small, the proportion $y_1^*$ of substrate filled by polyps is small, the ratio $\Pi_3$ of the (negative) death rate of potential competitors to the proportional rate at which polyps on substrate fill area by settlement is large, and the ratio $\Pi_8$ of polyp budding rate onto substrate by polyps on potential competitors to the proportional rate at which polyps on substrate fill area by settlement is small. In other words, low input rates of polyps on potential competitors and a high death rate of potential competitors (both scaled relative to the rate of settlement on substrate), or a high density of polyps on potential competitors, are likely to result in a negative effect of potential competitors on polyps growing on potential competitors.

### S11 Community matrices for models with positive effects of polyps on potential competitors

Here, we write models for community dynamics in which polyps may have positive effects on potential competitors in dimensionless form, and give the community matrices for each. For reference, the basic model is

$$\begin{aligned} \frac{\mathrm{d}x}{\mathrm{d}t^*} &= \Pi_1(1 - x - y_1^*) + \Pi_2 x(1 - x - y_1^*) + \Pi_3 x, \\ \frac{\mathrm{d}y_1^*}{\mathrm{d}t^*} &= (1 - x - y_1^*) + \Pi_4 y_1^*(1 - x - y_1^*) + \Pi_5 y_1^*, \end{aligned} \tag{S12}$$

Dimensionless parameters that are not new are as in Equation S12, and community matrices are obtained as in the main text. In all cases, the signs of the elements in the community matrix are

$$\begin{pmatrix} - & ? \\ - & - \end{pmatrix},$$

with the effect of polyps on potential competitors positive for some but not all compositions and parameter values. We give the condition for $c_{1,2} > 0$ in each model.

#### S11.1 Facilitation of settlement

For reference, the model with facilitation of settlement is

$$\frac{\mathrm{d}x}{\mathrm{d}t} = (a_0 + m_0\delta y_1)(1 - x - \delta y_1) + a_1 x(1 - x - \delta y_1) + a_2 x. \tag{S13}$$

The dimensionless form of Equation S13 is

$$\frac{\mathrm{d}x}{\mathrm{d}t^*} = (\Pi_1 + \Pi_{11} y_1^*)(1 - x - y_1^*) + \Pi_2 x(1 - x - y_1^*) + \Pi_3 x,$$

where $\Pi_{11} = m_0/(\delta b_0) > 0$. The community matrix is

$$\mathbf{C} = \begin{pmatrix} -(\Pi_1 + \Pi_{11} y_1^*)(1 - y_1^*)/x^2 - \Pi_2 & -\Pi_1/x - \Pi_2 + (\Pi_{11}/x)(1 - x - 2y_1^*) \\ -1/y_1^* - \Pi_4 & -(1 - x)/(y_1^*)^2 - \Pi_4 \end{pmatrix}$$

with $c_{1,2} > 0$ if $\Pi_{11}(1 - x - 2y_1^*) > \Pi_1 + \Pi_2 x$.

#### S11.2 Facilitation of growth

For reference, the model for facilitation of growth is

$$\frac{\mathrm{d}x}{\mathrm{d}t} = a_0(1 - x - \delta y_1) + (a_1 + m_1\delta y_1)x(1 - x - \delta y_1) + a_2 x. \tag{S14}$$

The dimensionless form of Equation S14 is

$$\frac{\mathrm{d}x}{\mathrm{d}t^*} = \Pi_1(1 - x - y_1^*) + (\Pi_2 + \Pi_{12} y_1^*)x(1 - x - y_1^*) + \Pi_3 x,$$

where $\Pi_{12} = m_1/(\delta b_0) > 0$. The community matrix is

$$\mathbf{C} = \begin{pmatrix} -\Pi_1(1 - y_1^*)/x^2 - \Pi_2 - \Pi_{12} y_1^* & -\Pi_1/x - \Pi_2 + \Pi_{12}(1 - x) - 2\Pi_{12} y_1^* \\ -1/y_1^* - \Pi_4 & -(1 - x)/(y_1^*)^2 - \Pi_4 \end{pmatrix}$$

with $c_{1,2} > 0$ if $\Pi_{12}(1 - x - 2y_1^*) > \Pi_1/x + \Pi_2$.

##### S11.3 Overgrowth of polyps by potential competitors

For reference, the model for overgrowth of polyps by potential competitors is

$$\frac{\mathrm{d}x}{\mathrm{d}t} = a_0\left(1 - x - \delta y_1\right) + a_1 x\left(1 - x - \delta y_1\right) + a_{1,y_1} x y_1 + a_2 x, \tag{S15}$$

$$\frac{\mathrm{d}y_1}{\mathrm{d}t} = b_0\left(1 - x - \delta y_1\right) + b_1 y_1\left(1 - x - \delta y_1\right) - \frac{a_{1,y_1}}{\delta} x y_1 + b_2 y_1. \tag{S16}$$

The dimensionless form of Equations S15 and S16 is

$$\begin{aligned}\frac{\mathrm{d}x}{\mathrm{d}t^*} &= \Pi_1\left(1 - x - y_1^*\right) + \Pi_2 x\left(1 - x - y_1^*\right) + \Pi_{13} x y_1^* + \Pi_3 x,\\ \frac{\mathrm{d}y_1^*}{\mathrm{d}t^*} &= \left(1 - x - y_1^*\right) + \Pi_4 y_1^*\left(1 - x - y_1^*\right) - \Pi_{13} x y_1^* + \Pi_5 y_1^*,\end{aligned}$$

where $\Pi_{13} = a_{1,y_1^*}/(\delta b_0) > 0$. The community matrix is

$$\mathbf{C} = \begin{pmatrix} -\Pi_1(1-y_1^*)/x^2 - \Pi_2 & -\Pi_1/x - \Pi_2 + \Pi_{13} \\ -1/y_1^* - \Pi_4 - \Pi_{13} & -(1-x)/(y_1^*)^2 - \Pi_4 \end{pmatrix}$$

with $c_{1,2} > 0$ if $\Pi_{13} > \Pi_1/x + \Pi_2$.

##### S11.4 Protection from predators

For reference, the model for protection from predators is

$$\frac{\mathrm{d}x}{\mathrm{d}t} = a_0\left(1 - x - \delta y_1\right) + a_1 x\left(1 - x - \delta y_1\right) + a_2 e^{-m_2 \delta y_1} x. \tag{S17}$$

The dimensionless form of Equation S17 is

$$\frac{\mathrm{d}x}{\mathrm{d}t^*} = \Pi_1\left(1 - x - y_1^*\right) + \Pi_2 x\left(1 - x - y_1^*\right) + \Pi_3 e^{-m_2 y_1^*} x,$$

where $m_2$ is already dimensionless. The community matrix is

$$\mathbf{C} = \begin{pmatrix} -\Pi_1(1-y_1^*)/x^2 - \Pi_2 & -\Pi_1/x - \Pi_2 - m_2\Pi_3 e^{-m_2 y_1^*} \\ -1/y_1^* - \Pi_4 & -(1-x)/(y_1^*)^2 - \Pi_4 \end{pmatrix},$$

with $c_{1,2} > 0$ if $-m_2\Pi_3 e^{-m_2 y_1^*} > \Pi_1/x + \Pi_2$ (note that $\Pi_3 < 0$).

#### S12 Fitting dynamic models to experimental data

We fitted versions of Equations 3 and 4, with each of the modifications in section 2.3.2 in turn, to the experimental data from all weeks and panels. As explained below, our underlying model for dynamics is deterministic, so we will only obtain a caricature of the true dynamics. Nevertheless, this will give a qualitative understanding of the interactions in the community. We wrote Equations 3 and 4 in terms of proportions of space occupied, $x$ and $y_1^*$, but retained time in dimensioned form (measured in weeks since panels were put in the water). Thus the basic model is:

$$\frac{\mathrm{d}x}{\mathrm{d}t} = a_0 (1 - x - y_1^*) + a_1 x (1 - x - y_1^*) + a_2 x, \tag{S18}$$

$$\frac{\mathrm{d}y_1^*}{\mathrm{d}t} = \delta b_0 (1 - x - y_1^*) + b_1 y_1^* (1 - x - y_1^*) + b_2 y_1^*. \tag{S19}$$

In the version with overgrowth of polyps by potential competitors, the overgrowth effect is then $+a_{1,y_1^*} x y_1^*$ in $\frac{\mathrm{d}x}{\mathrm{d}t}$, and $-a_{1,y_1^*} x y_1^*$ in $\frac{\mathrm{d}y_1^*}{\mathrm{d}t}$, where $a_{1,y_1^*} = a_{1,y_1}/\delta$.

Let $\mathbf{y}_{jkl,t}^{(3)}$ be the point counts of *A. aurita* on panel, bare panel and potential competitors (*Botrylloides spp.*, *Bugula spp* and *M. tubifera*, as in the models for final composition, but amalgamated into a single part) on the panel from depth $j$, treatment $k$, block $l$ at time $t$, pre-treatment in those cases where a treatment was applied. We model these counts using:

$$\mathbf{y}_{jkl,t}^{(3)} \sim \text{multinomial}(n_{jkl,t}^{(3)}, \boldsymbol{\rho}_{jkl,t}^{(3)}), \tag{S20}$$

$$\boldsymbol{\rho}_{jkl,t}^{(3)} = \begin{pmatrix} y_{1,jkl}^*(t) \\ 1 - x_{jkl}(t) - y_{1,jkl}^*(t) \\ x_{jkl}(t) \end{pmatrix}, \tag{S21}$$

where $x_{jkl}(t)$ and $y_{1,jkl}^*(t)$ are the solutions to Equations S18 and S19 (or modified versions as in section 2.3.2) for the panel with depth $j$, treatment $k$ in block $l$, obtained using a fourth and fifth order Runge-Kutta method, with initial conditions $x_{jkl}(0) = 0, y_{1,jkl}^*(0) = 0$ (the empty panel). We estimated all the parameters (other than removal effects, as described below) separately at depths 1 m and 3 m.

We did not include block and panel effects. These should really be the outcome of block- and panel-specific temporal variation in parameters such as settlement and growth rates in a stochastic differential equation version of Equations S18 and S19. We did not attempt to fit such a model because it presents substantial technical difficulties. First, each parameter has a fixed sign (for example, settlement rates $a_0$ and $\delta b_0$ must always be positive). Care must be taken to respect such conditions when specifying a stochastic differential equation. The usual stochastic differential equation models for community dynamics avoid the problem by allowing only proportional population growth rates at low density (which do not have a sign constraint) to be stochastic (Kloeden and Platen, 1999, p. 254). Second, we cannot obtain an explicit solution for a stochastic differential equation version of Equations S18 and S19, and would therefore have to rely on numerical methods such as the Euler-Maruyama algorithm. However, if the time steps in such a method are small, standard Markov chain methods for Bayesian estimation can converge arbitrarily slowly, and although alternative algorithms exist (Fuchs, 2013, section 7.3), they cannot be implemented in Stan, and dealing with the experimental design in existing software designed for estimating parameters of stochastic differential equations appears difficult. With large time steps, estimation bias may be substantial, and there is a risk that the estimated composition might fall outside the simplex. These problems are not insoluble, but are outside the scope of this paper.

Where a treatment was applied, we model the post-treatment counts $\mathbf{y}^{(3)}_{jkl,t,\text{post}}$ as

$$\mathbf{y}^{(3)}_{jkt,\text{post}} \sim \text{multinomial}(n^{(3)}_{jkt,\text{post}}, \boldsymbol{\rho}^{(3)}_{jkt,\text{post}}), \tag{S22}$$

$$\boldsymbol{\rho}^{(3)}_{jkt,\text{post}} = \begin{pmatrix} (1 - r_A \mathbf{1}_{A,jkt}) y_1^*(t) \\ 1 - (1 - r_O \mathbf{1}_{O,jkt}) x(t) - (1 - r_A \mathbf{1}_{A,jkt}) y_1^*(t) \\ (1 - r_O \mathbf{1}_{O,jkt}) x(t) \end{pmatrix}, \tag{S23}$$

where $\mathbf{1}_{A,jkt}$ and $\mathbf{1}_{O,jkt}$ are indicator variables for the application of treatments $A$ and $O$ respectively on the panel from treatment $k$, depth $j$, time $t$, and $r_A$ and $r_O$ are the proportions of *A. aurita* and potential competitors removed in the $A$ and $O$ treatments respectively (both intended to be 1/2, and the same at both depths). If a treatment was applied, we used the post-treatment state to initialize the differential equation solver for the next time interval.

We fitted each version of the dynamic model using `cmdstan 2.25.0` (Hoffman and Gelman, 2014; Carpenter et al., 2017). Priors are described in the supporting information, section S13. Fitting, checking, calibration and model comparison using approximate leave-one-out cross-validation are described in the supporting information, section S14.

### S13 Priors for models of community dynamics

In models for community dynamics, all parameters other than $r_A$ and $r_O$ were estimated separately at each depth. Priors were generally based on some biological knowledge from previous experiments. We used the same priors at both depths because we did not have strong prior knowledge of how each parameter might depend on depth.

The proportional population growth rate of *A. aurita* polyps at low density in the absence of settlement is $b_1 - b_2$. From a lab experiment conducted in winter, with ample food and a low density of polyps, this was estimated as $0.1\,\text{week}^{-1}$, with a standard deviation of $0.06\,\text{week}^{-1}$ (data from 9 tanks at ambient temperature, Goggins, 2018). Previous experiments with similar settlement panels at the same site suggest that few *A. aurita* polyps will be visible until panels have been in the water for at least 2 weeks, and that *A. aurita* polyps might cover about 0.05 of the available space after 5 weeks (Maxatova, 2016). A positive half-normal $N(0, 0.02)$ prior for $\delta b_0$ (these two parameters only appear as a product in this form of the model), together with positive half-normal $N(0, 0, 0.2)$ for $b_1$ and negative half-normal $N(0, 0.05)$ for $b_2$ gives a distribution for $b_1 - b_2$ with mean 0.1, standard deviation 0.1, and a distribution of trajectories that is consistent with previous experiments (Figure S4a).

Settlement and growth rates of potential competitors may be somewhat higher than those of *A. aurita.* In a previous experiment, potential competitors filled a mean of 0.06 of the available space after 1 week, and mean 0.24, range 0.04 to 0.97 after 5 weeks (Maxatova, 2016). A positive half-normal $N(0, 0.05)$ prior for $a_0$, positive half-normal $N(0, 1)$ for $a_1$ and negative half-normal $N(0, 0.05)$ for $a_2$ gives mean 0.06 after 1 week, and proportional cover over almost the entire range from 0 to 1 (with mean 0.42) after 5 weeks (Figure S4c).

In the removal treatments, we aimed to remove half of the target taxon. However, this was done by eye, so the actual proportion removed may differ. We treated $r_A$ and $r_O$ as parameters to be estimated, with beta$(2, 2)$ priors, which are moderately concentrated around $1/2$. We did not allow $r_A$ and $r_O$ to differ between depths, because we have no reason to believe that the proportions removed will depend on depth.

In the settlement facilitation model, it seems plausible that facilitation might double the settlement rate of potential competitors when polyps are very abundant (i.e. $y_1^*$ close to 1), so $m_0$ could be of similar size to $a_0$. We therefore chose a positive half-normal $N(0, 0.05)$ prior for $m_0$. In the growth facilitation model, it seems plausible that facilitation might double the proportional growth rate of potential competitors when polyps are very abundant, so $m_1$ could be of a similar size to $a_1$. We therefore chose a positive half-normal $N(0, 1)$ prior for $m_1$. In the overgrowth model, it seems plausible that the rate at which potential competitors overgrow space occupied by polyps might be similar to the rate at which they grow into empty space, so $a_{1,y_1^*}$ could be of a similar size to $a_1$. We therefore chose a positive half-normal $N(0, 1)$ prior for $a_{1,y_1^*}$. In the protection model, it seems plausible that protection might halve the proportional death rate of potential competitors when polyps are very abundant, so that $e^{-m_2} = 1/2$ should be a plausible value. This corresponds to $m_2 = -\log(1/2)$. We therefore chose a positive half-normal $N(0, -(1/2)\log(1/2))$ prior for $m_2$.

### S14 Fitting, checking, calibration and model comparison for models of community dynamics

We fitted each version of the dynamic model using `cmdstan 2.25.0` (Hoffman and Gelman, 2014; Carpenter et al., 2017). We ran 4 chains for 2500 warmup and 2500 samping iterations. This took up to 7546 s on a 64-bit Ubuntu 18.04 system wih 4 Intel Xeon 3.2 GHz cores and 16 GiB RAM. Effective sample sizes were always at least 1572, potential scale reduction factors were always no greater than 1.0026, and inspection of trace plots did not reveal any evidence of failures to converge. Processing and visualization of samples from the posterior was done in R version 4.0.3 (R Core Team, 2020), with the packages `rstan 2.21.2` (Stan Development Team, 2020) and `compositions 2.0-0` (van den Boogaart et al., 2020).

We compared models using approximate leave-one-out cross-validation (Vehtari et al., 2017). Because we did not include block or panel effects, this could be done directly using Pareto-smoothed importance sampling via the function `loo_compare()` in the R package `loo 2.4.1` (Vehtari et al., 2020).

We carried out a graphical posterior predictive check for the best-fitting version of the dynamic model. We generated 100 simulated data sets drawn from the multinomial distribution for each time point on each panel, given by Equations S20 and S21 in the main text, plotted the time series of simulated observations, and visually compared with the time series of real observations.

We checked for gross errors in coding and obtained a rough estimate of performance of the estimation method by fitting the best-performing model to 10 data sets simulated under this best-performing model, with the same structure (number of replicates, pattern of sampling and treatment applications, multinomial sample sizes) as the real data. Simulated counts were generated from Equations S20, S21 S22 and S23 in the main text. The model was fitted to each simulated data set in exactly the same way as for the real data set. This took approximately 10.5 h on a 64-bit Ubuntu 18.04 system wih 4 Intel Xeon 3.2 GHz cores and 16 GiB RAM. We plotted the true parameter values, prior densities, and posterior densities from each simulated data set, and recorded the number of simulated data sets for which the estimated 95% HPD interval included the true parameter value. Unlike simulation-based calibration, this approach cannot tell us whether the method is correctly sampling from the posterior distribution for each parameter, because the true posterior densities are unknown. However, simulation-based calibration would have been too time-consuming for this model. Nevertheless, gross errors may be revealed by implausible posterior densities. If the proportion of simulated data sets for which the 95% HPD interval includes the true parameter value is low, possible interpretations include biased sampling from the posterior or a strong influence of the prior.

Table S1: Manova parameter estimates for the selected model (with no interaction): overall mean $\boldsymbol{\mu}$, depth effect $\boldsymbol{\alpha}$, removal treatment effects $\boldsymbol{\beta}_1$, $\boldsymbol{\beta}_2$, rows of lower triangle of panel covariance matrix $\boldsymbol{\Sigma}$, rows of lower triangle of block covariance matrix $\mathbf{Z}$. Columns are ilr coordinates. Each cell contains the posterior mean, with marginal 95 % credible highest density regions in parentheses. For some elements of $\mathbf{Z}$, the highest density region consists of multiple disjoint intervals.

| | 1 | 2 | 3 | 4 |
|---|---|---|---|---|
| $\boldsymbol{\mu}$ | -2.00 (-2.27, -1.72) | -1.18 (-1.64, -0.71) | 0.36 (0.00, 0.72) | -1.33 (-1.68, -0.99) |
| $\boldsymbol{\alpha}$ | 0.22 (0.08, 0.36) | -1.59 (-1.85, -1.34) | 0.31 (0.07, 0.54) | 0.29 (0.02, 0.55) |
| $\boldsymbol{\beta}_1$ | 0.29 (0.13, 0.45) | -1.33 (-1.64, -1.03) | -0.16 (-0.44, 0.13) | -0.02 (-0.36, 0.31) |
| $\boldsymbol{\beta}_2$ | 0.07 (-0.12, 0.27) | 1.08 (0.74, 1.41) | 0.21 (-0.09, 0.51) | -0.14 (-0.49, 0.20) |
| $\boldsymbol{\sigma}_1$ | 0.10 (0.00, 0.21) | | | |
| $\boldsymbol{\sigma}_2$ | -0.02 (-0.16, 0.10) | 0.64 (0.23, 1.04) | | |
| $\boldsymbol{\sigma}_3$ | 0.10 (-0.01, 0.22) | -0.08 (-0.31, 0.15) | 0.43 (0.11, 0.76) | |
| $\boldsymbol{\sigma}_4$ | 0.02 (-0.10, 0.15) | 0.25 (-0.03, 0.53) | 0.10 (-0.14, 0.35) | 0.66 (0.26, 1.08) |
| $\boldsymbol{\zeta}_1$ | 0.13 (-0.02, 0.36) | | | |
| $\boldsymbol{\zeta}_2$ | -0.03 (-0.20, 0.12) | 0.36 (-0.18, 0.96) | | |
| $\boldsymbol{\zeta}_3$ | 0.04 (-0.05, 0.17) | 0.01 (-0.15, 0.19) | 0.14 (-0.02, 0.46) | |
| $\boldsymbol{\zeta}_4$ | -0.01 (-0.10, 0.07) | 0.03 (-0.12, 0.20) | 0.00 (-0.09, 0.10) | 0.10 (-0.01, 0.35) |

Table S2: Parameter estimates for ordinary differential equation models. Columns are models. Each cell contains the posterior mean, with marginal 95 % credible highest density regions in parentheses. For some parameters, the highest density region consists of multiple disjoint intervals. Where a boundary of the highest density region has a different sign from the posterior mean, this is an artefact of smoothing in density estimation, and all sample values had the same sign. All parameters other than $r_A$ and $r_O$ had separate estimated values at 1 m and 3 m. Spaces indicate parameters that were not included in a given model.

| | basic | settlement facilitation | growth facilitation | overgrowth | protection |
|---|---|---|---|---|---|
| $a_0$, 1 m | 0.014 (0.012, 0.015) | 0.014 (0.012, 0.015) | 0.014 (0.012, 0.015) | 0.013 (0.012, 0.015) | 0.014 (0.012, 0.015) |
| $a_0$, 3 m | 0.0036 (0.003, 0.0041) | 0.0033 (0.0026, 0.004) | 0.0036 (0.0031, 0.0041) | 0.0036 (0.0031, 0.0042) | 0.0036 (0.0031, 0.0041) |
| $a_1$, 1 m | 1.4 (1.3, 1.4) | 1.4 (1.3, 1.4) | 1.4 (1.3, 1.4) | 1.4 (1.3, 1.5) | 1.4 (1.3, 1.4) |
| $a_1$, 3 m | 0.59 (0.49, 0.68) | 0.58 (0.49, 0.68) | 0.56 (0.45, 0.67) | 0.42 (0.3, 0.54) | 0.59 (0.49, 0.69) |
| $a_2$, 1 m | -0.39 (-0.42, -0.35) | -0.39 (-0.42, -0.35) | -0.39 (-0.43, -0.35) | -0.4 (-0.44, -0.36) | -0.38 (-0.42, -0.35) |
| $a_2$, 3 m | -0.067 (-0.14, 0.0091) | -0.067 (-0.14, 0.0055) | -0.072 (-0.14, 0.00042) | -0.087 (-0.17, -0.0048) | -0.068 (-0.14, 0.00069) |
| $\delta b_0$, 1 m | 0.0009 (0.00067, 0.0011) | 0.00091 (0.00068, 0.0012) (0.0012, 0.0012) | 0.00092 (0.00068, 0.0012) | 0.0009 (0.00066, 0.00066) (0.00067, 0.0012) | 0.00091 (0.00067, 0.00067) (0.00068, 0.0012) |
| $\delta b_0$, 3 m | 0.049 (0.044, 0.053) | 0.049 (0.045, 0.053) | 0.049 (0.045, 0.054) | 0.046 (0.042, 0.051) | 0.049 (0.045, 0.053) |
| $b_1$, 1 m | 0.69 (0.58, 0.79) | 0.68 (0.57, 0.79) | 0.68 (0.57, 0.79) | 0.69 (0.6, 0.77) | 0.68 (0.57, 0.79) |
| $b_1$, 3 m | 0.012 (-0.0006, 0.032) (0.032, 0.035) | 0.012 (-0.00086, 0.035) (0.036, 0.037) | 0.012 (-0.00091, 0.035) | 0.014 (-0.0011, 0.04) (0.04, 0.041) | 0.012 (-0.00079, 0.034) |
| $b_2$, 1 m | -0.081 (-0.15, -0.008) | -0.081 (-0.15, -0.0079) | -0.079 (-0.15, -0.0066) | -0.035 (-0.088, 0.0017) | -0.079 (-0.15, -0.0077) |
| $b_2$, 3 m | -0.34 (-0.39, -0.3) | -0.35 (-0.4, -0.3) | -0.35 (-0.4, -0.3) | -0.26 (-0.31, -0.21) | -0.35 (-0.39, -0.3) |
| $r_A$ | 0.28 (0.25, 0.32) | 0.28 (0.25, 0.32) | 0.29 (0.25, 0.32) | 0.2 (0.16, 0.25) | 0.29 (0.25, 0.32) |
| $r_O$ | 0.41 (0.4, 0.43) | 0.41 (0.4, 0.43) | 0.41 (0.4, 0.43) | 0.42 (0.41, 0.43) | 0.41 (0.4, 0.43) |
| $m_0$, 1 m | | 0.032 (-0.0023, 0.084) | | | |
| $m_0$, 3 m | | 0.0062 (-0.00026, 0.017) | | | |
| $m_1$, 1 m | | | 0.21 (-0.021, 0.61) | | |
| $m_1$, 3 m | | | 0.49 (-0.018, 0.98) | | |
| $a_{1,y_1^*}$, 1 m | | | | 0.34 (0.17, 0.51) | |
| $a_{1,y_1^*}$, 3 m | | | | 2.2 (1.6, 2.8) | |
| $m_2$, 1 m | | | | | 0.25 (-0.02, 0.63) |
| $m_2$, 3 m | | | | | 0.28 (-0.021, 0.7) |

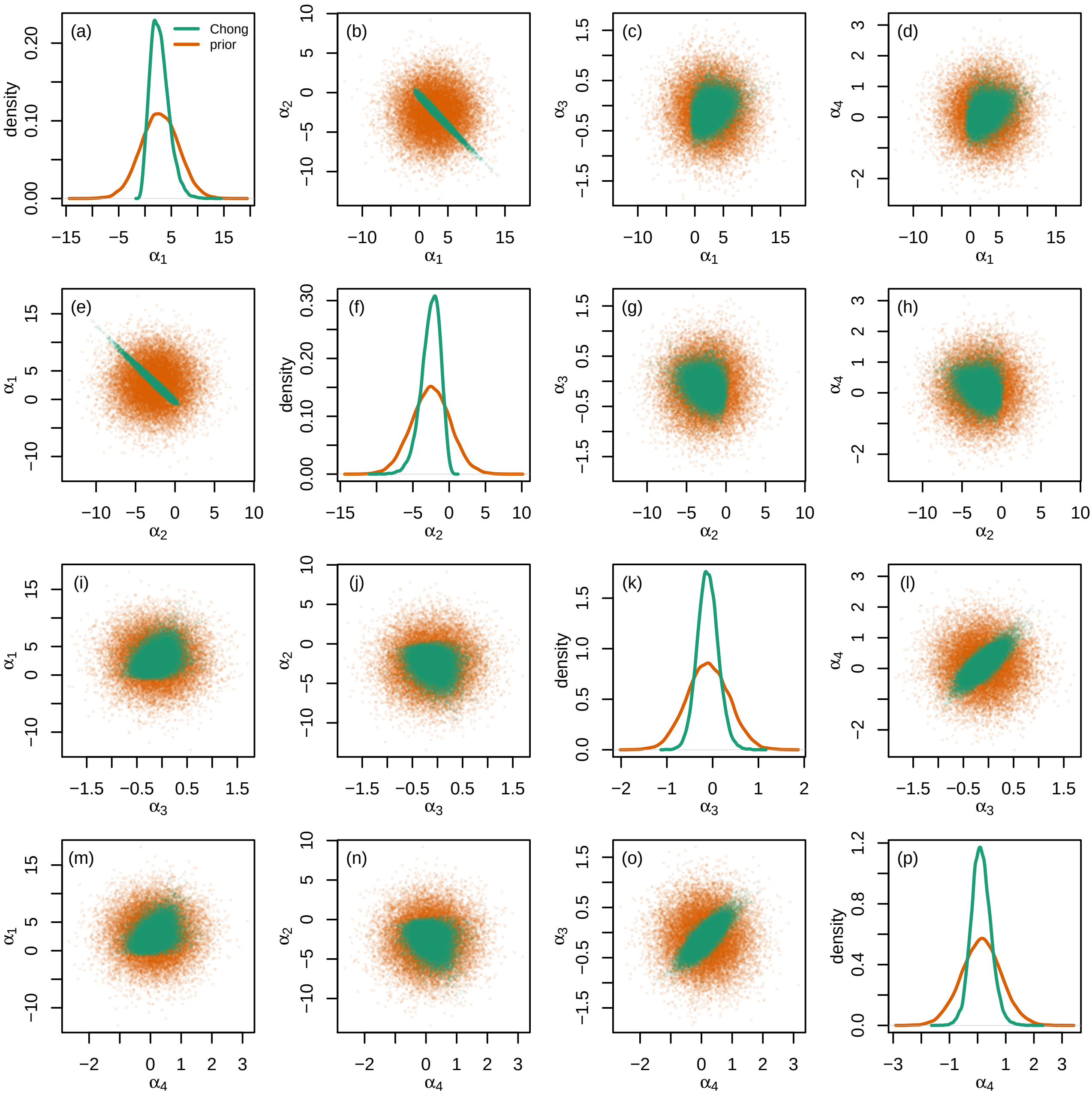


Figure S1: Prior for the depth effect on final subcomposition in ilr coordinates, $\boldsymbol{\alpha}$ (orange), and the posterior distribution of the difference in subcomposition between dock wall communities at 3 m and 1 m from Chong and Spencer (2018), on which it was based (green). Diagonal panels are kernel density estimates for individual components, and off-diagonal panels are pairwise scatter plots of pairs of components. Both based on samples of size 20000.

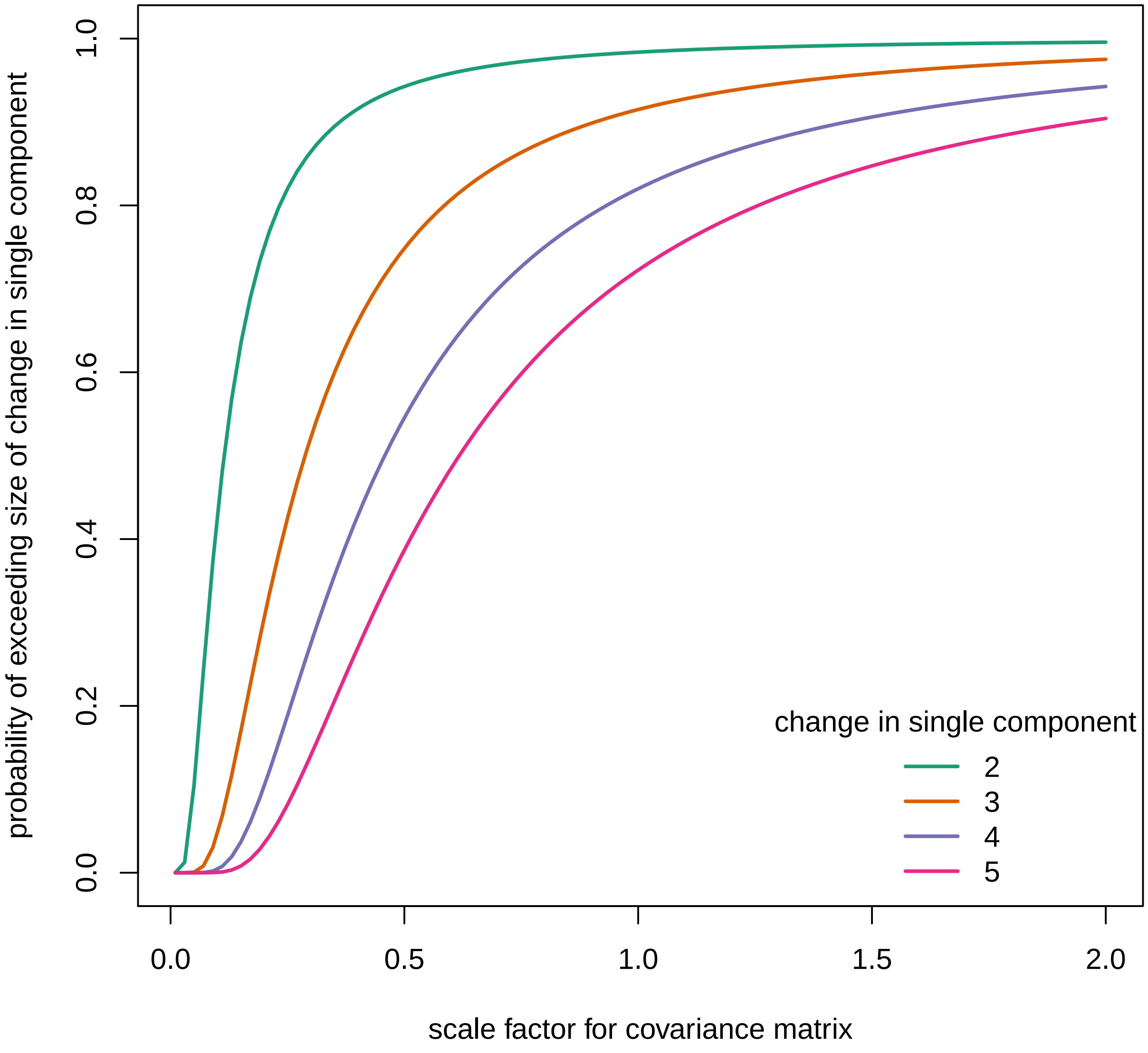


Figure S2: Choice of scale factor $c$ for covariance matrix $c\mathbf{I}_4$ of scraping treatment effects, scraping treatment by depth interaction and block effects. Each line represents a change corresponding to multiplying or dividing a single component by 2 (green), 3 (orange), 4 (purple) or 5 (pink). The $y$-axis is the probability of drawing a random vector at least as far from the origin as this, for scale factor $c$ on the $x$-axis.

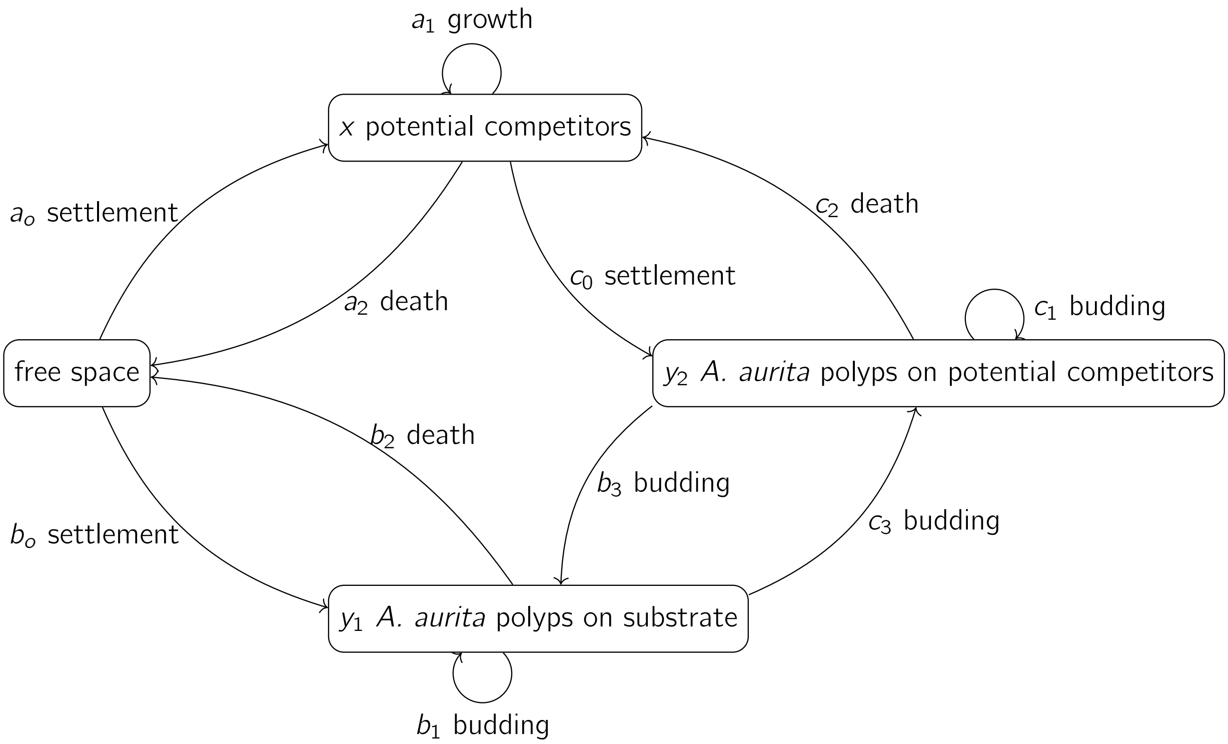


Figure S3: A model for the dynamics of polyps and potential competitors, including polyps growing on potential competitors, as in Equations S6, S7 and S8.

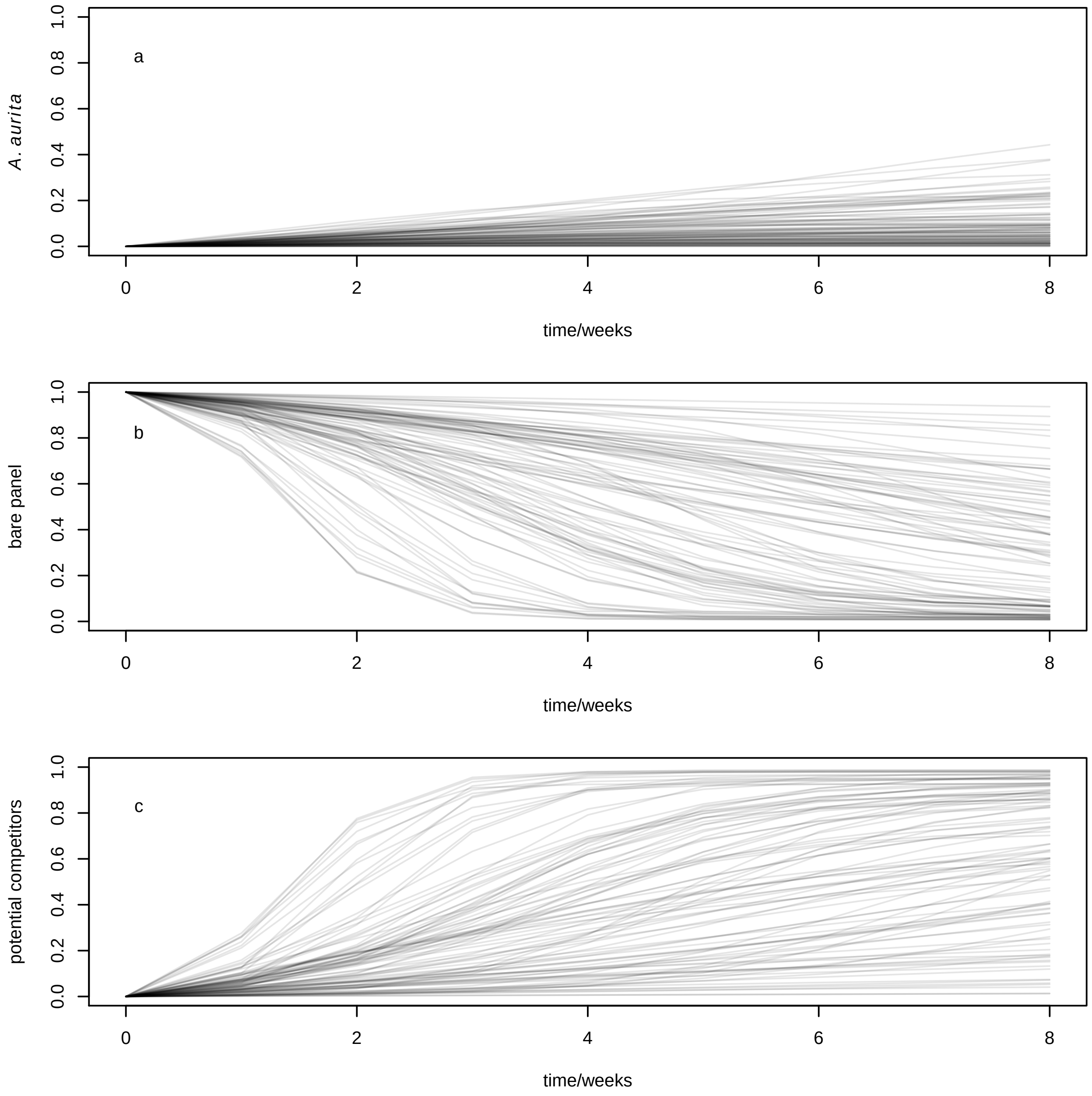


Figure S4: Prior distribution for time series of proportions of space filled by (a) *A. aurita*, (b) bare panel and (c) potential competitors, from the model specified by Equations S12 (with time redimensionalized by multiplying by the polyp settlement rate $\delta b_0$). Each of the 100 lines on each panel represents a simulated time series for a single draw from the priors.

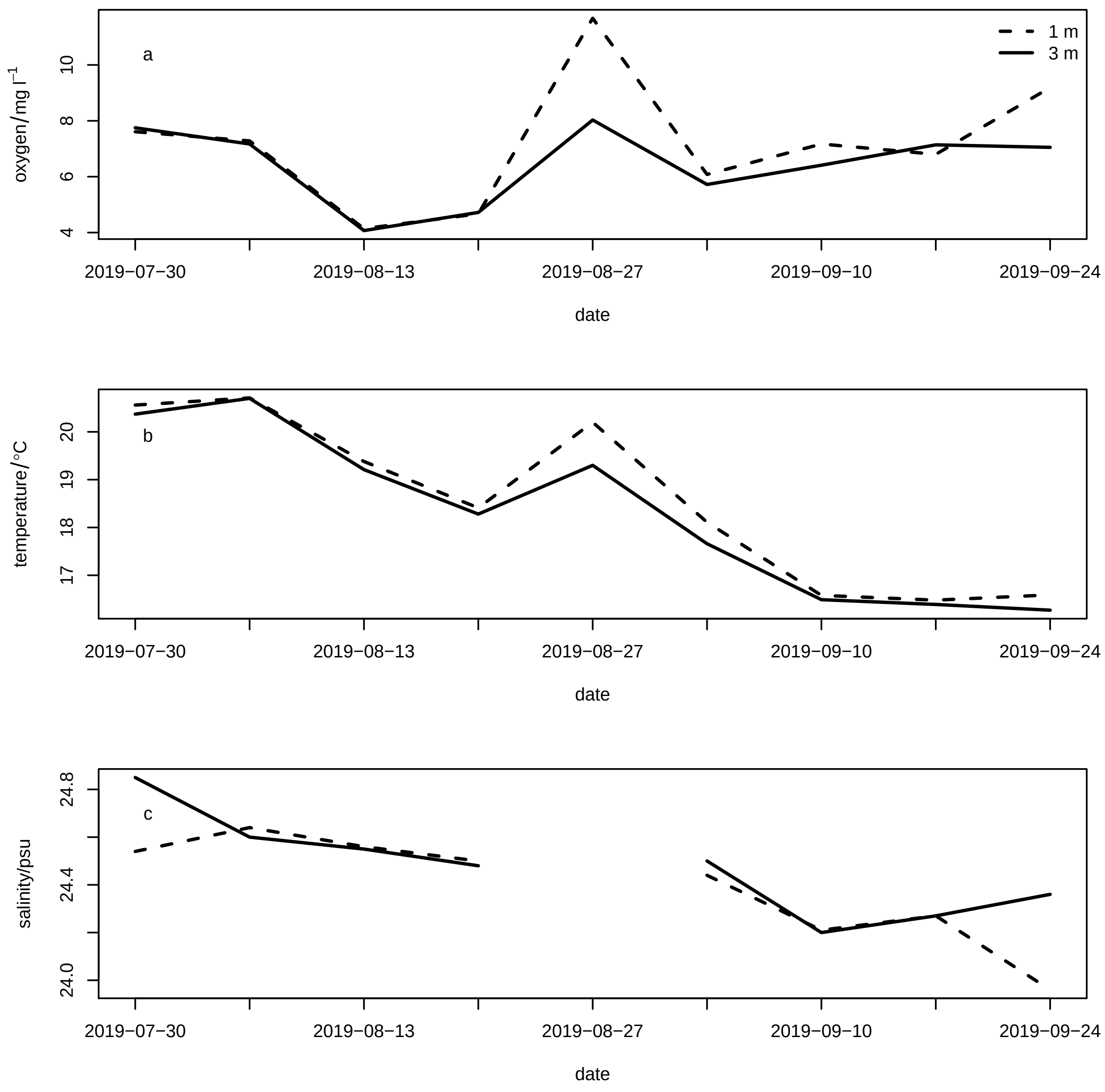


Figure S5: Dissolved oxygen (a, mg l$^{-1}$), temperature (b, °C) and salinity (c, psu) at 1 m (dashed lines) and 3 m (solid lines) over the course of the experiment. Ther were no salinity data on 2019-08-27.

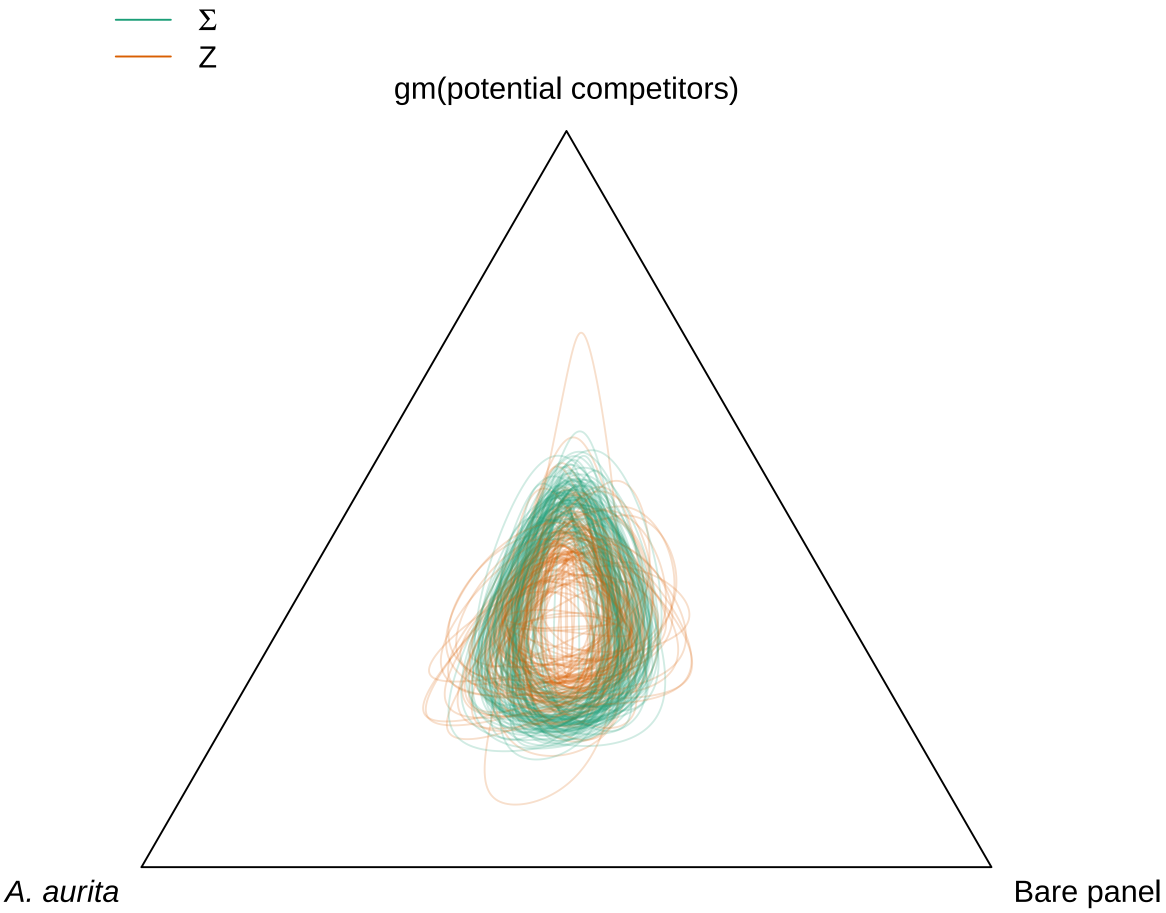


Figure S6: Visualization of panel ($\mathbf{\Sigma}$, green) and block ($\mathbf{Z}$, orange) covariance submatrices. Ellipses of unit Mahalanobis distance from the origin in the submatrix corresponding to the first two ilr coordinates, back-transformed into the 2-simplex with parts representing *A. aurita*, bare panel and gm(potential competitors). Samples of 100 ellipses from posterior distributions.

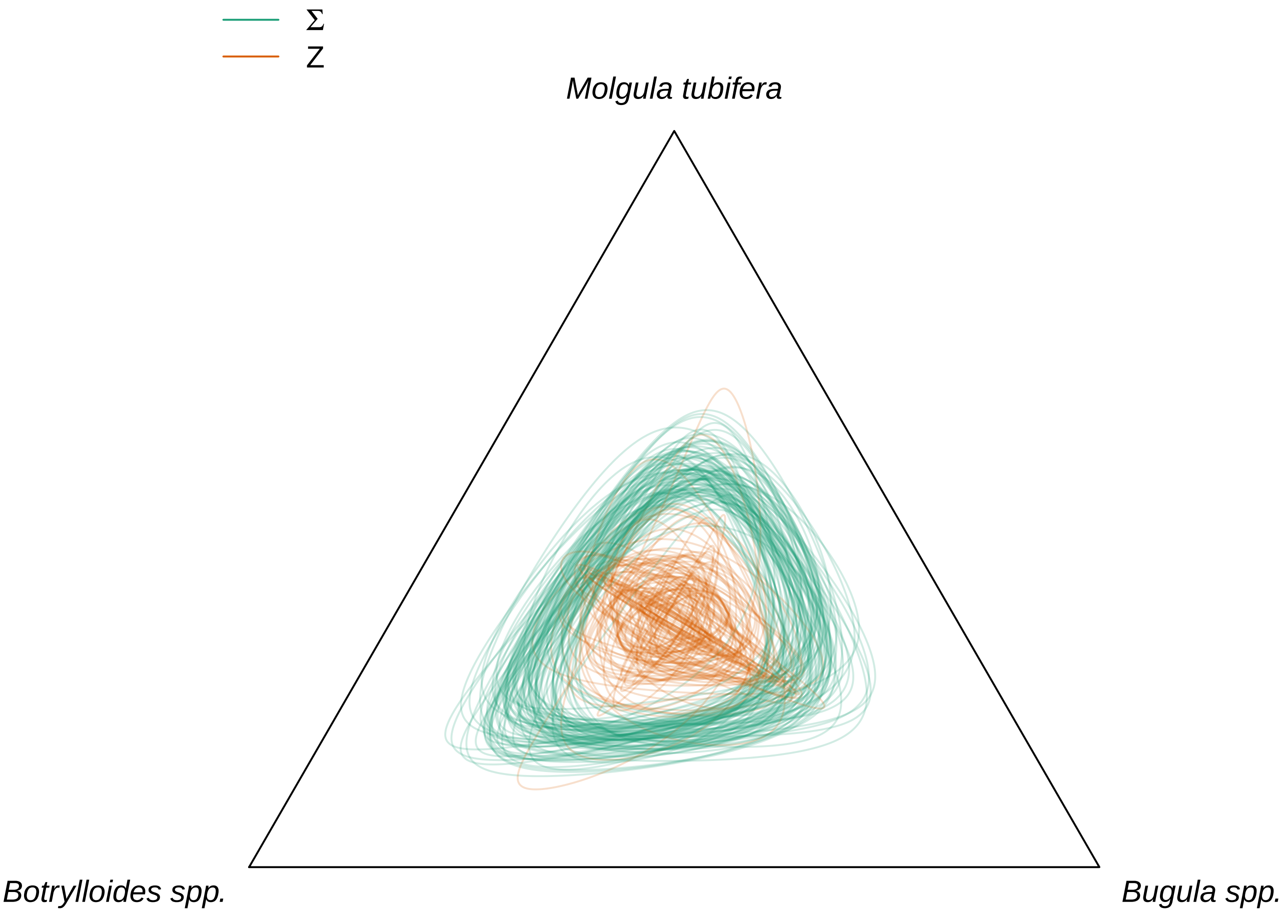


Figure S7: Visualization of panel ($\mathbf{\Sigma}$, green) and block ($\mathbf{Z}$, orange) covariance submatrices. Ellipses of unit Mahalanobis distance from the origin in the submatrix corresponding to the third and fourth ilr coordinates, back-transformed into the 2-simplex with parts representing potential competitors. Samples of 100 ellipses from posterior distributions.

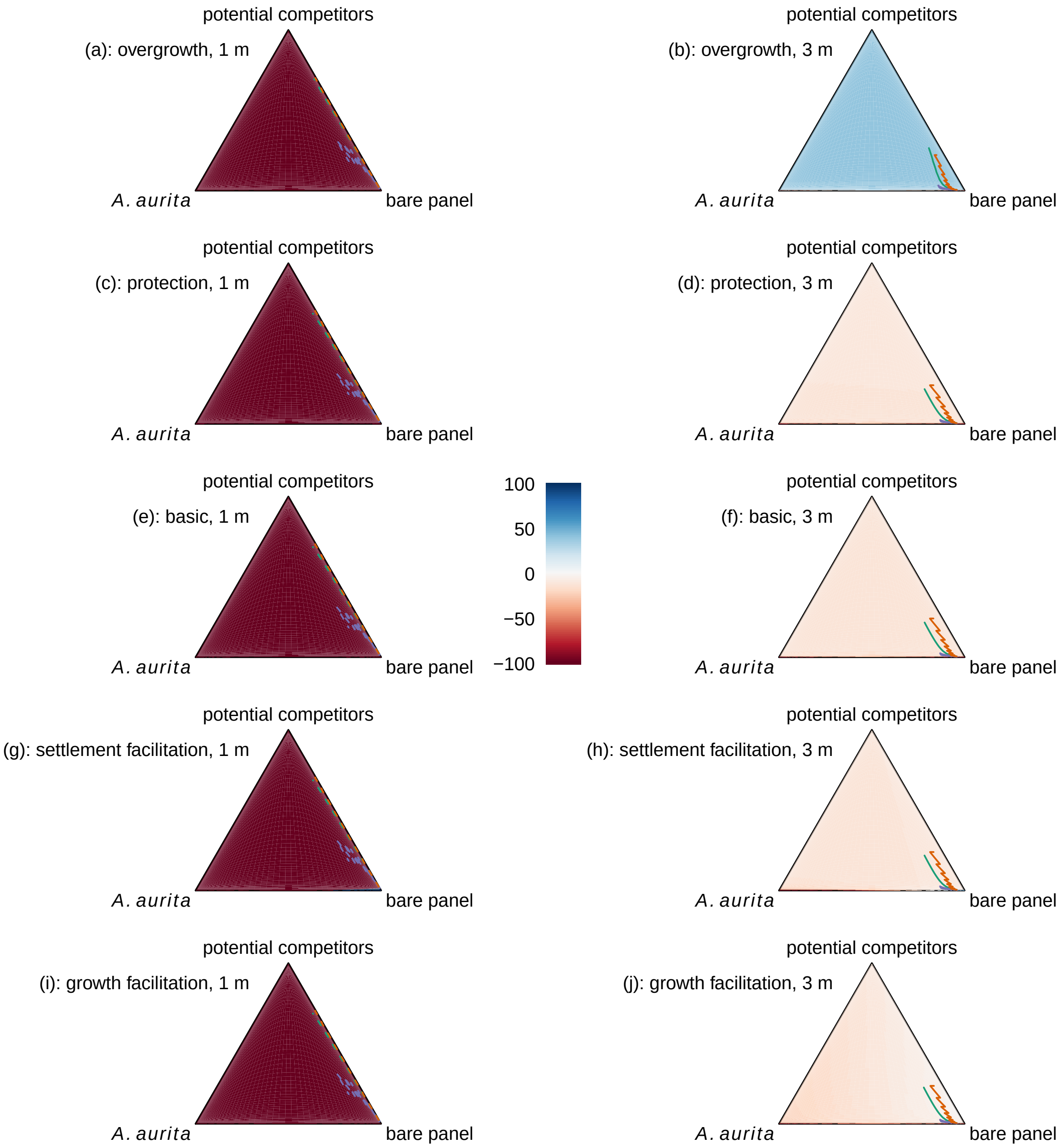


Figure S8: Effect of *A. aurita* polyps on proportional growth rate of potential competitors (colours represent posterior mean effects: positive blue, negative red, truncated at ±100) in the overgrowth (a, b), protection (c, d), basic (e, f), settlement facilitation (g, h) and growth facilitation (i, j) models, at 1 m (a, c, e, g, i) and 3 m (b, d, f, h, j). Models are ordered by $\text{elpd}_{\text{loo}}$, with the best model (overgrowth) at the top. Lines are posterior mean trajectories for typical panels. Dashed lines represent panels at 1 m, solid lines panels at 3 m. Line colours represent treatments: control ($C$) green, *A. aurita* removal ($A$) orange, potential competitor removal ($O$) purple.

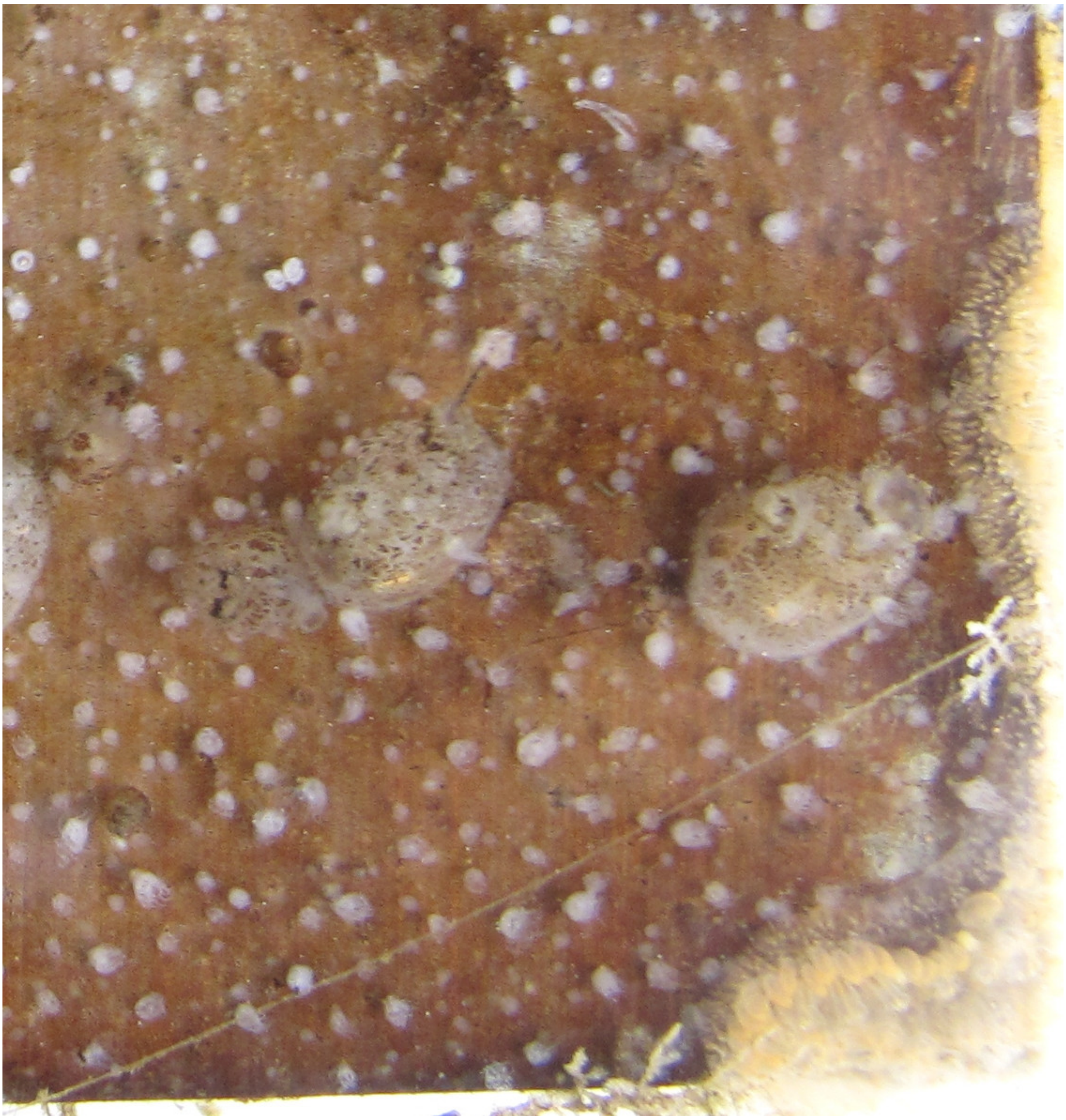

Figure S9: Apparent overgrowth of *A. aurita* polyps (small white translucent objects) by a colony of *Botrylloides sp.* (right, yellow). Some polyps can be seen apparently partially underneath the translucent margin of the colony. This is a closeup of the lower right corner of panel 3 (1 m depth, potential competitor removal treatment, 2019-09-24, pre-treatment, for which the whole panel is shown in Figure 2b in the main text.

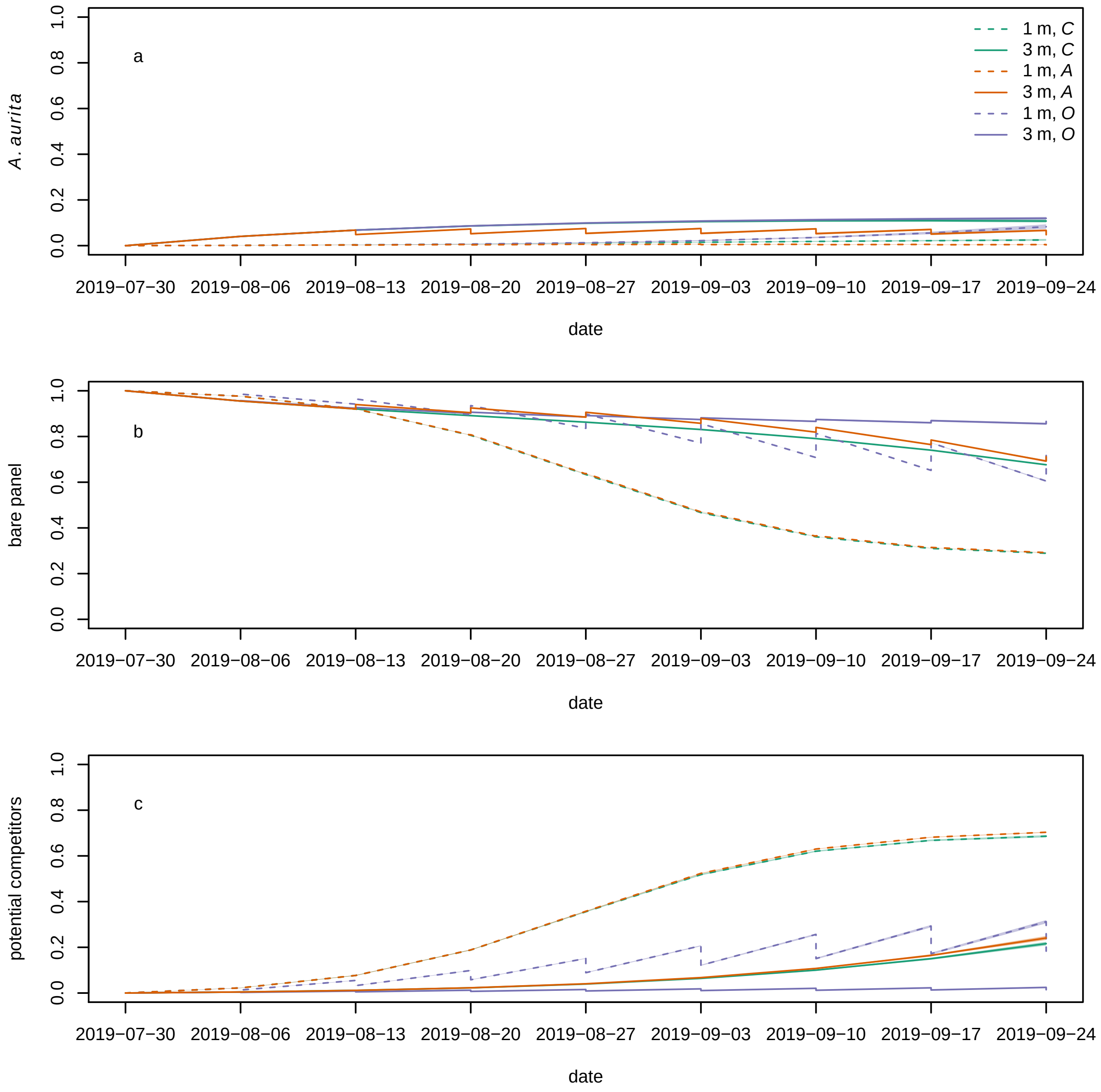


Figure S10: Modelled time series (basic model) for proportional cover of (a) *A. aurita*, (b) bare panel and (c) potential competitors. Each line is the posterior mean for a typical panel from a combination of treatment and depth. Dashed lines represent panels at 1 m, and solid lines panels at 3 m. Colours represent treatments: control ($C$) green, *A. aurita* removal ($A$) orange, potential competitor removal ($O$) purple. 95 % highest posterior density credible bands are shown, but are usually too narrow to be visible. Panels were put in the water on 2019-07-30.

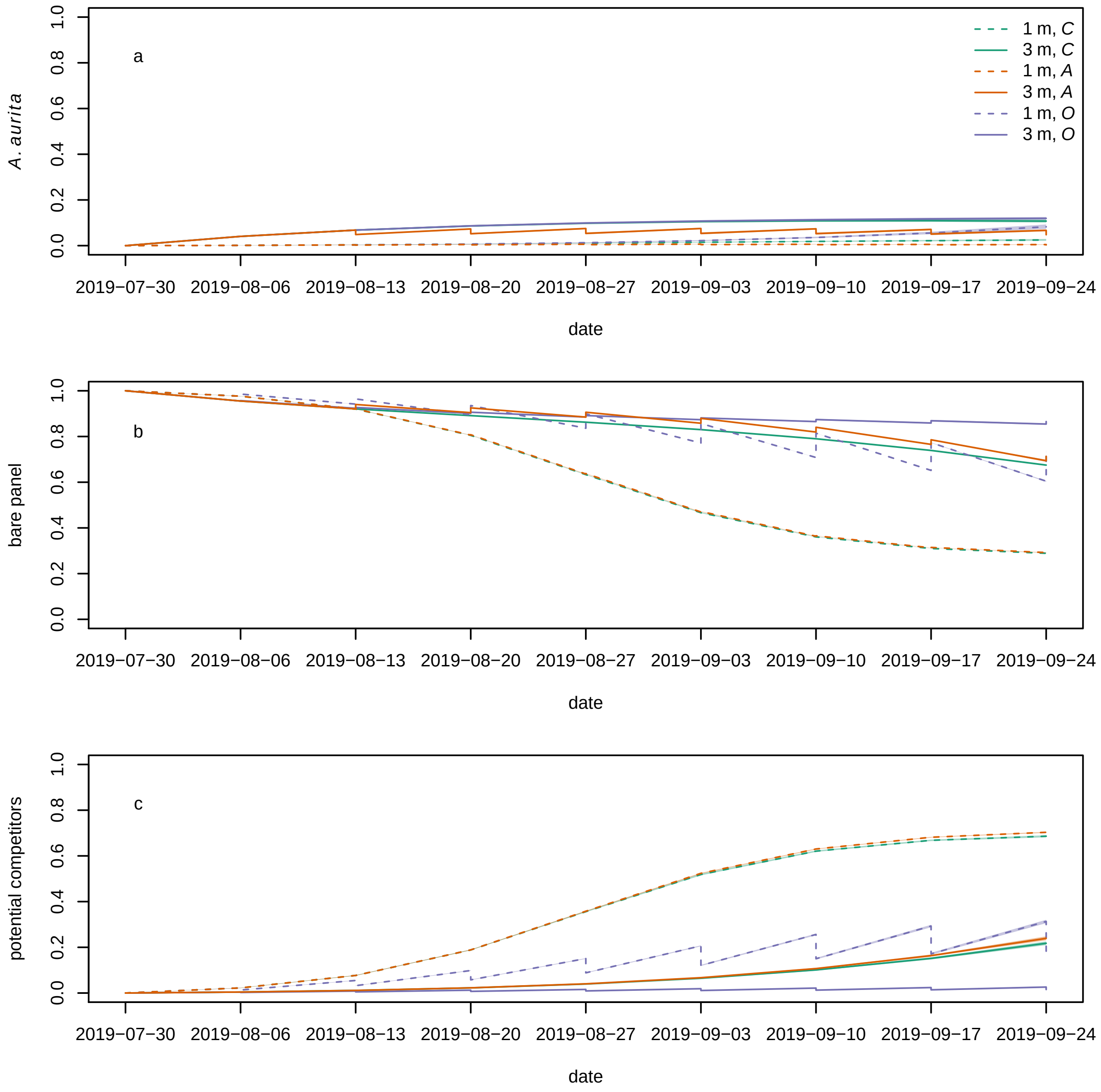


Figure S11: Modelled time series (model with settlement facilitation) for proportional cover of (a) *A. aurita*, (b) bare panel and (c) potential competitors. Each line is the posterior mean for a typical panel from a combination of treatment and depth. Dashed lines represent panels at 1 m, and solid lines panels at 3 m. Colours represent treatments: control ($C$) green, *A. aurita* removal ($A$) orange, potential competitor removal ($O$) purple. 95 % highest posterior density credible bands are shown, but are usually too narrow to be visible. Panels were put in the water on 2019-07-30.

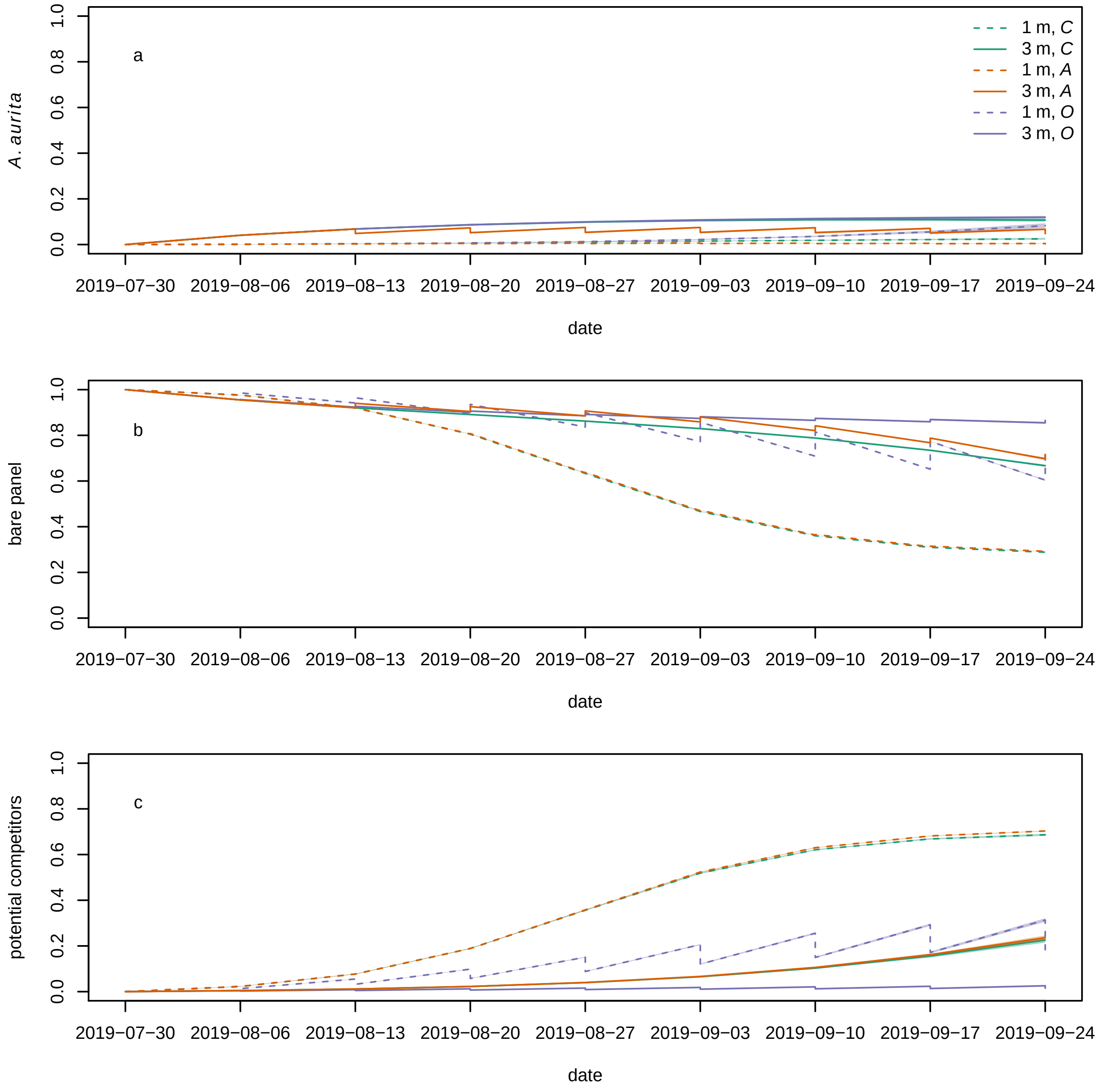


Figure S12: Modelled time series (model with growth facilitation) for proportional cover of (a) *A. aurita*, (b) bare panel and (c) potential competitors. Each line is the posterior mean for a typical panel from a combination of treatment and depth. Dashed lines represent panels at 1 m, and solid lines panels at 3 m. Colours represent treatments: control ($C$) green, *A. aurita* removal ($A$) orange, potential competitor removal ($O$) purple. 95 % credible bands are shown, but are usually too narrow to be visible. Panels were put in the water on 2019-07-30.

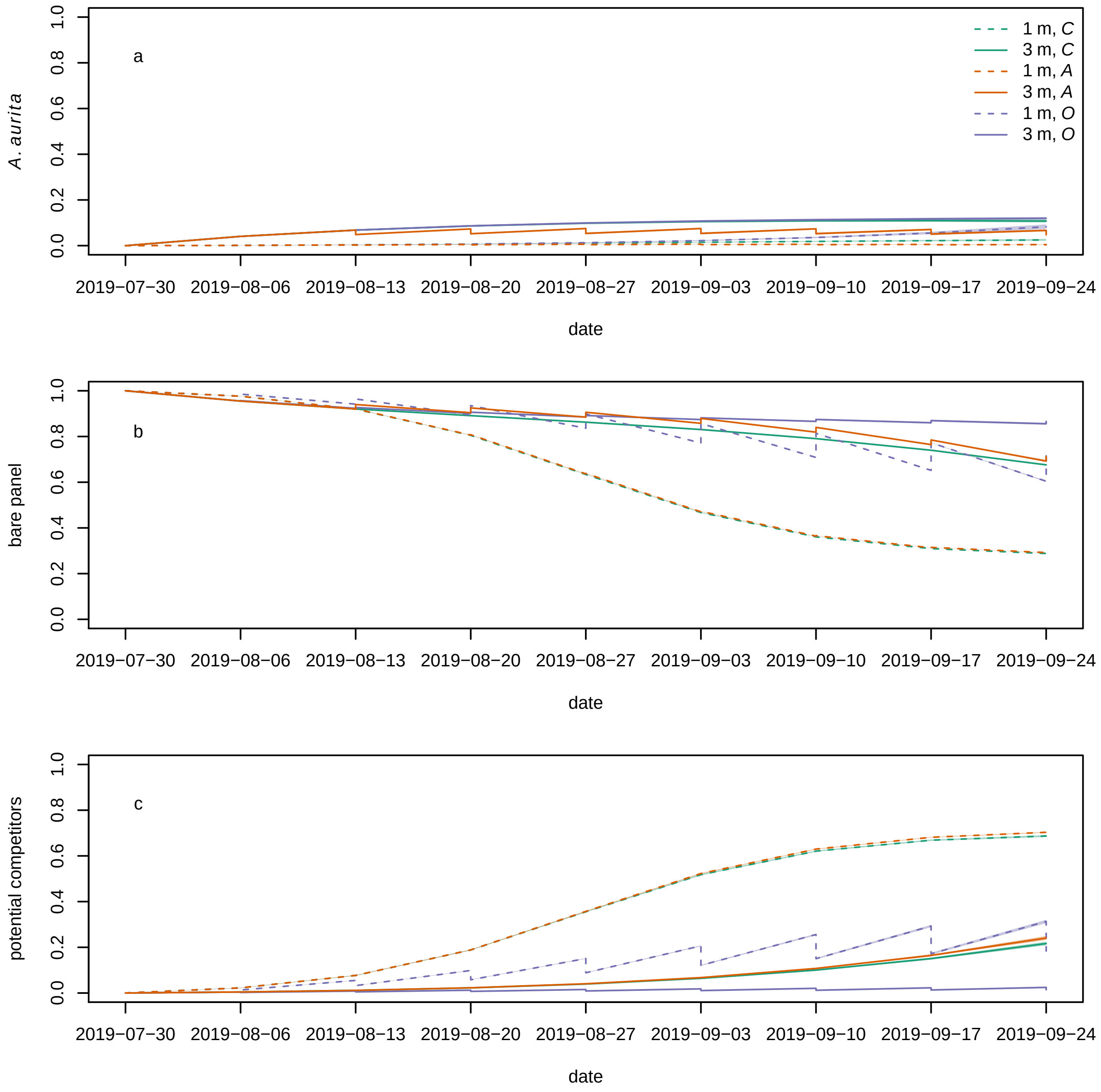


Figure S13: Modelled time series (model with protection from predators) for proportional cover of (a) *A. aurita*, (b) bare panel and (c) potential competitors. Each line is the posterior mean for a typical panel from a combination of treatment and depth. Dashed lines represent panels at 1 m, and solid lines panels at 3 m. Colours represent treatments: control ($C$) green, *A. aurita* removal ($A$) orange, potential competitor removal ($O$) purple. 95 % credible bands are shown, but are usually too narrow to be visible. Panels were put in the water on 2019-07-30.

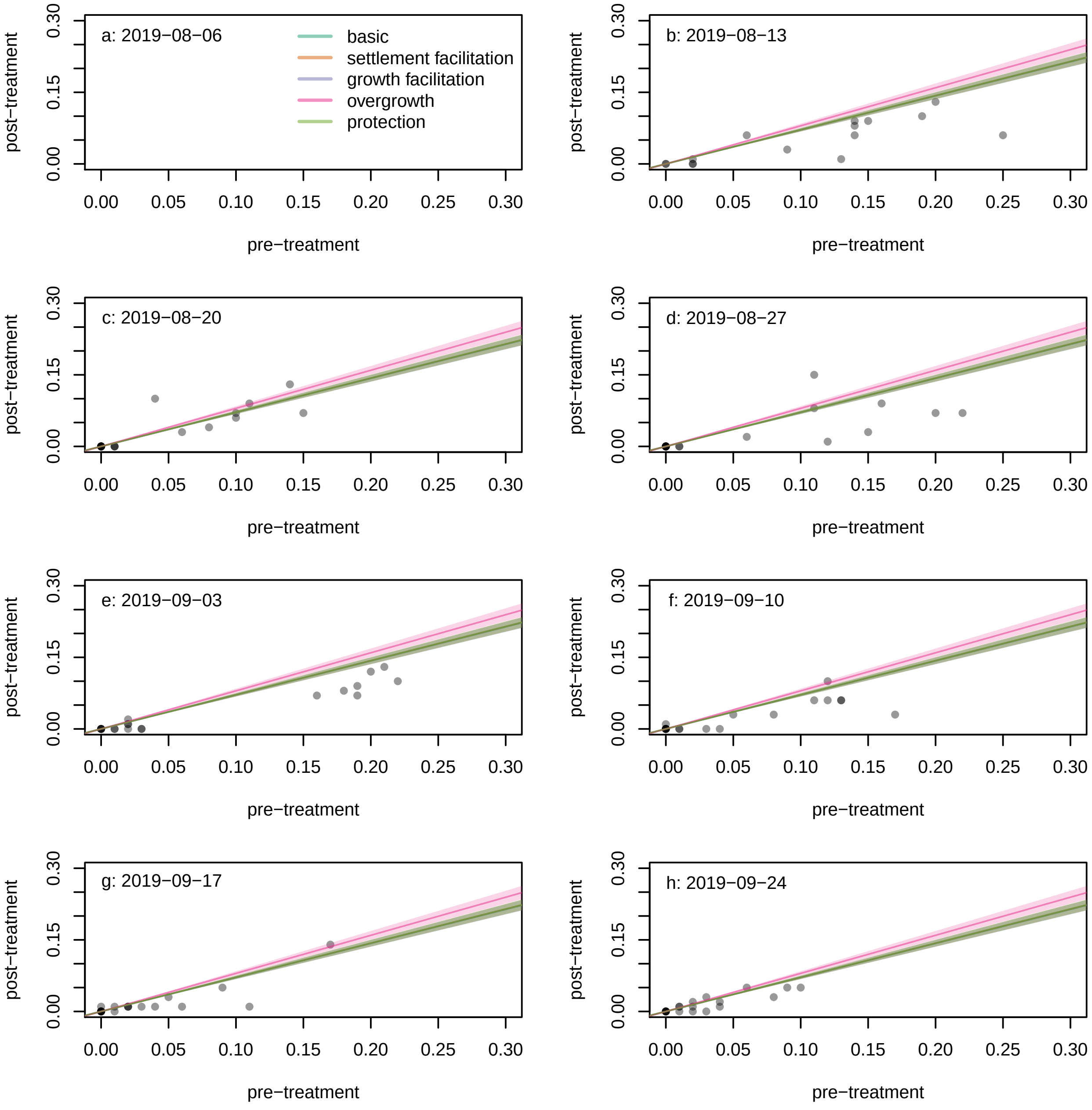


Figure S14: Comparison between post-treatment and pre-treatment sample proportions of space filled by *A. aurita* on panels from the *A* (*A. aurita* removal) treatment on each sample date. Each dot represents a single panel from the *A* treatment in a given week. Lines have slopes $1 - r_A$ and represent predictions from each dynamic model, with 95 % highest posterior density credible bands. Predictions from all models other than overgrowth are essentially indistinguishable and give overlapping lines. No post-treatment samples were taken on the first sample date because no *A. aurita* were observed.

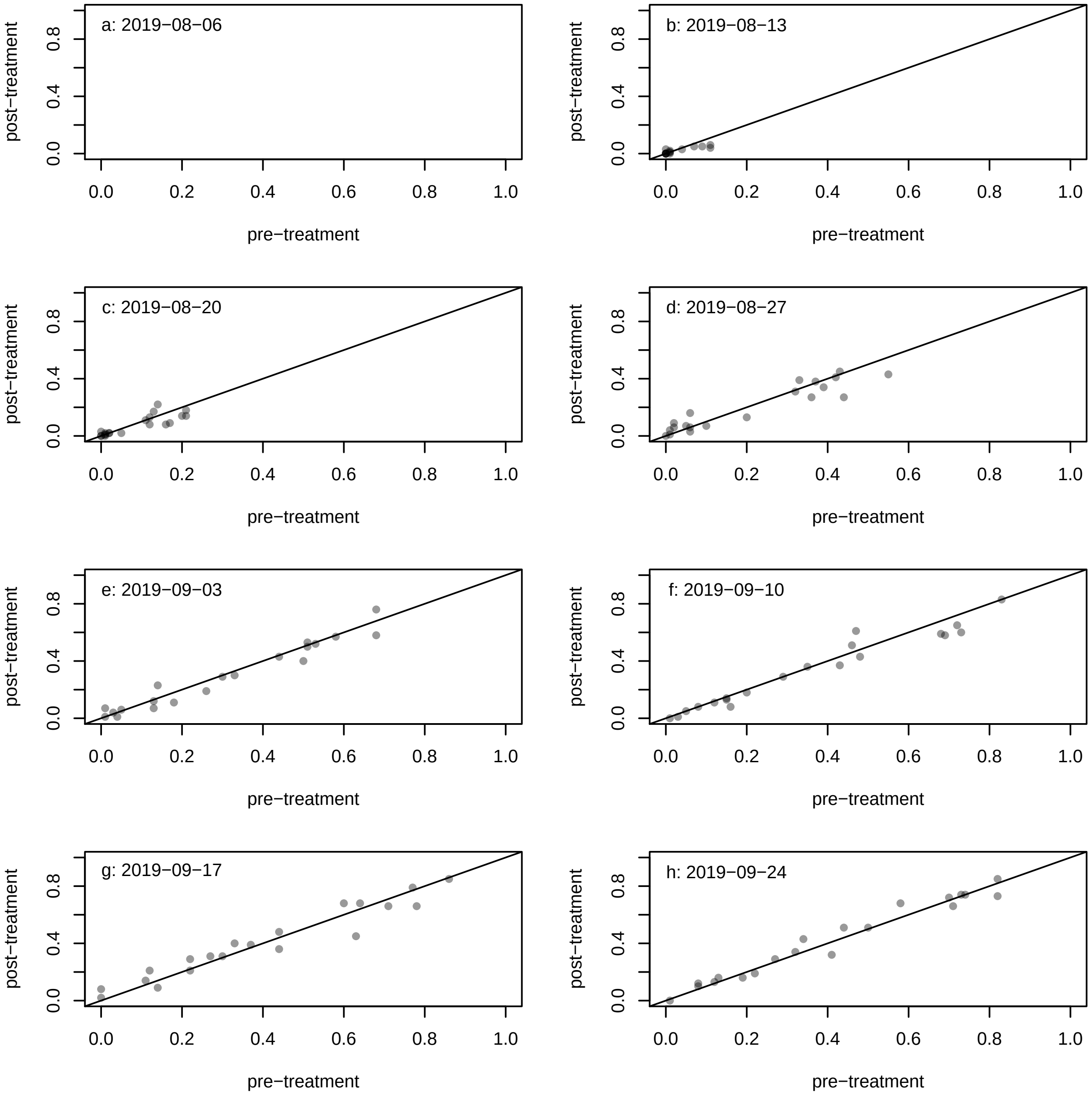


Figure S15: Comparison between post-treatment and pre-treatment sample proportions of space filled by potential competitors on panels from the $A$ (*A. aurita* removal) treatment on each sample date. Each dot represents a single panel from the $A$ treatment in a given week, and the line has intercept 0, slope 1. No post-treatment samples were taken on the first sample date because no *A. aurita* were observed.

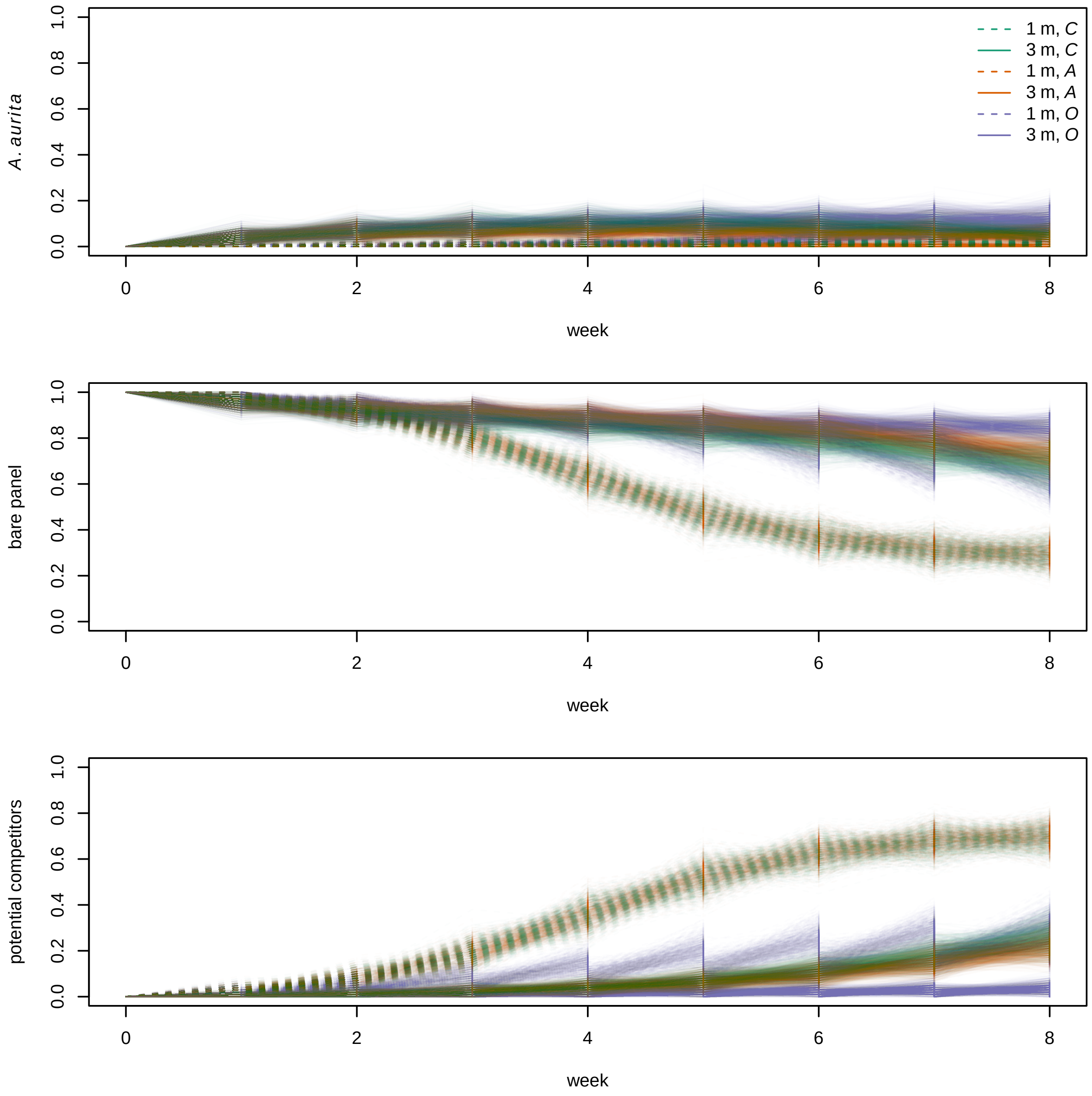


Figure S16: Posterior predictive simulation from the overgrowth model for proportional cover of (a) *A. aurita*, (b) bare panel and (c) potential competitors. Each line is a single posterior predictive simulation for a single panel, with parameters drawn from their posterior distributions, and observations generated by multinomial sampling. Dashed lines represent panels at 1 m, and solid lines panels at 3 m. Colours represent treatments: control ($C$) green, *A. aurita* removal ($A$) orange, potential competitor removal ($O$) purple. 100 simulated data sets are shown.

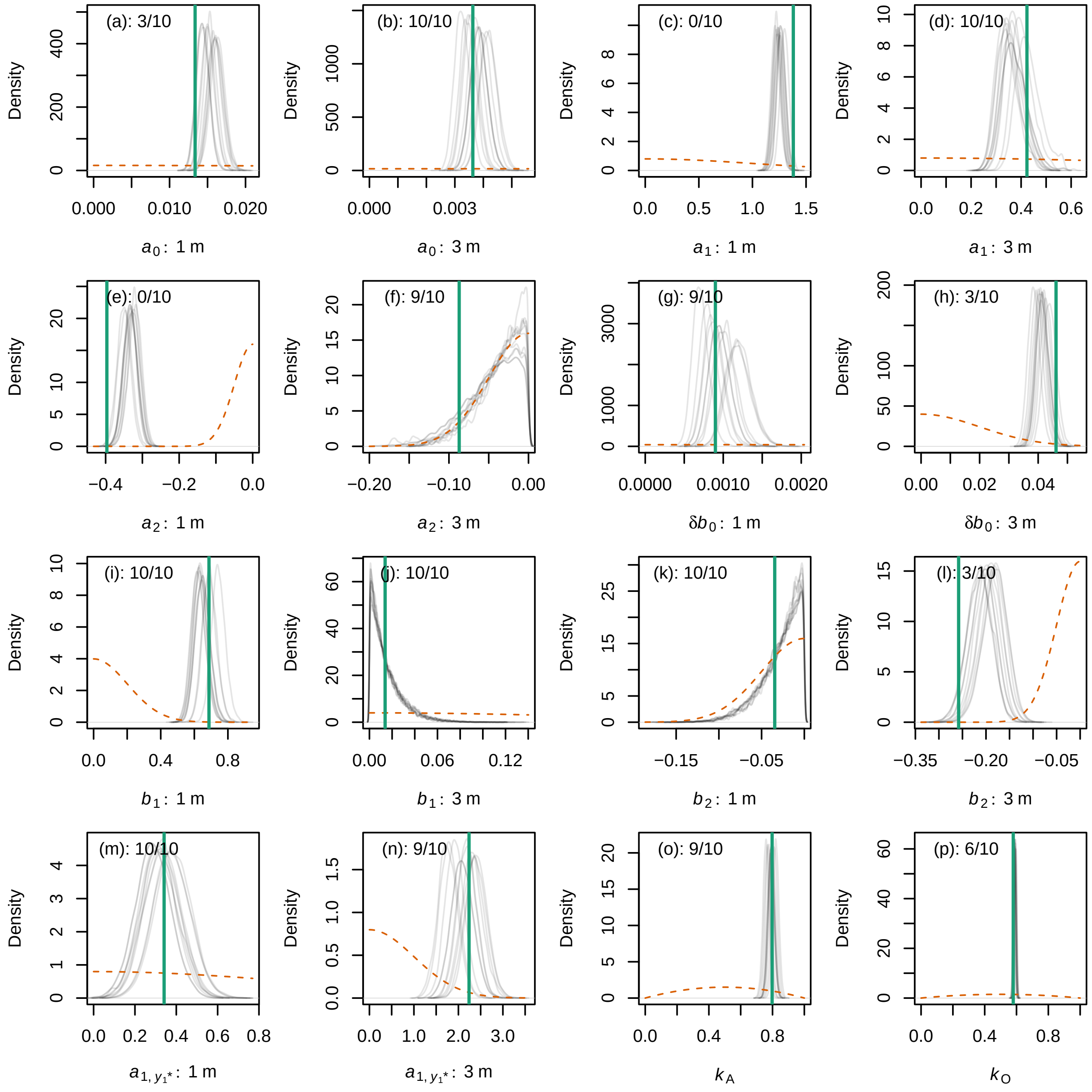


Figure S17: Performance of the overgrowth model on simulated data sets. Each panel represents a single parameter at a single depth, as labelled on the $x$-axis. Green vertical lines: true parameter values (posterior mean estimates from the real data, overgrowth model). Orange dashed lines: prior densities. Thin grey lines: posterior densities from each of 10 simulated data sets simulated under the overgrowth model, with the same structure as the real data. Numbers on panels: proportion of simulated data sets for which the posterior 95% HPD interval for the parameter contained the true value. Note that both $x$- and $y$-axis scales differ between panels.
