## Supplementary material for "Negative and positive interspecific interactions involving jellyfish polyps in marine sessile communities": Data and code: aurelia.R

```
rm(list = ls())
library(compositions)
library(RColorBrewer)
library(coda)
library(rstan)
library(mvtnorm)
library(rethinking)
library(hdrcde)
library(scatterplot3d)
library(mvQuad)
library(matrixStats)
library(loo)
library(xtable)
library(deSolve)
library(numDeriv)

source("aureliafuncs.R")

options(mc.cores = parallel::detectCores())
rstan_options(auto_write = TRUE)

makenewdata <- FALSE #rebuild data set from csv files?
choosepriors <- FALSE #choose some sensible priors and print out what they look like
setupmanova <- FALSE #set up data for the manova in stan?
readmanovaresults <- FALSE #extract results for manova from stan output files
loadmanovaresults <- FALSE #load already-extraced manova results from .RData
posteriorpredictive <- FALSE #posterior predictive checks
doSBC <- FALSE #simulation-based calibration of manova
summarizeSBC  <- FALSE #summarize results of simulation-based calibration of manova
plotmanovaresults <- FALSE #plot results from manova (and parameter tables)
hierarchicaldiagnostics <- FALSE #diagnostics for hierarchical parts of model
makeleaveoneoutdata <- FALSE #create leave-one-out cross validation data sets for Stan
processleaveoneoutdata <- FALSE #process stan leave-one-out output
processfull <- FALSE #which manova models to process within processleaveoneoutdata block
processnointeract <- FALSE
processnoscrape <- FALSE
processnodepth <- FALSE
comparecv <- FALSE #compare manova models using cross-validation
setupdynamicmodel <- FALSE #write data for dynamic models in stan
readoderesults <- FALSE #extract results for ODE model from stan output files
readodefacilresults <- FALSE #extract results for ODE model with facilitation and protection from stan output files
readodeFPFOresults <- FALSE #extract results for ODE model with FPFO from stan output files
readodeovergrowthresults <- FALSE #extract results for ODE model with overgrowth from stan output files
readodefrequencyresults <- FALSE #extract results for ODE model with frequency dependence from stan output files
readodefacilitationresults <- FALSE #extract results for ODE model with facilitation (only) from stan output files
readodeprotectionresults <- FALSE #extract results for ODE model with protection (only) from stan output files
readodegrowthfacilresults <- FALSE #extract results for ODE model with growth facilitation from stan output files

###########################
#only do one of these at a time:
loadoderesults <- FALSE #load already-extraded ODE results
loadodefacilresults <- FALSE #load already-extraded ODE results with facilitation and protection
loadodeFPFOresults <- FALSE #load already-extraded ODE results with FPFO
loadodeovergrowthresults <- FALSE #load already-extraded ODE results with overgrowth
loadodefrequencyresults <- FALSE #load already-extraded ODE results with frequency dependence
loadodefacilitationresults <- FALSE #load already-extraded ODE results with facilitation (only)
loadodeprotectionresults <- FALSE #load already-extraded ODE results with protection (only)
loadodegrowthfacilresults <- FALSE #load already-extraded ODE results with growth facilitation
################################

plotodepriors <- FALSE #plot solutions to ODE model with sensible priors
buildtypicalrho <- FALSE #ODE solution for typical panels from each combination
plotoderesults <- FALSE #plot results from ODE model (requires buildtypicalrho to have been run first)
checkAremoval  <- FALSE #check whether A treatment was also removing potential competitors
compareodemodels <- FALSE #compare fitted ODE models
checkcmatrix  <- FALSE #check calculation of community matrices
printODEtable <- FALSE #print ODE parameters to LaTeX
analyzeenvironmentaldata  <- FALSE #salinity, temperature, oxygen analyses
doSBCODE <- FALSE #SBC on ODE model
setupcondorSBCODE <- FALSE #write data for SBC on ODE model, for Condor
posteriorpredictiveODE <- FALSE #posterior predictive checks for ODE model
plotODEeffects <- FALSE #ternary plots of effect of A. aurita on proportional population growth rate of potential competitors
setupsimODEdata <- FALSE #generate simulated data sets for checking performance of estimation
checksimODEdata <- FALSE #process Stan output from simulated data sets
readrawsimoutput <- FALSE #read the raw Stan output from simulated data sets (if FALSE, load processed output from .RData file)

manovainputname <- "../data/manovainputs.RData" #input data for manova
ddinputname <- "../data/ddinputs.RData" #input data for dynamic models
nointeract <- TRUE #exclude interaction in manova model used in manuscript?

##################
#only set at most one of these to TRUE:
usefacil <- FALSE #use facilitation in ODE model?
useFPFO  <- FALSE #use facilitation, protection, frequency dependence and overgrowth?
useovergrowth <- TRUE #use overgrowth in ODE model?
usefrequency <- FALSE #use frequency dependence in ODE model?
usefacilitation <- FALSE #use facilitation (only) in ODE model?
useprotection <- FALSE #use protection (only) in ODE model?
usegrowthfacil <- FALSE #use growth facilitation (only) in ODE model?
###################

if(nointeract == TRUE){
    manovaoutputname <- "../../manovaresults/manovanointeract" #base of name for manova output files
} else {
    manovaoutputname <- "../../manovaresults/manova" #base of name for manova output files
}
manovastanfile <- "../stan/aureliamanova_sbc.stan" #stan file for manova, used only for SBC
odestanfile <- "../stan/aureliaode_overgrowth_sbc.stan" #stan file for ode model, used only for SBC
SBCODEdatafile <- "../condor_sbcode/sbcode_inputs.RData"
leaveoneoutpath <- "../../leaveoneoutdata/" #directory for leave-one-out cross-validation data
llminteractcvname <- "../../leaveoneoutdata/llminteractcv.RData" #save Monte Carlo marginal likelihood info here for manova with interaction, estimated by cross-validation
llmnointeractcvname <- "../../leaveoneoutdata/llmnointeractcv.RData" #save Monte Carlo marginal likelihood info here for manova without interaction, estimated by cross-validation
llmnoscrapecvname <- "../../leaveoneoutdata/llmnoscrapecv.RData" #save Monte Carlo marginal likelihood info here for manova without interaction or scrape, estimated by cross-validation
llmnodepthcvname <- "../../leaveoneoutdata/llmnodepthcv.RData" #save Monte Carlo marginal likelihood info here for manova without interaction or depth, estimated by cross-validation
odeoutputname <- "../../oderesults/ode" #base of name for ode output files
odefaciloutputname <- "../../oderesults/ode_facil" #base of name for ode output files with facilitation and protection
odeFPFOoutputname <- "../../oderesults/ode_FPFO" #base of name for ode output files with FPFO
odeovergrowthoutputname <- "../../oderesults/ode_overgrowth" #base of name for ode output files with overgrowth
odefrequencyoutputname <- "../../oderesults/ode_frequency" #base of name for ode output files with frequency dependence
odefacilitationoutputname <- "../../oderesults/ode_facilitation" #base of name for ode output files with facilitation (only)
odeprotectionoutputname <- "../../oderesults/ode_protection" #base of name for ode output files with protection (only)
odegrowthfaciloutputname <- "../../oderesults/ode_growthfacil" #base of name for ode output files with growth facilitation
envdataname  <- "../data/environmentaldata.txt" #file containing temperature, oxygen, salinity
simodename <- "../../simodedata/simode" #base of name for simulated ode data files
simodeprocessedname <- "../../simodedata/simode_processed.RData" #name for processed simulated ode results
if(makenewdata == TRUE){
    compiledata()
}

alldata <- read.table(file = "../data/alldata.txt", header = TRUE, sep = "\t", stringsAsFactors = TRUE)
taxa <- c("Aurelia.aurita.on.panel", "Bare.Panel", "Botrylloides.spp.", "Bugula.spp.", "Molgula.tubifera") #taxa included in analysis
ntaxa <- length(taxa)
V <- makeV(D = ntaxa, choice = "B1") #ilr basis matrix
lastdate <- "2019-09-24" #final week

if(choosepriors == TRUE){
    mytaxa <- c(8, 1, 9, 2, 3) #Aurelia, bare wall, colonial ascidians, Bugula, solitary ascidians in Chong and Spencer 2018
    makedepthprior(Chongfilename = "../../stanresults/Chong18coefficients.RData", mytaxa = mytaxa, V = V, figname = "../latex/depthprior.pdf")
    makemuprior(V = V, vmu = 4)
    makescrapeprior(changesize = c(2, 3, 4, 5), V = V, fname = "../latex/scrapeprior.pdf")
}

if(setupmanova == TRUE){
    manovainputs <- aureliamanovasetup(alldata = alldata, date = lastdate, preorpost = "Pre", taxa = taxa, tVinv = V, standatafilename = "../data/manovadata.txt")
    save(manovainputs, file = manovainputname)
}

if(readmanovaresults == TRUE){
    fnames <- c(paste(manovaoutputname, "1.csv", sep = ""), paste(manovaoutputname, "2.csv", sep = ""), paste(manovaoutputname, "3.csv", sep = ""), paste(manovaoutputname, "4.csv", sep = ""))
    manovadata <- read_stan_csv(csvfiles = fnames)
    if(nointeract == TRUE){
        chainsummary <- summary(manovadata, pars = c("mu", "alpha", "beta1", "beta2", "zpanel", "zblock", "LSigma", "LZeta", "lp__"))$summary
        plot(rstan::traceplot(manovadata, pars = c("mu", "alpha", "beta1", "beta2", "LSigma", "LZeta", "lp__"), inc_warmup = FALSE))
    } else {
        chainsummary <- summary(manovadata, pars = c("mu", "alpha", "beta1", "beta2", "gamma1", "gamma2", "zpanel", "zblock", "LSigma", "LZeta", "lp__"))$summary
        plot(rstan::traceplot(manovadata, pars = c("mu", "alpha", "beta1", "beta2", "gamma1", "gamma2", "LSigma", "LZeta", "lp__"), inc_warmup = FALSE))
    }
    print(chainsummary)
    print("Summary of n_eff:")
    print(summary(chainsummary[, 9]))
    print("Summary of Rhat:")
    print(summary(chainsummary[, 10]))
          
    X11()
    plot(rstan::traceplot(manovadata, pars = c("zblock"), inc_warmup = FALSE))
    X11()
    plot(rstan::traceplot(manovadata, pars = c("zpanel"), inc_warmup = FALSE))
    mu <- extract(manovadata, pars = "mu")[[1]]
    alpha <- extract(manovadata, pars = "alpha")[[1]]
    beta1 <- extract(manovadata, pars = "beta1")[[1]]
    beta2 <- extract(manovadata, pars = "beta2")[[1]]
    LSigma <- extract(manovadata, pars = "LSigma")[[1]]
    LZeta <- extract(manovadata, pars = "LZeta")[[1]]
    zpanel <- extract(manovadata, pars = "zpanel")[[1]]
    zblock <- extract(manovadata, pars = "zblock")[[1]]    

    if(nointeract == TRUE){
        gamma1 <- array(0, dim = dim(alpha))
        gamma2 <- array(0, dim = dim(alpha))
    } else {
        gamma1 <- extract(manovadata, pars = "gamma1")[[1]]
        gamma2 <- extract(manovadata, pars = "gamma2")[[1]]    
    }
    savelist <- c("mu", "alpha", "beta1", "beta2", "gamma1", "gamma2", "LSigma", "LZeta", "zpanel", "zblock")
    save(list = savelist, file = paste(manovaoutputname, ".RData", sep = ""))
}

if(loadmanovaresults == TRUE) {
    load(file = paste(manovaoutputname, ".RData", sep = ""))
    load(file = manovainputname)
    manovapredictions <- panelpredict(LZeta = LZeta, zblock = zblock, LSigma = LSigma, zpanel = zpanel, blockindex = manovainputs$blockindex, d1 = manovainputs$d1, d2 = manovainputs$d2, d3 = manovainputs$d3, d4 = manovainputs$d4, d5 = manovainputs$d5, mu = mu, alpha = alpha, beta1 = beta1, beta2 = beta2, gamma1 = gamma1, gamma2 = gamma2, V = V)
}

if(posteriorpredictive == TRUE) {
    ppsample <- posteriorsample(rho = manovapredictions$panelsimplexprediction, niter = 1e3, counts = manovainputs$counts)
    manovapredvsobsplot(ppsample$pp, manovainputs$counts, ntaxa, fname = "../diagnosticplots/predproportions.pdf")
    rawcounteffectspp(alldata = alldata, date = lastdate, preorpost = "Pre", pp = ppsample, niter = 1e3, fname = "../diagnosticplots/predteststats.pdf")
}   

if(doSBC == TRUE){
    sbcoutput <- SBC(manovastanfile = manovastanfile, manovainputs = manovainputs, ntaxa = ntaxa, V = V, M = 500, iter = 5000)
    save(sbcoutput, file = "../../manovaresults/SBC.RData")
}

if(summarizeSBC == TRUE){
    load(file = "../../manovaresults/SBC.RData")
    sbcranks <- getsbcranks(sbcoutput)
    myplot.sbc(sbcranks = sbcranks, manovaresultspath = "../../manovaresults")
}

if(plotmanovaresults == TRUE){
    plotblockpredictions(manovapredictions = manovapredictions, fname = "../diagnosticplots/blockpredictions.pdf")
    plotpanelpredictions(manovapredictions = manovapredictions, fname = "../diagnosticplots/panelpredictions.pdf")
    pdf(file = "../latex/manovaplot.pdf", width = 7, height = 7)
    par(xpd = NA)
    par(mfrow = c(2, 1))
    manovaplot(manovainputs = manovainputs, alldata = alldata, mu = mu, alpha = alpha, beta1 = beta1, beta2 = beta2, gamma1 = gamma1, gamma2 = gamma2, date = lastdate, preorpost = "Pre", V = V, centre = TRUE, coords = c(1:2), plotcoords = c(1:3), labels = c(expression(italic("A. aurita")),"Bare panel", "gm(potential competitors)"), fname = NA, paneltext = "a", plotlegend = TRUE)
    manovaplot(manovainputs = manovainputs, alldata = alldata, mu = mu, alpha = alpha, beta1 = beta1, beta2 = beta2, gamma1 = gamma1, gamma2 = gamma2, date = lastdate, preorpost = "Pre", V = V, centre = TRUE, coords = c(3:4), plotcoords = c(3:5), labels = c(expression(italic("Botrylloides spp.")), expression(italic("Bugula spp.")), expression(italic("Molgula tubifera"))), fname = NA, paneltext = "b", plotlegend = FALSE)
    dev.off()
    testeffects(mu = mu, alpha = alpha, beta1 = beta1, beta2 = beta2, gamma1 = gamma1, gamma2 = gamma2, V = V, fname = "../latex/logitresponses.pdf")
    plothyperellipsoids(LSigma = LSigma, LZeta = LZeta, n = 100, m = 1e3, cov12fname = "../latex/cov12.pdf", cov34fname = "../latex/cov34.pdf", labels12 = c(expression(italic("A. aurita")),"Bare panel", "gm(potential competitors)"), labels34 = c(expression(italic("Botrylloides spp.")), expression(italic("Bugula spp.")), expression(italic("Molgula tubifera"))), alpha = 0.2, method = "2d")
    printmanovaparameters(mu = mu, alpha = alpha, beta1 = beta1, beta2 = beta2, gamma1 = gamma1, gamma2 = gamma2, LSigma = LSigma, LZeta = LZeta, fname = "../latex/manovaparameters.tex", includeinteraction = !nointeract)

}

if(hierarchicaldiagnostics == TRUE){
    hierarchicalqqplots(y = manovapredictions$paneleffect, LSigma = LSigma, fname = "../diagnosticplots/panelqq.pdf", n = 1e3)
    hierarchicalqqplots(y = manovapredictions$blockeffect, LSigma = LZeta, fname = "../diagnosticplots/blockqq.pdf", n = 1e3)
}

if(makeleaveoneoutdata == TRUE){
    leaveoneoutname <- paste(leaveoneoutpath, "leaveoneout", sep = "")
    writeleaveoneout(alldata = alldata, date = lastdate, preorpost = "Pre", taxa = taxa, tVinv = V, leaveoneoutname = leaveoneoutname)
}

if(processleaveoneoutdata == TRUE){
    load(file = manovainputname)
    niter <- 1e3 #number of stan iterations to use
    mciter <- 1e3 #number of Monte Carlo iterations to use to approximate integrals over block and panel effects
    nblocks <- length(unique(manovainputs$blockindex))
    if(processfull == TRUE){
        llminteract <- array(dim = c(niter, nblocks)) #stan iterations x blocks: marginal log likelihood for new block.

        for(l in 1:nblocks){#number in file name must indicate the block that was left out
            ipattern <- paste("leaveoneout_", l, ".txt[[:digit:]].csv", sep = "")
            csvfiles <- list.files(path = leaveoneoutpath, pattern = ipattern)
            print(csvfiles)
            llminteract[, l] <- getcv(leaveoneoutpath = leaveoneoutpath, csvfiles = csvfiles, l = l, mciter = mciter, niter = niter, model = "interaction", V = V, manovainputs = manovainputs)
        }
        save(llminteract, file = llminteractcvname)
    }
    if(processnointeract == TRUE){
        llmnointeract <- array(dim = c(niter, nblocks)) #stan iterations x blocks: marginal log likelihood for new block.
        for(l in 1:nblocks){#number in file name must indicate the block that was left out
            ipattern <- paste("leaveoneout_", l, ".txt[[:digit:]].nointeraction.csv", sep = "")
            csvfiles <- list.files(path = leaveoneoutpath, pattern = ipattern)
            print(csvfiles)
            llmnointeract[, l] <- getcv(leaveoneoutpath = leaveoneoutpath, csvfiles = csvfiles, l = l, mciter = mciter, niter = niter, model = "nointeraction", V = V, manovainputs = manovainputs)
        }
        save(llmnointeract, file = llmnointeractcvname)
    }
    if(processnoscrape == TRUE){
        llmnoscrape <- array(dim = c(niter, nblocks)) #stan iterations x blocks: marginal log likelihood for new block.
        for(l in 1:nblocks){#number in file name must indicate the block that was left out
            ipattern <- paste("leaveoneout_", l, ".txt[[:digit:]].noscrape.csv", sep = "")
            csvfiles <- list.files(path = leaveoneoutpath, pattern = ipattern)
            print(csvfiles)
            llmnoscrape[, l] <- getcv(leaveoneoutpath = leaveoneoutpath, csvfiles = csvfiles, l = l, mciter = mciter, niter = niter, model = "noscrape", V = V, manovainputs = manovainputs)
        }
        save(llmnoscrape, file = llmnoscrapecvname)
    }
    if(processnodepth == TRUE){
        llmnodepth <- array(dim = c(niter, nblocks)) #stan iterations x blocks: marginal log likelihood for new block.
        for(l in 1:nblocks){#number in file name must indicate the block that was left out
            ipattern <- paste("leaveoneout_", l, ".txt[[:digit:]].nodepth.csv", sep = "")
            csvfiles <- list.files(path = leaveoneoutpath, pattern = ipattern)
            print(csvfiles)
            llmnodepth[, l] <- getcv(leaveoneoutpath = leaveoneoutpath, csvfiles = csvfiles, l = l, mciter = mciter, niter = niter, model = "nodepth", V = V, manovainputs = manovainputs)
        }
        save(llmnodepth, file = llmnodepthcvname)
    }
    
}

if(comparecv == TRUE){
    load(llminteractcvname)
    load(llmnointeractcvname)
    load(llmnoscrapecvname)
    load(llmnodepthcvname)
    cvinteract <- kfold(llminteract)
    cvnointeract <- kfold(llmnointeract)
    cvnoscrape <- kfold(llmnoscrape)
    cvnodepth <- kfold(llmnodepth)
    print("interaction vs no interaction:")
    print(loo_compare(cvinteract, cvnointeract))
    print("no interaction vs no interaction, no scrape:")
    print(loo_compare(cvnointeract, cvnoscrape))
    print("no interaction vs no interaction, no depth:")
    print(loo_compare(cvnointeract, cvnodepth))
}

if(setupdynamicmodel == TRUE){
    dynamicdata <- datafordynamicmodel(alldata = alldata, potentialcompetitors = c("Botrylloides.spp.", "Bugula.spp.", "Molgula.tubifera"), treatmentappliedfile = "../data/treatmentapplied.txt", taxaforstan = c("Aurelia.aurita.on.panel", "Bare.Panel", "potentialcomp"), standatafilename = "../data/dynamicdata.txt")
    save(dynamicdata, file = ddinputname)
}

if(readoderesults == TRUE){
    fnames <- c(paste(odeoutputname, "1.csv", sep = ""), paste(odeoutputname, "2.csv", sep = ""), paste(odeoutputname, "3.csv", sep = ""), paste(odeoutputname, "4.csv", sep = ""))
    odedata <- read_stan_csv(csvfiles = fnames)
    chainsummary <- summary(odedata, pars = c("a0", "a1", "a2", "deltab0", "b1", "b2", "kA", "kO", "lp__"))$summary
    plot(rstan::traceplot(odedata, pars = c("a0", "a1", "a2", "deltab0", "b1", "b2", "kA", "kO", "lp__"), inc_warmup = FALSE))

    print(chainsummary)
    print("Summary of n_eff for dynamic parameters:")
    print(summary(chainsummary[, 9]))
    print("Summary of Rhat for dynamic parameters:")
    print(summary(chainsummary[, 10]))

    rhosummary <- summary(odedata, pars = c("rho"))$summary
    print("Summary of n_eff for rho:")
    print(summary(rhosummary[, 9]))
    print("Summary of Rhat for rho:")
    print(summary(rhosummary[, 10]))
    
    a0 <- extract(odedata, pars = "a0")[[1]]
    a1 <- extract(odedata, pars = "a1")[[1]]
    a2 <- extract(odedata, pars = "a2")[[1]]
    deltab0 <- extract(odedata, pars = "deltab0")[[1]]
    b1 <- extract(odedata, pars = "b1")[[1]]
    b2 <- extract(odedata, pars = "b2")[[1]]
    kA <- extract(odedata, pars = "kA")[[1]]
    kO <- extract(odedata, pars = "kO")[[1]]
    rho <- extract(odedata, pars = "rho")[[1]]
    looode <- loo(odedata, save_psis = TRUE)
    print(looode)
    savelist <- c("a0", "a1", "a2", "deltab0", "b1", "b2", "kA", "kO", "rho", "looode")
    save(list = savelist, file = paste(odeoutputname, ".RData", sep = ""))
}

if(readodefacilresults == TRUE){
    fnames <- c(paste(odefaciloutputname, "1.csv", sep = ""), paste(odefaciloutputname, "2.csv", sep = ""), paste(odefaciloutputname, "3.csv", sep = ""), paste(odefaciloutputname, "4.csv", sep = ""))
    odefacildata <- read_stan_csv(csvfiles = fnames)
    chainsummary <- summary(odefacildata, pars = c("a0", "a1", "a2", "deltab0", "b1", "b2", "kA", "kO", "m0", "m2", "lp__"))$summary
    plot(rstan::traceplot(odefacildata, pars = c("a0", "a1", "a2", "deltab0", "b1", "b2", "kA", "kO", "m0", "m2", "lp__"), inc_warmup = FALSE))

    print(chainsummary)
    print("Summary of n_eff for dynamic parameters:")
    print(summary(chainsummary[, 9]))
    print("Summary of Rhat for dynamic parameters:")
    print(summary(chainsummary[, 10]))

    rhosummary <- summary(odefacildata, pars = c("rho"))$summary
    print("Summary of n_eff for rho:")
    print(summary(rhosummary[, 9]))
    print("Summary of Rhat for rho:")
    print(summary(rhosummary[, 10]))
    
    a0 <- extract(odefacildata, pars = "a0")[[1]]
    a1 <- extract(odefacildata, pars = "a1")[[1]]
    a2 <- extract(odefacildata, pars = "a2")[[1]]
    deltab0 <- extract(odefacildata, pars = "deltab0")[[1]]
    b1 <- extract(odefacildata, pars = "b1")[[1]]
    b2 <- extract(odefacildata, pars = "b2")[[1]]
    kA <- extract(odefacildata, pars = "kA")[[1]]
    kO <- extract(odefacildata, pars = "kO")[[1]]
    rho <- extract(odefacildata, pars = "rho")[[1]]
    m0 <- extract(odefacildata, pars = "m0")[[1]]
    m2 <- extract(odefacildata, pars = "m2")[[1]]
    savelist <- c("a0", "a1", "a2", "deltab0", "b1", "b2", "kA", "kO", "rho", "m0", "m2")
    save(list = savelist, file = paste(odefaciloutputname, ".RData", sep = ""))
}

if(readodeFPFOresults == TRUE){
    fnames <- c(paste(odeFPFOoutputname, "1.csv", sep = ""), paste(odeFPFOoutputname, "2.csv", sep = ""), paste(odeFPFOoutputname, "3.csv", sep = ""), paste(odeFPFOoutputname, "4.csv", sep = ""))
    odeFPFOdata <- read_stan_csv(csvfiles = fnames)
    chainsummary <- summary(odeFPFOdata, pars = c("a0", "a1", "a1y1star", "a2", "deltab0", "b1", "b2", "kA", "kO", "m0", "m2", "psi", "lp__"))$summary
    plot(rstan::traceplot(odeFPFOdata, pars = c("a0", "a1", "a1y1star", "a2", "deltab0", "b1", "b2", "kA", "kO", "m0", "m2", "psi", "lp__"), inc_warmup = FALSE))
    print(chainsummary)
    print("Summary of n_eff for dynamic parameters:")
    print(summary(chainsummary[, 9]))
    print("Summary of Rhat for dynamic parameters:")
    print(summary(chainsummary[, 10]))

    rhosummary <- summary(odeFPFOdata, pars = c("rho"))$summary
    print("Summary of n_eff for rho:")
    print(summary(rhosummary[, 9]))
    print("Summary of Rhat for rho:")
    print(summary(rhosummary[, 10]))
    
    a0 <- extract(odeFPFOdata, pars = "a0")[[1]]
    a1 <- extract(odeFPFOdata, pars = "a1")[[1]]
    a1y1star <- extract(odeFPFOdata, pars = "a1y1star")[[1]]
    a2 <- extract(odeFPFOdata, pars = "a2")[[1]]
    deltab0 <- extract(odeFPFOdata, pars = "deltab0")[[1]]
    b1 <- extract(odeFPFOdata, pars = "b1")[[1]]
    b2 <- extract(odeFPFOdata, pars = "b2")[[1]]
    kA <- extract(odeFPFOdata, pars = "kA")[[1]]
    kO <- extract(odeFPFOdata, pars = "kO")[[1]]
    rho <- extract(odeFPFOdata, pars = "rho")[[1]]
    m0 <- extract(odeFPFOdata, pars = "m0")[[1]]
    m2 <- extract(odeFPFOdata, pars = "m2")[[1]]
    psi <- extract(odeFPFOdata, pars = "psi")[[1]]

    looFPFO <- loo(odeFPFOdata, save_psis = TRUE)
    print(looFPFO)
    savelist <- c("a0", "a1", "a1y1star", "a2", "deltab0", "b1", "b2", "kA", "kO", "rho", "m0", "m2", "psi", "looFPFO")
    save(list = savelist, file = paste(odeFPFOoutputname, ".RData", sep = ""))
}

if(readodeovergrowthresults == TRUE){
    fnames <- c(paste(odeovergrowthoutputname, "1.csv", sep = ""), paste(odeovergrowthoutputname, "2.csv", sep = ""), paste(odeovergrowthoutputname, "3.csv", sep = ""), paste(odeovergrowthoutputname, "4.csv", sep = ""))
    odeovergrowthdata <- read_stan_csv(csvfiles = fnames)
    chainsummary <- summary(odeovergrowthdata, pars = c("a0", "a1", "a1y1star", "a2", "deltab0", "b1", "b2", "kA", "kO", "lp__"))$summary
    plot(rstan::traceplot(odeovergrowthdata, pars = c("a0", "a1", "a1y1star", "a2", "deltab0", "b1", "b2", "kA", "kO", "lp__"), inc_warmup = FALSE))
    print(chainsummary)
    print("Summary of n_eff for dynamic parameters:")
    print(summary(chainsummary[, 9]))
    print("Summary of Rhat for dynamic parameters:")
    print(summary(chainsummary[, 10]))

    rhosummary <- summary(odeovergrowthdata, pars = c("rho"))$summary
    print("Summary of n_eff for rho:")
    print(summary(rhosummary[, 9]))
    print("Summary of Rhat for rho:")
    print(summary(rhosummary[, 10]))
    
    a0 <- extract(odeovergrowthdata, pars = "a0")[[1]]
    a1 <- extract(odeovergrowthdata, pars = "a1")[[1]]
    a1y1star <- extract(odeovergrowthdata, pars = "a1y1star")[[1]]
    a2 <- extract(odeovergrowthdata, pars = "a2")[[1]]
    deltab0 <- extract(odeovergrowthdata, pars = "deltab0")[[1]]
    b1 <- extract(odeovergrowthdata, pars = "b1")[[1]]
    b2 <- extract(odeovergrowthdata, pars = "b2")[[1]]
    kA <- extract(odeovergrowthdata, pars = "kA")[[1]]
    kO <- extract(odeovergrowthdata, pars = "kO")[[1]]
    rho <- extract(odeovergrowthdata, pars = "rho")[[1]]
    looovergrowth <- loo(odeovergrowthdata, save_psis = TRUE)
    print(looovergrowth)
    savelist <- c("a0", "a1", "a1y1star", "a2", "deltab0", "b1", "b2", "kA", "kO", "rho", "looovergrowth")
    save(list = savelist, file = paste(odeovergrowthoutputname, ".RData", sep = ""))
}

if(readodefrequencyresults == TRUE){
    fnames <- c(paste(odefrequencyoutputname, "1.csv", sep = ""), paste(odefrequencyoutputname, "2.csv", sep = ""), paste(odefrequencyoutputname, "3.csv", sep = ""), paste(odefrequencyoutputname, "4.csv", sep = ""))
    odefrequencydata <- read_stan_csv(csvfiles = fnames)
    chainsummary <- summary(odefrequencydata, pars = c("a0", "a1", "a2", "deltab0", "b1", "b2", "kA", "kO", "psi", "lp__"))$summary
    plot(rstan::traceplot(odefrequencydata, pars = c("a0", "a1", "a2", "deltab0", "b1", "b2", "kA", "kO", "psi", "lp__"), inc_warmup = FALSE))
    print(chainsummary)
    print("Summary of n_eff for dynamic parameters:")
    print(summary(chainsummary[, 9]))
    print("Summary of Rhat for dynamic parameters:")
    print(summary(chainsummary[, 10]))

    rhosummary <- summary(odefrequencydata, pars = c("rho"))$summary
    print("Summary of n_eff for rho:")
    print(summary(rhosummary[, 9]))
    print("Summary of Rhat for rho:")
    print(summary(rhosummary[, 10]))
    
    a0 <- extract(odefrequencydata, pars = "a0")[[1]]
    a1 <- extract(odefrequencydata, pars = "a1")[[1]]
    a2 <- extract(odefrequencydata, pars = "a2")[[1]]
    deltab0 <- extract(odefrequencydata, pars = "deltab0")[[1]]
    b1 <- extract(odefrequencydata, pars = "b1")[[1]]
    b2 <- extract(odefrequencydata, pars = "b2")[[1]]
    kA <- extract(odefrequencydata, pars = "kA")[[1]]
    kO <- extract(odefrequencydata, pars = "kO")[[1]]
    rho <- extract(odefrequencydata, pars = "rho")[[1]]
    psi <- extract(odefrequencydata, pars = "psi")[[1]]
    loofrequency <- loo(odefrequencydata, save_psis = TRUE)
    print(loofrequency)
    savelist <- c("a0", "a1", "a2", "deltab0", "b1", "b2", "kA", "kO", "psi", "rho", "loofrequency")
    save(list = savelist, file = paste(odefrequencyoutputname, ".RData", sep = ""))
}

if(readodefacilitationresults == TRUE){
    fnames <- c(paste(odefacilitationoutputname, "1.csv", sep = ""), paste(odefacilitationoutputname, "2.csv", sep = ""), paste(odefacilitationoutputname, "3.csv", sep = ""), paste(odefacilitationoutputname, "4.csv", sep = ""))
    odefacilitationdata <- read_stan_csv(csvfiles = fnames)
    chainsummary <- summary(odefacilitationdata, pars = c("a0", "a1", "a2", "deltab0", "b1", "b2", "kA", "kO", "m0", "lp__"))$summary
    plot(rstan::traceplot(odefacilitationdata, pars = c("a0", "a1", "a2", "deltab0", "b1", "b2", "kA", "kO", "m0", "lp__"), inc_warmup = FALSE))
    print(chainsummary)
    print("Summary of n_eff for dynamic parameters:")
    print(summary(chainsummary[, 9]))
    print("Summary of Rhat for dynamic parameters:")
    print(summary(chainsummary[, 10]))

    rhosummary <- summary(odefacilitationdata, pars = c("rho"))$summary
    print("Summary of n_eff for rho:")
    print(summary(rhosummary[, 9]))
    print("Summary of Rhat for rho:")
    print(summary(rhosummary[, 10]))
    
    a0 <- extract(odefacilitationdata, pars = "a0")[[1]]
    a1 <- extract(odefacilitationdata, pars = "a1")[[1]]
    a2 <- extract(odefacilitationdata, pars = "a2")[[1]]
    deltab0 <- extract(odefacilitationdata, pars = "deltab0")[[1]]
    b1 <- extract(odefacilitationdata, pars = "b1")[[1]]
    b2 <- extract(odefacilitationdata, pars = "b2")[[1]]
    kA <- extract(odefacilitationdata, pars = "kA")[[1]]
    kO <- extract(odefacilitationdata, pars = "kO")[[1]]
    rho <- extract(odefacilitationdata, pars = "rho")[[1]]
    m0 <- extract(odefacilitationdata, pars = "m0")[[1]]
    loofacilitation <- loo(odefacilitationdata, save_psis = TRUE)
    print(loofacilitation)
    savelist <- c("a0", "a1", "a2", "deltab0", "b1", "b2", "kA", "kO", "m0", "rho", "loofacilitation")
    save(list = savelist, file = paste(odefacilitationoutputname, ".RData", sep = ""))
}

if(readodeprotectionresults == TRUE){
    fnames <- c(paste(odeprotectionoutputname, "1.csv", sep = ""), paste(odeprotectionoutputname, "2.csv", sep = ""), paste(odeprotectionoutputname, "3.csv", sep = ""), paste(odeprotectionoutputname, "4.csv", sep = ""))
    odeprotectiondata <- read_stan_csv(csvfiles = fnames)
    chainsummary <- summary(odeprotectiondata, pars = c("a0", "a1", "a2", "deltab0", "b1", "b2", "kA", "kO", "m2", "lp__"))$summary
    plot(rstan::traceplot(odeprotectiondata, pars = c("a0", "a1", "a2", "deltab0", "b1", "b2", "kA", "kO", "m2", "lp__"), inc_warmup = FALSE))
    print(chainsummary)
    print("Summary of n_eff for dynamic parameters:")
    print(summary(chainsummary[, 9]))
    print("Summary of Rhat for dynamic parameters:")
    print(summary(chainsummary[, 10]))

    rhosummary <- summary(odeprotectiondata, pars = c("rho"))$summary
    print("Summary of n_eff for rho:")
    print(summary(rhosummary[, 9]))
    print("Summary of Rhat for rho:")
    print(summary(rhosummary[, 10]))
    
    a0 <- extract(odeprotectiondata, pars = "a0")[[1]]
    a1 <- extract(odeprotectiondata, pars = "a1")[[1]]
    a2 <- extract(odeprotectiondata, pars = "a2")[[1]]
    deltab0 <- extract(odeprotectiondata, pars = "deltab0")[[1]]
    b1 <- extract(odeprotectiondata, pars = "b1")[[1]]
    b2 <- extract(odeprotectiondata, pars = "b2")[[1]]
    kA <- extract(odeprotectiondata, pars = "kA")[[1]]
    kO <- extract(odeprotectiondata, pars = "kO")[[1]]
    rho <- extract(odeprotectiondata, pars = "rho")[[1]]
    m2 <- extract(odeprotectiondata, pars = "m2")[[1]]
    looprotection <- loo(odeprotectiondata, save_psis = TRUE)
    print(looprotection)
    savelist <- c("a0", "a1", "a2", "deltab0", "b1", "b2", "kA", "kO", "m2", "rho", "looprotection")
    save(list = savelist, file = paste(odeprotectionoutputname, ".RData", sep = ""))
}

if(readodegrowthfacilresults == TRUE){
    fnames <- c(paste(odegrowthfaciloutputname, "1.csv", sep = ""), paste(odegrowthfaciloutputname, "2.csv", sep = ""), paste(odegrowthfaciloutputname, "3.csv", sep = ""), paste(odegrowthfaciloutputname, "4.csv", sep = ""))
    odegrowthfacildata <- read_stan_csv(csvfiles = fnames)
    chainsummary <- summary(odegrowthfacildata, pars = c("a0", "a1", "a2", "deltab0", "b1", "b2", "kA", "kO", "m1", "lp__"))$summary
    plot(rstan::traceplot(odegrowthfacildata, pars = c("a0", "a1", "a2", "deltab0", "b1", "b2", "kA", "kO", "m1", "lp__"), inc_warmup = FALSE))
    print(chainsummary)
    print("Summary of n_eff for dynamic parameters:")
    print(summary(chainsummary[, 9]))
    print("Summary of Rhat for dynamic parameters:")
    print(summary(chainsummary[, 10]))

    rhosummary <- summary(odegrowthfacildata, pars = c("rho"))$summary
    print("Summary of n_eff for rho:")
    print(summary(rhosummary[, 9]))
    print("Summary of Rhat for rho:")
    print(summary(rhosummary[, 10]))
    
    a0 <- extract(odegrowthfacildata, pars = "a0")[[1]]
    a1 <- extract(odegrowthfacildata, pars = "a1")[[1]]
    a2 <- extract(odegrowthfacildata, pars = "a2")[[1]]
    deltab0 <- extract(odegrowthfacildata, pars = "deltab0")[[1]]
    b1 <- extract(odegrowthfacildata, pars = "b1")[[1]]
    b2 <- extract(odegrowthfacildata, pars = "b2")[[1]]
    kA <- extract(odegrowthfacildata, pars = "kA")[[1]]
    kO <- extract(odegrowthfacildata, pars = "kO")[[1]]
    rho <- extract(odegrowthfacildata, pars = "rho")[[1]]
    m1 <- extract(odegrowthfacildata, pars = "m1")[[1]]
    loogrowthfacil <- loo(odegrowthfacildata, save_psis = TRUE)
    print(loogrowthfacil)
    savelist <- c("a0", "a1", "a2", "deltab0", "b1", "b2", "kA", "kO", "m1", "rho", "loogrowthfacil")
    save(list = savelist, file = paste(odegrowthfaciloutputname, ".RData", sep = ""))
}

if(loadoderesults == TRUE) {
    load(file = paste(odeoutputname, ".RData", sep = ""))
    load(file = ddinputname)
}

if(loadodefacilresults == TRUE) {
    load(file = paste(odefaciloutputname, ".RData", sep = ""))
    load(file = ddinputname)
}

if(loadodeFPFOresults == TRUE) {
    load(file = paste(odeFPFOoutputname, ".RData", sep = ""))
    load(file = ddinputname)
}

if(loadodeovergrowthresults == TRUE) {
    load(file = paste(odeovergrowthoutputname, ".RData", sep = ""))
    load(file = ddinputname)
}

if(loadodefrequencyresults == TRUE) {
    load(file = paste(odefrequencyoutputname, ".RData", sep = ""))
    load(file = ddinputname)
}

if(loadodefacilitationresults == TRUE) {
    load(file = paste(odefacilitationoutputname, ".RData", sep = ""))
    load(file = ddinputname)
}

if(loadodeprotectionresults == TRUE) {
    load(file = paste(odeprotectionoutputname, ".RData", sep = ""))
    load(file = ddinputname)
}

if(loadodegrowthfacilresults == TRUE) {
    load(file = paste(odegrowthfaciloutputname, ".RData", sep = ""))
    load(file = ddinputname)
}

if(plotodepriors == TRUE) {
    chooseodepriors(niter = 1e2, nweeks = 8, fname = "../latex/odepriors.pdf")
}

if(buildtypicalrho == TRUE) {
        typicalpanels <- c(7, 4, 3, 6, 9, 5) #panels with typical treatment application in order C1, C3, O1, O3, A1, A3
        dayzero <- as.Date("2019-07-30")
        tC1dates <- c(dayzero, as.Date(dynamicdata$dynamicdata$date[dynamicdata$dynamicdata$panel == typicalpanels[1]]))
        tC3dates <- c(dayzero, as.Date(dynamicdata$dynamicdata$date[dynamicdata$dynamicdata$panel == typicalpanels[2]]))
        tO1dates <- c(dayzero, as.Date(dynamicdata$dynamicdata$date[dynamicdata$dynamicdata$panel == typicalpanels[3]]))
        tO3dates <- c(dayzero, as.Date(dynamicdata$dynamicdata$date[dynamicdata$dynamicdata$panel == typicalpanels[4]]))
        tA1dates <- c(dayzero, as.Date(dynamicdata$dynamicdata$date[dynamicdata$dynamicdata$panel == typicalpanels[5]]))
        tA3dates <- c(dayzero, as.Date(dynamicdata$dynamicdata$date[dynamicdata$dynamicdata$panel == typicalpanels[6]]))
        savelist = c("typicalrhoset", "typicalpanels", "tC1dates", "tC3dates", "tO1dates", "tO3dates", "tA1dates", "tA3dates")
        if(usegrowthfacil == TRUE){
            typicalrhoset <- typicalrho(a0 = a0, a1 = a1, a2 = a2, deltab0 = deltab0, b1 = b1, b2 = b2, kA = kA, kO = kO, m1 = m1, niter = 1e4)
            save(list = savelist, file = paste(odegrowthfaciloutputname, "_typicalrho.RData", sep = ""))
        } else if(useprotection == TRUE){
            typicalrhoset <- typicalrho(a0 = a0, a1 = a1, a2 = a2, deltab0 = deltab0, b1 = b1, b2 = b2, kA = kA, kO = kO, m2 = m2, niter = 1e4)
            save(list = savelist, file = paste(odeprotectionoutputname, "_typicalrho.RData", sep = ""))
        } else if(usefacilitation == TRUE){
            typicalrhoset <- typicalrho(a0 = a0, a1 = a1, a2 = a2, deltab0 = deltab0, b1 = b1, b2 = b2, kA = kA, kO = kO, m0 = m0, niter = 1e4)
            save(list = savelist, file = paste(odefacilitationoutputname, "_typicalrho.RData", sep = ""))
        } else if(usefrequency == TRUE){
            typicalrhoset <- typicalrho(a0 = a0, a1 = a1, a2 = a2, deltab0 = deltab0, b1 = b1, b2 = b2, kA = kA, kO = kO, psi = psi, niter = 1e4)
            save(list = savelist, file = paste(odefrequencyoutputname, "_typicalrho.RData", sep = ""))
        } else if(useovergrowth == TRUE){
            typicalrhoset <- typicalrho(a0 = a0, a1 = a1, a2 = a2, deltab0 = deltab0, b1 = b1, b2 = b2, kA = kA, kO = kO, a1y1star = a1y1star, niter = 1e4)
            save(list = savelist, file = paste(odeovergrowthoutputname, "_typicalrho.RData", sep = ""))
        } else if(useFPFO == TRUE){
            typicalrhoset <- typicalrho(a0 = a0, a1 = a1, a2 = a2, deltab0 = deltab0, b1 = b1, b2 = b2, kA = kA, kO = kO, m0 = m0, m2 = m2, a1y1star = a1y1star, psi = psi, niter = 1e4)
            save(list = savelist, file = paste(odeFPFOoutputname, "_typicalrho.RData", sep = ""))
        } else if(usefacil == TRUE){
            typicalrhoset <- typicalrho(a0 = a0, a1 = a1, a2 = a2, deltab0 = deltab0, b1 = b1, b2 = b2, kA = kA, kO = kO, m0 = m0, m2 = m2, niter = 1e4)
        save(list = savelist, file = paste(odefaciloutputname, "_typicalrho.RData", sep = ""))
    } else {
        typicalrhoset <- typicalrho(a0 = a0, a1 = a1, a2 = a2, deltab0 = deltab0, b1 = b1, b2 = b2, kA = kA, kO = kO, niter = 1e4)
        save(list = savelist, file = paste(odeoutputname, "_typicalrho.RData", sep = ""))
    }
}

if(plotoderesults == TRUE) {
        if(usegrowthfacil == TRUE){
        load(file = paste(odegrowthfaciloutputname, "_typicalrho.RData", sep = ""))
        timeseriesplot(alldata = alldata, potentialcompetitors = c("Botrylloides.spp.", "Bugula.spp.", "Molgula.tubifera"), dayzero = as.Date("2019-07-30"), fname = "../latex/timeseries.pdf", rhoC1summary = typicalrhoset$rhoC1summary, rhoC3summary = typicalrhoset$rhoC3summary, rhoO1summary = typicalrhoset$rhoO1summary, rhoO3summary = typicalrhoset$rhoO3summary, rhoA1summary = typicalrhoset$rhoA1summary, rhoA3summary = typicalrhoset$rhoA3summary, tC1dates = tC1dates, tC3dates = tC3dates, tO1dates = tO1dates, tO3dates = tO3dates, tA1dates = tA1dates, tA3dates = tA3dates, showobservations = TRUE, showpredictions = FALSE)
        timeseriesplot(alldata = alldata, potentialcompetitors = c("Botrylloides.spp.", "Bugula.spp.", "Molgula.tubifera"), dayzero = as.Date("2019-07-30"), fname = "../latex/timeseries_odegrowthfacil.pdf", rhoC1summary = typicalrhoset$rhoC1summary, rhoC3summary = typicalrhoset$rhoC3summary, rhoO1summary = typicalrhoset$rhoO1summary, rhoO3summary = typicalrhoset$rhoO3summary, rhoA1summary = typicalrhoset$rhoA1summary, rhoA3summary = typicalrhoset$rhoA3summary, tC1dates = tC1dates, tC3dates = tC3dates, tO1dates = tO1dates, tO3dates = tO3dates, tA1dates = tA1dates, tA3dates = tA3dates, showobservations = FALSE, showpredictions = TRUE)
    } else if(usefacilitation == TRUE){
        load(file = paste(odefacilitationoutputname, "_typicalrho.RData", sep = ""))
        timeseriesplot(alldata = alldata, potentialcompetitors = c("Botrylloides.spp.", "Bugula.spp.", "Molgula.tubifera"), dayzero = as.Date("2019-07-30"), fname = "../latex/timeseries.pdf", rhoC1summary = typicalrhoset$rhoC1summary, rhoC3summary = typicalrhoset$rhoC3summary, rhoO1summary = typicalrhoset$rhoO1summary, rhoO3summary = typicalrhoset$rhoO3summary, rhoA1summary = typicalrhoset$rhoA1summary, rhoA3summary = typicalrhoset$rhoA3summary, tC1dates = tC1dates, tC3dates = tC3dates, tO1dates = tO1dates, tO3dates = tO3dates, tA1dates = tA1dates, tA3dates = tA3dates, showobservations = TRUE, showpredictions = FALSE)
        timeseriesplot(alldata = alldata, potentialcompetitors = c("Botrylloides.spp.", "Bugula.spp.", "Molgula.tubifera"), dayzero = as.Date("2019-07-30"), fname = "../latex/timeseries_odefacilitation.pdf", rhoC1summary = typicalrhoset$rhoC1summary, rhoC3summary = typicalrhoset$rhoC3summary, rhoO1summary = typicalrhoset$rhoO1summary, rhoO3summary = typicalrhoset$rhoO3summary, rhoA1summary = typicalrhoset$rhoA1summary, rhoA3summary = typicalrhoset$rhoA3summary, tC1dates = tC1dates, tC3dates = tC3dates, tO1dates = tO1dates, tO3dates = tO3dates, tA1dates = tA1dates, tA3dates = tA3dates, showobservations = FALSE, showpredictions = TRUE)
    } else if(usefrequency == TRUE){
            load(file = paste(odefrequencyoutputname, "_typicalrho.RData", sep = ""))
            timeseriesplot(alldata = alldata, potentialcompetitors = c("Botrylloides.spp.", "Bugula.spp.", "Molgula.tubifera"), dayzero = as.Date("2019-07-30"), fname = "../latex/timeseries.pdf", rhoC1summary = typicalrhoset$rhoC1summary, rhoC3summary = typicalrhoset$rhoC3summary, rhoO1summary = typicalrhoset$rhoO1summary, rhoO3summary = typicalrhoset$rhoO3summary, rhoA1summary = typicalrhoset$rhoA1summary, rhoA3summary = typicalrhoset$rhoA3summary, tC1dates = tC1dates, tC3dates = tC3dates, tO1dates = tO1dates, tO3dates = tO3dates, tA1dates = tA1dates, tA3dates = tA3dates, showobservations = TRUE, showpredictions = FALSE)
            timeseriesplot(alldata = alldata, potentialcompetitors = c("Botrylloides.spp.", "Bugula.spp.", "Molgula.tubifera"), dayzero = as.Date("2019-07-30"), fname = "../latex/timeseries_odefrequency.pdf", rhoC1summary = typicalrhoset$rhoC1summary, rhoC3summary = typicalrhoset$rhoC3summary, rhoO1summary = typicalrhoset$rhoO1summary, rhoO3summary = typicalrhoset$rhoO3summary, rhoA1summary = typicalrhoset$rhoA1summary, rhoA3summary = typicalrhoset$rhoA3summary, tC1dates = tC1dates, tC3dates = tC3dates, tO1dates = tO1dates, tO3dates = tO3dates, tA1dates = tA1dates, tA3dates = tA3dates, showobservations = FALSE, showpredictions = TRUE)
        } else if(useovergrowth == TRUE){
            load(file = paste(odeovergrowthoutputname, "_typicalrho.RData", sep = ""))
            timeseriesplot(alldata = alldata, potentialcompetitors = c("Botrylloides.spp.", "Bugula.spp.", "Molgula.tubifera"), dayzero = as.Date("2019-07-30"), fname = "../latex/timeseries.pdf", rhoC1summary = typicalrhoset$rhoC1summary, rhoC3summary = typicalrhoset$rhoC3summary, rhoO1summary = typicalrhoset$rhoO1summary, rhoO3summary = typicalrhoset$rhoO3summary, rhoA1summary = typicalrhoset$rhoA1summary, rhoA3summary = typicalrhoset$rhoA3summary, tC1dates = tC1dates, tC3dates = tC3dates, tO1dates = tO1dates, tO3dates = tO3dates, tA1dates = tA1dates, tA3dates = tA3dates, showobservations = TRUE, showpredictions = FALSE)
             timeseriesplot(alldata = alldata, potentialcompetitors = c("Botrylloides.spp.", "Bugula.spp.", "Molgula.tubifera"), dayzero = as.Date("2019-07-30"), fname = "../latex/timeseries_odeovergrowth.pdf", rhoC1summary = typicalrhoset$rhoC1summary, rhoC3summary = typicalrhoset$rhoC3summary, rhoO1summary = typicalrhoset$rhoO1summary, rhoO3summary = typicalrhoset$rhoO3summary, rhoA1summary = typicalrhoset$rhoA1summary, rhoA3summary = typicalrhoset$rhoA3summary, tC1dates = tC1dates, tC3dates = tC3dates, tO1dates = tO1dates, tO3dates = tO3dates, tA1dates = tA1dates, tA3dates = tA3dates, showobservations = TRUE, showpredictions = TRUE, predlwd = 2) #for best model (in main text), overlay predictions on plot of observations
        } else if(useFPFO == TRUE){
            load(file = paste(odeFPFOoutputname, "_typicalrho.RData", sep = ""))
            timeseriesplot(alldata = alldata, potentialcompetitors = c("Botrylloides.spp.", "Bugula.spp.", "Molgula.tubifera"), dayzero = as.Date("2019-07-30"), fname = "../latex/timeseries.pdf", rhoC1summary = typicalrhoset$rhoC1summary, rhoC3summary = typicalrhoset$rhoC3summary, rhoO1summary = typicalrhoset$rhoO1summary, rhoO3summary = typicalrhoset$rhoO3summary, rhoA1summary = typicalrhoset$rhoA1summary, rhoA3summary = typicalrhoset$rhoA3summary, tC1dates = tC1dates, tC3dates = tC3dates, tO1dates = tO1dates, tO3dates = tO3dates, tA1dates = tA1dates, tA3dates = tA3dates, showobservations = TRUE, showpredictions = FALSE)
            timeseriesplot(alldata = alldata, potentialcompetitors = c("Botrylloides.spp.", "Bugula.spp.", "Molgula.tubifera"), dayzero = as.Date("2019-07-30"), fname = "../latex/timeseries_odeFPFO.pdf", rhoC1summary = typicalrhoset$rhoC1summary, rhoC3summary = typicalrhoset$rhoC3summary, rhoO1summary = typicalrhoset$rhoO1summary, rhoO3summary = typicalrhoset$rhoO3summary, rhoA1summary = typicalrhoset$rhoA1summary, rhoA3summary = typicalrhoset$rhoA3summary, tC1dates = tC1dates, tC3dates = tC3dates, tO1dates = tO1dates, tO3dates = tO3dates, tA1dates = tA1dates, tA3dates = tA3dates, showobservations = FALSE, showpredictions = TRUE)
        } else if(usefacil == TRUE){
            load(file = paste(odefaciloutputname, "_typicalrho.RData", sep = ""))
            timeseriesplot(alldata = alldata, potentialcompetitors = c("Botrylloides.spp.", "Bugula.spp.", "Molgula.tubifera"), dayzero = as.Date("2019-07-30"), fname = "../latex/timeseries.pdf", rhoC1summary = typicalrhoset$rhoC1summary, rhoC3summary = typicalrhoset$rhoC3summary, rhoO1summary = typicalrhoset$rhoO1summary, rhoO3summary = typicalrhoset$rhoO3summary, rhoA1summary = typicalrhoset$rhoA1summary, rhoA3summary = typicalrhoset$rhoA3summary, tC1dates = tC1dates, tC3dates = tC3dates, tO1dates = tO1dates, tO3dates = tO3dates, tA1dates = tA1dates, tA3dates = tA3dates, showobservations = TRUE, showpredictions = FALSE)
            timeseriesplot(alldata = alldata, potentialcompetitors = c("Botrylloides.spp.", "Bugula.spp.", "Molgula.tubifera"), dayzero = as.Date("2019-07-30"), fname = "../latex/timeseries_odefacil.pdf", rhoC1summary = typicalrhoset$rhoC1summary, rhoC3summary = typicalrhoset$rhoC3summary, rhoO1summary = typicalrhoset$rhoO1summary, rhoO3summary = typicalrhoset$rhoO3summary, rhoA1summary = typicalrhoset$rhoA1summary, rhoA3summary = typicalrhoset$rhoA3summary, tC1dates = tC1dates, tC3dates = tC3dates, tO1dates = tO1dates, tO3dates = tO3dates, tA1dates = tA1dates, tA3dates = tA3dates, showobservations = FALSE, showpredictions = TRUE)
        } else {
            load(file = paste(odeoutputname, "_typicalrho.RData", sep = ""))
            timeseriesplot(alldata = alldata, potentialcompetitors = c("Botrylloides.spp.", "Bugula.spp.", "Molgula.tubifera"), dayzero = as.Date("2019-07-30"), fname = "../latex/timeseries.pdf", rhoC1summary = typicalrhoset$rhoC1summary, rhoC3summary = typicalrhoset$rhoC3summary, rhoO1summary = typicalrhoset$rhoO1summary, rhoO3summary = typicalrhoset$rhoO3summary, rhoA1summary = typicalrhoset$rhoA1summary, rhoA3summary = typicalrhoset$rhoA3summary, tC1dates = tC1dates, tC3dates = tC3dates, tO1dates = tO1dates, tO3dates = tO3dates, tA1dates = tA1dates, tA3dates = tA3dates, showobservations = TRUE, showpredictions = FALSE)
            timeseriesplot(alldata = alldata, potentialcompetitors = c("Botrylloides.spp.", "Bugula.spp.", "Molgula.tubifera"), dayzero = as.Date("2019-07-30"), fname = "../latex/timeseries_ode.pdf", rhoC1summary = typicalrhoset$rhoC1summary, rhoC3summary = typicalrhoset$rhoC3summary, rhoO1summary = typicalrhoset$rhoO1summary, rhoO3summary = typicalrhoset$rhoO3summary, rhoA1summary = typicalrhoset$rhoA1summary, rhoA3summary = typicalrhoset$rhoA3summary, tC1dates = tC1dates, tC3dates = tC3dates, tO1dates = tO1dates, tO3dates = tO3dates, tA1dates = tA1dates, tA3dates = tA3dates, showobservations = FALSE, showpredictions = TRUE)
            plotodeparameters(a0 = a0, a1 = a1, a2 = a2, deltab0 = deltab0, b1 = b1, b2 = b2, kA = kA, kO = kO, fname = "../latex/odeparameters.pdf")
        }
}

if(checkAremoval == TRUE){
    load(file = ddinputname)
    plotAremovaleffects(dynamicdata = dynamicdata$dynamicdata, fname = "../latex/checkAremovaleffects.pdf")
    plotAremovalproportion(dynamicdata = dynamicdata$dynamicdata, fname = "../latex/checkAremovalproportion.pdf", odeoutputname = odeoutputname, odefacilitationoutputname = odefacilitationoutputname, odeprotectionoutputname = odeprotectionoutputname, odeovergrowthoutputname = odeovergrowthoutputname, odegrowthfaciloutputname = odegrowthfaciloutputname)
}

if(compareodemodels == TRUE){
    odeloo(odeoutputname = odeoutputname, odefacilitationoutputname = odefacilitationoutputname, odeprotectionoutputname = odeprotectionoutputname, odeovergrowthoutputname = odeovergrowthoutputname, odegrowthfaciloutputname = odegrowthfaciloutputname)
}

if(checkcmatrix == TRUE){
    checkcommunitymatrix(x = 0.75, y1star = 0.1, odeoutputname = odeoutputname, odefacilitationoutputname = odefacilitationoutputname, odegrowthfaciloutputname = odegrowthfaciloutputname, odeovergrowthoutputname = odeovergrowthoutputname, odeprotectionoutputname = odeprotectionoutputname)
}

if(printODEtable == TRUE){
    printODEparameters(odeoutputname = odeoutputname, odefacilitationoutputname = odefacilitationoutputname, odegrowthfaciloutputname = odegrowthfaciloutputname, odeovergrowthoutputname = odeovergrowthoutputname, odeprotectionoutputname = odeprotectionoutputname, fname = "../latex/ODEparameters.tex")
}

if(analyzeenvironmentaldata == TRUE){
    envdata <- read.table(file = envdataname, header = TRUE, sep = "\t")
    envanalyses(envdata = envdata, envfigname = "../latex/envdata.pdf")
}

if(doSBCODE == TRUE){
    load(file = ddinputname)
        sbcodeoutput <- SBCODE(odestanfile = odestanfile, dynamicdata = dynamicdata, M = 1, iter = 50) #test with few draws from prior
#    sbcodeoutput <- SBCODE(odestanfile = odestanfile, dynamicdata = dynamicdata, M = 500, iter = 5000)
    save(sbcodeoutput, file = "../../oderesults/SBCODE.RData")
}

if(setupcondorSBCODE == TRUE){
    load(file = ddinputname)
    odemodel <- stan_model(odestanfile)
    save(list = c("odemodel", "dynamicdata"), file = SBCODEdatafile, version = 2) #need version = 2 to run with old R version on Condor
}

if(posteriorpredictiveODE == TRUE){
    niter <- 1e2
    simdata <- simulatefromode(dynamicdata = dynamicdata, rho = rho, niter = niter)
    plotposteriorode(dynamicdata = dynamicdata, simdata = simdata, alldata = alldata, niter = niter, fname = "../latex/posteriorpredictiveovergrowth.pdf")
}

if(plotODEeffects == TRUE){
    plotallAureliaeffects(odeoutputname = odeoutputname, odeovergrowthoutputname = odeovergrowthoutputname, odeprotectionoutputname = odeprotectionoutputname, odefacilitationoutputname = odefacilitationoutputname, odegrowthfaciloutputname = odegrowthfaciloutputname, grid = 5, gridstep = 0.1, rangelim = 100, fname = "../latex/Aureliaeffects.pdf")
}

if(setupsimODEdata == TRUE){
    rhofrommean <- simfrommeanparameters(dynamicdata = dynamicdata, a0 = a0, a1 = a1, a2 = a2, deltab0 = deltab0, b1 = b1, b2 = b2, kA = kA, kO = kO, a1y1star = a1y1star, niter = 10, fname = simodename) #only implemented for overgrowth model
}

if (checksimODEdata == TRUE){#only implemented for overgrowth model
    if(readrawsimoutput == TRUE){#read the raw data files?
        readrawsimdata(fname = simodename, outname = simodeprocessedname, a0 = a0, a1 = a1, a2 = a2, deltab0 = deltab0, b1 = b1, b2 = b2, kA = kA, kO = kO, a1y1star = a1y1star)
    }
    load(simodeprocessedname)
    priorsds <- c(0.05, 0.05, 1, 1, 0.05, 0.05, 0.02, 0.02, 0.2, 0.2, 0.05, 0.05, 1, 1) #prior st devs in order a0, a1, a2, deltab0, b1, b2, a1y1star (each at two depths)
    signs <- c(1, 1, 1, 1, -1, -1, 1, 1 ,1, 1, -1, -1, 1, 1) #signs for half-normals, order as above
    priorhalfnorms <- cbind(priorsds, signs)
    priorbetaparms <- array(c(2, 2, 2, 2), dim = c(2, 2)) #beta(2, 2) prior for each of kA and kO: each row is for one parameter
    plotsimdens(truetheta = truetheta, densities = densities, priorhalfnorms = priorhalfnorms, priorbetaparms = priorbetaparms, inHPDs = inHPDs, fname = "../latex/simODEperformance.pdf")
}
```
