## Supplementary material for "Negative and positive interspecific interactions involving jellyfish polyps in marine sessile communities": Data and code: aureliafuncs.R

```
#extract data from a single point-counting csv file exported from JMicroVision
#Arguments:
#dirname: path to the directory containing the file
#fname: name of the csv file (note these are actually semicolon-separated). Format example: Panel1_Pre_6Aug2019.csv (we split at the _ to extract panel number, pre/post, date)
#Value: a list containing:
#csvdata: a data frame containing the variables Class and Whole.Image (number of points out of 100 occupied by each class)
#panelmetadata: a list containing photodate, prepost, panelnumber, from getpanelmetadata()
readonecsv <- function(dirname, fname){
    csvdata <- read.table(paste(dirname, fname, sep = "/"), header = TRUE, sep = ";")
    if(any(names(csvdata) == "Whole.Image")){#we have "Classes" format
        csvdata <- csvdata[, c("Class", "Whole.Image")]
    } else {#we have "Graphics" format
        csvdata <- as.data.frame(table(csvdata$Class))
        names(csvdata)  <- c("Class", "Whole.Image")
    }
    panelmetadata <- getpanelmetadata(fname)
    return(list(csvdata = csvdata, panelmetadata = panelmetadata))
}

#extract panel info from file name
#Arguments:
#fname: name of a .csv file, format Panel#_[Pre or Post]_[date e.g. 6Aug2019].csv
#Value:
#list containing:
#photodate: date on which photo was taken
#prepost: Pre or Post
#panelnumber: integer, panel number
getpanelmetadata <- function(fname){
    panelinfo <- strsplit(fname, "_")
    photodate <- as.Date(strsplit(panelinfo[[1]][3], "\\.")[[1]][1], "%d%B%Y")
    prepost <- panelinfo[[1]][2]
    panelnumber <- as.integer(strsplit(panelinfo[[1]][1], "Panel")[[1]][2])
    return(list(photodate = photodate, prepost = prepost, panelnumber = panelnumber))
}

#add any missing levels to csv data from a single photo
#Arguments:
#csvdata: first item in output list from readonecsv, a data frame with variables Class, Whole.Image
#classlist: list of all the classes we want
#Value:
#data frame with same structure as csvdata, but with new rows appended, with the missing levels and count of zero, all in alphabetical order
addlevels <- function(csvdata, classlist){
    newclasses <- setdiff(classlist, levels(csvdata$Class))
    alllevels <-  unique(c(levels(csvdata$Class), classlist))
    alllevels <- alllevels[order(alllevels)]
    csvdata$Class <- factor(csvdata$Class, levels = alllevels)
    nnew <- length(newclasses)
    newrows <- data.frame(Class = newclasses, Whole.Image = rep(0, nnew))
    csvdata <- rbind(csvdata, newrows)
    return(csvdata)
}

#extract one row of counts from output of readonecsv
#Arguments:
#csvdata: first item in output list from readonecsv, a data frame with variables Class, Whole.Image
#classlist: list of all the classes we want
#Value:
#vector containing count for each class in alphabetical order
getcounts <- function(csvdata, classlist){
    fulllevels <- addlevels(csvdata, classlist)
    fulllevels <- fulllevels[order(fulllevels$Class), ]
    return(as.integer(t(fulllevels)[2, ]))
}

#add treatment information to master data frame
#Arguments:
#alldata: data frame containing panel, date, prepost and count for each class
#fname: name of tab-separated text file containing Block, Depthm, Panel, Treatment
#Value:
#alldata with block, depthm, treatment added for each panel.
addtreatments <- function(alldata, fname){
    treatments <- read.table(file = fname, header = TRUE, sep = "\t")
    nrows <- dim(alldata)[1]
    block <- numeric(nrows)
    depthm <- numeric(nrows)
    treatment <- factor(rep(NA, nrows), levels = c("A", "O", "C"))
    alldata <- cbind(block, depthm, treatment, alldata)
    for(i in 1:nrows){
        mypanel <- alldata$panel[i]
        mytreatment <- which(treatments$Panel == mypanel)
        alldata$block[i] <- treatments$Block[mytreatment]
        alldata$depthm[i] <- treatments$Depthm[mytreatment]
        alldata$treatment[i] <- treatments$Treatment[mytreatment]
    }
    return(alldata)
}

#extract all the csv files in a given list of directories
#Arguments: a list of all the directories containing JMicroVision csv files
#Value:
#A data frame alldata with variables panel (integer panel number), date, prepost (levels Pre, Post, meaning before or after treatment application each week), and then counts of each class, in alphabetical order 
#Note: the raw data contain some files that are called Polyp or PostPolyp (polyp counts rather than point counts) rather than Pre or Post. These will be read and will give warnings, but won't appear as rows in alldata
readdirectories <- function(dirnames, treatmentdata){
    classlist <- character(0)
    nrows <- 0
    for(dirname in dirnames){#first pass: construct list of taxa
        files <- list.files(dirname, pattern = "\\.csv")
        for(file in files){
            paneldata <- readonecsv(dirname = dirname, fname = file)
            classlist <- unique(c(classlist, levels(paneldata$csvdata$Class)))
            nrows <- nrows + 1
        }
    }
    classlist <- classlist[order(classlist)]
    nclasses <- length(classlist)
    alldata <- data.frame(panel = numeric(nrows), date = integer(nrows), prepost = factor(rep(NA, nrows), levels = c("Pre", "Post")))
    counts <- array(dim = c(nrows, nclasses))
    colnames(counts) <- gsub(" ", "\\.", classlist)
    class(alldata$date) <- "Date"
    row <- 1
    for(dirname in dirnames){#second pass: assemble data frame
        files <- list.files(dirname, pattern = "\\.csv")
        for(file in files){
            paneldata <- readonecsv(dirname = dirname, fname = file)
            alldata[row, "panel"] <- paneldata$panelmetadata$panelnumber
            alldata[row, "date"] <- paneldata$panelmetadata$photodate
            alldata[row, "prepost"] <- paneldata$panelmetadata$prepost
            counts[row, ]  <- getcounts(paneldata$csvdata, classlist)
            row <- row + 1
        }
    }
    alldata <- cbind(alldata, counts)
    alldata <- alldata[!is.na(alldata$prepost), ] #remove rows from csv files that were Polyp or PostPolyp rather than Pre or Post
    alldata <- addtreatments(alldata = alldata, fname = treatmentdata)
    alldata <- alldata[with(alldata, order(date, prepost, panel)), ]
    return(alldata)
}

#correct some typos in taxon names, merge some taxa that we couldn't distinguish reliably, remove taxa we never saw
fixtyposandmerge <- function(alldata){
    alldata$Botrylloides.spp. <- alldata$Botrylloides + alldata$Botrylloides.leachii + alldata$Botrylloides.spp. + alldata$Botrylloides.violaceus
    drops <- c("Botrylloides", "Botrylloides.leachii", "Botrylloides.violaceus")
    alldata  <- alldata[, !names(alldata) %in% drops]    
    colnames(alldata)[which(colnames(alldata) == "Botryllus.schlossoi")] <- "Botryllus.schlosseri"
    colnames(alldata)[which(colnames(alldata) == "Botryllus.schlossoion.competitor")] <- "Botryllus.schlosseri.on.competitor"
    maxabund <- colSums(alldata[, -c(1:6)])
    alldata <- alldata[, c(1:6, which(maxabund > 0) + 6)]
    return(alldata)
}

#assemble a data frame for all panels, all weeks
#Arguments: none
#Value: no return value, but tab-separated text file alldata.txt produced 
compiledata <- function(){
    dirnames <- c("../../photos/Panels_6Aug2019", "../../photos/Panels_13Aug2019", "../../photos/Panels_20Aug2019", "../../photos/Panels27Aug2019", "../../photos/Panels3Sep2019", "../../photos/Panels10Sep2019", "../../photos/Panels17Sep2019", "../../photos/Panels24Sep2019")
    alldata <- readdirectories(dirnames = dirnames, treatmentdata = "../data/treatments.txt")
    alldata <- fixtyposandmerge(alldata)
    write.table(alldata, file = "../data/alldata.txt", sep = "\t", quote = FALSE, row.names = FALSE)
}

#ODE model for polyps, potential competitors, free space (no polyps on potential competitors) in nondimensionalized form
#Arguments:
#t is dimensionless time
#state is xstar, y1star (proportion of space filled by potential competitors, proportion of space filled by polyps)
#parameters is dimensionless parameters Pi1, ... , Pi5
#Value: rates of change of xstar, y1star wrt dimensionless time
#Example:
#parameters <- c(Pi1 = 1, Pi2 = 2, Pi3 = -1, Pi4 = 1, Pi5 = -1)
#times <- seq(from = 0, to = 1, by = 0.001)
#state <- c(xstar = 0, y1star = 0)
#aureliasol <- ode(y = state, times = times, func = aurelia2dODE, parms = parameters)
aurelia2dODE <- function(t, state, parameters){
    with(as.list(c(state, parameters)),{
        freespace  <- 1 - xstar - y1star
        dxstar <- Pi1 * freespace + Pi2 * xstar * freespace + Pi3 * xstar
        dy1star <- freespace + Pi4 * y1star * freespace + Pi5 * y1star
        list(c(dxstar, dy1star))
    })
}

#ODE model for polyps, potential competitors, free space (no polyps on potential competitors) in "dimensional" form : parameters are all scaled by sampling interval
#Arguments:
#t is time in units of sampling interval (e.g. one week)
#state is xstar, y1star (proportion of space filled by potential competitors, proportion of space filled by polyps)
#parameters is a0, a1, a2, m0, m2, deltab0, b1, b2, psi, a1y1star, m1
#Value: rates of change of xstar, y1star wrt time in sampling units
aurelia2dODEdim <- function(t, state, parameters){
    with(as.list(c(state, parameters)),{
        freespace  <- 1 - xstar - y1star
        dxstar <- (a0 + m0 * y1star) * freespace +  (a1 + m1 * y1star) * xstar * freespace^(1 - psi) * y1star^psi + a1y1star * xstar * y1star + (a2 * exp(-m2 * y1star)) * xstar
        dy1star <- deltab0 * freespace + b1 * y1star * freespace - a1y1star * xstar * y1star + b2 * y1star
        list(c(dxstar, dy1star))
    })
}

#simulate 2-dimensionsal ODE model for aurelia and potential competitors, with treatment applied at regular time intervals
#Arguments:
#treatment: vector of proportion remaining after treatment applied at end of each interval, in order (potential competitors, A. aurita), e.g. c(0.5, 1) to remove half of potential competitors
#Value: list containing:
#alltimes: vector of times. End of each time interval is repeated, because we have before and after treatment values at this time.
#aureliacomp: composition from numerical solution of ODEs: potential competitors, A. aurita, free space
#TO DO: this version has the parameters hard coded, assumes 8 time intervals, and hasn't properly dealt with redimensionalizing time
aurelia2dwithtreatment <- function(treatment){
    parameters <- c(Pi1 = 1, Pi2 = 2, Pi3 = -1, Pi4 = 1, Pi5 = -1)
    state <- c(xstar = 0, y1star = 0)
    tend <- 1 #time units are weeks
    times <- seq(from = 0, to = tend, by = 0.1)
    nweeks <- 8
    ntimes <- length(times)
    alltimes <- numeric(ntimes * nweeks)
    aurelia <- array(dim = c(ntimes * nweeks, 3))
    for(week in 1:nweeks){
        aureliasol <- ode(y = state, times = times, func = aurelia2dODE, parms = parameters)
        state <- aureliasol[dim(aureliasol)[1], 2:3] * treatment
        aurelia[(1 + (week - 1) * ntimes):(week * ntimes), ] <- aureliasol
        alltimes[(1 + (week - 1) * ntimes):(week * ntimes)] <- aureliasol[, 1] + week - 1
    }
    aureliacomp <- acomp(cbind(aurelia[, 2:3], 1 - aurelia[, 2] - aurelia[, 3]))
    names(aureliacomp) <- c("potential competitors", "A. aurita", "free space")
    return(list(alltimes = alltimes, aureliacomp = aureliacomp))
}

#simulate three treatments in 2-dimensional Aurelia model: control, remove half Aurelia once a week, removfe half potential competitors once a week
#Arguments: none
#Value: numerical solutions to the ODEs plotted in the simplex for each treatment
aureliatreatmentplot <- function(){
    treatmentC <- aurelia2dwithtreatment(treatment = c(1, 1))
    treatmentA <- aurelia2dwithtreatment(treatment = c(1, 0.5))
    treatmentO <- aurelia2dwithtreatment(treatment = c(0.5, 1))
    mycolors <- brewer.pal(3, 'Dark2')
    plot(treatmentC$aureliacomp, col = mycolors[1], type = "l")
    plot(treatmentA$aureliacomp, col = mycolors[2], type = "l", add = TRUE)
    plot(treatmentO$aureliacomp, col = mycolors[3], type = "l", add = TRUE)
}

#Rough plot of last week's data
#Currently just adds half detection limit to zeros. Different symbols for 1 m and 3 m. Different colours for treatments. Optionally, centre.
#e.g. lastweekplot(alldata = aurelia, date = "2019-09-24", preorpost = "Pre", centre = FALSE)
lastweekplot <- function(alldata, date, preorpost, centre = FALSE){
    lastweek <- alldata[alldata$date == date & alldata$prepost == preorpost, ]
    lastweek$potentialcompetitors <- lastweek$Ascidiella.aspersa + lastweek$Botrylloides.spp. + lastweek$Botryllus.schlosseri + lastweek$Bugula.spp. + lastweek$Molgula.tubifera
    lastweekcomp <- cbind(lastweek$Bare.Panel, lastweek$Aurelia.aurita.on.panel, lastweek$potentialcompetitors)
    colnames(lastweekcomp)  <- c("Bare panel", "A. aurita", "Potential competitors")
    lastweekcomp[lastweekcomp == 0] <- 0.5 #rough here: adding half detection limit to plot
    lastweekcomp <- acomp(lastweekcomp)
    if(centre == TRUE) lastweekcomp <- lastweekcomp - mean(lastweekcomp) #rough: centre by sample mean
    mycolors <- brewer.pal(3, "Dark2")
    mypch <- c(1, 16)
    plot(acomp(lastweekcomp[lastweek$treatment == "C", ]), pch = mypch[factor(lastweek$depthm)], col = adjustcolor(mycolors[1], 0.7)) #green
    plot(acomp(lastweekcomp[lastweek$treatment == "A", ]), pch = mypch[factor(lastweek$depthm)], col = adjustcolor(mycolors[2], 0.7), add = TRUE) #orange
    plot(acomp(lastweekcomp[lastweek$treatment == "O", ]), pch = mypch[factor(lastweek$depthm)], col = adjustcolor(mycolors[3], 0.7), add = TRUE) #blue
}

#build a set of dummy variables representing contrasts between depths and scrape treatments
#Arguments: depth (1 or 3) and treatment (O, C or A)
#Value: a vector containing values of dummy variables:
#d1: depth contrast
#d2: scrape contrast between O and A
#d3: scrape contrast between C and O & A
#d4: interaction contrast: product of d1 and d2
#d5: interaction contrast: product of d1 and d3
buildcontrastbasis <- function(depth, treatment){
    if(depth == 1) d1  <- -1 else if (depth == 3) d1 <- 1 else stop("depth must be 1 or 3")
    d2 <- switch(as.character(treatment), "O" =  1,
                 "C" = 0,
                 "A" = -1,
                 stop("treatment must be O, C or A"))
    d3 <- switch(as.character(treatment), "O" =  -1 / 2,
                 "C" = 1,
                 "A" = -1 / 2,
                 stop("treatment must be O, C or A"))
    d4 <- d1 * d2
    d5 <- d1 * d3
    return(list(d1 = d1, d2 = d2, d3 = d3, d4 = d4, d5 = d5))
}

#set up for manova on data from a single (e.g. final) week
#Arguments:
#alldata: data frame from alldata.txt
#date: the date we want to analyse
#preorpost: use the pre- or post-scrape data from this week?
#taxa: names of the taxa to include in the response vector
#tVinv: generalized inverse of transpose of basis matrix for ilr coordinates
#standatafilename: name of file to which input data for stan will be written
#Value:
#writes to standatafilename using stan_rdump and returns list containing
#d1 to d5: vectors of values of dummy variables representing treatment contrasts for each panel
#counts: array of counts by panel for the taxa we include
#blockindex: vector labelling panels by block
aureliamanovasetup <- function(alldata, date, preorpost, taxa, tVinv, standatafilename){
    ntaxa  <- length(taxa)
    lastweek <- alldata[alldata$date == date & alldata$prepost == preorpost, ]
    npanels <- dim(lastweek)[1]
    d1 <- numeric(npanels)
    d2 <- d1
    d3 <- d1
    d4 <- d1
    d5 <- d1
    for(i in 1:npanels){
        contrasts <- buildcontrastbasis(depth = alldata$depthm[i], treatment = alldata$treatment[i])
        d1[i] <- contrasts$d1
        d2[i] <- contrasts$d2
        d3[i] <- contrasts$d3
        d4[i] <- contrasts$d4
        d5[i] <- contrasts$d5
    }
    counts  <- lastweek[, taxa]
    blockindex = lastweek$block
    nblocks <- length(unique(lastweek$block))
    paneldata <- c("d1", "d2", "d3", "d4", "d5", "counts", "tVinv", "ntaxa", "npanels", "nblocks", "blockindex")
    stan_rdump(list = paneldata, file = standatafilename, quiet = FALSE, append = FALSE)
    return(list(d1 = d1, d2 = d2, d3 = d3, d4 = d4, d5 = d5, counts = counts, blockindex = blockindex))
}

#create a suitable basis matrix
#Arguments:
#D: number of taxa
#choice: default is the default basis from ilrBase.
#B1 has log contrasts: Aurelia vs bare panel, potential competitors vs Aurelia and bare panel, ascidians vs bryozoans, colonial vs solitary bryozoans.
#Value:
#a D x (D - 1) ilr basis matrix
#Notes: ginv(t(V)) = V provided we have an orthonormal basis, so can use this as tVinv (generalized inverse of transpose of basis matrix) where needed
makeV <- function(D, choice = "default"){
    if(choice == "default"){
        V <- ilrBase(D = D)
    } else if (choice == "B1"){
        V <- matrix(nrow = 5, ncol = 4, c(1 / sqrt(2), -1 / sqrt(2), 0, 0, 0, -sqrt(3 / 10), -sqrt(3 / 10), sqrt(2 / 15), sqrt(2 / 15), sqrt(2 / 15), 0, 0, 1 / sqrt(6), -2 / sqrt(6), 1 / sqrt(6), 0, 0, 1 / sqrt(2), 0, -1 / sqrt(2)))
    } else {
        stop("This choice of basis matrix not implemented")
    }
return(V)
}

#load data on depth effects extracted from Chong and Spencer 2018 and extract a suitable prior based on these
#Arguments:
#Chongfilename: name of .RData file containing coefficients of intercept and centred and scaled depth and depth^2 (not orthogonal) from Chong and Spencer 2018
#mytaxa: the columns in back-transformed parameters from Chong corresponding to the subcomposition we want
#V: the ilr basis matrix we want for our subcomposition
#depth1 and depth3: the depths for our two treatments (defaults to 1 m and 3 m)
#figname: name of figure showing our chosen prior and the posterior from Chong 18
#Value: prints out mean and covariance matrix of chosen prior, creates pdf figure fname with kernel densities for marginal distributions and pairwise scatter plots of ilr components
makedepthprior <- function(Chongfilename, mytaxa = mytaxa, V = V, depth1 = 1, depth3 = 3, figname){
    load(file = Chongfilename)
    ChongV <- makeV(D = dim(alpha0)[2] + 1) #we used the V from ilrBase() in Chong 18
    gamma0 <- ilrInv(alpha0, V = ChongV)
    gamma1 <- ilrInv(alpha1, V = ChongV)
    gamma2 <- ilrInv(alpha2, V = ChongV)
    z1 <- (depth1 - mean(depth)) / sd(depth) #centre and scale the depths we want
    z3 <- (depth3 - mean(depth)) / sd(depth)
    depth3vs1 <- (z3 - z1) * gamma1 + (z3^2 - z1^2) * gamma2
    depth3vs1 <- acomp(depth3vs1[, mytaxa]) #the subcomposition we want
    d31ilr <- ilr(depth3vs1, V = V)
    epd <- empiricaldepthprior(d31ilr = d31ilr, rscale = 1 / 2, sscale = 2)
    priorsample <- rmvnorm(n = dim(d31ilr)[1], mean = epd$mu, sigma = epd$Sigma)
    plotdepthprior(posteriorsample = d31ilr, priorsample = priorsample, figname = figname)
    print("ilr mean vector for prior on depth effect:")
    print(epd$mu)
    print("ilr covariance matrix for prior on depth effect:")
    print(epd$Sigma)
    print(paste("based on posterior sample of size", dim(d31ilr)[1], "from posterior from Chong 18"))
}

#generate a simple empirical prior for depth effect, based on posterior from Chong '18
#Arguments:
#d31ilr: iterations x ilr components, sample from posterior for depth effect
#rscale (default 1): value by which we multiply the off-diagonals in the correlation matrix in our proposed prior. Decreasing this makes the correlations among ilr components weaker.
#sscale (default 1): value by which we multiply elements of sample standard deviation vector when calculating prior covariance matrix: increasing this makes the prior more diffuse
#Value: a list containing
#mu: mean vector for posterior (sample estimate from posterior in Chong 18),
#r: mean off-diagonal of sample correlation matrix,
#s: vector of sample standard deviations
#Sigma: covariance matrix for prior. Calculate as diag(S) * R * diag(S), where
#S = sscale * s, R is a correlation matrix with all off-diagonals equal to r * rscale
empiricaldepthprior <- function(d31ilr, rscale = 1, sscale = 1){
    mu <- colMeans(d31ilr) #mean vector is sample means from posterior
    s <- apply(d31ilr, 2, sd)
    R <- cor(d31ilr)
    diag(R) <- NA
    r <- mean(R, na.rm = TRUE) #single value for off-diagonal of correlation matrix
    RR <- matrix(r * rscale, nrow = length(mu), ncol = length(mu))
    diag(RR) <- 1
    S <- s * sscale
    Sigma <- diag(S) %*% RR %*% diag(S)
    return(list(mu = mu, r = r, s = s, Sigma = Sigma))
}

#pairwise plots of components of sample from posterior mean estimate of depth effect from Chong '18, and of a sample drawn from a proposed prior distribution based on this
#Arguments:
#posteriorsample: iterations x ilr components, drawn from posterior mean estimate of depth effect
#priorsample: iterations x ilr components, drawn from a proposed prior distribution
#figname: name of pdf file to which plot will be written
#Value: pairwise plots of posterior sample (green) and proposed prior based on this (orange)
plotdepthprior <- function(posteriorsample, priorsample, figname = figname){
    k <- dim(posteriorsample)[2]
    pdf(file = figname, width = 7, height = 7)
    par(mfrow = c(k , k))
    par(mar = c(3, 3, 1, 1))
    par(mgp = c(2, 1, 0))
    mycolors <- brewer.pal(3, "Dark2")
    for(i in 1:k){
        for(j in (1):k){
            if(i ==j){
                plot(density(priorsample[, i]), col = mycolors[2], xlab = bquote(alpha[.(i)]), ylab = "density", main = "", lwd = 2, ylim = c(0, max(density(priorsample[, i])$y, density(posteriorsample[, i])$y)))
                lines(density(posteriorsample[, i]), col = mycolors[1], lwd = 2)
            } else {
                plot(priorsample[, c(i, j)], pch = ".", col = adjustcolor(mycolors[2], 0.1), xlab = bquote(alpha[.(i)]), ylab = bquote(alpha[.(j)]))
                points(posteriorsample[, c(i, j)], pch = ".", col = adjustcolor(mycolors[1], 0.1))
            }
            axls <- par("usr")
            text(axls[1] + 0.1 * (axls[2] - axls[1]), axls[4] - 0.1 * (axls[4] - axls[3]), paste("(", letters[(i - 1) * k + j], ")", sep = ""))
            if(i == 1 & j == 1){
                legend("topright", bty = "n", lwd = 2, col = mycolors[1:2], legend = c("Chong", "prior"), cex = 0.75)
            }
        }
    }
    dev.off()
}

#plot histograms on diagonal panels of pairs plot, based on ?panel.hist
panel.hist <- function(x, ...)
{
    usr <- par("usr"); on.exit(par(usr))
    par(usr = c(usr[1:2], 0, 1.5) )
    h <- hist(x, plot = FALSE)
    breaks <- h$breaks; nB <- length(breaks)
    y <- h$counts; y <- y/max(y)
    rect(breaks[-nB], 0, breaks[-1], y, ...)
}

#extract standard deviation vector tau from Cholesky factor LSigma of covariance matrix
#Arguments: Cholesky factor LSigma of covariance matrix
#Value: vector of standard deviations
taufromLSigma <- function(LSigma){
    Sigma <- LSigma %*% t(LSigma)
    tau <- sqrt(diag(Sigma))
    return(tau)
}

#load data on random image effects extracted from Chong and Spencer 2018
#Arguments:
#Chongfilename: name of .RData file containing posterior distribution of LSigma, Cholesky factor of image effect covariance matrix, from Chong and Spencer 2018
#mytaxa: the columns in back-transformed parameters from Chong corresponding to the subcomposition we want
#V: the ilr basis matrix we want for our subcomposition
makeLSigmaprior <- function(Chongfilename, mytaxa = mytaxa, V = V){
    load(file = Chongfilename)
    ChongD <- dim(alpha0)[2] + 1
    ChongV <- makeV(D = ChongD) #we used the V from ilrBase() in Chong 18
    mycolors <- brewer.pal(8, "Dark2")
    reps <- 8
    samplesize <- 1e3
    z2 <- array(dim = c(reps, samplesize, 4))
    for(i in 1:reps){#a few draws from the posterior
        x2 <- rmvnorm(n = samplesize, mean = rep(0, 8), sigma = LSigma[i, ,] %*% t(LSigma[i, ,])) #LSigma is Cholesky factor
        y2 <- ilrInv(x2, V = ChongV)[, mytaxa] #back to simplex, keep only our taxa in the order we want
        z2[i, , ] <- ilr(y2, V = V) #back to ilr
    }
    par(mfrow = c(4, 4))
        for(j in 1:4){
            for(k in 1:4){
                if(j == k){
                    plot(density(z2[1, , j]), col = mycolors[1], main = "")
                    for(i in 2:reps){
                        lines(density(z2[i, , j]), col = mycolors[i])
                    }
                } else {
                    plot(z2[1, , j], z2[1, , k], col = adjustcolor(mycolors[1], 0.4), pch = ".")
                    for(i in 2:reps){
                        points(z2[i, , j], z2[i, , k], col = adjustcolor(mycolors[i], 0.4), pch = ".")
                    }
                }
            }
        }
}

#simulate some possible compositions from independent normal priors in ilr coordinates on overall composition
#Arguments:
#V is basis matrix for ilr transformation
#vmu is variance (same for all components) for ilr components
#Value:
#print out summaries for each component
makemuprior <- function(V, vmu){
    D = dim(V)[1]
    mu <- rmvnorm(n = 1000, mean = rep(0, D - 1), sigma = vmu * diag(D - 1))
    muback <- ilrInv(mu, V = V)
    print(paste("Simulations from independent normal in ilr, with variance", vmu))
    print(summary(unclass(muback)))
    print(apply(unclass(muback), 2, FUN = quantile, probs = c(0.1, 0.9)))
}

#choose scalar multiplier for identity covariance matrix so that a given size of change in a single component is sufficiently plausible (in ilr coordinates)
#Arguments:
#changesize: vector of sizes of change in a single component due to treatment that we want to be plausible (e.g. 2 means we want doubling/halving of a component to be plausible)
#V: basis matrix for ilr
#simulate (default FALSE): if TRUE, check calculation by simulation
#Value: plot probability of getting a vector at least as far from origin as one in which one of the components is multiplied/divided by changesize against scale factor for covariance matrix
makescrapeprior <- function(changesize, V, simulate = FALSE, fname){
    mycolors <- brewer.pal(length(changesize), "Dark2")
    D <- dim(V)[1]
    c <- seq(from = 0.01, to = 2, length.out = 100)
    pdf(file = fname, width = 7, height = 7)
    plot(c, rep(0, length(c)), type = "n", xlab = "scale factor for covariance matrix", ylab = "probability of exceeding size of change in single component", ylim = c(0, 1))
    for(i in 1:length(changesize)){
        r <- acomp(c(changesize[i], rep(1, D - 1)))
        d2 <- sum(r %*% r) #squared Aitchison distance from origin

        dM2 <- 1 / c * d2 #squared Mahalanobis distance in ilr coordinates, assuming covariance matrix is c times identity matrix
        pdM2 <- pchisq(q = dM2, df = D - 1, lower.tail = FALSE)
        lines(c, pdM2, col = mycolors[i], lwd = 2)
    
        if(simulate == TRUE){#check by simulation
            reps <- 1e4
            csim <- seq(from = 0.01, to = 2, length.out = 10)
            for(cc in csim){
                sigma <- cc * diag(D - 1)
                x <- rmvnorm(n = reps, mean = rep(0, D - 1), sigma = sigma)
                dx <- mahalanobis(x, center = rep(0, D - 1), cov = diag(D - 1))
                points(cc, sum(dx > d2) / reps, col = "red")
            }
        }
    }
    legend("bottomright", bty = "n", lwd = 2, pch = NA, col = mycolors, legend = changesize, title = "change in single component")
    dev.off()
}

#Predictions for each panel from fitted manova
#Arguments:
#LZeta: cholesky factor of block covariance matrix (iterations x ilr components x ilr components)
#zblock: standardized block effect (iterations x blocks x ilr components)
#LSigma: cholesky factor of panel covariance matrix (iterations x ilr components x ilr components)
#zblock: standardized panel effect (iterations x panels x ilr components)
#blockindex: vector (one element per panel) indexing the block for each panel
#d1, ..., d5: dummy variables representing treatment effects and interactions (each a vector with one element per panel)
#parameter arrays (each iterations x ilr components):
#mu: overall mean
#alpha: depth contrast
#beta1, beta2: treatment contrasts
#gamma1, gamma2: interaction contrasts
#V: ilr basis matrix
#Value: list containing:
#blockeffect: block effects (iterations x blocks x ilr components)
#paneleffect: panel effects (iterations x panels x ilr components)
#indexedblockeffect: block effects indexed to panels (iterations x panels x ilr components)
#fixedeffect: "fixed" part of fitted model (iterations x panels x ilr components)
#panelprediction: overall prediction (iterations x panels x ilr components)
#panelsimplexprediction: back transformation of panelprediction to simplex (iterations x panels x taxa)
panelpredict <- function(LZeta, zblock, LSigma, zpanel, blockindex, d1, d2, d3, d4, d5, mu, alpha, beta1, beta2, gamma1, gamma2, V){
    niter <- dim(LZeta)[1]
    nblock <- dim(zblock)[2]
    npanel <- dim(zpanel)[2]
    s <- dim(LZeta)[2]
    blockeffect <- array(dim = c(niter, nblock, s))
    paneleffect <- array(dim = c(niter, npanel, s))
    fixedeffect <- array(dim = c(niter, npanel, s))
    D <- cbind(rep(1, npanel), d1, d2, d3, d4, d5)
    for(i in 1:niter){
        blockeffect[i, , ] <- zblock[i, , ] %*% t(LZeta[i, , ])
        paneleffect[i, , ] <- zpanel[i, , ] %*% t(LSigma[i, , ])
        fixedeffect[i, , ] <- D %*% rbind(mu[i, ], alpha[i, ], beta1[i, ], beta2[i, ], gamma1[i, ], gamma2[i, ])
    }
    indexedblockeffect <- blockeffect[, blockindex, ]
    panelprediction <- fixedeffect + paneleffect + indexedblockeffect
    panelsimplexprediction <- array(dim = c(niter, npanel, s + 1))
    for(i in 1:niter){
        panelsimplexprediction[i, , ] <- ilrInv(panelprediction[i, , ], V = V)
    }
    return(list(blockeffect = blockeffect, paneleffect = paneleffect, indexedblockeffect = indexedblockeffect, fixedeffect = fixedeffect, panelprediction = panelprediction, panelsimplexprediction = panelsimplexprediction))
}

#rough plot of observations against predictions from manova
#Arguments:
#pp: predicted proportions (iterations x panels x taxa)
#observed counts: panels x taxa
#ntaxa: number of taxa
#fname: name of pdf file for plot
#Value: pdf scatter plots of predicted against sample proportions, one panel per taxon
manovapredvsobsplot <- function(pp, counts, ntaxa, fname){
    pdf(file = fname, width = 7, height = 7)
    par(mfrow = c(3, 2))
    par(mar = c(4, 4, 2, 2))
    niter <- dim(pp)[1]
    for(i in 1:ntaxa){
        sampleprop <- t(matrix(rep(counts[, i] / rowSums(counts), niter), ncol = niter))
        plot(sampleprop, pp[, , i], pch = 16, col = adjustcolor("black", 0.005), xlim = c(0, 1), ylim = c(0, 1), xlab = "sample", ylab = "predicted")
        text(0, 0.9, paste(letters[i], ": ", names(counts)[i], sep = ""), pos = 4)
    }
    dev.off()
}

#generate posterior predictive sample data sets
#Arguments:
#rho: predicted proportions (iterations x panels x taxa)
#niter: number of iterations to use
#counts: observed counts
#Value: list containing arrays (iterations x panels x taxa
#pcounts: multinomial samples from rho
#pp: sample proportions from pcounts
posteriorsample <- function(rho, niter, counts){
    npanels <- dim(rho)[2]
    pcounts <- array(dim = c(niter, npanels, dim(rho)[3]))
    pp <- pcounts
    npoints <- rowSums(counts)
    for(i in 1:niter){
        for(j in 1:npanels){
            pcounts[i, j, ] <- rmultinom(n = 1, prob = rho[i, j, ], size = npoints[j])
            pp[i, j, ] <- pcounts[i, j, ] / npoints[j]
        }
    }
    return(list(pcounts = pcounts, pp = pp))
}

#set up data for SBCRS
#Arguments:
#seed: seed for pseudorandom number generator
#Value: list containing all variables in the data block of aureliamanova.stan other than counts
#Note: I think seed is the only argument we are allowed, so assumes the required variables are in the main workspace
gendata <- function(seed){
    set.seed(seed + 1e6)
    ntaxa <- ntaxa
    npanels <- dim(manovainputs$counts)[1]
    nblocks <- max(manovainputs$blockindex)
    tVinv <- V
    blockindex <- manovainputs$blockindex
    d1 <- manovainputs$d1
    d2 <- manovainputs$d2
    d3 <- manovainputs$d3
    d4 <- manovainputs$d4
    d5 <- manovainputs$d5
    return(list(ntaxa = ntaxa, npanels = npanels, nblocks = nblocks, tVinv = tVinv, blockindex = blockindex, d1 = d1, d2 = d2, d3 = d3, d4 = d4, d5 = d5))
}

#sample from priors for SBCRS
#Arguments:
#seed: seed for pseudorandom number generator
#data: output from gendata()
#Value: samples from priors for all the parameters appearing in the parameters block in aureliamanova.stan
#Note: priors are hard coded here: need to make sure they match what's hard coded in aureliamanova.stan
genparams <- function(seed, data){
    set.seed(seed + 2e6)
    s <- data$ntaxa - 1
    mu <- rnorm(n = s, mean = 0, sd = 2)
    mudepth  <-  c(2.8, -2.6, -0.1, 0.1)
    Sigmadepth  <- matrix(c(12.7, 3.2e-4, 5.5e-5, 8.5e-5, 3.2e-4, 7.1, 4.1e-5, 6.4e-5, 5.5e-5, 4.1e-5, 2.1e-1, 1.1e-5, 8.5e-5, 6.4e-5, 1.1e-5, 5.0e-1), nrow = 4, ncol = 4)
    alpha <- as.numeric(rmvnorm(n = 1, mean = mudepth, sigma = Sigmadepth))
    beta1 <- rnorm(n = s, mean = 0, sd = 1)
    beta2 <- rnorm(n = s, mean = 0, sd = 1)
    gamma1 <- rnorm(n = s, mean = 0, sd = 1)
    gamma2 <- rnorm(n = s, mean = 0, sd = 1)
    zpanel <- array(rnorm(n = data$npanels * s, mean = 0, sd = 1), dim = c(data$npanels, s))
    zblock <- array(rnorm(n = data$nblocks * s, mean = 0, sd = 1), dim = c(data$nblocks, s))
    LOmegaSigma <- chol((rlkjcorr(n = 1, K = s, eta = 2)))
    tauSigma <- abs(rcauchy(n = s, location = 0, scale = 2.5))
    LOmegaZeta <- chol((rlkjcorr(n = 1, K = s, eta = 2)))
    tauZeta <- abs(rcauchy(n = s, location = 0, scale = 2.5))
    return(list(mu = mu, alpha = alpha, beta1 = beta1, beta2 = beta2, gamma1 = gamma1, gamma2 = gamma2, zpanel = zpanel, zblock = zblock, LOmegaSigma = LOmegaSigma, tauSigma = tauSigma, LOmegaZeta = LOmegaZeta, tauZeta = tauZeta))
}

#generated a simulated manova data set from the prior
#Arguments:
#seed: pseudorandom number seed
#data: output from gendata()
#params: output from genparams()
#Value:
#list containing counts, multinomial sample, npanels x ntaxa, with taxon probabilities generated from priors for manova model. Number of points in each row comes from rowSums(manovainputs$counts), assumed to be available in main workspace
genmodeleddata <- function(seed, data, params){
    set.seed(seed + 3e6)
    LSigma <- diag(params$tauSigma) %*% params$LOmegaSigma
    LZeta <- diag(params$tauZeta) %*% params$LOmegaZeta
    blockeffect <- params$zblock %*% LZeta
    blockeffect <- blockeffect[data$blockindex, ]
    D <- cbind(rep(1, data$npanel), data$d1, data$d2, data$d3, data$d4, data$d5)
    x <- D %*% rbind(params$mu, params$alpha, params$beta1, params$beta2, params$gamma1, params$gamma2) + params$zpanel %*% LSigma + blockeffect
    rho <- ilrInv(x, V = data$tVinv)
    counts <- array(dim = c(data$npanels, data$ntaxa))
    points <- rowSums(manovainputs$counts)
    for(i in 1:data$npanels){
        counts[i, ] <- rmultinom(n = 1, size = points[i], prob = rho[i, ])
    }
    return(list(counts = counts))
}

#function to call rstan for SBC
#Arguments:
#seed: pseudorandom number seed
#data: output from gendata()
#params: output from genparams()
#modeled_data: output from genmodeleddata
#iters: number of iterations
#Note: manovamodel is compiled manova model, assumed to exist in main workspace
samplefrommodel <- function(seed, data, params, modeled_data, iters){
    dataforstan <- c(data, modeled_data)
    rstan::sampling(manovamodel, chains = 1, iter = 2 * iters, warmup = iters, data = dataforstan, seed = seed, refresh = 1000, open_progress = FALSE, show_messages = FALSE)
}


#debug SBCRS code: are we generating NA values anywhere?
testSBCRS <- function(){
    seed <- 12345
    iter <- 1e4
    for(i in 1:iter){
        data <- gendata(seed = seed)
        params <- genparams(seed = seed, data = data)
        modeleddata <- genmodeleddata(seed = seed, data = data, params = params)
        testparams <- unlist(lapply(params, is.na))
        if(any(testparams)) print(params)
        if(any(is.na(modeleddata$counts))) print(modeleddata)
     }
}

#simulation-based calibration of final week manova using rstan::sbc
#Arguments:
#manovastanfile: name of .stan file specifying model
#manovainputs: list returned by aureliamanovasetup
#ntaxa: number of taxa
#V: ilr basis matrix
#M: number of times to condition on draws from prior predictive distribution
#iter: number of iterations (half sampling, half warmup) per simulated data set
#Value: object returned by rstan::sbc
SBC <- function(manovastanfile, manovainputs, ntaxa, V, M, iter){
    manovamodel <- stan_model(manovastanfile)
    npanels <- dim(manovainputs$counts)[1]
    nblocks <- max(manovainputs$blockindex)
    n <- rowSums(manovainputs$counts)
    sbcoutput <- sbc(manovamodel, data = list(ntaxa = ntaxa, npanels = npanels, tVinv = V, blockindex = manovainputs$blockindex, n = n, d1 = manovainputs$d1, d2 = manovainputs$d2, d3 = manovainputs$d3, d4 = manovainputs$d4, d5 = manovainputs$d5), M = M, iter = iter, refresh = 0)
    return(sbcoutput)
}

#extract ranks from rstan::sbc, for use in plotting. Based on rstan:plot.sbc
#Arguments:
#x: object returned by rstan::sbc
#thin: thinning level for ranks (default 3)
#Value: matrix (simulated data sets x parameters) of sbc ranks
getsbcranks <- function(x, thin = 3, ...){
    thinner <- seq(from = 1, to = nrow(x$ranks[[1]]), by = thin)
    u <- t(sapply(x$ranks, FUN = function(r) 1L + colSums(r[thinner, , drop = FALSE])))
    return(u)
}

#plot function for rstan sbc output, based on rstan::plot.sbc
#Arguments:
#sbcranks: matrix (simulated data sets x parameters) of sbc ranks
#manovaresultspath: path to directory where figures will be written
#Value: pdfs of sbc rank plots for sets of parameters
#Note: indexing would need to change if number or order of parameters in stan code changes
myplot.sbc <- function(sbcranks, manovaresultspath) {
    sbchist(fname = paste(manovaresultspath, "/SBC_mualphabeta1beta2.pdf", sep = ""), startindex = 1, endindex = 16, sbcranks = sbcranks)
    sbchist(fname = paste(manovaresultspath, "/SBC_gamma1gamma2.pdf", sep = ""), startindex = 17, endindex = 24, sbcranks = sbcranks)
    for(j in 1:15){
        startpanel <- 1 + (j - 1) * 4
        endpanel <- j * 4
        startindex <- 25 + (startpanel - 1) * 4
        endindex <- endpanel * 4 + 24
        sbchist(fname = paste(manovaresultspath, "/SBC_zpanel_", startpanel, "_", endpanel, ".pdf", sep = ""), startindex = startindex, endindex = endindex, sbcranks = sbcranks)
    }
    sbchist(fname = paste(manovaresultspath, "/SBC_zblock_1_4.pdf", sep = ""), startindex = 265, endindex = 280, sbcranks = sbcranks)
    sbchist(fname = paste(manovaresultspath, "/SBC_zblock_5_8.pdf", sep = ""), startindex = 281, endindex = 296, sbcranks = sbcranks)
    sbchist(fname = paste(manovaresultspath, "/SBC_zblock_9_10.pdf", sep = ""), startindex = 297, endindex = 304, sbcranks = sbcranks)
    sbchist(fname = paste(manovaresultspath, "/SBC_LSigma.pdf", sep = ""), startindex = 305, endindex = 320, sbcranks = sbcranks)
    sbchist(fname = paste(manovaresultspath, "/SBC_LZeta.pdf", sep = ""), startindex = 321, endindex = 336, sbcranks = sbcranks)
}

#produce a set of up to 16 histograms of SBC ranks
#Arguments:
#fname: name of pdf file for plot
#startindex, endindex: column indices of first and last parameters to plot in matrix sbcranks
#sbcranks: matrix (simulated data sets x parameters) of SBC ranks
#Value: pdf file, histograms of SBC ranks
sbchist <- function(fname, startindex, endindex, sbcranks){
    pdf(file = fname, width = 7, height = 7)
    par(mfrow = c(4, 4))
    par(mar = c(4, 2, 1, 1))
    for(i in startindex:endindex){
        hist(sbcranks[, i], xlab = colnames(sbcranks)[i], main = "", breaks = 12)
    }
    dev.off()
}

#ternary plot of manova results and sample data from final week in simplex using orthogonal projection
#Arguments:
#manovainputs: object returned by aureliamanovasetup
#alldata: raw data
#mu: posterior sample from overall mean (iterations x ilr components)
#alpha: posterior sample from depth effect (iterations x ilr components)
#beta1, beta2: posterior samples from treatment contrasts (iterations x ilr components)
#gamma1, gamma2: posterior samples from interaction contrasts (iterations x ilr components)
#date: date for which we want subset of data (the last week of experiment)
#preorpost: "Pre" or "Post" scrape
#V: ilr basis matrix
#centre: logical (default FALSE): centre on overall mean vector from manova?
#coords: set of ilr coordinates to use in projection (e.g. c(1, 2) for first two)
#plotcoords: which columns of orthogonal projection we want to plot (e.g. c(1, 2, 3): will need one more than the number in coords, but choice depends on V
#labels: labels for the parts on the ternary plot
#fname: name of pdf file to plot to (if NA, don't create a file)
#paneltext: label for panel (if NA, no label)
#plotlegend (logical, default TRUE): add a legend?
#Value: plot of raw data for last week, with treatment means and 95% HPD credible regions from manova, written to fname
manovaplot <- function(manovainputs, alldata, mu, alpha, beta1, beta2, gamma1, gamma2, date, preorpost, V, centre = FALSE, coords, plotcoords, labels, fname, paneltext = NA, plotlegend = TRUE){
    lastweekcomp <- manovainputs$counts
    lastweekcomp[lastweekcomp == 0] <- 0.5 #adding half detection limit to plot
    lastweekcomp <- acomp(lastweekcomp)
    ilrlastweek <- ilr(lastweekcomp, V = V)
    meanmu <- colMeans(mu)
    if(centre == TRUE) ilrlastweek <- sweep(ilrlastweek, 2, meanmu, "-")
    lastweekproject <- acomp(getorthogonalprojection(x = ilrlastweek, coords = coords, V = V)$xprojsimplex[, plotcoords])
    lastweek <- alldata[alldata$date == date & alldata$prepost == preorpost, ]
    mycolors <- brewer.pal(3, "Dark2")
    mypch <- c(1, 16)
    mylty <- c("dashed", "solid")
    if(!is.na(fname)){
        pdf(file = fname, width = 7, height = 7)
    }
    par(mar = c(1, 6, 1, 4))
    plot(acomp(lastweekproject[lastweek$treatment == "C", ]), pch = mypch[factor(lastweek$depthm)], col = adjustcolor(mycolors[1], 0.7), labels = labels) #green
    plot(acomp(lastweekproject[lastweek$treatment == "A", ]), pch = mypch[factor(lastweek$depthm)], col = adjustcolor(mycolors[2], 0.7), add = TRUE) #orange
    plot(acomp(lastweekproject[lastweek$treatment == "O", ]), pch = mypch[factor(lastweek$depthm)], col = adjustcolor(mycolors[3], 0.7), add = TRUE) #blue
    depths = c(1, 3)
    treats <- c("C", "A" ,"O") 
    for(i in 1:2){
        for(j in 1:3){
            D <- as.numeric(cbind(1, t(buildcontrastbasis(depth = depths[i], treatment = treats[j]))))
            treatfixedeff <- getfixedeffect(mu = mu, alpha = alpha, beta1 = beta1, beta2 = beta2, gamma1 = gamma1, gamma2 = gamma2, D = D)
            if(centre == TRUE) treatfixedeff <- sweep(treatfixedeff, 2, meanmu, "-")
            treatmean <- colMeans(treatfixedeff)
            treatcontour <- gethdrcontour(x = treatfixedeff[, coords], prob = 95)
            treatmeansimplex <- acomp(getorthogonalprojection(x = treatmean, coords = coords, V = V)$xprojsimplex[plotcoords])
            plot(treatmeansimplex, pch = mypch[i], col = adjustcolor(mycolors[j], 0.7), cex = 2, add = TRUE)
            for(k in 1:length(treatcontour)){
                plotilrcontour(contour = treatcontour[[k]], s = dim(ilrlastweek)[2], V = V, col = adjustcolor(mycolors[j], 0.7), lty = mylty[i], coords = coords, plotcoords = plotcoords)
            }
        }
    }
    if(plotlegend){
        legend(x = 1, y = 1, pch = rep(mypch, 3), col = mycolors[c(1, 1, 2, 2, 3, 3)], lty = rep(mylty, 3), legend = c(expression(paste(1~m, ",", ~italic(C))), expression(paste(3~m, ",", ~italic(C))), expression(paste(1~m, ",", ~italic(A))), expression(paste(3~m, ",", ~italic(A))), expression(paste(1~m, ",", ~italic(O))), expression(paste(3~m, ",", ~italic(O)))), bty = "n")
    }
    if(!is.na(paneltext)){
        text(0, 0.925, paneltext)
    }
    if(!is.na(fname)){
        dev.off()
    }
}

#get the fixed effect for a treatment combination
#Arguments:
#mu: overall mean in ilr (iterations x components)
#alpha: depth effect in ilr (iterations x components)
#beta1: first treatment contrast in ilr (iterations x components)
#beta2: second treatment contrast in ilr (iterations x components)
#D: vector of (dummy) explanatory variables for specified treatment combination: overall mean, contrasts d1 to d5
#Value:
#matrix of fixed effects for the specified treatment combinatin: iterations x components
getfixedeffect <- function(mu, alpha, beta1, beta2, gamma1, gamma2, D){
    niter <- dim(mu)[1]
    s <- dim(mu)[2]
    fixedeffect <- array(dim = c(niter, s))
    for(i in 1:niter){
        fixedeffect[i, ] <- D %*% rbind(mu[i, ], alpha[i, ], beta1[i, ], beta2[i, ], gamma1[i, ], gamma2[i, ])
    }
    return(fixedeffect)
}


#orthogonal projection of x onto the subspace spanned by a given set of coordinates
#Arguments:
#matrix x (in an orthogonal coordinate system)
#coords: indices of the coordinates onto which we want to project x
#V: ilr basis matrix
#Value: list containing:
#xproj: x projected onto the given coordinates
#xprojsimplex: back-transformation of xproj to simplex
#note: still have to choose by hand which columns of the projection are of interest, e.g. if x has 4 columns, coords is 1:2, and first two columns of V contrast part 1 with 2 and geomean of 1 and 2 with geomean of 3 and 4, then columns 1:3 of xprojsimplex are probably the ones of interest
getorthogonalprojection <- function(x, coords, V){
    xproj <- x
    if(is.vector(xproj)){
        xproj[-coords] <- 0
    } else {
        xproj[, -coords] <- 0
    }
    xprojsimplex <- ilrInv(xproj, V = V)
    return(list(xproj = xproj, xprojsimplex = xprojsimplex))
}

#get contours of highest posterior density of orthogonal projection in first two coordinates
#Arguments:
#x: array of ilr coordinates
#prob: percentage at which we want the contour e.g. 95
#Value:
#a list, each element contains variables x and y, the contour in first two coordinates
gethdrcontour <- function(x, prob){
    myhdr <- hdr.2d(x = x[, 1], y = x[, 2], prob = prob) #first two coordinates: orthogonal projection
    mycontour <- contourLines(myhdr$den, levels = myhdr$falpha)
    return(mycontour)
}

#plot a contour in the simplex from ilr coordinates
#Arguments:
#contour: an object containing variables x and y, first two ilr coordinates
#s: number of coordinates (those other than first two are set to zero (assuming we have an orthogonal basis, this gives orthogonal projection)
#V: ilr basis matrix
#col: colour for contour
#lty: line style for contour
plotilrcontour <- function(contour, s, V, col, lty, coords, plotcoords){
    ilrcontour <- array(0, dim = c(length(contour$x), s)) #zeros in remaining coordinates
    ilrcontour[, coords[1]] <- contour$x
    ilrcontour[, coords[2]] <- contour$y
    simplexcontour <- acomp(ilrInv(ilrcontour, V = V)[, plotcoords])
    plot(simplexcontour, type = "l", add = TRUE, col = col, lty = lty)
}

#what does a null hypothesis of the form a / (not a) = constant look like when we do orthogonal projection into lower dimension?
testamalgamatednull <- function(){
    a <- 0.2
    n <- 1e5
    eps <- 1e-6
    myvec <- rmvnorm(n = n, mean = rep(0, 3), sigma = 10 * diag(3)) #sample ilr of last 4 parts of 5-part composition
    mycomp <- acomp(cbind(rep(a, n), (1 - a) * unclass(ilrInv(myvec)))) #construct a 5-part composition in simplex with constant first part a
    print(mycomp[1:10, ])
    myilr <- ilr(mycomp, V = V) #transform to ilr
    myilr[, 3:4] <- 0 #orthogonal projection onto first two coordinates
    myback <- acomp(ilrInv(myilr, V = V)[, 1:3]) #back-transform orthogonal projection and produce marginal plot
    plot(myback, pch = ".")
    isoPortionLines()
    point1 <- acomp(c(a, 1 - a, eps)) #calculated corners of the region representing the null hypothesis
    point2 <- acomp(c(a, eps, eps))
    point3 <- acomp(c(a, eps, (1 - a) / 3))
    plot(point1, col = "red", pch = 16, add = TRUE)
    plot(point2, col = "green", pch = 16, add = TRUE)
    plot(point3, col = "blue", pch = 16, add = TRUE)
          
}

#Calculate distributions of test statistics relating to logit change in Aurelia (O vs C) and potential competitors (A vs C) at each depth
#Arguments:
#mu, alpha, beta1, beta2, gamma1, gamma2: posterior samples from manova parameters (each iterations x ilr components)
#V: ilr basis matrix
#fname: name of pdf file to which plot will be written
#Value:
#pdf file with distribution of each test statistic at each depth
#and print out proportion positive responses in sample
testeffects <- function(mu, alpha, beta1, beta2, gamma1, gamma2, V, fname){
    mycolors <- brewer.pal(3, 'Dark2')
    depths = c(1, 3)
    pdf(file = fname, width = 7, height = 7)
    par(mfrow = c(2, 1))
    par(mar = c(4.5, 4, 2, 1))
    pls <- c("a: 1 m", "b: 3 m")
    legends  <- list(c(expression(Potential~competitors*","~italic(A)~vs~italic(C)), expression(italic(A.~aurita)*","~italic(O)~vs~italic(C))), NULL)
    xlim <- c(-3, 3) #chose these values by hand to put all density plots on same scale
    ylim <- c(0, 2)
    for(i in 1:2){
        DC <- as.numeric(cbind(1, t(buildcontrastbasis(depth = depths[i], treatment = "C"))))
        treatfixedeffC <- getfixedeffect(mu = mu, alpha = alpha, beta1 = beta1, beta2 = beta2, gamma1 = gamma1, gamma2 = gamma2, D = DC)
        DO <- as.numeric(cbind(1, t(buildcontrastbasis(depth = depths[i], treatment = "O"))))
        treatfixedeffO <- getfixedeffect(mu = mu, alpha = alpha, beta1 = beta1, beta2 = beta2, gamma1 = gamma1, gamma2 = gamma2, D = DO)
        Oresponse <- logitresponse(fixedeff1 = treatfixedeffO, fixedeff2 = treatfixedeffC, taxon = 1, V = V)
        DA <- as.numeric(cbind(1, t(buildcontrastbasis(depth = depths[i], treatment = "A"))))
        treatfixedeffA <- getfixedeffect(mu = mu, alpha = alpha, beta1 = beta1, beta2 = beta2, gamma1 = gamma1, gamma2 = gamma2, D = DA)
        Aresponse <- logitresponse(fixedeff1 = treatfixedeffA, fixedeff2 = treatfixedeffC, taxon = c(3, 4, 5), V = V)
        print(paste("Depth :", depths[i], sep = ""))
        print("Response of A. aurita to O treatment:")
        print(summary(Oresponse$logitchange))
        print(hdr(Oresponse$logitchange)$hdr)
        print(paste("Proportion positive: ", Oresponse$p, sep = ""))
        print("Response of potential competitors to A treatment:")
        print(summary(Aresponse$logitchange))
        print(hdr(Aresponse$logitchange)$hdr)
        print(paste("Proportion positive: ", Aresponse$p, sep = ""))
        plotoneresponse(logitchange1 = Aresponse$logitchange, logitchange2 = Oresponse$logitchange, cols = mycolors[c(2, 3)], panellabel = pls[i], xlim = xlim, ylim = ylim, legend = legends[[i]])
    }
    dev.off()
}

#calculate difference in logit response of a set of taxa between two treatments
#Arguments:
#fixedeff1, fixedeff2: fixed effects of two treatments 1 and 2 (iterations x ilr components)
#taxon: the taxa of interest: may be a single index or a vector
#V: ilr basis matrix
#Value: a list containing:
#logitchange: the posterior distribution of difference in log(sum of parts in taxon / (1 - sum of parts in taxon) between treatment 1 and treatment 2
#p: proportion of positive entries in logitchange
logitresponse <- function(fixedeff1, fixedeff2, taxon, V){
    simplex1 <- unclass(ilrInv(fixedeff1, V = V))
    simplex2 <- unclass(ilrInv(fixedeff2, V = V))
    if(length(taxon) > 1){
        rst1 <- rowSums(simplex1[, taxon])
        rst2 <- rowSums(simplex2[, taxon])
    } else {
        rst1 <- simplex1[, taxon]
        rst2 <- simplex2[, taxon]
    }
    logit1 <- log(rst1 / (1 - rst1))
    logit2 <- log(rst2 / (1 - rst2))
    logitchange <- logit1 - logit2
    p <- sum(logitchange > 0) / (length(logitchange))
    return(list(logitchange = logitchange, p = p))
}

#plot logit responses to treatments at one depth
#Arguments:
#logitchange1, logitchange2: posterior samples from difference in logit of potential competitors (A vs C treatment) and A. aurita (O vs C treatment) respectively
#cols: vector of colours
#panellabel: label for top left of panel
#xlim and ylim: axis limits
#legend: labels for legend: logitchange1, logitchange2 in that order. If NULL (default), no legend
#Value: kernel density plots of logitchange1 and logitchange 2, with vertical line at 0
plotoneresponse <- function(logitchange1, logitchange2, cols, panellabel, xlim, ylim, legend = NULL){
    d1 <- density(logitchange1)
    plot(d1, main = "", type = "n", xlab = "logit difference", ylab = "density", xlim = xlim, ylim = ylim)
    polygon(d1, col = adjustcolor(cols[1], alpha = 0.4), border = cols[1])
    d2 <- density(logitchange2)
    polygon(d2, col = adjustcolor(cols[2], alpha = 0.4), border = cols[2])
    abline(v = 0, lwd = 2, col = "grey")
    axlims <- par("usr")
    text(axlims[1] + 0.05 * (axlims[2] - axlims[1]), axlims[4] - 0.1 * (axlims[4] - axlims[3]), panellabel, pos = 4)
    if(!is.null(legend))
        legend("topright", pch = c(15, 15), col = adjustcolor(cols, 0.4), legend = legend, bty = "n")
}

#Calculate a 4-dimensional hyperellipsoid of concentration
#Arguments:
#r: Mahalanobis distance (default 1)
#sigma: covariance matrix (default diag(4))
#mu: mean vector (default rep(0, 4)
#n: number of points to generate (default 100)
#plot (default FALSE): produce a set of marginal 3d scatterplots?
#Value:
#an n x 4 matrix of coordinates of the required hyperellipsoid, all points at specified Mahalanobis distance from mean vector mu, under covariance matrix sigma
mhhyperellipsoid <- function(r = 1, sigma = diag(4), mu = rep(0, 4), n = 100, plot = FALSE){
    phi1 <- seq(from = 0, to = 2 * pi, length.out = n)
    phi2 <- seq(from = 0, to = pi, length.out = n)
    phi3 <- seq(from = 0, to = pi, length.out = n)
    phis <- expand.grid(phi1, phi2, phi3)
    x1 <- sin(phis[, 1]) #construct as a hypersurface of revolution: phi3 is the angle through which we rotate the parametric form of a sphere
    x2 <- cos(phis[, 1]) * sin(phis[, 2])
    x3 <- cos(phis[, 1]) * cos(phis[, 2]) * sin(phis[, 3])
    x4 <- cos(phis[, 1]) * cos(phis[, 2]) * cos(phis[, 3])
    x <- cbind(x1, x2, x3, x4)
    dm <- sqrt(mahalanobis(x, center = rep(0, 4), cov = sigma))
    xm <- r * x / dm
    xm <- sweep(xm, 2, mu, "+")
    if(plot == TRUE){
        par(mfrow = c(2, 2))
        alpha <- 0.01
        scatterplot3d(xm[, c(1, 2, 3)], pch = ".", xlab = "x1", ylab = "x2", zlab = "x3", color = adjustcolor("black", alpha = alpha))
        scatterplot3d(xm[, c(1, 2, 4)], pch = ".", xlab = "x1", ylab = "x2", zlab = "x4", color = adjustcolor("black", alpha = alpha))
        scatterplot3d(xm[, c(1, 3, 4)], , pch = ".", xlab = "x1", ylab = "x3", zlab = "x3", color = adjustcolor("black", alpha = alpha))
        scatterplot3d(xm[, c(2, 3, 4)], , pch = ".", xlab = "x2", ylab = "x3", zlab = "x4", color = adjustcolor("black", alpha = alpha))
    }
    return(xm)
}

#plot projections of hyperellipsoids (or marginal ellipses) of unit Mahalanobis distance based on panel and block covariance matrices onto two 2-simplexes. Hard-coded for 5-part composition.
#Arguments:
#LSigma, LZeta: Cholesky factors of panel and block covariance matrices (each iterations x 4 x 4)
#n: number of hyperellipsoids to sample from posterior distribution
#m: number of points per hyperellipsoid
#cov12fname, cov34fname: file names for pdf files of projections onto coordinates 1,2 and 3,4
#labels12, labels34: labels for the parts of the projections
#alpha: transparency level (default 0.1)
#method: options are "hyper" (calculate hyperellipsoid in four dimensions and project) or "2d" (default, calculate two-dimensional marginal ellipsoids)
#Value: pdf plots cov12fname, cov34fname: projections of hyperellipsoids (or marginal ellipses) based on panel and block covariance matrices
plothyperellipsoids <- function(LSigma, LZeta, n = 100, m = 10, cov12fname, cov34fname, labels12, labels34, alpha = 0.1, method = "2d"){
    if(method == "hyper"){
        xsigma12projsimplex <- array(dim = c(n, m^3, 3))
        xsigma34projsimplex <- array(dim = c(n, m^3, 3))
        xzeta12projsimplex <- array(dim = c(n, m^3, 3))
        xzeta34projsimplex <- array(dim = c(n, m^3, 3))
        for(i in 1:n){
            Sigma <- LSigma[i, , ] %*% t(LSigma[i, , ])
            Zeta <- LZeta[i, , ] %*% t(LZeta[i, , ])
            xsigma <- mhhyperellipsoid(r = 1, mu = rep(0, 4), sigma = Sigma, n = m)
            xsigma12 <- getorthogonalprojection(x = xsigma, coords = c(1, 2), V = V)
            xsigma34 <- getorthogonalprojection(x = xsigma, coords = c(3, 4), V = V)
            xsigma12projsimplex[i, , ] <- xsigma12$xprojsimplex[, c(1, 2, 3)]
            xsigma34projsimplex[i, , ] <- xsigma34$xprojsimplex[, c(3, 4, 5)]
            xzeta <- mhhyperellipsoid(r = 1, mu = rep(0, 4), sigma = Zeta, n = m)
            xzeta12 <- getorthogonalprojection(x = xzeta, coords = c(1, 2), V = V)
            xzeta34 <- getorthogonalprojection(x = xzeta, coords = c(3, 4), V = V)
            xzeta12projsimplex[i, , ] <- xzeta12$xprojsimplex[, c(1, 2, 3)]
            xzeta34projsimplex[i, , ] <- xzeta34$xprojsimplex[, c(3, 4, 5)]
    }
    } else if(method == "2d"){
        xsigma12projsimplex <- array(dim = c(n, m, 3))
        xsigma34projsimplex <- array(dim = c(n, m, 3))
        xzeta12projsimplex <- array(dim = c(n, m, 3))
        xzeta34projsimplex <- array(dim = c(n, m, 3))
        for(i in 1:n){
            Sigma <- LSigma[i, , ] %*% t(LSigma[i, , ])
            Zeta <- LZeta[i, , ] %*% t(LZeta[i, , ])
            xsigma12 <- cbind(mhellipse(mu = c(0, 0), sigma = Sigma[1:2, 1:2], r = 1, n = m), 0, 0)
            xsigma12projsimplex[i, , ] <- ilrInv(xsigma12, V = V)[, c(1, 2, 3)]
            xsigma34 <- cbind(0, 0, mhellipse(mu = c(0, 0), sigma = Sigma[3:4, 3:4], r = 1, n = m))
            xsigma34projsimplex[i, , ] <- ilrInv(xsigma34, V = V)[, c(3, 4, 5)]
            xzeta12 <- cbind(mhellipse(mu = c(0, 0), sigma = Zeta[1:2, 1:2], r = 1, n = m), 0, 0)
            xzeta12projsimplex[i, , ] <- ilrInv(xzeta12, V = V)[, c(1, 2, 3)]
            xzeta34 <- cbind(0, 0, mhellipse(mu = c(0, 0), sigma = Zeta[3:4, 3:4], r = 1, n = m))
            xzeta34projsimplex[i, , ] <- ilrInv(xzeta34, V = V)[, c(3, 4, 5)]
        }
    } else stop(paste("Method ", method, " not implemented", sep = ""))
    mycolors <- brewer.pal(3, "Dark2")
    if(method == "hyper"){
        pch  <-  "."
        legendpch <- 16
        lty  <- NA
    } else {
        pch = NA
        legendpch <- NA
        lty  <- "solid"
    }
    pdf(file = cov12fname, width = 7, height = 7)
    par(mar = c(1, 6, 1, 4))
    plot(acomp(c(1, 1, 1)), labels = labels12, type = "n")
    if(method == "hyper"){
        for(i in 1:n){
            plot(acomp(xsigma12projsimplex[i, , ]), pch = pch, add = TRUE, col = adjustcolor(mycolors[1], alpha = alpha))
            plot(acomp(xzeta12projsimplex[i, , ]), pch = pch, add = TRUE, col = adjustcolor(mycolors[2], alpha = alpha))
        }
        legend("topleft", pch = rep(legendpch, 2), col = mycolors[1:2], legend = c(expression(Sigma), expression(Z)), bty = "n")
    } else {
        for(i in 1:n){
            plot(acomp(xsigma12projsimplex[i, , ]), type = "l", lty = lty, add = TRUE, col = adjustcolor(mycolors[1], alpha = alpha))
            plot(acomp(xzeta12projsimplex[i, , ]), type = "l", lty = lty, add = TRUE, col = adjustcolor(mycolors[2], alpha = alpha))
        }
        legend("topleft", pch = rep(legendpch, 2), lty = rep(lty, 2), col = mycolors[1:2], legend = c(expression(Sigma), expression(Z)), bty = "n")
    }
    dev.off()
    pdf(file = cov34fname, width = 7, height = 7)
    par(mar = c(1, 6, 1, 4))
    plot(acomp(c(1, 1, 1)), labels = labels34, type = "n")
    if(method == "hyper"){
        for(i in 1:n){
            plot(acomp(xsigma34projsimplex[i, , ]), pch = pch, add = TRUE, col = adjustcolor(mycolors[1], alpha = alpha))
            plot(acomp(xzeta34projsimplex[i, , ]), pch = pch, add = TRUE, col = adjustcolor(mycolors[2], alpha = alpha))
        }
        legend("topleft", pch = rep(legendpch, 2), col = mycolors[1:2], legend = c(expression(Sigma), expression(Z)), bty = "n")
    } else {
        for(i in 1:n){
            plot(acomp(xsigma34projsimplex[i, , ]), type = "l", lty = lty, add = TRUE, col = adjustcolor(mycolors[1], alpha = alpha))
            plot(acomp(xzeta34projsimplex[i, , ]), type = "l", lty = lty, add = TRUE, col = adjustcolor(mycolors[2], alpha = alpha))
        }
        legend("topleft", pch = rep(legendpch, 2), lty = rep(lty, 2), col = mycolors[1:2], legend = c(expression(Sigma), expression(Z)), bty = "n")
    }

    dev.off()
}


#Calculate distributions of test statistics relating to logit change in Aurelia (O vs C) and potential competitors (A vs C) at each depth from raw data, rather than from manova outputs
#Arguments:
#alldata: data frame containing panel, date, prepost and count for each class
#date: the date we want (last sampling week)
#preorpost: "Pre" or "Post" (before or after scrape
#Value:
#plot of distribution of each test statistic at each depth
#Example:
#rawcounteffects(alldata = alldata, date = lastdate, preorpost = "Pre")
rawcounteffects <- function(alldata, date, preorpost){
    lastweek <- alldata[alldata$date == date & alldata$prepost == preorpost, ]
    counts <- lastweek[, taxa] + 1 #add-one pseudocount
    props <- sweep(counts, 1, rowSums(counts), "/")
    logitAurelia <- logit(props[, 1])
    potentialcomp <- rowSums(props[, 3:5])
    logitcomp <- logit(potentialcomp)
    Oresponse1m <- logitAurelia[lastweek$treatment == "O" & lastweek$depthm == 1] - logitAurelia[lastweek$treatment == "C" & lastweek$depthm == 1]#difference in logit Aurelia between O and C treatments at 1 m
    Oresponse3m <- logitAurelia[lastweek$treatment == "O" & lastweek$depthm == 3] - logitAurelia[lastweek$treatment == "C" & lastweek$depthm == 3]#difference in logit Aurelia between O and C treatments at 3 m
    Aresponse1m <- logitcomp[lastweek$treatment == "O" & lastweek$depthm == 1] - logitcomp[lastweek$treatment == "C" & lastweek$depthm == 1]#difference in logit potential competitors between A and C treatments at 1 m
    Aresponse3m <- logitcomp[lastweek$treatment == "O" & lastweek$depthm == 3] - logitcomp[lastweek$treatment == "C" & lastweek$depthm == 3]#difference in logit potential competitors between A and C treatments at 3 m
    mycols <- brewer.pal(3, "Dark2")
    legends  <- list(c(expression(Potential~competitors*","~italic(A)~vs~italic(C)), expression(italic(A.~aurita)*","~italic(O)~vs~italic(C))), NULL)
    depths <- c(1, 3)
    par(mfrow = c(2, 1))
    par(mar = c(4.5, 4, 2, 1))
    plotoneresponse(logitchange1 = Aresponse1m, logitchange2 = Oresponse1m, cols = mycols[2:3], panellabel = "a: 1 m", xlim = c(-5, 5), ylim = c(0, 0.8), legend = legends[[1]])
    plotoneresponse(logitchange1 = Aresponse3m, logitchange2 = Oresponse3m, cols = mycols[2:3], panellabel = "b: 3 m", xlim = c(-5, 5), ylim = c(0, 0.8), legend = legends[[2]])
}

#Calculate posterior predictive distributions of test statistics relating to logit change in Aurelia (O vs C) and potential competitors (A vs C) at each depth from raw data, rather than from manova outputs
#Arguments:
#alldata: data frame containing panel, date, prepost and count for each class
#date: the date we want (last sampling week)
#preorpost: "Pre" or "Post" (before or after scrape
#ppcounts: posterior predictive counts (iterations x panels x taxa)
#niter: number of iterations to plot
#fname: name of file for pdf plot
#Value:
#pdf plot of distribution of each test statistic at each depth: filled version from observed counts, unfilled from posterior predictive
#Example:
#rawcounteffects(alldata = alldata, date = lastdate, preorpost = "Pre")
rawcounteffectspp <- function(alldata, date, preorpost, pp, niter, fname){
    lastweek <- alldata[alldata$date == date & alldata$prepost == preorpost, ]
    logits <- getrawcounteffect(counts = lastweek[, taxa], lastweek = lastweek)
    mycols <- brewer.pal(3, "Dark2")
    legends  <- list(c(expression(Potential~competitors*","~italic(A)~vs~italic(C)), expression(italic(A.~aurita)*","~italic(O)~vs~italic(C))), NULL)
    depths <- c(1, 3)
    pdf(file = fname, width = 7, height = 7)
    par(mfrow = c(2, 1))
    par(mar = c(4.5, 4, 2, 1))
    plotoneresponse(logitchange1 = logits$Aresponse1m, logitchange2 = logits$Oresponse1m, cols = mycols[2:3], panellabel = "a: 1 m", xlim = c(-5, 5), ylim = c(0, 2), legend = legends[[1]])
    for(i in 1:niter){
        simlogits <- getrawcounteffect(counts = ppsample$pcounts[i, , ], lastweek = lastweek)
        lines(density(simlogits$Aresponse1m), col = adjustcolor(mycols[2], 0.1))
        lines(density(simlogits$Oresponse1m), col = adjustcolor(mycols[3], 0.1))
    }
    plotoneresponse(logitchange1 = logits$Aresponse3m, logitchange2 = logits$Oresponse3m, cols = mycols[2:3], panellabel = "b: 3 m", xlim = c(-5, 5), ylim = c(0, 2), legend = legends[[2]])
    for(i in 1:niter){
        simlogits <- getrawcounteffect(counts = ppsample$pcounts[i, , ], lastweek = lastweek)
        lines(density(simlogits$Aresponse3m), col = adjustcolor(mycols[2], 0.1))
        lines(density(simlogits$Oresponse3m), col = adjustcolor(mycols[3], 0.1))
    }
    dev.off()
}

#get logit responses from raw data
#Arguments: counts (panels x taxa)
#Value: list containing logits of Aurelia and amalgamated potential competitors (using add-one pseudocounts)
#and values of the test statistics Oresponse and Aresponse at 1 m and at 3 m
getrawcounteffect <- function(counts, lastweek){
    counts <- counts + 1 #add-one pseudocount
    props <- sweep(counts, 1, rowSums(counts), "/")
    logitAurelia <- logit(props[, 1])
    potentialcomp <- rowSums(props[, 3:5])
    logitcomp <- logit(potentialcomp)
    Oresponse1m <- logitAurelia[lastweek$treatment == "O" & lastweek$depthm == 1] - logitAurelia[lastweek$treatment == "C" & lastweek$depthm == 1]#difference in logit Aurelia between O and C treatments at 1 m
    Oresponse3m <- logitAurelia[lastweek$treatment == "O" & lastweek$depthm == 3] - logitAurelia[lastweek$treatment == "C" & lastweek$depthm == 3]#difference in logit Aurelia between O and C treatments at 3 m
    Aresponse1m <- logitcomp[lastweek$treatment == "O" & lastweek$depthm == 1] - logitcomp[lastweek$treatment == "C" & lastweek$depthm == 1]#difference in logit potential competitors between A and C treatments at 1 m
    Aresponse3m <- logitcomp[lastweek$treatment == "O" & lastweek$depthm == 3] - logitcomp[lastweek$treatment == "C" & lastweek$depthm == 3]#difference in logit potential competitors between A and C treatments at 3 m
    return(list(logitAurelia = logitAurelia, logitcomp = logitcomp, Oresponse1m = Oresponse1m, Oresponse3m = Oresponse3m, Aresponse1m = Aresponse1m, Aresponse3m = Aresponse3m))
}

#ellipse of given Mahalanobis distance (two dimensions)
#Arguments:
#mu: mean vector
#sigma: covariance matrix
#r: target distance
#n: number of points on ellipse
#Value:
#coordinates of points on ellipse
mhellipse <- function(mu, sigma, r, n = 1e2){
    theta <- seq(from = 0,to = 2 * pi, length.out = n)
    x <- cbind(cos(theta), sin(theta))
    dm <- sqrt(mahalanobis(x, center = c(0, 0), cov = sigma))
    xm <- r * x / dm
    xm <- sweep(xm, 2, mu, "+")
    return(xm)
}

#rough plot of posterior mean block effects
#Arguments: manovapredictions: object returned by panelpredict()
#fname: file to plot to
#Value: pdf of ternary plots of block means in first two and last two coordinates of basis
#Note: labelled by block number, but with "x" as label for block 10 (only single-character labels are easy using pch
plotblockpredictions <- function(manovapredictions, fname){
    pdf(file = fname, width = 7, height = 14)
    blockmean <- apply(manovapredictions$blockeffect, c(2, 3), "mean")
    bm12 <- getorthogonalprojection(x = blockmean, coords = c(1, 2), V = V)$xprojsimplex[, 1:3]
    bm34 <- getorthogonalprojection(x = blockmean, coords = c(3, 4), V = V)$xprojsimplex[, 3:5]
    par(mfrow = c(2, 1))
    par(mar = c(3, 3, 1, 3))
    plot(acomp(bm12), pch = c(as.character(1:9), "x"), col = adjustcolor("black", 0.4), labels = c(expression(italic("A. aurita")),"Bare panel", "gm(potential competitors)"))
    text(x = 0, y = 1, "posterior mean block effects", pos = 4) 
    plot(acomp(bm34), pch = c(as.character(1:9), "x"), col = adjustcolor("black", 0.4), labels = c(expression(italic("Botrylloides spp.")), expression(italic("Bugula spp.")), expression(italic("Molgula tubifera"))))
    dev.off()
}


#rough plot of posterior mean panel effects
#Arguments: manovapredictions: object returned by panelpredict()
#fname: file to plot to
#Value: pdf of ternary plots of panel means in first two and last two coordinates of basis
plotpanelpredictions <- function(manovapredictions, fname){
    pdf(file = fname, width = 7, height = 14)
    panelmean <- apply(manovapredictions$paneleffect, c(2, 3), "mean")
    pm12 <- getorthogonalprojection(x = panelmean, coords = c(1, 2), V = V)$xprojsimplex[, 1:3]
    pm34 <- getorthogonalprojection(x = panelmean, coords = c(3, 4), V = V)$xprojsimplex[, 3:5]
    par(mfrow = c(2, 1))
    par(mar = c(3, 3, 1, 3))
    plot(acomp(pm12), pch = 16, col = adjustcolor("black", 0.4), labels = c(expression(italic("A. aurita")),"Bare panel", "gm(potential competitors)"))
    text(x = 0, y = 1, "posterior mean panel effects", pos = 4)
    plot(acomp(pm34), pch = 16, col = adjustcolor("black", 0.4), labels = c(expression(italic("Botrylloides spp.")), expression(italic("Bugula spp.")), expression(italic("Molgula tubifera"))))
    dev.off()
}

#points for qq-plot for multivariate normal
#Arguments:
#y: data (observations x components)
#Sigma: covariance matrix (assumed mean vector 0)
#logscale: if TRUE (default is FALSE), plotted on log scale
#Value: x and y coordinates of points on qq-plot: squared Mahalanobis distances against chi-square quantiles
nqq <- function(y, Sigma, logscale = FALSE){
    k <- dim(y)[2]
    n <- dim(y)[1]
    delta2 <- mahalanobis(y, center = rep(0, k), cov = Sigma)
    delta2 <- rev(delta2)
    if(logscale == TRUE){
        qqpoints <- qqplot(log(qchisq((1:n) / (n + 1), k)), log(delta2), plot.it = FALSE)
    } else {
        qqpoints <- qqplot(qchisq((1:n) / (n + 1), k), delta2, plot.it = FALSE)
    }
    return(qqpoints)
}

#multivariate normal qq plots for a hierarchical part of the manova
#Arguments:
#y: hierarchical variable (iterations x observations x ilr components, where "observations" are either blocks or panels)
#LSigma: Cholesky factor of covariance matrix (iterations x ilr components x ilr components)
#fname: file name of pdf file to which plots will be written
#n: number of MCMC iterations
#Value: pdf fname, n multivariate normal qq plots from posterior distribution of y, and n from simulated multivariate normal
hierarchicalqqplots <- function(y, LSigma, fname, n){
    pdf(file = fname, width = 7, height = 7)
    par(mar = c(4, 4, 2, 2))
    par(mfrow = c(2, 1))
    s <- dim(y)[3]
    m <- dim(y)[2]
    myx <- array(dim = c(m, n))
    myy <- myx
    mysimx <- myx
    mysimy <- myy
    for(i in 1:n){
        simy <- rmvnorm(n = m, mean = rep(0, s), sigma = diag(s))
        qqp <- nqq(y = simy, Sigma = diag(s), logscale = FALSE)
        mysimx[, i] <- qqp$x
        mysimy[, i] <- qqp$y            
        qqp <- nqq(y = y[i, , ], Sigma = LSigma[i, , ] %*% t(LSigma[i, , ]), logscale = FALSE)
        myx[, i] <- qqp$x
        myy[, i] <- qqp$y
    }
    plot(myx[, 1], myy[, 1], xlim = range(c(myx, mysimx)), ylim = range(c(myy, mysimy)), xlab = "theoretical", ylab = "sample", type = "n")
    for(i in 1:n){
        lines(myx[, i], myy[, i], col = adjustcolor("black", 0.1))
    }
    axl <- par("usr")
    text(axl[1] + 0.1 * (axl[2] - axl[1]), axl[4] - 0.1 * (axl[4] - axl[3]), "a: posterior sample", pos = 4)
    plot(myx[, 1], myy[, 1], xlim = range(c(myx, mysimx)), ylim = range(c(myy, mysimy)), xlab = "theoretical", ylab = "sample", type = "n")
    for(i in 1:n){
        lines(mysimx[, i], mysimy[, i], col = adjustcolor("black", 0.1))
    }
    text(axl[1] + 0.1 * (axl[2] - axl[1]), axl[4] - 0.1 * (axl[4] - axl[3]), "b: simulated", pos = 4)
    dev.off()
}


#test quadrature
#Arguments
#s: number of dimensions (needs to be 4 for this test case)
#level: number of quadrature points in each dimension
#muepsilon: mean vector for epsilon
#mudelta: mean vector for delta
#LSigma: Cholesky factor of covariance matrix for epsilon
#LZeta: Cholesky factor of covariance matrix for delta
#counts: an observation from a multinomial
#V: basis matrix for ilr transformation
#mucounts: ilr mean vector for counts
#Value: integral over epsilon and delta of innerfunction(), where both epsilon and delta are multivariate normal. Test version of innerfunction is a multinomial probability for counts, with probability vector ilrInv(epsilon + delta + mucounts)
#Example: testquad(s = 4, level = 2, muepsilon = zpanel[1, 1, ], mudelta = zblock[1, 1, ], LSigma = LSigma[1, , ], LZeta = LZeta[1, , ], counts = manovainputs$counts[1, ], V = V, mucounts = manovapredictions$fixedeffect[1, 1, ])
testquad <- function(s, level, muepsilon, mudelta, LSigma, LZeta, counts, V, mucounts){
    epsilonGrid <- createNIGrid(dim = s, type = "GHe", level = level)
    deltaGrid <- createNIGrid(dim = s, type = "GHe", level = level)
    Sigma <- LSigma %*% t(LSigma)
    Zeta <- LZeta %*% t(LZeta)
    rescale(epsilonGrid, m = muepsilon, C = Sigma, dec.type = 2)
    rescale(deltaGrid, m = mudelta, C = Zeta, dec.type = 2)
    innerfunction <- function(epsilon, delta, counts, V, mucounts){
        mu <- sweep(epsilon + delta, 2, mucounts, "+")
        prob  <- ilrInv(mu, V = V)
        multinomprob <- function(prob, counts){dmultinom(x = counts, prob = prob)}
        return(apply(prob, 1, FUN = multinomprob, counts = counts))
    }
    
    epsilonfun <- function(x, muepsilon, Sigma, delta, counts, V, mucounts){
        dmvnorm(x, mean = muepsilon, sigma = Sigma) * innerfunction(epsilon = x, delta = delta, counts = counts, V = V, mucounts = mucounts)
    }

    deltafun <- function(x, muepsilon, mudelta, Sigma, Zeta, epsilonGrid, counts, V, mucounts){
        epsilonintegral <- dmvnorm(x, mean = mudelta, sigma = Zeta) * quadrature(epsilonfun, grid = epsilonGrid, mu = muepsilon, Sigma = Sigma, delta = x, counts = counts, V = V, mucounts = mucounts)
    }

    print(quadrature(deltafun, grid = deltaGrid, muepsilon = muepsilon, mudelta = mudelta, Sigma = Sigma, Zeta = Zeta, epsilonGrid = epsilonGrid, counts = counts, V = V, mucounts = mucounts))

 }

#test Monte Carlo integration
#Arguments
#muepsilon: mean vector for epsilon
#mudelta: mean vector for delta
#LSigma: Cholesky factor of covariance matrix for epsilon
#LZeta: Cholesky factor of covariance matrix for delta
#counts: an observation from a multinomial
#V: basis matrix for ilr transformation
#mucounts: ilr mean vector for counts
#Value: integral over epsilon and delta of multinomial probability for counts, with probability vector ilrInv(epsilon + delta + mucounts)
#Example: testquad(muepsilon = zpanel[1, 1, ], mudelta = zblock[1, 1, ], LSigma = LSigma[1, , ], LZeta = LZeta[1, , ], counts = manovainputs$counts[1, ], V = V, mucounts = manovapredictions$fixedeffect[1, 1, ], niter = 1e3)
testMonteCarlo <- function(muepsilon, mudelta, LSigma, LZeta, counts, V, mucounts, niter){
    y <- numeric(niter)
    Sigma <- LSigma %*% t(LSigma)
    Zeta <- LZeta %*% t(LZeta)
    for(i in 1:niter){
        epsilon  <- rmvnorm(n = 1, mean = muepsilon, sigma = Sigma)
        delta  <- rmvnorm(n = 1, mean = mudelta, sigma = Zeta)
        mu <- epsilon + delta + mucounts
        prob <- unclass(ilrInv(mu, V = V))
        y[i] <- dmultinom(x = counts, prob = prob)
    }
    return(mean(y))
}

#set up leave-one-out data files for Stan
#Arguments:
#alldata: data frame from alldata.txt
#date: the date we want to analyse
#preorpost: use the pre- or post-scrape data from this week?
#taxa: names of the taxa to include in the response vector
#tVinv: generalized inverse of transpose of basis matrix for ilr coordinates
#leaveoneoutname: base for name of file to which input data for stan will be written
#Value:
#writes to standatafilename using stan_rdump and returns list containing
#d1 to d5: vectors of values of dummy variables representing treatment contrasts for each panel
#counts: array of counts by panel for the taxa we include
#blockindex: vector labelling panels by block
writeleaveoneout <- function(alldata, date, preorpost, taxa, tVinv, leaveoneoutname){
    ntaxa  <- length(taxa)
    lastweek <- alldata[alldata$date == date & alldata$prepost == preorpost, ]
    npanels <- dim(lastweek)[1]
    d1all <- numeric(npanels)
    d2all <- d1all
    d3all <- d1all
    d4all <- d1all
    d5all <- d1all
    for(i in 1:npanels){
        contrasts <- buildcontrastbasis(depth = alldata$depthm[i], treatment = alldata$treatment[i])
        d1all[i] <- contrasts$d1
        d2all[i] <- contrasts$d2
        d3all[i] <- contrasts$d3
        d4all[i] <- contrasts$d4
        d5all[i] <- contrasts$d5
    }
    countsall  <- lastweek[, taxa]
    blockindexall = lastweek$block
    nblocksall <- length(unique(lastweek$block))
    for(i in 1:nblocksall){
        minusi <- which(blockindexall != i)
        blockindex <- blockindexall[minusi]
        d1 <- d1all[minusi]
        d2 <- d2all[minusi]
        d3 <- d3all[minusi]
        d4 <- d4all[minusi]
        d5 <- d5all[minusi]
        counts <- countsall[minusi, ]
        nblocks <- nblocksall #one of the blocks excluded: no data to estimate parameters, but still need it there for indexing in stan code
        npanels <- dim(counts)[1]
        fnamei <- paste(leaveoneoutname, "_", i, ".txt", sep = "")
        paneldatai <- c("d1", "d2", "d3", "d4", "d5", "counts", "tVinv", "ntaxa", "npanels", "nblocks", "blockindex")
        stan_rdump(list = paneldatai, file = fnamei, quiet = FALSE, append = FALSE)
    }
}

#read data from a set of .csv Stan output files and compute leave-one-block-out cross-validation score
#Arguments:
#leaveoneoutpath: path to directory containing stan output files
#csvfiles: character vector of file names
#mciter: number of Monte Carlo iterations to use in Monte Carlo integration over block and panel effects
#niter: number of Stan iterations run
#model: which model ("interaction", "nointeraction", "noscrape" or "nodepth)
#V: ilr basis matrix
#manovainputs: object returned by aureliamanovasetup()
#Value:
#llmcontrib: marginal log likelihood distribution for lth block, given parameters estimated from all other blocks
getcv <- function(leaveoneoutpath, csvfiles, l, mciter, niter, model, V, manovainputs){
    gc()
    manovadata <- read_stan_csv(csvfiles = paste(leaveoneoutpath, csvfiles, sep = ""))
    mu <- extract(manovadata, pars = "mu")[[1]]
    LSigma <- extract(manovadata, pars = "LSigma")[[1]]
    LZeta <- extract(manovadata, pars = "LZeta")[[1]]
    zpanel <- extract(manovadata, pars = "zpanel")[[1]]
    zblock <- extract(manovadata, pars = "zblock")[[1]]    
    if(model == "interaction"){
        alpha <- extract(manovadata, pars = "alpha")[[1]]
        beta1 <- extract(manovadata, pars = "beta1")[[1]]
        beta2 <- extract(manovadata, pars = "beta2")[[1]]
        gamma1 <- extract(manovadata, pars = "gamma1")[[1]]
        gamma2 <- extract(manovadata, pars = "gamma2")[[1]]    
    } else if (model == "nointeraction"){#set interaction parameters to zero
        alpha <- extract(manovadata, pars = "alpha")[[1]]
        beta1 <- extract(manovadata, pars = "beta1")[[1]]
        beta2 <- extract(manovadata, pars = "beta2")[[1]]
        gamma1 <- array(0, dim = dim(mu))
        gamma2 <- array(0, dim = dim(mu))
    } else if(model == "noscrape"){#set interaction and scrape parameters to zero
        alpha <- extract(manovadata, pars = "alpha")[[1]]
        beta1 <- array(0, dim = dim(mu))
        beta2 <- array(0, dim = dim(mu))
        gamma1 <- array(0, dim = dim(mu))
        gamma2 <- array(0, dim = dim(mu))
    } else if (model == "nodepth"){#set interaction and depth parameters to zero
        alpha <- array(0, dim = dim(mu))
        beta1 <- extract(manovadata, pars = "beta1")[[1]]
        beta2 <- extract(manovadata, pars = "beta2")[[1]]
        gamma1 <- array(0, dim = dim(mu))
        gamma2 <- array(0, dim = dim(mu))
    } else {
        stop(paste("getcv(): model ", model, "not implemented", sep = ""))
    }
    rm(manovadata)
    gc()
    s <- dim(V)[2]
    llmcontrib <- numeric(niter)
    for(i in 1:niter){#loop over Stan iterations
        delta <- rmvnorm(n = mciter, mean = rep(0, s), sigma = LZeta[i, , ] %*% t(LZeta[i, , ])) #sample from distribution of delta for a new block, given the Zeta that was drawn in this Stan iteration
        mymu <- matrix(rep(mu[i, ], mciter), nrow = mciter, byrow = TRUE)
        myalpha <- matrix(rep(alpha[i, ], mciter), nrow = mciter, byrow = TRUE)
        mybeta1 <- matrix(rep(beta1[i, ], mciter), nrow = mciter, byrow = TRUE)
        mybeta2 <- matrix(rep(beta2[i, ], mciter), nrow = mciter, byrow = TRUE)
        mygamma1 <- matrix(rep(gamma1[i, ], mciter), nrow = mciter, byrow = TRUE)
        mygamma2 <- matrix(rep(gamma2[i, ], mciter), nrow = mciter, byrow = TRUE)
        mycounts <- manovainputs$counts[manovainputs$blockindex == l, ]
        myd1 <- manovainputs$d1[manovainputs$blockindex == l]
        myd2 <- manovainputs$d2[manovainputs$blockindex == l]
        myd3 <- manovainputs$d3[manovainputs$blockindex == l]
        myd4 <- manovainputs$d4[manovainputs$blockindex == l]
        myd5 <- manovainputs$d5[manovainputs$blockindex == l]
        llc <- array(dim = c(length(myd1), mciter))
        for(j in 1:length(myd1)){#each panel in new block
            epsilon <- rmvnorm(n = mciter, mean = rep(0, s), sigma = LSigma[i, , ] %*% t(LSigma[i, , ])) #sample from distribution of epsilon for a new panel within block, given the Sigma that was drawn in this Stan iteration
            llc[j, ] <- llmcontrib[i] + mean(likelihoodcontribution(epsilon = epsilon, delta = delta, mu = mymu, alpha = myalpha, beta1 = mybeta1, beta2 = mybeta2, gamma1 = mygamma1, gamma2 = mygamma2, V = V, panel = j, counts = mycounts, d1 = myd1, d2 = myd2, d3 = myd3, d4 = myd4, d5 = myd5, uselog = TRUE))
        }
        llmcontrib[i] <- log(mean(exp(colSums(llc))))
    }
    return(llmcontrib)
}


#contribution to likelihood or log likelihood for a single panel within a block over all iterations
#Arguments:
#epsilon: s x iterations vector of panel effects (we will be integrating over this)
#delta: s x iterations vector of block effects (we will be integrating over this)
#parameter arrays (each iterations x ilr components):
#mu: overall mean
#alpha: depth contrast
#beta1, beta2: treatment contrasts
#gamma1, gamma2: interaction contrasts
#V: ilr basis matrix
#counts: matrix of observed counts (panels x taxa)
#d1 to d5: dummy variables associated with alpha, beta1, beta2, gamma1, gamma2 respectively
#uselog (logical): return log likelihood or likelihood?
#Value:
#contribution to likelihood or log likelihood for given panel in block in each iteration
likelihoodcontribution <- function(epsilon, delta, mu, alpha, beta1, beta2, gamma1, gamma2, V, panel, counts, d1, d2, d3, d4, d5, uselog){
    x <- mu + alpha * d1[panel] + beta1 * d2[panel] + beta2 * d3[panel] + gamma1 * d4[panel] + gamma2 * d5[panel] + epsilon + delta 
    n <- counts[panel, ]
    p <- ilrInv(x, V = V)
    multinomprob <- function(prob, counts, uselog){dmultinom(x = counts, prob = prob, log = uselog)}
    f <- apply(p, 1, FUN = multinomprob, counts = n, uselog = uselog)
    return(f)

#C++ version faster, but Rstan doesn't currently like having other compiled code. Needs Vt to be transpose of V    
#    p <- cilrinv(x = x, Vt = Vt) #defined in marginal.cpp
#    if(uselog == FALSE){
#        f <- cmultinomialmatrix(p = p, n = n) #defined in marginal.cpp
#    } else {
#        f <- cmultinomialmatrixlog(p = p, n = n) #defined in marginal.cpp
#    }
#    return(f)
}

#function for using k-fold cross-validation output with loo_compare() from package loo. Modified from https://github.com/stan-dev/stancon_talks/blob/master/2017/Contributed-Talks/07_nicenboim/kfold.Rmd
#returns an object of class c("kfold", "loo"), which can be used with loo_compare()
#Arguments:
#log_lik_heldout: array (stan iterations x blocks) of leave-one-block-out cross validation scores 
#Value: object of class c("kfold", "loo"): first item contains elpd_kfold, columns Estimate, SE; second item contains elpd_kfold for each block
kfold <- function(log_lik_heldout){
  logColMeansExp <- function(x){
    # should be more stable than log(colMeans(exp(x)))
    S <- nrow(x)
    colLogSumExps(x) - log(S)
  }
  # See equation (20) of @VehtariEtAl2016
  pointwise <- matrix(logColMeansExp(log_lik_heldout), ncol = 1)
  colnames(pointwise) <- "elpd_kfold"
  # See equation (21) of @VehtariEtAl2016
  elpd_kfold <- sum(pointwise)
  se_elpd_kfold <- sqrt(ncol(log_lik_heldout) * var(pointwise))
  estimates <- matrix(c(elpd_kfold, se_elpd_kfold), byrow = TRUE, nrow = 1, ncol = 2)
  rownames(estimates) <- "elpd_kfold"
  colnames(estimates) <- c("Estimate", "SE")
  out <- list(estimates = estimates, pointwise = pointwise)
  return(structure(out, dims = dim(out$pointwise), class = c("kfold", "loo")))
}

#get posterior mean and 95% highest density region for a vector-valued parameter
#Arguments: x (iterations x components)
#lambda: Box-Cox transformation parameter between 0 and 1 (default 1). Values less than 1 mean we get positive density estimate only on the positive real line
#Value: list containing mean and an array in which rows are lower and upper bounds of 95% HDR region for each component of x
posteriorsummaryvector <- function(x, lambda = 1){
    if(length(dim(x)) == 2){#x really is a vector-valued parameter
        xmean <- colMeans(x)
        s <- dim(x)[2]
        myhdr <- function(y){
            hdr(y, prob = 95, lambda = lambda)$hdr
        }
        xhdr <- apply(x, 2, FUN = myhdr)
    } else {#x is actually a scalar-valued parameter
        xmean <- mean(x)
        xhdr <- hdr(x, prob = 95, lambda = lambda)$hdr
    }
    return(list(mean = xmean, hdr = xhdr))
}

#extract posterior mean and hdr region for a parameter in a form we can print using xtable
#Arguments: x: object returned by posteriorsummaryvector
#i: index of the component within x for which we want summary
#format: optional format string (default "%.2f")
#Value: a string giving posterior mean and all the intervals making up the credible region
extractposteriorsummary <- function(x, i, format = "%.2f"){
    if(class(x$hdr)[1] == "matrix"){
        if(dim(x$hdr)[1] == 1){#parameter is really a scalar
            psumm <- c(sprintf(paste(format, " (", format, ", ", format, ")\n", sep = ""), x$mean[i], x$hdr[1], x$hdr[2]))
            } else {
                psumm <- c(sprintf(paste(format, " (", format, ", ", format, ")\n", sep = ""), x$mean[i], x$hdr[1, i], x$hdr[2, i]))
            }
    } else {#hdr is more than one disjoint interval, in list
        nint <- length(x$hdr[[i]]) / 2
        psumm <- sprintf(format, x$mean[i])
        for(j in 1:nint){
            psumm <- paste(psumm, sprintf(paste("(", format, ", ", format, ")", sep = ""), x$hdr[[i]][2 * j - 1], x$hdr[[i]][2 * j]), sep = " ")
        }
        psumm <- paste(psumm, "\n", sep = "")
    }
    return(psumm)
}

#produce a LaTeX table summarizing posterior distributions of manova parameters
#Arguments:
#mu, alpha, beta1, beta2, gamma1, gamma2: arrays of parameters (iterations x ilr components): mean, depth effect, treatment contrast 1, treatment contrast2, interaction contrast 1, interaction contrast 2
#LSigma, LZeta: Cholesky factors of panel and block covariance matrices respectively (iterations x ilr components x ilr components)
#fname: name of file to which LaTeX output should be written
#includeinteraction: logical: include the interaction parameters gamma1, gamma2 in table?
#Value:
#LaTeX code for table of posterior means and 95% HPD intervals, written to fname
#Note: will currently only work for 4 ilr coordinates: indices of covariance matrices hard coded
printmanovaparameters <- function(mu, alpha, beta1, beta2, gamma1, gamma2, LSigma, LZeta, fname, includeinteraction){
    musummary <- posteriorsummaryvector(mu)
    alphasummary <- posteriorsummaryvector(alpha)
    beta1summary <- posteriorsummaryvector(beta1)
    beta2summary <- posteriorsummaryvector(beta2)
    mfun <- function(x){x %*% t(x)}
    Sigma <- t(apply(LSigma, 1, mfun)[c(1, 5, 6, 9, 10, 11, 13, 14, 15, 16), ])#gets the lower triangle of Sigma by rows: will only work for s = 4
    Zeta <- t(apply(LZeta, 1, mfun)[c(1, 5, 6, 9, 10, 11, 13, 14, 15, 16), ])
    sigmasummary <- posteriorsummaryvector(Sigma)
    zetasummary <- posteriorsummaryvector(Zeta)
    if(includeinteraction == TRUE){
        gamma1summary <- posteriorsummaryvector(gamma1)
        gamma2summary <- posteriorsummaryvector(gamma2)
    }
    s <- dim(mu)[2]
    if(includeinteraction == TRUE){
        mytable <- array(dim = c(14, s)) #will only work for s = 4
        rownames(mytable) <- c("$\\bm \\mu$", "$\\bm \\alpha$", "$\\bm \\beta_1$", "$\\bm \\beta_2$", "$\\bm \\gamma_1$", "$\\bm \\gamma_2$", "$\\bm \\sigma_1$", "$\\bm \\sigma_2$", "$\\bm \\sigma_3$", "$\\bm \\sigma_4$", "$\\bm \\zeta_1$", "$\\bm \\zeta_2$", "$\\bm \\zeta_3$", "$\\bm \\zeta_4$") #will only work for s = 4
    } else {
        mytable <- array(dim = c(12, s))
        rownames(mytable) <- c("$\\bm \\mu$", "$\\bm \\alpha$", "$\\bm \\beta_1$", "$\\bm \\beta_2$", "$\\bm \\sigma_1$", "$\\bm \\sigma_2$", "$\\bm \\sigma_3$", "$\\bm \\sigma_4$", "$\\bm \\zeta_1$", "$\\bm \\zeta_2$", "$\\bm \\zeta_3$", "$\\bm \\zeta_4$")
    }
    mindex <- 1
    for(i in 1:s){
        mytable[1, i] <- extractposteriorsummary(musummary, i)
        mytable[2, i] <- extractposteriorsummary(alphasummary, i)
        mytable[3, i] <- extractposteriorsummary(beta1summary, i)
        mytable[4, i] <- extractposteriorsummary(beta2summary, i)
        rowindex <- 4
        if(includeinteraction == TRUE){
            mytable[5, i] <- extractposteriorsummary(gamma1summary, i)
            mytable[6, i] <- extractposteriorsummary(gamma2summary, i)
            rowindex <- 6
        }
        for(j in 1:i){
            mytable[rowindex + i, j] <- extractposteriorsummary(sigmasummary, mindex)
            mytable[rowindex + i + s, j] <- extractposteriorsummary(zetasummary, mindex)
            mindex <- mindex + 1
        }
    }
    xt <- xtable(mytable)
    align(xt) <- c("r", rep("p {3cm}", s))
    print(xt, sanitize.text.function = function(x){x}, file = fname)
}

#print out which panels had identical pre and post counts and were not controls, and which panels were controls but had different pre and post counts
#Arguments: data frame alldata
#Value: just prints panels as above
checkidentical <- function(alldata){
    print("Identical pre/post and not C:")
    for(i in unique(alldata$date)){
        for(j in unique(alldata$panel)){
            predata <- alldata[alldata$date == i & alldata$panel == j & alldata$prepost == "Pre", ]
            postdata <- alldata[alldata$date == i & alldata$panel == j & alldata$prepost == "Post", ]
            if(max(abs(predata[, 7:15] - postdata[, 7:15])) == 0 & predata$treatment != "C"){
                print(predata[c(3, 4, 5, 7:15)])
            }
        }
    }

        print("Different pre/post and C:")
    for(i in unique(alldata$date)){
        for(j in unique(alldata$panel)){
            predata <- alldata[alldata$date == i & alldata$panel == j & alldata$prepost == "Pre", ]
            postdata <- alldata[alldata$date == i & alldata$panel == j & alldata$prepost == "Post", ]
            if(max(abs(predata[, 7:15] - postdata[, 7:15])) != 0 & predata$treatment == "C"){
                print(predata[c(3, 4, 5)])
            }
        }
    }
}

#plot of sample time series of A. aurita, bare panel, sum of potential competitors on each panel
#Arguments:
#alldata: data frame including date, treatment, depthm, counts for each taxon
#potentialcompetitors: names of taxa to include in the amalgamated "potential competitors" part (default is Botrylloides.spp. Bugula.spp., Molgula.tubifera)
#dayzero: day on which panels went into water e.g. as.Date("2019-07-30")
#fname: name of pdf file to which to write plot
#rhoC1summary, ..., rhoA3summary: summaries of posterior distributions predicted from ODE model for each treatment and depth combination (output from typicalrhoset())
#tC1dates, ..., tA3dates: observation dates for typical panels of each combination
#showobservations (logical, default TRUE): plot the observations
#showpredictions (logical, default TRUE): plot the predictions
#predlwd (default 1): line width for predictions
#obsalpha (default 0.3): transparency alpha for observations
#Value: pdf plot of time series for each panel for A. aurita, bare panel, potential competitors, coloured by treatment. Solid lines for 3 m, dashed lines for 1 m
#Note: circle added (hard coded) to indicate treatment wrongly applied to panel 2 on second sampling date
timeseriesplot <- function(alldata, potentialcompetitors = c("Botrylloides.spp.", "Bugula.spp.", "Molgula.tubifera"), dayzero, fname, rhoC1summary, rhoC3summary, rhoO1summary, rhoO3summary, rhoA1summary, rhoA3summary, typicalpanels, dynamicdata, tC1dates, tC3dates, tO1dates, tO3dates, tA1dates, tA3dates, showobservations = TRUE, showpredictions = TRUE, predlwd = 1, obsalpha = 0.3){
    pdf(file = fname, width = 7, height = 7)
    par(mfrow = c(3, 1))
    par(mar = c(4, 4, 2, 2))
    mycolors <- brewer.pal(3, "Dark2")[c(2, 1, 3)]
    mylty <- c("dashed", NA, "solid")
    potentialcomp <- rowSums(alldata[, potentialcompetitors])
    panelcomp <- acomp(cbind(alldata$Aurelia.aurita.on.panel, alldata$Bare.Panel, potentialcomp))
    date <- as.Date(alldata$date)
    plot(c(dayzero, date[alldata$panel == 1]), c(0, panelcomp[alldata$panel == 1, 1]), ylim =c(0, 1), xlab = "date", ylab = expression(italic(A.~aurita)), type = "n", xaxt = "n")
    axis(side = 1, at = c(dayzero, unique(date)), labels = c(dayzero, unique(date)))
    legend("topright", bty = "n", legend = c(expression(paste(1~m, ",", ~italic(C))), expression(paste(3~m, ",", ~italic(C))), expression(paste(1~m, ",", ~italic(A))), expression(paste(3~m, ",", ~italic(A))), expression(paste(1~m, ",", ~italic(O))), expression(paste(3~m, ",", ~italic(O))), "observed", "predicted"), col = c(mycolors[c(2, 2, 1, 1, 3, 3)], adjustcolor("black", obsalpha), "black"), lty = c(rep(mylty[c(1, 3)], 3), "solid", "solid"), pch = NA, lwd = c(1, 1, 1, 1, 1, 1, 1, predlwd))
    axl <- par("usr")
    text(axl[1] + 0.05 * (axl[2] - axl[1]), axl[4] - 0.2 * (axl[4] - axl[3]), "a")
    if(showobservations == TRUE){
        for(i in unique(alldata$panel)){
         lines(c(dayzero, date[alldata$panel == i]), c(0, panelcomp[alldata$panel == i, 1]), col = adjustcolor(mycolors[alldata$treatment[i]], obsalpha), lty = mylty[alldata$depthm[i]])
        }
        points(as.Date("2019-08-13"), panelcomp[alldata$panel == 2, 1][2], col = adjustcolor(mycolors[2], 1))
    }
    if(showpredictions == TRUE){
        drawtimepolygon(dates = tC1dates, summary = rhoC1summary, part = 1, lty = "dashed", color = mycolors[2], lwd = predlwd)
        drawtimepolygon(dates = tC3dates, summary = rhoC3summary, part = 1, lty = "solid", color = mycolors[2], lwd = predlwd)
        drawtimepolygon(dates = tO1dates, summary = rhoO1summary, part = 1, lty = "dashed", color = mycolors[3], lwd = predlwd)
        drawtimepolygon(dates = tO3dates, summary = rhoO3summary, part = 1, lty = "solid", color = mycolors[3], lwd = predlwd)
        drawtimepolygon(dates = tA1dates, summary = rhoA1summary, part = 1, lty = "dashed", color = mycolors[1], lwd = predlwd)
        drawtimepolygon(dates = tA3dates, summary = rhoA3summary, part = 1, lty = "solid", color = mycolors[1], lwd = predlwd)
    }
    
    plot(c(dayzero, date[alldata$panel == 1]), c(1, panelcomp[alldata$panel == 1, 2]), ylim =c(0, 1), xlab = "date", ylab = "bare panel", type = "n", xaxt = "n")
    axis(side = 1, at = c(dayzero, unique(date)), labels = c(dayzero, unique(date)))
    text(axl[1] + 0.05 * (axl[2] - axl[1]), axl[4] - 0.2 * (axl[4] - axl[3]), "b")
    if(showobservations == TRUE){
        for(i in unique(alldata$panel)){
            lines(c(dayzero, date[alldata$panel == i]), c(1, panelcomp[alldata$panel == i, 2]), col = adjustcolor(mycolors[alldata$treatment[i]], obsalpha), lty = mylty[alldata$depthm[i]])
        }
        points(as.Date("2019-08-13"), panelcomp[alldata$panel == 2, 2][2], col = adjustcolor(mycolors[2], 1))
    }
    if(showpredictions == TRUE){
        drawtimepolygon(dates = tC1dates, summary = rhoC1summary, part = 2, lty = "dashed", color = mycolors[2], lwd = predlwd)
        drawtimepolygon(dates = tC3dates, summary = rhoC3summary, part = 2, lty = "solid", color = mycolors[2], lwd = predlwd)
        drawtimepolygon(dates = tO1dates, summary = rhoO1summary, part = 2, lty = "dashed", color = mycolors[3], lwd = predlwd)
        drawtimepolygon(dates = tO3dates, summary = rhoO3summary, part = 2, lty = "solid", color = mycolors[3], lwd = predlwd)
        drawtimepolygon(dates = tA1dates, summary = rhoA1summary, part = 2, lty = "dashed", color = mycolors[1], lwd = predlwd)
        drawtimepolygon(dates = tA3dates, summary = rhoA3summary, part = 2, lty = "solid", color = mycolors[1], lwd = predlwd)
    }

    plot(c(dayzero, date[alldata$panel == 1]), c(0, panelcomp[alldata$panel == 1, 3]), ylim =c(0, 1), xlab = "date", ylab = "potential competitors", type = "n", xaxt = "n")
    axis(side = 1, at = c(dayzero, unique(date)), labels = c(dayzero, unique(date)))
    text(axl[1] + 0.05 * (axl[2] - axl[1]), axl[4] - 0.2 * (axl[4] - axl[3]), "c")

    if(showobservations == TRUE){
        for(i in unique(alldata$panel)){
            lines(c(dayzero, date[alldata$panel == i]), c(0, panelcomp[alldata$panel == i, 3]), col = adjustcolor(mycolors[alldata$treatment[i]], obsalpha), lty = mylty[alldata$depthm[i]])
        }
        points(as.Date("2019-08-13"), panelcomp[alldata$panel == 2, 3][2], col = adjustcolor(mycolors[2], 1))
    }
    if(showpredictions == TRUE){
        drawtimepolygon(dates = tC1dates, summary = rhoC1summary, part = 3, lty = "dashed", color = mycolors[2], lwd = predlwd)
        drawtimepolygon(dates = tC3dates, summary = rhoC3summary, part = 3, lty = "solid", color = mycolors[2], lwd = predlwd)
        drawtimepolygon(dates = tO1dates, summary = rhoO1summary, part = 3, lty = "dashed", color = mycolors[3], lwd = predlwd)
        drawtimepolygon(dates = tO3dates, summary = rhoO3summary, part = 3, lty = "solid", color = mycolors[3], lwd = predlwd)
        drawtimepolygon(dates = tA1dates, summary = rhoA1summary, part = 3, lty = "dashed", color = mycolors[1], lwd = predlwd)
        drawtimepolygon(dates = tA3dates, summary = rhoA3summary, part = 3, lty = "solid", color = mycolors[1], lwd = predlwd)
    }
    dev.off()
}

#draw polygon representing credible band, with line representing metric centre, on a time series plot
#Arguments:
#dates: observation times, including zero time
#summary: object returned by getrhosummary
#part: which part of composition? If part is 2 (bare panel), append initial value 1, otherwise append initial value 0
#lty: line style for metric centre
#color: color for line and polygon
#lwd (default 1): line width for metric centre
#Value: adds polygon and line to time series plot
drawtimepolygon <- function(dates, summary, part, lty, color, lwd = 1){
    if(part == 2) initial = 1 else initial = 0
    polygon(c(dates, rev(dates)), c(initial, summary$simplexhdr[part, 1, ], rev(summary$simplexhdr[part, 2, ]), initial), col = adjustcolor(color, 0.4), border = NA)
    lines(dates, c(initial, summary$metriccentre[, part]), col = color, lty = lty, lwd = lwd)
}

#build data set for ODE/SDE model in Stan: drop post rows in which no treatment was applied, work out time blocks for ODE solution
#Arguments:
#alldata: data frame written by compiledata(): needs to be in time order
#potentialcompetitors: names of taxa to include in the amalgamated "potentialcomp" part (default is Botrylloides.spp. Bugula.spp., Molgula.tubifera)
#treatmentappliedfile: name of tab-separated text file containing variables panel, date, applied: applied is blank if no treatment applied, otherwise A or O (and records what was actually done: e.g. one case where the wrong treatment was applied to a panel on one data is recorded here
#taxaforstan: names of taxa to include in counts written to Stan file (default: Aurelia.aurita.on.panel, Bare.Panel, potentialcomp)
#standatafilename: name of file to which input data for stan will be written
#Value:
#list containing:
#dynamicdata: data frame with variables as in alldata, but post rows with no treatment applied removed, and new variables treatmentapplied (from treatmentappliedfile, NA in pre rows or post rows where no treatment applied) and potentialcomp (sum of the counts in variables listed in potentialcompetitors)
#sumnt: sum of values in nt
#counts: counts for the taxa in taxaforstan
#npanels: number of panels
#ntaxa: number of taxa included for stan (after amalgamation)
#times: time in weeks
#tb: output of maketimeblocks: number of time blocks per panel, number of times per time block in panel, time indices in each time block for each panel
#depthcode: coded depth, 1 for 1 m, 2 for 3 m
datafordynamicmodel <- function(alldata, potentialcompetitors = c("Botrylloides.spp.", "Bugula.spp.", "Molgula.tubifera"), treatmentappliedfile, taxaforstan = c("Aurelia.aurita.on.panel", "Bare.Panel", "potentialcomp"), standatafilename){
    npanels <- length(unique(alldata$panel))
    depthcode <- numeric(npanels)
    depthcode[alldata$depthm[1:npanels] == 1]  <- 1 #assumes one observation from each panel comes first
    depthcode[alldata$depthm[1:npanels] == 3]  <- 2
    alldata <- alldata[order(alldata$panel), ] #have assumed already ordered by date and prepost
    treatmentapplied <- read.table(file = treatmentappliedfile, header = TRUE, sep = "\t")
    treatmentapplied$applied <- factor(treatmentapplied$applied)
    treatmentapplied <- treatmentapplied[order(treatmentapplied$panel), ]
    alldata$treatmentapplied <- factor(NA, levels = levels(treatmentapplied$applied))
    alldata$treatmentapplied[alldata$prepost == "Post"] <- treatmentapplied$applied
    keep <- alldata$prepost == "Pre" | (alldata$prepost == "Post" & alldata$treatmentapplied != "")
    dynamicdata <- alldata[keep, ]
    dynamicdata$potentialcomp <- rowSums(dynamicdata[, potentialcompetitors])
    nt <- numeric(npanels)
    for(i in 1:npanels){
        nt[i] <- sum(dynamicdata$panel == i)
    }
    sumnt <- sum(nt)
    counts <- dynamicdata[, taxaforstan]
    ntaxa = length(taxaforstan)
    times <- as.numeric(factor(dynamicdata$date))
    tb <- maketimeblocks(dynamicdata = dynamicdata, npanels = npanels, times = times)
    ntimeblocks <- tb$ntimeblocks
    ntimesinblock <- tb$ntimesinblock
    timeblocks <- tb$timeblocks
    maxtimeblocks <- tb$maxtimeblocks
    maxtimes <- tb$maxtimes
    treatmentapplied <- as.numeric(dynamicdata$treatmentapplied)
    treatmentapplied[is.na(treatmentapplied)] <- 0
    ddata <- c("counts", "ntaxa", "npanels", "nt", "sumnt", "times", "ntimeblocks", "ntimesinblock", "timeblocks", "maxtimeblocks", "maxtimes", "treatmentapplied", "depthcode")
    stan_rdump(list = ddata, file = standatafilename, quiet = FALSE, append = FALSE)
    return(list(dynamicdata = dynamicdata, nt = nt, sumnt = sumnt, counts = counts, npanels = npanels, ntaxa = ntaxa, times = times, tb = tb, treatmentapplied = treatmentapplied, depthcode = depthcode))
}

#how many time blocks for a given panel?
#Arguments:
#dynamicdata: data frame containing variable panel (with one row per observation, stacked by panels and weeks)
#panel: panel index
#times: vector of times (stacked by panel and week)
#Value: number of time blocks needed in ODE model for this panel
counttimeblocks <- function(dynamicdata, panel, times){
    maxtime <- max(times)
    ptimes <- times[dynamicdata$panel == panel]
    nmaxtime <- sum(ptimes == maxtime) #if last time appears twice because of treatment, we don't need an extra time block for it
    return(length(ptimes) - length(unique(ptimes)) + 2 - nmaxtime)
}

#build array of time blocks for ODE model
#Arguments:
#dynamicdata: data frame containing variable panel (with one row per observation, stacked by panels and weeks)
#npanels: number of panels
#times: vector of times (stacked by panel and week)
#Value:
#ntimeblocks (vector, one entry per panel): number of time blocks for panel
#timeblocks: array (panels x time blocks x max possible number of times). The layer in row i, column j gives the observation times for the jth time block in panel i (unused values are -1: note initial time is not included in this)
#ntimesinblock: array (panels x time blocks): row i, column j gives number of time points in jth time block for panel i. Unused values are -1.
#maxtimeblocks: max number of time blocks per panel
#maxtimes: max possible number of observation times per time block
maketimeblocks <- function(dynamicdata, npanels, times){
    ntimeblocks <- numeric(npanels)
    for(i in 1:npanels){
        ntimeblocks[i] <- counttimeblocks(dynamicdata = dynamicdata, panel = i, times = times)
    }
    maxtimeblocks <- max(ntimeblocks)
    maxtimes <- length(unique(times))
    timeblocks <- array(-1, dim = c(npanels, maxtimeblocks, maxtimes))
    ntimesinblock <- array(-1, dim = c(npanels, maxtimeblocks))
    for(i in 1:npanels){
        for(j in 1:ntimeblocks[i]){
            tb <- gettimeblock(dynamicdata = dynamicdata, panel = i, times = times, j = j)
            timeblocks[i, j, 1:length(tb)] <- tb
            ntimesinblock[i, j] <- length(tb)
            
        }
    }
    return(list(ntimeblocks = ntimeblocks, timeblocks = timeblocks, ntimesinblock = ntimesinblock, maxtimeblocks = maxtimeblocks, maxtimes = maxtimes))
}

#get time points in jth time block for ODE model
#Arguments:
#dynamicdata: data frame containing variable panel (with one row per observation, stacked by panels and weeks)
#panel: panel index
#times: vector of times (stacked by panel and week)
#j: index of time block we want
#Value: vector of times in weeks in jth time block for given panel
gettimeblock <- function(dynamicdata, panel, times, j){
    ptimes <- times[dynamicdata$panel == panel]
    blockstarts <- which(duplicated(ptimes))
    if(j == 1){
        b1 <- 1
    } else b1  <- ptimes[blockstarts[j - 1] + 1]
    if(j <= length(blockstarts)){
        b2 <- ptimes[blockstarts[j]]
    } else b2 <- max(ptimes)
    timeblock <- seq(from = b1, to = b2)
    return(timeblock)
}

#check that numerical solution of ODEs in aureliaode.stan is close to what we get in R
#Arguments:
#a0: potential competitor settlement rate (iterations)
#a1: potential competitor growth rate (iterations)
#a2: potential competitor death rate (iterations)
#deltab0: polyp settlement rate * polyp area (iterations)
#b1: polyp budding rate
#b2: polyp death rate
#rho: Stan estimate of composition (iterations x (panels stacked by time) x parts, in order Aurelia, bare space, potential competitors
#Value: prints out composition over time from ode solution in R and Stan
#Notes: rough version, currently no treatments or depth
checkstanode <- function(a0, a1, a2, deltab0, b1, b2, rho){
    parameters <- c(Pi1 = a0[1] / deltab0[1], Pi2 = a1[1] / deltab0[1], Pi3 = a2[1] / deltab0[1], Pi4 = b1[1] / deltab0[1], Pi5 = b2[1] / deltab0[1]) #takes just a single stan iteration
    times <- c(0:8) * deltab0[1]
    state <- c(xstar = 0, y1star = 0)
    aureliasol <- ode(y = state, times = times, func = aurelia2dODE, parms = parameters)
    print("From R ode:")
    print(cbind(c(1:8), aureliasol[-1, 3], 1 - aureliasol[-1, 2] - aureliasol[-1, 3], aureliasol[-1, 2]))
    print("From stan ode:")
    print(rho[1, 1:8, ])
}

#plot the solution to ODE model from Stan
#Arguments:
#rho: predicted composition from ODE (iterations x stacked panels x parts)
#times: vector of time points (iterations x stacked panels)
#dynamicdata: data frame created by datafordynamicmodel(): rows are iterations x stacked panels: includes panel variable
#panels: vector: which panels do we want to plot?
#Value: plots each component against time, metric centre and marginal 95% credible band, for requested panels
#Notes: currently very rough. 
plotstanode <- function(rho, times, dynamicdata, panels){
    par(mfrow = c(3, 1))
    par(mar = c(4, 4, 2, 2))
#    times <- times[1:14] #ROUGH: FIRST PANEL ONLY
#   rhosummary <- getrhosummary(rho[, 1:14, ])
    rhosummary <- getrhosummary(rho)
    npanels <- length(panels)
    panelindex <- vector(mode = "list", length = npanels)
    for(i in 1:npanels){
        panelindex[[i]] <- which(dynamicdata$panel == panels[i])
    }
    plot(times[panelindex[[1]]], rhosummary$metriccentre[panelindex[[1]], 1], ylim = c(0, 1), ylab = expression(italic(A.~aurita)), xlab = "time/weeks", type = "n")
    for(i in 1:npanels){
        polygon(c(times[panelindex[[i]]], rev(times[panelindex[[i]]])), c(rhosummary$simplexhdr[1, 1, panelindex[[i]]], rev(rhosummary$simplexhdr[1, 2, panelindex[[i]]])), col = adjustcolor("black", 0.4), border = NA)
    lines(times[panelindex[[i]]], rhosummary$metriccentre[panelindex[[i]], 1], col = "black")
    }

    plot(times[panelindex[[1]]], rhosummary$metriccentre[panelindex[[1]], 2], ylim = c(0, 1), ylab = "bare panel", xlab = "time/weeks", type = "n")
    for(i in 1:npanels){
        polygon(c(times[panelindex[[i]]], rev(times[panelindex[[i]]])), c(rhosummary$simplexhdr[2, 1, panelindex[[i]]], rev(rhosummary$simplexhdr[2, 2, panelindex[[i]]])), col = adjustcolor("black", 0.4), border = NA)
    lines(times[panelindex[[i]]], rhosummary$metriccentre[panelindex[[i]], 2], col = "black")
    }

    plot(times[panelindex[[1]]], rhosummary$metriccentre[panelindex[[1]], 3], ylim = c(0, 1), ylab = "potential competitors", xlab = "time/weeks", type = "n")
    for(i in 1:npanels){
        polygon(c(times[panelindex[[i]]], rev(times[panelindex[[i]]])), c(rhosummary$simplexhdr[3, 1, panelindex[[i]]], rev(rhosummary$simplexhdr[3, 2, panelindex[[i]]])), col = adjustcolor("black", 0.4), border = NA)
    lines(times[panelindex[[i]]], rhosummary$metriccentre[panelindex[[i]], 3], col = "black")
    }

    
#    plot(times, rep(0, length(times)), ylim = c(0, 1), ylab = "bare panel", xlab = "time/weeks", type = "n")
#    polygon(c(times, rev(times)), c(rhosummary$simplexhdr[2, 1, ], rev(rhosummary$simplexhdr[2, 2, ])), col = adjustcolor("black", 0.4), border = NA)
#    lines(times, rhosummary$metriccentre[, 2], col = "black")
#    plot(times, rep(0, length(times)), ylim = c(0, 1), ylab = "potential competitors", xlab = "time/weeks", type = "n")
#    polygon(c(times, rev(times)), c(rhosummary$simplexhdr[3, 1, ], rev(rhosummary$simplexhdr[3, 2, ])), col = adjustcolor("black", 0.4), border = NA)
#    lines(times, rhosummary$metriccentre[, 3], col = "black")
}

#get metric centre and marginal 95% credible bands for ODE prediction of composition
#Arguments: rho: predicted composition, iterations x stacked panels over time x parts
#Value: list containing
#metriccentre: metric centre (stacked panels over time x parts)
#simplexhdr: marginal 95% credible bands: parts x [lower, upper] x stacked panels over time
getrhosummary <- function(rho){
    rhoilr <- apply(rho, c(1, 2), "ilr")
    rhoilrmean <- t(apply(rhoilr, c(1, 3), "mean"))
    metriccentre <- ilrInv(rhoilrmean)

    myhdr <- function(y){
        hdr(y, prob = 95)$hdr
    }
    
    xhdr <- apply(rhoilr, c(1, 3), FUN = myhdr)
    simplexhdr <- apply(xhdr, c(1, 3), "ilrInv")
    return(list(metriccentre = metriccentre, simplexhdr = simplexhdr))
}

#some rough simulations of ODE model to choose sensible priors
#Arguments:
#niter: number of iterations in which to sample from prior (default 1e2)
#nweeks: number of weeks for which to solve ODEs (default 8)
#fname: name of pdf file to write to
#Value: pdf plots of A. aurita, bare panel, potential competitors in each week, for each iteration
chooseodepriors <- function(niter = 1e2, nweeks = 8, fname){
    pdf(file = fname, width = 7, height = 7)
    odesol <- array(dim = c(niter, nweeks + 1, 3))
    dimnames(odesol) <- list(NULL, NULL, c("time", "potentialcomp", "Aaurita"))
    growthrates <- array(dim = c(niter, 2))
    colnames(growthrates) <- c("Aauritagrowthperweek", "potentialcompgrowthperweek")
    for(i in 1:niter){
        a0 <- abs(rnorm(n = 1, mean = 0, sd = 0.05))
        a1 <- abs(rnorm(n = 1, mean = 0, sd = 1))
        a2 <- -abs(rnorm(n = 1, mean = 0, sd = 0.05))
        deltab0 <- abs(rnorm(n = 1, mean = 0, sd = 0.02))
        b1 <- abs(rnorm(n = 1, mean = 0, sd = 0.2))
        b2 <- -abs(rnorm(n = 1, mean = 0, sd = 0.05))
        growthrates[i, 1] <- b1 + b2
        growthrates[i, 2] <- a1 + a2
        parameters <- c(Pi1 = a0 / deltab0, Pi2 = a1 / deltab0, Pi3 = a2 / deltab0, Pi4 = b1 / deltab0, Pi5 = b2 / deltab0)
        times <- c(0:nweeks) * deltab0
        state <- c(xstar = 0, y1star = 0)
        odesol[i, , ] <- ode(y = state, times = times, func = aurelia2dODE, parms = parameters)
    }
    print(summary(growthrates))
    print("standard deviations of growth rates:")
    print(apply(growthrates, 2, "sd"))

    print("mean cover by time")
    print(apply(odesol[, , 2:3], c(2, 3), "mean"))

    print("sd cover by time")
    print(apply(odesol[, , 2:3], c(2, 3), "sd"))

    
    par(mfrow = c(3, 1))
    par(mar = c(4, 4, 2, 2))
    plot(0:nweeks, rep(0, nweeks + 1), type = "n", xlab = "time/weeks", ylab = expression(italic(A.~aurita)), ylim = c(0, 1))
    for(i in 1:niter){
        lines(0:nweeks, odesol[i, , 3], col = adjustcolor("black", 0.1))
    }
    axl <- par("usr")
    text(axl[1] + 0.05 * (axl[2] - axl[1]), axl[4] - 0.2 * (axl[4] - axl[3]), "a")
    plot(0:nweeks, rep(0, nweeks + 1), type = "n", xlab = "time/weeks", ylab = "bare panel", ylim = c(0, 1))
    for(i in 1:niter){
        lines(0:nweeks, 1 - odesol[i, , 2] - odesol[i, , 3], col = adjustcolor("black", 0.1))
    }
    axl <- par("usr")
    text(axl[1] + 0.05 * (axl[2] - axl[1]), axl[4] - 0.2 * (axl[4] - axl[3]), "b")
    plot(0:nweeks, rep(0, nweeks + 1), type = "n", xlab = "time/weeks", ylab = "potential competitors", ylim = c(0, 1))
    for(i in 1:niter){
        lines(0:nweeks, odesol[i, , 2], col = adjustcolor("black", 0.1))
    }
    axl <- par("usr")
    text(axl[1] + 0.05 * (axl[2] - axl[1]), axl[4] - 0.2 * (axl[4] - axl[3]), "c")
    dev.off()
}

#Build rho for a given treatment, depth and removal pattern
#Arguments:
#a0: potential competitor settlement rate
#a1: potential competitor growth rate
#a2: potential competitor death rate
#deltab0: polyp settlement rate * polyp area
#b1: polyp budding rate
#b2: polyp death rate
#kA: proportion A. aurita not removed in A treatment
#kO: proportion potential competitors not removed in O treatment
#m0: facilitation effect of polyps on potential comp settlement
#m2: reduction in potential comp mortality due to polyps
#psi: frequency-dependence exponent
#a1y1star: overgrowth rate of polyps by potential competitors
#treatmentapplied: vector of treatment applied at each time point: code 0 for nothing, 2 for A treatment, 3 for O treatment
#m1: growth facilitation effect of polyps on potential competitors
#timeblocks: array (ntimeblocks x max number of time blocks) of observation times in each block (between treatment applications). -1 for unused values
#ntimeblocks: number of time blocks for this panel
#ntimesinblock: vector (max number of time blocks): number of observation times in each time block
#controltruncate (default TRUE): shorten because no final post-treatment observation? Set to TRUE when calling from typicalrho(), but FALSE when calling using timeblocks ... ntimesinblock as supplied to Stan
#Value:
#array rho: rows are time points, columns are A. aurita, bare panel, potential competitors
buildrho <- function(a0, a1, a2, deltab0, b1, b2, kA, kO, m0, m2, psi, a1y1star,m1, treatmentapplied, timeblocks, ntimeblocks, ntimesinblock, controltruncate = TRUE){
#    parameters <- c(Pi1 = a0 / deltab0, Pi2 = a1 / deltab0, Pi3 = a2 / deltab0, Pi4 = b1 / deltab0, Pi5 = b2 / deltab0) #parameters scaled by deltab0
    parameters <- c(a0 = a0, a1 = a1, a2 = a2, m0 = m0, m2 = m2, deltab0 = deltab0, b1 = b1, b2 = b2, psi = psi, a1y1star = a1y1star, m1 = m1) #parameters scaled by sampling interval
    state <- c(xstar = 0, y1star = 0)
    t0 <- 0
    tindex <- 1
    nrows <- length(treatmentapplied)
    if(treatmentapplied[nrows] == 0 & controltruncate == TRUE) nrows <- nrows - 1 #control panel: no final post-treatment observation
    rho <- array(dim = c(nrows, 3))
    for(k in 1:ntimeblocks){
#        aureliasol <- ode(y = state, times = c(t0, timeblocks[k, 1:ntimesinblock[k]]) * deltab0, func = aurelia2dODEdim, parms = parameters) #time scaled by deltab0
        aureliasol <- ode(y = state, times = c(t0, timeblocks[k, 1:ntimesinblock[k]]), func = aurelia2dODEdim, parms = parameters) #time scaled by sampling interval
        rho[tindex:(tindex + ntimesinblock[k] - 1), ] <- c(aureliasol[-1, 3], 1 - aureliasol[-1, 2] - aureliasol[-1, 3], aureliasol[-1, 2])
        t0 <- timeblocks[k, ntimesinblock[k]]
        state <- aureliasol[ntimesinblock[k] + 1, 2:3]

        if(!(tindex + ntimesinblock[k] > length(treatmentapplied))){#indexing problem with last time block in control treatment
            if(treatmentapplied[tindex + ntimesinblock[k]] == 2){
                state <- Atreatment(state = state, kA = kA)
            } else if(treatmentapplied[tindex + ntimesinblock[k]] == 3){
                state <- Otreatment(state = state, kO = kO)
            }
            if(treatmentapplied[tindex + ntimesinblock[k]] != 0){
                rho[tindex + ntimesinblock[k], ] <- c(state[2], 1 - sum(state), state[1])
                tindex <- tindex + 1
            }
        }
        tindex <- tindex + ntimesinblock[k]
    }
    return(rho)
}

#remove proportion 1 - kA of A. aurita from vector state
Atreatment <- function(state, kA){
    state[2] <- kA * state[2]
    return(state)
}

#remove proportion 1 - kO of potential competitors from vector state
Otreatment <- function(state, kO){
    state[1] <- kO * state[1]
    return(state)
}


#build predicted composition for typical panel from each treatment combination
#Arguments:
#a0: potential competitor settlement rate
#a1: potential competitor growth rate
#a2: potential competitor death rate
#deltab0: polyp settlement rate * polyp area
#b1: polyp budding rate
#b2: polyp death rate
#kA: proportion A. aurita not removed in A treatment
#kO: proportion potential competitors not removed in O treatment
#m0: facilitation effect of polyps on potential comp settlement (default NA: no effect)
#m2: reduction in potential comp mortality due to polyps (default NA: no effect)
#niter: number of iterations to use
#psi: frequency-dependence exponent (default NA no effect)
#a1y1star: overgrowth rate (default NA: no effect)
#m1: growth facilitation effect (default NA: no effect)
#Value: list of arrays rhoC1, ..., rhoA3: each iterations x time points x 3: predicted composition for typical panel from each treatment and depth. Note that the number of time points differs between combinations. Typical treatment application pattern for each combination hard coded.
#and objects rhoC1summary, ..., rhoA3summary: lists containing metric centre and 95% credible bands for each combination
typicalrho <- function(a0, a1, a2, deltab0, b1, b2, kA, kO, m0 = NA, m2 = NA, psi = NA, a1y1star = NA, m1 = NA, niter){
    rhoC1 <- array(dim = c(niter, 8, 3)) #control at 1 m
    depthcode <- 1
    treatmentapplied <- rep(0, 9)
    timeblocks <- array(1:8, dim = c(1, 8))
    ntimesinblock <- 8
    ntimeblocks <- 1
    for(i in 1:niter){
        if (is.na(m0[1])){
            m0i <- 0
        } else {
            m0i <- m0[i, depthcode]
        }
        if (is.na(m2[1])){
            m2i <- 0
        } else {
            m2i <- m2[i, depthcode]
        }
        if (is.na(a1y1star[1])){
            a1y1stari <- 0
        } else {
            a1y1stari <- a1y1star[i, depthcode]
        }
        if (is.na(psi[1])){
            psii <- 0
        } else {
            psii <- psi[i, depthcode]
        }
        if (is.na(m1[1])){
            m1i <- 0
        } else {
            m1i <- m1[i, depthcode]
        }

        rhoC1[i, , ] <-  buildrho(a0 = a0[i, depthcode], a1 = a1[i, depthcode], a2 = a2[i, depthcode], deltab0 = deltab0[i, depthcode], b1 = b1[i, depthcode], b2 = b2[i, depthcode], kA = kA[i], kO = kO[i], m0 = m0i, m2 = m2i, a1y1star = a1y1stari, psi = psii, m1 = m1i, treatmentapplied = treatmentapplied, timeblocks = timeblocks, ntimeblocks = ntimeblocks, ntimesinblock = ntimesinblock)
    }

    rhoC3 <- array(dim = c(niter, 8, 3)) #control at 3 m
    depthcode <- 2
    treatmentapplied <- rep(0, 9)
    timeblocks <- array(1:8, dim = c(1, 8))
    ntimesinblock <- 8
    ntimeblocks <- 1
    for(i in 1:niter){
        if (is.na(m0[1])){
            m0i <- 0
        } else {
            m0i <- m0[i, depthcode]
        }
        if (is.na(m2[1])){
            m2i <- 0
        } else {
            m2i <- m2[i, depthcode]
        }
        if (is.na(a1y1star[1])){
            a1y1stari <- 0
        } else {
            a1y1stari <- a1y1star[i, depthcode]
        }
        if (is.na(psi[1])){
            psii <- 0
        } else {
            psii <- psi[i, depthcode]
        }
        if (is.na(m1[1])){
            m1i <- 0
        } else {
            m1i <- m1[i, depthcode]
        }
        rhoC3[i, , ] <-  buildrho(a0 = a0[i, depthcode], a1 = a1[i, depthcode], a2 = a2[i, depthcode], deltab0 = deltab0[i, depthcode], b1 = b1[i, depthcode], b2 = b2[i, depthcode], kA = kA[i], kO = kO[i], m0 = m0i, m2 = m2i, a1y1star = a1y1stari, psi = psii, m1 = m1i, treatmentapplied = treatmentapplied, timeblocks = timeblocks, ntimeblocks = ntimeblocks, ntimesinblock = ntimesinblock)
    }

    rhoO1 <- array(dim = c(niter, 16, 3)) #O at 1 m
    depthcode <- 1
    treatmentapplied <- rep(c(0, 3), 8)
    timeblocks <- array(-1, dim = c(8, 8))
    timeblocks[, 1] <- 1:8
    ntimesinblock <- rep(1, 8)
    ntimeblocks <- 8
    for(i in 1:niter){
        if (is.na(m0[1])){
            m0i <- 0
        } else {
            m0i <- m0[i, depthcode]
        }
        if (is.na(m2[1])){
            m2i <- 0
        } else {
            m2i <- m2[i, depthcode]
        }
        if (is.na(a1y1star[1])){
            a1y1stari <- 0
        } else {
            a1y1stari <- a1y1star[i, depthcode]
        }
        if (is.na(psi[1])){
            psii <- 0
        } else {
            psii <- psi[i, depthcode]
        }
        if (is.na(m1[1])){
            m1ii <- 0
        } else {
            m1i <- m1[i, depthcode]
        }
        rhoO1[i, , ] <-  buildrho(a0 = a0[i, depthcode], a1 = a1[i, depthcode], a2 = a2[i, depthcode], deltab0 = deltab0[i, depthcode], b1 = b1[i, depthcode], b2 = b2[i, depthcode], kA = kA[i], kO = kO[i], m0 = m0i, m2 = m2i, a1y1star = a1y1stari, psi = psii, m1 = m1i, treatmentapplied = treatmentapplied, timeblocks = timeblocks, ntimeblocks = ntimeblocks, ntimesinblock = ntimesinblock)
    }

    rhoO3 <- array(dim = c(niter, 16, 3)) #O at 3 m
    depthcode <- 2
    treatmentapplied <- rep(c(0, 3), 8)
    timeblocks <- array(-1, dim = c(8, 8))
    timeblocks[, 1] <- 1:8
    ntimesinblock <- rep(1, 8)
    ntimeblocks <- 8
    for(i in 1:niter){
        if (is.na(m0[1])){
            m0i <- 0
        } else {
            m0i <- m0[i, depthcode]
        }
        if (is.na(m2[1])){
            m2i <- 0
        } else {
            m2i <- m2[i, depthcode]
        }
        if (is.na(a1y1star[1])){
            a1y1stari <- 0
        } else {
            a1y1stari <- a1y1star[i, depthcode]
        }
        if (is.na(psi[1])){
            psii <- 0
        } else {
            psii <- psi[i, depthcode]
        }
        if (is.na(m1[1])){
            m1ii <- 0
        } else {
            m1i <- m1[i, depthcode]
        }
        rhoO3[i, , ] <-  buildrho(a0 = a0[i, depthcode], a1 = a1[i, depthcode], a2 = a2[i, depthcode], deltab0 = deltab0[i, depthcode], b1 = b1[i, depthcode], b2 = b2[i, depthcode], kA = kA[i], kO = kO[i], m0 = m0i, m2 = m2i, a1y1star = a1y1stari, psi = psii, m1 = m1i, treatmentapplied = treatmentapplied, timeblocks = timeblocks, ntimeblocks = ntimeblocks, ntimesinblock = ntimesinblock)
    }

    rhoA1 <- array(dim = c(niter, 15, 3)) #A at 1 m
    depthcode <- 1
    treatmentapplied <- c(0, rep(c(0, 2), 7)) #A treatment typically not applied in week 1
    timeblocks <- array(-1, dim = c(7, 8))
    timeblocks[1, 1:2] <- 1:2
    timeblocks[2:7, 1] <- 3:8
    ntimesinblock <- c(2, rep(1, 6))
    ntimeblocks <- 7
    for(i in 1:niter){
        if (is.na(m0)[1]){
            m0i <- 0
        } else {
            m0i <- m0[i, depthcode]
        }
        if (is.na(m2[1])){
            m2i <- 0
        } else {
            m2i <- m2[i, depthcode]
        }
        if (is.na(a1y1star[1])){
            a1y1stari <- 0
        } else {
            a1y1stari <- a1y1star[i, depthcode]
        }
        if (is.na(psi[1])){
            psii <- 0
        } else {
            psii <- psi[i, depthcode]
        }
        if (is.na(m1[1])){
            m1ii <- 0
        } else {
            m1i <- m1[i, depthcode]
        }
        rhoA1[i, , ] <-  buildrho(a0 = a0[i, depthcode], a1 = a1[i, depthcode], a2 = a2[i, depthcode], deltab0 = deltab0[i, depthcode], b1 = b1[i, depthcode], b2 = b2[i, depthcode], kA = kA[i], kO = kO[i], m0 = m0i, m2 = m2i, a1y1star = a1y1stari, psi = psii, m1 = m1i, treatmentapplied = treatmentapplied, timeblocks = timeblocks, ntimeblocks = ntimeblocks, ntimesinblock = ntimesinblock)
    }

   rhoA3 <- array(dim = c(niter, 15, 3)) #A at 3 m
    depthcode <- 2
    treatmentapplied <- c(0, rep(c(0, 2), 7)) #A treatment typically not applied in week 1
    timeblocks <- array(-1, dim = c(7, 8))
    timeblocks[1, 1:2] <- 1:2
    timeblocks[2:7, 1] <- 3:8
    ntimesinblock <- c(2, rep(1, 6))
    ntimeblocks <- 7
    for(i in 1:niter){
        if (is.na(m0[1])){
            m0i <- 0
        } else {
            m0i <- m0[i, depthcode]
        }
        if (is.na(m2[1])){
            m2i <- 0
        } else {
            m2i <- m2[i, depthcode]
        }
        if (is.na(a1y1star[1])){
            a1y1stari <- 0
        } else {
            a1y1stari <- a1y1star[i, depthcode]
        }
        if (is.na(psi[1])){
            psii <- 0
        } else {
            psii <- psi[i, depthcode]
        }
        if (is.na(m1[1])){
            m1ii <- 0
        } else {
            m1i <- m1[i, depthcode]
        }
        rhoA3[i, , ] <-  buildrho(a0 = a0[i, depthcode], a1 = a1[i, depthcode], a2 = a2[i, depthcode], deltab0 = deltab0[i, depthcode], b1 = b1[i, depthcode], b2 = b2[i, depthcode], kA = kA[i], kO = kO[i], m0 = m0i, m2 = m2i, a1y1star = a1y1stari, psi = psii, m1 = m1i, treatmentapplied = treatmentapplied, timeblocks = timeblocks, ntimeblocks = ntimeblocks, ntimesinblock = ntimesinblock)
    }
    rhoC1summary <- getrhosummary(rhoC1)
    rhoC3summary <- getrhosummary(rhoC3)
    rhoO1summary <- getrhosummary(rhoO1)
    rhoO3summary <- getrhosummary(rhoO3)
    rhoA1summary <- getrhosummary(rhoA1)
    rhoA3summary <- getrhosummary(rhoA3)
    
    return(list(rhoC1 = rhoC1, rhoC3 = rhoC3, rhoO1 = rhoO1, rhoO3 = rhoO3, rhoA1 = rhoA1, rhoA3 = rhoA3, rhoC1summary = rhoC1summary, rhoC3summary = rhoC3summary, rhoO1summary = rhoO1summary, rhoO3summary = rhoO3summary, rhoA1summary = rhoA1summary, rhoA3summary = rhoA3summary))
}

#plot posterior estimates of parameters from ODE model
#Arguments:
#a0: potential competitor settlement rate
#a1: potential competitor growth rate
#a2: potential competitor death rate
#deltab0: polyp settlement rate * polyp area
#b1: polyp budding rate
#b2: polyp death rate
#kA: proportion A. aurita not removed in A treatment
#kO: proportion potential competitors not removed in O treatment
#fname: name of pdf for plot
#Value: plots of posterior density for each parameter. All except kA and kO: two densities (for 1 m, solid filled, and 3 m, dashed filled). Prior densities (all except kA and kO half normal, kA and kO beta) solid unfilled
plotodeparameters <- function(a0, a1, a2, deltab0, b1, b2, kA, kO, fname){
    pdf(file = fname, width = 7, height = 7)
    par(mfrow = c(3, 3))
    par(mar = c(4, 4, 2, 2))
    da01 <- density(a0[, 1])
    da02 <- density(a0[, 2])
    da11 <- density(a1[, 1])
    da12 <- density(a1[, 2])
    da21 <- density(a2[, 1])
    da22 <- density(a2[, 2])
    ddeltab01 <- density(deltab0[, 1])
    ddeltab02 <- density(deltab0[, 2])
    db11 <- density(b1[, 1])
    db12 <- density(b1[, 2])
    db21 <- density(b2[, 1])
    db22 <- density(b2[, 2])
    dkA <- density(kA)
    dkO <- density(kO)
    xrange <- NULL
    yrange <- NULL
    plotoneodehalfnparameter(d1 = da01, d2 = da02, xrange = xrange, yrange = yrange, priorsd = 0.05, label = "a", xlab = expression(paste(italic(a)[0]/weeks^-1)))
    plotoneodehalfnparameter(d1 = da11, d2 = da12, xrange = xrange, yrange = yrange, priorsd = 1, label = "b", xlab = expression(paste(italic(a)[1]/weeks^-1)))
    plotoneodehalfnparameter(d1 = da21, d2 = da22, xrange = xrange, yrange = yrange, priorsd = 0.05, label = "c", xlab = expression(paste(italic(a)[2]/weeks^-1)))
    plotoneodehalfnparameter(d1 = ddeltab01, d2 = ddeltab02, xrange = xrange, yrange = yrange, priorsd = 0.02, label = "d", xlab = expression(paste(delta * italic(b)[0]/weeks^-1)))
    plotoneodehalfnparameter(d1 = db11, d2 = db12, xrange = xrange, yrange = yrange, priorsd = 0.2, label = "e", xlab = expression(paste(italic(b)[1]/weeks^-1)))
    plotoneodehalfnparameter(d1 = db21, d2 = db22, xrange = xrange, yrange = yrange, priorsd = 0.05, label = "f", xlab = expression(paste(italic(b)[2]/weeks^-1)))
    plotoneodebetaparameter(d1 = dkA, yrange = NULL, prior1 = 2, prior2 = 2, label = "g", xlab = expression(paste(italic(k[A]))))
    plotoneodebetaparameter(d1 = dkO, yrange = NULL, prior1 = 2, prior2 = 2, label = "h", xlab = expression(paste(italic(k[O]))))
    dev.off()
}

#plot a single parameter from ODE model (from two different depths) with half-normal prior
#Arguments: d1, d2: density for the parameter at 1 m and 3 m
#xrange, yrange: ranges for x and y axes. If null (default) calculate from the densities d1 and d2
#priorsd: standard deviation for half-normal prior
#label:text for panel label
#xlab: label for x axis
#Value: plots of densities at 1 m (dashed line), 3 m (solid line) and prior (unfilled solid: assumed half-normal, with mode at zero)
plotoneodehalfnparameter <- function(d1, d2, xrange = NULL, yrange = NULL, priorsd, label, xlab){
    if(is.null(xrange)) xrange <- range(c(0, d1$x, d2$x))
    if(is.null(yrange)) yrange <- range(c(d1$y, d2$y))
    plot(d1, xlim = xrange, ylim = yrange, type = "n", main = "", xlab = xlab)
    polygon(d1, col = adjustcolor("black", 0.4), lty = "dashed")
    polygon(d2, col = adjustcolor("black", 0.4), lty = "solid")
    pxr <- xrange
    if(pxr[1] < 0) pxr[2] <- 0
    if(pxr[2] > 0) pxr[1]  <- 0
    priorx <- seq(from = pxr[1], to = pxr[2], length.out = 100)
    dprior <- 2 * dnorm(x = priorx, sd = priorsd) #multiply by 2 because half-normal
    lines(priorx, dprior)
    text(xrange[1] + 0.45 * (xrange[2] - xrange[1]), yrange[2] - 0.1 * (yrange[2] - yrange[1]), label)
}

#plot a single parameter from ODE model with beta prior
#Arguments: d1: density for the parameter
#yrange: range for y axis. If null (default) calculate from the density d1
#prior1, prior2: parameters for prior beta distribution
#label:text for panel label
#xlab: text for x axis label
#Value: plot of posterior density (filled solid) and prior (unfilled solid)
plotoneodebetaparameter <- function(d1, yrange = NULL, prior1, prior2, label, xlab){
    if(is.null(yrange)) yrange <- range(d1$y)
    xrange <- c(0, 1)
    kx <- seq(from = 0, to = 1, length.out = 100)
    kprior <- dbeta(x = kx, shape1 = prior1, shape2 = prior2)
    plot(d1, xlim = xrange, ylim = yrange, type = "n", main = "", xlab = xlab)
    lines(kx, kprior)
    polygon(d1, col = adjustcolor("black", 0.4))
    text(xrange[1] + 0.45 * (xrange[2] - xrange[1]), yrange[2] - 0.1 * (yrange[2] - yrange[1]), label)
}

#calculate logit test statistics from ODE output
#Arguments: typicalrhoset: object returned by typicalrho()
#Value: plots of difference in logit(potential competitors) between A and C treatments, and in logit(A. aurita) between O and C treatments, at 1 m and 3 m.
#Example:
#load(file = paste(odeoutputname, "_typicalrho.RData", sep = ""))
#ODElogitstatistics(typicalrhoset = typicalrhoset)
ODElogitstatistics <- function(typicalrhoset){
    logitA1 <- log(typicalrhoset$rhoA1[, 14, 3] / (1 - typicalrhoset$rhoA1[, 14, 3])) #logit potential competitors in A treatment. Second index is hard coded to be last week pre-treatment in each typical treatment pattern
    logitA3 <- log(typicalrhoset$rhoA3[, 14, 3] / (1 - typicalrhoset$rhoA3[, 14, 3]))
    logitC1a <- log(typicalrhoset$rhoC1[, 8, 3] / (1 - typicalrhoset$rhoC1[, 8, 3])) #logit potential competitors in C treatment
    logitC3a <- log(typicalrhoset$rhoC3[, 8, 3] / (1 - typicalrhoset$rhoC3[, 8, 3]))
    logitC1o <- log(typicalrhoset$rhoC1[, 8, 1] / (1 - typicalrhoset$rhoC1[, 8, 1])) #logit A. aurita in C treatment
    logitC3o <- log(typicalrhoset$rhoC3[, 8, 1] / (1 - typicalrhoset$rhoC3[, 8, 1]))
    logitO1 <- log(typicalrhoset$rhoO1[, 15, 1] / (1 - typicalrhoset$rhoO1[, 15, 1])) #logit A. aurita in O treatment
    logitO3 <- log(typicalrhoset$rhoO3[, 15, 1] / (1 - typicalrhoset$rhoO3[, 15, 1]))
    logitC1o <- log(typicalrhoset$rhoC1[, 8, 1] / (1 - typicalrhoset$rhoC1[, 8, 1])) #logit A. aurita in C treatment
    logitC3o <- log(typicalrhoset$rhoC3[, 8, 1] / (1 - typicalrhoset$rhoC3[, 8, 1]))
    Aresponse1 <- logitA1 - logitC1a
    Aresponse3 <- logitA3 - logitC3a
    Oresponse1 <- logitO1 - logitC1o
    Oresponse3 <- logitO3 - logitC3o
    mycolors <- brewer.pal(3, 'Dark2')
    pls <- c("a: 1 m", "b: 3 m")
    legends  <- list(c(expression(Potential~competitors*","~italic(A)~vs~italic(C)), expression(italic(A.~aurita)*","~italic(O)~vs~italic(C))), NULL)
    xlim <- c(-3, 3) #chose these values by hand to put all density plots on same scale
    ylim <- c(0, 100)
    par(mfrow = c(2, 1))
    plotoneresponse(logitchange1 = Aresponse1, logitchange2 = Oresponse1, cols = mycolors[c(2, 3)], panellabel = pls[1], xlim = xlim, ylim = ylim, legend = legends[[1]])
    plotoneresponse(logitchange1 = Aresponse3, logitchange2 = Oresponse3, cols = mycolors[c(2, 3)], panellabel = pls[2], xlim = xlim, ylim = ylim, legend = legends[[2]])
}

#test code for Euler-Maruyama
EMtest <- function(){
    parameters <- c(Pi1 = a0[1, 1] / deltab0[1, 1], Pi2 = a1[1, 1] / deltab0[1, 1], Pi3 = a2[1, 1] / deltab0[1, 1], Pi4 = b1[1, 1] / deltab0[1, 1], Pi5 = b2[1, 1] / deltab0[1, 1])
    times <- c(0:8) * deltab0[1, 1]
    state <- c(xstar = 0, y1star = 0)
    aureliasol <- ode(y = state, times = times, func = aurelia2dODE, parms = parameters)
#    print("Using ode(), 1 m")
    xode1 <- cbind(aureliasol[, 3], 1 - aureliasol[, 2] - aureliasol[, 3], aureliasol[, 2])
#    print(xode1)

    parameters <- c(Pi1 = a0[1, 2] / deltab0[1, 2], Pi2 = a1[1, 2] / deltab0[1, 2], Pi3 = a2[1, 2] / deltab0[1, 2], Pi4 = b1[1, 2] / deltab0[1, 2], Pi5 = b2[1, 2] / deltab0[1, 2])
    times <- c(0:8) * deltab0[1, 2]
    state <- c(xstar = 0, y1star = 0)
    aureliasol <- ode(y = state, times = times, func = aurelia2dODE, parms = parameters)
 #   print("Using ode(), 3 m")
    xode3 <- cbind(aureliasol[, 3], 1 - aureliasol[, 2] - aureliasol[, 3], aureliasol[, 2])
  #  print(xode3)

    Vdyn <- matrix(c(-1 / sqrt(2), 0, 1 / sqrt(2), 1 / sqrt(6), - 2 / sqrt(6), 1 / sqrt(6)), nrow = 3, ncol = 2)
    R <- 0.2 * diag(2) #arbitrary choice for test
    #epsilon <- 1e-2
    #init <- c(epsilon / 2, 1 - epsilon, epsilon / 2)
    t0 <- 0
    t1 <- 1
    nsteps  <- 1
    xout1 <- array(dim = c(9, 3))
    xout1[1, ] <- NA #on boundary of simplex
    xout1[2, ] <- xode1[2, ] #initialize with week 1 of ode() solution
    for(i in 2:8){
        xout1[i + 1, ] <- EMapproximation(a0 = a0[1, 1], a1 = a1[1, 1], a2 = a2[1, 1], deltab0 = deltab0[1, 1], b1 = b1[1, 1], b2 = b2[1, 1], init = acomp(xout1[i, ]), t0 = t0, t1 = t1, nsteps = nsteps, V = Vdyn, R = R)
    }
#    print("Using EM, 1 m:")
#    print(xout1)

    xout3 <- array(dim = c(9, 3))
    xout3[1, ] <- NA #on boundary of simplex
    xout3[2, ] <- xode3[2, ] #initialize with week 1 of ode() solution
    for(i in 2:8){
        xout3[i + 1, ] <- EMapproximation(a0 = a0[1, 2], a1 = a1[1, 2], a2 = a2[1, 2], deltab0 = deltab0[1, 2], b1 = b1[1, 2], b2 = b2[1, 2], init = acomp(xout3[i, ]), t0 = t0, t1 = t1, nsteps = nsteps, V = Vdyn, R = R)
    }
 #   print("Using EM, 3 m:")
 #   print(xout3)

    time <- c(0:8)
    par(mfrow = c(3, 2))
    plot(time, xode1[, 1], type = "l", xlab = "time/weeks", ylab = expression(italic(A.~aurita)), ylim = c(0, 1))
    lines(time[-1], xout1[-1, 1], col = "red")
    legend("topright", bty = "n", col = c("red", "black"), legend = c("Euler", "ode()"), pch = c(NA, NA), lty = c("solid", "solid"))
    axl <- par("usr")
    text(axl[1] + 0.1 * (axl[2] - axl[1]), axl[4] - 0.1 * (axl[4] - axl[3]), "a: 1 m")
    plot(time, xode3[, 1], type = "l", xlab = "time/weeks", ylab = expression(italic(A.~aurita)), ylim = c(0, 1))
    lines(time[-1], xout3[-1, 1], col = "red")
    axl <- par("usr")
    text(axl[1] + 0.1 * (axl[2] - axl[1]), axl[4] - 0.1 * (axl[4] - axl[3]), "b: 3 m")

    plot(time, xode1[, 2], type = "l", xlab = "time/weeks", ylab = "bare panel", ylim = c(0, 1))
    lines(time[-1], xout1[-1, 2], col = "red")
    axl <- par("usr")
    text(axl[1] + 0.1 * (axl[2] - axl[1]), axl[4] - 0.1 * (axl[4] - axl[3]), "c: 1 m")

    plot(time, xode3[, 2], type = "l", xlab = "time/weeks", ylab = "bare panel", ylim = c(0, 1))
    lines(time[-1], xout3[-1, 2], col = "red")
    axl <- par("usr")
    text(axl[1] + 0.1 * (axl[2] - axl[1]), axl[4] - 0.1 * (axl[4] - axl[3]), "d: 3 m")

    plot(time, xode1[, 3], type = "l", xlab = "time/weeks", ylab = "potential competitors", ylim = c(0, 1))
    lines(time[-1], xout1[-1, 3], col = "red")
    axl <- par("usr")
    text(axl[1] + 0.1 * (axl[2] - axl[1]), axl[4] - 0.1 * (axl[4] - axl[3]), "e: 1 m")

    plot(time, xode3[, 3], type = "l", xlab = "time/weeks", ylab = "potential competitors", ylim = c(0, 1))
    lines(time[-1], xout3[-1, 3], col = "red")
    axl <- par("usr")
    text(axl[1] + 0.1 * (axl[2] - axl[1]), axl[4] - 0.1 * (axl[4] - axl[3]), "f: 3 m")

}

#Euler-Maruyama approximation for SDE
#Arguments:
#a0: potential competitor settlement rate
#a1: potential competitor growth rate
#a2: potential competitor death rate
#deltab0: polyp settlement rate * polyp area
#b1: polyp budding rate
#b2: polyp death rate
#init: composition at start of time interval (A. aurita, bare panel, potential competitors)
#t0: initial time
#t1: final time
#nsteps: number of time steps
#V: ilr basis matrix
#R: Cholesky factor of noise covariance matrix
#Value:
#composition at end of time interval
EMapproximation <- function(a0, a1, a2, deltab0, b1, b2, init, t0, t1, nsteps, V, R){
    xk <- acomp(init)
    deltat <- (t1 - t0) / nsteps
    for(k in 1:nsteps){
        xk <- EMstep(a0 = a0, a1 = a1, a2 = a2, deltab0 = deltab0, b1 = b1, b2 = b2, x = xk, V = V, deltat = deltat, R = R)
    }
    return(xk)
}

#Euler-Maruyama step for SDE approximation
#Arguments:
#a0: potential competitor settlement rate
#a1: potential competitor growth rate
#a2: potential competitor death rate
#deltab0: polyp settlement rate * polyp area
#b1: polyp budding rate
#b2: polyp death rate
#x: composition at start of time step (A. aurita, bare panel, potential competitors)
#V: ilr basis matrix
#deltat: length of time step (in weeks)
#R: Cholesky factor of noise covariance matrix (lower triangular, positive diagonal elements)
#Value:
#xnew: composition at end of step
EMstep <- function(a0, a1, a2, deltab0, b1, b2, x, V, deltat, R){
    propchange <- c(deltab0 * x[2] / x[1] + b1 * x[2] + b2, -a0 - a1 * x[3] - a2 * x[3] / x[2] - deltab0 - b1 * x[1] - b2 * x[1] / x[2], a0 * x[2] / x[3] + a1 * x[2] + a2)
    print("EMstep propchange")
    print(propchange)

    dx <- a0 * x[2] +  a1 * x[3] * x[2] + a2 * x[3]
    dy1star <- deltab0 * x[2] + b1 * x[1] * x[2] + b2 * x[1]
    fx  <- c(dy1star, - dy1star - dx, dx)
    h <- 1e-6
    xth <- unclass(x) + h * fx
    zt <- ilr(x, V = V)
    zth <- ilr(xth, V = V)
    print("Numerical:")
    print((zth - zt) / h)
    mu <- propchange %*% V
    print("Calculated:")
    print(mu)
    noise <- rmvnorm(n = 1, mean = c(0, 0), sigma = deltat * diag(2)) %*% t(R)
    musimplex <- ilrInv(mu, V = V)
    noisesimplex <- ilrInv(noise, V = V)
    xnew <- x + deltat * musimplex + noisesimplex
    return(xnew)
}

#Euler-Maruyama step for SDE approximation that works even when initial condition is on boundary of simplex
#Arguments:
#a0: potential competitor settlement rate
#a1: potential competitor growth rate
#a2: potential competitor death rate
#deltab0: polyp settlement rate * polyp area
#b1: polyp budding rate
#b2: polyp death rate
#x: composition at start of time step (A. aurita, bare panel, potential competitors): can be of class acomp or just a vector
#V: ilr basis matrix
#deltat: length of time step (in weeks)
#R: Cholesky factor of noise covariance matrix (lower triangular, positive diagonal elements)
#Value:
#xnew: composition at end of step
EMstepboundary <- function(a0, a1, a2, deltab0, b1, b2, x, V, deltat, R){
    dx <- a0 * x[2] +  a1 * x[3] * x[2] + a2 * x[3]
    dy1star <- deltab0 * x[2] + b1 * x[1] * x[2] + b2 * x[1]
    fx  <- c(dy1star, - dy1star - dx, dx)
    print("EMstepboundary propchange:")
    print(c(dy1star / x[1], (- dy1star - dx) / x[2], dx / x[3]))
    xdet <- acomp(unclass(x) + fx * deltat)
    noise <- rmvnorm(n = 1, mean = c(0, 0), sigma = deltat * diag(2)) %*% t(R)
    noisesimplex <- ilrInv(noise, V = V)
    xnew <- xdet + noisesimplex
    return(xnew)
}

#debug difference between EMstep() and EMstep2()
EMstepdebug  <- function(){
    Vdyn <- matrix(c(-1 / sqrt(2), 0, 1 / sqrt(2), 1 / sqrt(6), - 2 / sqrt(6), 1 / sqrt(6)), nrow = 3, ncol = 2)
    R <- 0 * diag(2) #arbitrary choice for test
    a0 <- 0.1
    a1 <- 2
    a2  <- -0.5
    deltab0  <- 0.1
    b1 <- 0.8
    b2 <- -0.2
    deltat <- 1
    x <- acomp(c(0.7, 0.2, 0.1))
    test <- unclass(EMstep(a0 = a0, a1 = a1, a2 = a2, deltab0 = deltab0, b1 = b1, b2 = b2, x = x, V = Vdyn, deltat = deltat, R = R))
    test2 <- unclass(EMstepboundary(a0 = a0, a1 = a1, a2 = a2, deltab0 = deltab0, b1 = b1, b2 = b2, x = x, V = Vdyn, deltat = deltat, R = R))
    print("Difference EMstep vs EMstepboundary:")
    print(test - test2)
    print("EMstep:")
    print(test)
    print("EMstepboundary:")
    print(test2)

    parameters <- c(a0 = a0, a1 = a1, a2 = a2, m0 = 0, m2 = 0, deltab0 = deltab0, b1 = b1, b2 = b2, psi = 0, a1y1star = 0, m1 = 0) #parameters scaled by sampling interval
    state <- c(xstar = x[3], y1star = x[1])
    aureliasol <- ode(y = state, times = c(0, deltat), func = aurelia2dODEdim, parms = parameters) #time scaled by sampling interval
    test3 <- c(aureliasol[2, 3], 1 - aureliasol[2, 3] - aureliasol[2, 2], aureliasol[2, 2])
    print("ode:")
    print(test3)
                      


}

#Test code: how well does EM work with 1 step compared to larger numbers?
EMsteptest <- function(){
    parameters <- c(Pi1 = a0[1, 1] / deltab0[1, 1], Pi2 = a1[1, 1] / deltab0[1, 1], Pi3 = a2[1, 1] / deltab0[1, 1], Pi4 = b1[1, 1] / deltab0[1, 1], Pi5 = b2[1, 1] / deltab0[1, 1])
    times <- c(0:8) * deltab0[1, 1]
    state <- c(xstar = 0, y1star = 0)
    aureliasol <- ode(y = state, times = times, func = aurelia2dODE, parms = parameters)
    xode1 <- cbind(aureliasol[, 3], 1 - aureliasol[, 2] - aureliasol[, 3], aureliasol[, 2])

    Vdyn <- matrix(c(-1 / sqrt(2), 0, 1 / sqrt(2), 1 / sqrt(6), - 2 / sqrt(6), 1 / sqrt(6)), nrow = 3, ncol = 2)
    R <- 0.2 * diag(2) #arbitrary choice for test
    init <- xode1[2, ] #initialize with week 1 of ode() solution
    t0 <- 0
    t1 <- 1
    niter <- 100
    ntimes <- 8
    xout1 <- array(dim = c(ntimes, niter, 3))
    xout10 <- xout1
    xout100 <- xout1
    for(i in 1:niter){
        xout1[1, i, ] <- init
        xout10[1, i, ] <- init
        xout100[1, i, ] <- init
        for(j in 2:ntimes - 1){
            xout1[j + 1, i, ] <- EMapproximation(a0 = a0[1, 1], a1 = a1[1, 1], a2 = a2[1, 1], deltab0 = deltab0[1, 1], b1 = b1[1, 1], b2 = b2[1, 1], init = acomp(xout1[j, i, ]), t0 = t0, t1 = t1, nsteps = 1, V = Vdyn, R = R)
            xout10[j + 1, i, ] <- EMapproximation(a0 = a0[1, 1], a1 = a1[1, 1], a2 = a2[1, 1], deltab0 = deltab0[1, 1], b1 = b1[1, 1], b2 = b2[1, 1], init = acomp(xout10[j, i, ]), t0 = t0, t1 = t1, nsteps = 10, V = Vdyn, R = R)
            xout100[j + 1, i, ] <- EMapproximation(a0 = a0[1, 1], a1 = a1[1, 1], a2 = a2[1, 1], deltab0 = deltab0[1, 1], b1 = b1[1, 1], b2 = b2[1, 1], init = acomp(xout100[j, i, ]), t0 = t0, t1 = t1, nsteps = 100, V = Vdyn, R = R)
        }
    }
    par(mfrow = c(ntimes - 1, 3))
    par(mar = c(3, 3, 1, 2))
    mycolors <- brewer.pal(3, "Dark2")
    mylabs <- c("a: A. aurita, week 2", "b: bare panel, week 2", "c: potential comp., week 2", "d: A. aurita, week 3", "e: bare panel, week 3", "f: potential comp., week 3", "g: A. aurita, week 4", "h: bare panel, week 4", "i: potential comp., week 4", "j: A. aurita, week 5", "k: bare panel, week 5", "l: potential comp., week 5", "m: A. aurita, week 6", "n: bare panel, week 6", "o: potential comp., week 6", "p: A. aurita, week 7", "q: bare panel, week 7", "r: potential comp., week 7", "s: A. aurita, week 8", "t: bare panel, week 8", "u: potential comp., week 8")
    count <- 1
    for(j in 2:ntimes){
        d1 <- density(xout1[j, , 1])
        d10 <- density(xout10[j, , 1])
        d100 <- density(xout100[j, , 1])
        plot(d1, main = "", xlim = c(0, 1), ylim = c(0, max(c(d1$y, d10$y, d100$y))), col = mycolors[1], xlab = "", ylab = "")
        axl <- par("usr")
        text(axl[1] + 0.1 * (axl[2] - axl[1]), axl[4] - 0.1 * (axl[4] - axl[3]),mylabs[count], pos = 4, cex = 0.5)
        if(j == 2) legend("topright", bty = "n", lty = "solid", pch = NA, col = mycolors, legend = c("1 Euler step", "10 Euler steps", "100 Euler steps"), cex = 0.5)
        lines(d10, col = mycolors[2])
        lines(d100, col = mycolors[3])
        d1 <- density(xout1[j, , 2])
        d10 <- density(xout10[j, , 2])
        d100 <- density(xout100[j, , 2])
        plot(d1, main = "", xlim = c(0, 1), ylim = c(0, max(c(d1$y, d10$y, d100$y))), col = mycolors[1], xlab = "", ylab = "")
        lines(d10, col = mycolors[2])
        lines(d100, col = mycolors[3])
        axl <- par("usr")
        text(axl[1] + 0.1 * (axl[2] - axl[1]), axl[4] - 0.1 * (axl[4] - axl[3]),mylabs[count + 1], pos = 4, cex = 0.5)

        d1 <- density(xout1[j, , 3])
        d10 <- density(xout10[j, , 3])
        d100 <- density(xout100[j, , 3])
        plot(d1, main = "", xlim = c(0, 1), ylim = c(0, max(c(d1$y, d10$y, d100$y))), col = mycolors[1], xlab = "", ylab = "")
        lines(d10, col = mycolors[2])
        lines(d100, col = mycolors[3])
        axl <- par("usr")
        text(axl[1] + 0.1 * (axl[2] - axl[1]), axl[4] - 0.1 * (axl[4] - axl[3]),mylabs[count + 2], pos = 4, cex = 0.5)
        count <- count + 3
    }
}


#test: is facilitation model doing what it should? Plot ODE solutions for a few iterations, scaling m0 and m2
#Example:
#load(file = paste(odefaciloutputname, "_typicalrho.RData", sep = ""))
#checkfacilitation(a0, a1, a2, deltab0, b1, b2, kA, kO, m0, m2, niter = 1, m0scale = 1, m2scale = 1, tA1dates)
checkfacilitation <- function(a0, a1, a2, deltab0, b1, b2, kA, kO, m0, m2, niter, m0scale, m2scale, tA1dates){
    dayzero = as.Date("2019-07-30")
    rhoA1 <- array(dim = c(niter, 15, 3)) #A at 1 m, without facilitation
    rhoA1facil <- array(dim = c(niter, 15, 3)) #A at 1 m, with facilitation
    depthcode <- 1
    treatmentapplied <- c(0, rep(c(0, 2), 7)) #A treatment typically not applied in week 1
    timeblocks <- array(-1, dim = c(7, 8))
    timeblocks[1, 1:2] <- 1:2
    timeblocks[2:7, 1] <- 3:8
    ntimesinblock <- c(2, rep(1, 6))
    ntimeblocks <- 7
    for(i in 1:niter){
        m0i <- m0[i, depthcode] * m0scale
        m2i <- m2[i, depthcode] * m2scale

        print(m2i)
        
        rhoA1[i, , ] <-  buildrho(a0 = a0[i, depthcode], a1 = a1[i, depthcode], a2 = a2[i, depthcode], deltab0 = deltab0[i, depthcode], b1 = b1[i, depthcode], b2 = b2[i, depthcode], kA = kA[i], kO = kO[i], m0 = 0, m2 = 0, treatmentapplied = treatmentapplied, timeblocks = timeblocks, ntimeblocks = ntimeblocks, ntimesinblock = ntimesinblock)
        rhoA1facil[i, , ] <-  buildrho(a0 = a0[i, depthcode], a1 = a1[i, depthcode], a2 = a2[i, depthcode], deltab0 = deltab0[i, depthcode], b1 = b1[i, depthcode], b2 = b2[i, depthcode], kA = kA[i], kO = kO[i], m0 = m0i, m2 = m2i, treatmentapplied = treatmentapplied, timeblocks = timeblocks, ntimeblocks = ntimeblocks, ntimesinblock = ntimesinblock)
    }

    par(mfrow = c(3, 1))
    par(mar = c(4, 4, 2, 2))
    mycolors <- brewer.pal(5, "Dark2")[c(4, 5)]
    mylty <- c("dashed", NA, "solid")
    plot(tA1dates, rep(0.5, length(tA1dates)), ylim =c(0, 1), xlab = "date", ylab = expression(italic(A.~aurita)), type = "n")
    legend("topright", bty = "n", legend = c(expression(paste(1~m, ",", ~italic(A))), expression(paste(1~m, ",", ~italic(A)~facil))), col = mycolors, lty = rep(mylty[c(1, 3)], 1), pch = NA)
    axl <- par("usr")
    text(axl[1] + 0.05 * (axl[2] - axl[1]), axl[4] - 0.2 * (axl[4] - axl[3]), "a")
    for(i in 1:niter){
        lines(tA1dates, c(0, rhoA1[i, , 1]), lty = "dashed", col = mycolors[1])
        lines(tA1dates, c(0, rhoA1facil[i, , 1]), lty = "dashed", col = mycolors[2])
    }
    plot(tA1dates, rep(0.5, length(tA1dates)), ylim =c(0, 1), xlab = "date", ylab = "bare panel", type = "n")
    legend("topright", bty = "n", legend = c(expression(paste(1~m, ",", ~italic(A))), expression(paste(1~m, ",", ~italic(A)~facil))), col = mycolors, lty = rep(mylty[c(1, 3)], 1), pch = NA)
    
    axl <- par("usr")
    text(axl[1] + 0.05 * (axl[2] - axl[1]), axl[4] - 0.2 * (axl[4] - axl[3]), "b")
    for(i in 1:niter){
        lines(tA1dates, c(1, rhoA1[i, , 2]), lty = "dashed", col = mycolors[1])
        lines(tA1dates, c(1, rhoA1facil[i, , 2]), lty = "dashed", col = mycolors[2])
    }
    plot(tA1dates, rep(0.5, length(tA1dates)), ylim =c(0, 1), xlab = "date", ylab = "potential competitors", type = "n")
    legend("topright", bty = "n", legend = c(expression(paste(1~m, ",", ~italic(A))), expression(paste(1~m, ",", ~italic(A)~facil))), col = mycolors, lty = rep(mylty[c(1, 3)], 1), pch = NA)
    axl <- par("usr")
    text(axl[1] + 0.05 * (axl[2] - axl[1]), axl[4] - 0.2 * (axl[4] - axl[3]), "c")
    for(i in 1:niter){
        lines(tA1dates, c(0, rhoA1[i, , 3]), lty = "dashed", col = mycolors[1])
        lines(tA1dates, c(0, rhoA1facil[i, , 3]), lty = "dashed", col = mycolors[2])
    }
    print("No facil:")
    print(rhoA1[1, , ])
    print("Facil:")
    print(rhoA1facil[1, , ])
}

#check effect of A. aurita on population growth rate of potential competitors, for posterior mean parameters at 1 m
checkfacilrate <- function(a0, a1, a2, m0, m2, deltab0, b1, b2){
    ny1 <- 100
    y1star <- seq(from = 0, to = 1, length.out = ny1)
    xstarset <- c(0, 0.05, 0.1, 0.5)
    parametersfacil <- c(a0 = mean(a0[, 1]), a1 = mean(a1[, 1]), a2 = mean(a2[, 1]), m0 = mean(m0[, 1]), m2 = mean(m2[, 1]), deltab0 = mean(deltab0[, 1]), b1 = mean(b1[, 1]), b2 = mean(b2[, 1])) #parameters scaled by sampling interval
    parametersnofacil <- c(a0 = mean(a0[, 1]), a1 = mean(a1[, 1]), a2 = mean(a2[, 1]), m0 = 0, m2 = 0, deltab0 = mean(deltab0[, 1]), b1 = mean(b1[, 1]), b2 = mean(b2[, 1])) #parameters scaled by sampling interval
    mycolors <- brewer.pal(5, "Dark2")[c(4, 5)]
    par(mfrow = c(2, 2))
    for(xstar in xstarset){
        dxdtfacil <- array(dim = c(ny1, 1))
        dxdtnofacil <- array(dim = c(ny1, 1))
        for(i in 1:ny1){
            if(y1star[i] + xstar <= 1){
                state <- c(xstar = xstar, y1star = y1star[i])
                t <- 0
                rates <- aurelia2dODEdim(t, state, parametersfacil)[[1]]
                dxdtfacil[i] <- rates[1]
                rates <- aurelia2dODEdim(t, state, parametersnofacil)[[1]]
                dxdtnofacil[i] <- rates[1]
            }
        }
        plot(y1star, dxdtfacil, type = "l", col = mycolors[2], xlab = expression(italic(A.~aurita)), ylab = expression(dx / dt), ylim = c(-0.1, 0.4))
        lines(y1star, dxdtnofacil, col = mycolors[1])
        abline(h = 0, lty = "dashed")
        axl <- par("usr")
        text(axl[1] + 0.1 * (axl[2] - axl[1]), axl[3] + 0.1 * (axl[4] - axl[3]), paste("x = ", bquote(.(xstar)), sep = ""), pos = 4)
    }
    legend("topright", bty = "n", legend = c("no facilitation", "facilitation"), pch = NA, lty = c("solid", "solid"), col = mycolors)
}

#check effect of A. aurita on proportional settlement and death of potential competitors, for posterior mean parameters at 1 m
plotfacileffects <- function(a0, a1, a2, m0, m2, deltab0, b1, b2){
    ny1 <- 100
    y1star <- seq(from = 0, to = 1, length.out = ny1)
    meana2 <- mean(a2[, 1])
    meanm2 <- mean(m2[, 1])
    meana0 <- mean(a0[, 1])
    meanm0 <- mean(m0[, 1])
    propdeathfacil <- meana2 * exp(- meanm2 * y1star)
    par(mfrow = c(2, 1))
    mycolors <- brewer.pal(5, "Dark2")[c(4, 5)]
    plot(y1star, propdeathfacil, type = "l", col = mycolors[2], xlab = expression(italic(A.~aurita)), ylab = "proportional death rate")
    lines(y1star, rep(meana2, ny1), col = mycolors[1])
    legend("topleft", bty = "n", legend = c("no facilitation", "facilitation"), pch = NA, lty = c("solid", "solid"), col = mycolors)
    propsettlefacil <- meana0 + meanm0 * y1star
    plot(y1star, propsettlefacil, type = "l", col = mycolors[2], xlab = expression(italic(A.~aurita)), ylab = "proportional settlement rate")
    lines(y1star, rep(meana0, ny1), col = mycolors[1])
}

#check whether we were accidentally removing potential competitors in A treatment: plot Post vs Pre count of potential competitors in A treatment on each date
#Arguments:
#dynamicdata: dataframe: one of the items returned by datafordynamicmodel(): has treatmentapplied variable that tells whether a treatment was actually done, so that there are Pre and Post observations
#Value: pdf fname: scatter plot for each date: potential competitors in A treatment, post vs pre, with line of slope 1, intercept 0
plotAremovaleffects <- function(dynamicdata, fname){
    pdf(file = fname, width = 7, height = 7)
    par(mfrow = c(4, 2))
    par(mar = c(4, 4, 2, 2))
    dates <- unique(dynamicdata$date)
    for(i in 1:length(dates)){
        date <- dates[i]
        Adatapost <- dynamicdata[dynamicdata$treatmentapplied == "A" & dynamicdata$prepost == "Post" & dynamicdata$date == date, ]
        if(dim(Adatapost)[1] > 0){
            mypanels <- Adatapost$panel
            Adatapre <- dynamicdata[dynamicdata$prepost == "Pre" & dynamicdata$date == date, ]
            Adatapre <- Adatapre[Adatapre$panel %in% mypanels, ]
            presum <- rowSums(Adatapre[, 7:15])
            postsum <- rowSums(Adatapost[, 7:15])
            plot(Adatapre$potentialcomp / presum, Adatapost$potentialcomp / postsum, xlim = c(0, 1), ylim = c(0, 1), xlab = "pre-treatment", ylab = "post-treatment", pch = 16, col = adjustcolor("black", 0.4))
            abline(a = 0, b = 1)
        } else {
            plot(c(0, 1), c(0, 1), type = "n", xlab = "pre-treatment", ylab = "post-treatment")
        }
        text(0.15, 0.9, paste(letters[i], ": ", date, sep = ""))
    }
    dev.off()
}

#check whether proportion A. aurita removed matches model estimates
#Arguments:
#dynamicdata: dataframe: one of the items returned by datafordynamicmodel(): has treatmentapplied variable that tells whether a treatment was actually done, so that there are Pre and Post observations
#fname: name of pdf output file
#odeoutputname, odefacilitationoutputname, odeprotectionoutputname, odeovergrowthoutputname, odegrowthfaciloutputname: names of .RData files for ODE fits with basic, settlement facilitation, protection, overgrowth and growth facilitation models respectively
#Value: pdf fname: scatter plot for each date: A. aurita in A treatment, post vs pre, with lines representing estimated proportions removed under each model
plotAremovalproportion <- function(dynamicdata, fname, odeoutputname, odefacilitationoutputname, odeprotectionoutputname, odeovergrowthoutputname, odegrowthfaciloutputname){
    mycolors <- brewer.pal(5, "Dark2")
    pdf(file = fname, width = 7, height = 7)
    par(mfrow = c(4, 2))
    par(mar = c(4, 4, 2, 2))
    dates <- unique(dynamicdata$date)
    for(i in 1:length(dates)){
        date <- dates[i]
        Adatapost <- dynamicdata[dynamicdata$treatmentapplied == "A" & dynamicdata$prepost == "Post" & dynamicdata$date == date, ]
        if(dim(Adatapost)[1] > 0){
            mypanels <- Adatapost$panel
            Adatapre <- dynamicdata[dynamicdata$prepost == "Pre" & dynamicdata$date == date, ]
            Adatapre <- Adatapre[Adatapre$panel %in% mypanels, ]
            presum <- rowSums(Adatapre[, 7:15])
            postsum <- rowSums(Adatapost[, 7:15])
            plot(Adatapre$Aurelia.aurita.on.panel / presum, Adatapost$Aurelia.aurita.on.panel / postsum, xlim = c(0, 0.3), ylim = c(0, 0.3), xlab = "pre-treatment", ylab = "post-treatment", pch = 16, col = adjustcolor("black", 0.4))
            load(file = paste(odeoutputname, ".RData", sep = ""))
            abenvelope(kA = kA, color = mycolors[1])
            load(file = paste(odefacilitationoutputname, ".RData", sep = ""))
            abenvelope(kA = kA, color = mycolors[2])
            load(file = paste(odegrowthfaciloutputname, ".RData", sep = ""))
            abenvelope(kA = kA, color = mycolors[3])
            load(file = paste(odeovergrowthoutputname, ".RData", sep = ""))
            abenvelope(kA = kA, color = mycolors[4])
            load(file = paste(odeprotectionoutputname, ".RData", sep = ""))
            abenvelope(kA = kA, color = mycolors[5])
        } else {
            plot(c(0, 0.3), c(0, 0.3), type = "n", xlab = "pre-treatment", ylab = "post-treatment")
        }
        if(i == 1) {
            legend("topright", legend = c("basic", "settlement facilitation", "growth facilitation", "overgrowth", "protection"), pch = rep(NA, 5), lty = rep("solid", 5), col = adjustcolor(mycolors, 0.5), lwd = 2, bty = "n")
        }
        text(0.045, 0.27, paste(letters[i], ": ", date, sep = ""))
    }
    dev.off()
}

#plot a line through the origin with slope kA, and envelope representing uncertainty
#Arguments:
#kA: vector of length iterations: posterior sample from kA
#color: color for line
#Value: draws a line through the origin with slope kA, and an envelope representing all lines through origin with slopes in 95% hpd interval for kA
abenvelope  <- function(kA, color){
        xmean <- mean(kA)
        xhdr <- hdr(kA, prob = 95)$hdr
        polygon(c(0, 1, 1, 0), c(0, xhdr[1], xhdr[2], 0), col = adjustcolor(color, 0.2), border = NA)
        abline(a = 0, b = xmean, col = adjustcolor(color, 0.5))
}

#compare ode models using loo
#Arguments:
#odeoutputname: name of files containing basic ode model (the .RData will have this name plus .RData extension)
#odefacilitationoutputname: name of .RData file containing ode model with facilitation
#odeprotectionoutputname: name of .RData file containing ode model with protection
#odeovergrowthoutputname: name of .RData file containing ode model with overgrowth
#odegrowthfaciloutputname: name of .RData file containing ode model with growth facilitation
#Value: prints out table of model comparison using loo_compare(). Model order: the order in which the models are given as arguments to loo_compare().
odeloo <- function(odeoutputname, odefacilitationoutputname, odeprotectionoutputname, odeovergrowthoutputname, odegrowthfaciloutputname){
    load(file = paste(odeoutputname, ".RData", sep = ""))
    load(file = paste(odefacilitationoutputname, ".RData", sep = ""))
    load(file = paste(odeprotectionoutputname, ".RData", sep = ""))
    load(file = paste(odeovergrowthoutputname, ".RData", sep = ""))
    load(file = paste(odegrowthfaciloutputname, ".RData", sep = ""))
    print(loo_compare(looode, loofacilitation, looprotection, looovergrowth, loogrowthfacil), simplify = FALSE)
}

#Proportional rates of change in ODE model
#state: vector (x, y1star): proportions of space covered by potential competitors and polyps
#a0: potential competitor settlement rate
#a1: potential competitor growth rate
#a2: potential competitor death rate
#deltab0: polyp settlement rate * polyp area
#b1: polyp budding rate
#b2: polyp death rate
#m0: settlement facilitation (default zero)
#m1: growth facilitation rate (default zero)
#a1y1star: rate of overgrowth (default zero)
#m2: predator protection parameter (default zero)
#Value:
#Vector of proportional rates of change evaluated at given state and parameters
odeproportionalchange <- function(state, a0, a1, a2, deltab0, b1, b2, m0 = 0, m1 = 0, a1y1star = 0, m2 = 0){
    x  <-  state[1]
    y1star <- state[2]
    Pi1 <- a0 / deltab0
    Pi2  <- a1 / deltab0
    Pi3 <- a2 / deltab0
    Pi4 <- b1 / deltab0
    Pi5 <- b2 / deltab0
    Pi11 <- m0 / deltab0
    Pi12  <- m1 / deltab0
    Pi13 <- a1y1star / deltab0
    f <- 1 / x * ((Pi1 + Pi11 * y1star) * (1 - x - y1star) +  (Pi2 + Pi12 * y1star) * x * (1 - x - y1star) + Pi13 * x * y1star + Pi3 * exp(-m2 * y1star) * x)
    g <- 1 / y1star * ((1 - x - y1star) + Pi4 * y1star * (1 - x - y1star) - Pi13 * x * y1star + Pi5 * y1star)
    return(c(f, g))
}

#Proportional rates of change in ODE model with polyps on potential competitors
#state: vector (x, y1star, y2star): proportions of space covered by potential competitors and polyps, and area filled by polyps on potential competitors per unit substrate area
#a0: potential competitor settlement rate
#a1: potential competitor growth rate
#a2: potential competitor death rate
#deltab0: polyp settlement rate * polyp area
#b1: polyp budding rate
#b2: polyp death rate
#b3: budding by polyps on potential competitors onto substrate
#c0: settlement rate of polyps on potential competitors
#c1: budding of polyps on potential competitors onto potential competitors
#c2: polyps on potential competitors death rate
#c3: budding by polyps on substrate onto potential competitors
#psi: surface area of potential competitors available for settlement per unit substrate occupied by them
#Value:
#Vector of proportional rates of change evaluated at given state and parameters
odeproportionalchangethreestate <- function(state, a0, a1, a2, deltab0, b1, b2, b3, c0, c1, c2, c3, psi){
    x  <-  state[1]
    y1star <- state[2]
    y2star  <- state[3]
    Pi1 <- a0 / deltab0
    Pi2  <- a1 / deltab0
    Pi3 <- a2 / deltab0
    Pi4 <- b1 / deltab0
    Pi5 <- b2 / deltab0
    Pi6 <- b3 / deltab0
    Pi7 <- c0 / deltab0
    Pi8 <- c1 / deltab0
    Pi9 <- c2 / deltab0
    Pi10 <- c3 / deltab0
    f <- 1 / x * (Pi1 * (1 - x - y1star) +  Pi2 * x * (1 - x - y1star) + Pi3 * x)
    g <- 1 / y1star * ((1 - x - y1star) + Pi4 * y1star * (1 - x - y1star) + Pi5 * y1star + Pi6 * y2star * (1 - x - y1star))
    h <-  1 / y2star * (Pi7 * (psi * x - y2star) + Pi8 * y2star * (psi * x - y2star) + Pi3 * psi * x * y2star + Pi9 * y2star + Pi10 * y1star * (psi * x - y2star))
    return(c(f, g, h))
}


#community matrix for basic ODE model
#Arguments:
#x, y1star: proportions of space covered by potential competitors and polyps
#a0: potential competitor settlement rate
#a1: potential competitor growth rate
#a2: potential competitor death rate
#deltab0: polyp settlement rate * polyp area
#b1: polyp budding rate
#b2: polyp death rate
#Value:
#community matrix evaluated at given state and parameters
cmbasic <- function(x, y1star, a0, a1, a2, deltab0, b1, b2){
    Pi1 <- a0 / deltab0
    Pi2  <- a1 / deltab0
    Pi3 <- a2 / deltab0
    Pi4 <- b1 / deltab0
    Pi5 <- b2 / deltab0
    matrix(c(-Pi1 * (1 - y1star) / x^2 - Pi2, -1 / y1star - Pi4, -Pi1 / x - Pi2, - (1 - x) / y1star^2 - Pi4), nrow = 2, ncol = 2)
}

#community matrix for ODE model with settlement facilitation
#Arguments:
#x, y1star: proportions of space covered by potential competitors and polyps
#a0: potential competitor settlement rate
#a1: potential competitor growth rate
#a2: potential competitor death rate
#deltab0: polyp settlement rate * polyp area
#b1: polyp budding rate
#b2: polyp death rate
#m0: settlement facilitation
#Value:
#community matrix evaluated at given state and parameters
cmsettlement <- function(x, y1star, a0, a1, a2, deltab0, b1, b2, m0){
    Pi1 <- a0 / deltab0
    Pi2  <- a1 / deltab0
    Pi3 <- a2 / deltab0
    Pi4 <- b1 / deltab0
    Pi5 <- b2 / deltab0
    Pi11 <- m0 / deltab0
    matrix(c(-(Pi1 + Pi11 * y1star) * (1 - y1star) / x^2 - Pi2, -1 / y1star - Pi4, -Pi1 / x - Pi2 + Pi11 / x * (1 - x - 2 * y1star), - (1 - x) / y1star^2 - Pi4), nrow = 2, ncol = 2)
}

#community matrix for ODE model with growth facilitation
#Arguments:
#x, y1star: proportions of space covered by potential competitors and polyps
#a0: potential competitor settlement rate
#a1: potential competitor growth rate
#a2: potential competitor death rate
#deltab0: polyp settlement rate * polyp area
#b1: polyp budding rate
#b2: polyp death rate
#Value:
#community matrix evaluated at given state and parameters
cmgrowthfacil <- function(x, y1star, a0, a1, a2, deltab0, b1, b2, m1){
    Pi1 <- a0 / deltab0
    Pi2  <- a1 / deltab0
    Pi3 <- a2 / deltab0
    Pi4 <- b1 / deltab0
    Pi5 <- b2 / deltab0
    Pi12  <- m1 / deltab0
    matrix(c(-Pi1 * (1 - y1star) / x^2 - Pi2 - Pi12 * y1star, -1 / y1star - Pi4, -Pi1 / x - Pi2 + Pi12 * (1 - x) - 2 * Pi12 * y1star, - (1 - x) / y1star^2 - Pi4), nrow = 2, ncol = 2)
}

#community matrix for ODE model with overgrowth
#Arguments:
#x, y1star: proportions of space covered by potential competitors and polyps
#a0: potential competitor settlement rate
#a1: potential competitor growth rate
#a2: potential competitor death rate
#deltab0: polyp settlement rate * polyp area
#b1: polyp budding rate
#b2: polyp death rate
#a1y1star: overgrowth rate
#Value:
#community matrix evaluated at given state and parameters
cmovergrowth <- function(x, y1star, a0, a1, a2, deltab0, b1, b2, a1y1star){
    Pi1 <- a0 / deltab0
    Pi2  <- a1 / deltab0
    Pi3 <- a2 / deltab0
    Pi4 <- b1 / deltab0
    Pi5 <- b2 / deltab0
    Pi13  <- a1y1star / deltab0
    matrix(c(-Pi1 * (1 - y1star) / x^2 - Pi2, -1 / y1star - Pi4 - Pi13, -Pi1 / x - Pi2 + Pi13, - (1 - x) / y1star^2 - Pi4), nrow = 2, ncol = 2)
}

#community matrix for ODE model with protection from predators
#Arguments:
#x, y1star: proportions of space covered by potential competitors and polyps
#a0: potential competitor settlement rate
#a1: potential competitor growth rate
#a2: potential competitor death rate
#deltab0: polyp settlement rate * polyp area
#b1: polyp budding rate
#b2: polyp death rate
#m2: predator protection parameter
#Value:
#community matrix evaluated at given state and parameters
cmprotection <- function(x, y1star, a0, a1, a2, deltab0, b1, b2, m2){
    Pi1 <- a0 / deltab0
    Pi2  <- a1 / deltab0
    Pi3 <- a2 / deltab0
    Pi4 <- b1 / deltab0
    Pi5 <- b2 / deltab0
    matrix(c(-Pi1 * (1 - y1star) / x^2 - Pi2, -1 / y1star - Pi4, -Pi1 / x - Pi2 - m2 * Pi3 * exp(-m2 * y1star), - (1 - x) / y1star^2 - Pi4), nrow = 2, ncol = 2)
}

#community matrix for ODE model with polyps on potential competitors
#Arguments:
#x, y1star, y2star: proportions of space covered by potential competitors, polyps on substrate, polyps on potential competitors
#a0: potential competitor settlement rate
#a1: potential competitor growth rate
#a2: potential competitor death rate
#deltab0: polyp settlement rate * polyp area
#b1: polyp budding rate
#b2: polyp death rate
#b3: budding by polyps on potential competitors onto substrate
#c0: settlement rate of polyps on potential competitors
#c1: budding of polyps on potential competitors onto potential competitors
#c2: polyps on potential competitors death rate
#c3: budding by polyps on substrate onto potential competitors
#psi: surface area of potential competitors available for settlement per unit substrate occupied by them
#Value:
#community matrix evaluated at given state and parameters
cmthreestate <- function(x, y1star, y2star, a0, a1, a2, deltab0, b1, b2, b3, c0, c1, c2, c3, psi){
    Pi1 <- a0 / deltab0
    Pi2  <- a1 / deltab0
    Pi3 <- a2 / deltab0
    Pi4 <- b1 / deltab0
    Pi5 <- b2 / deltab0
    Pi6 <- b3 / deltab0
    Pi7 <- c0 / deltab0
    Pi8 <- c1 / deltab0
    Pi9 <- c2 / deltab0
    Pi10 <- c3 / deltab0
    matrix(c(-Pi1 * (1 - y1star) / x^2 - Pi2, - (1 + Pi6 * y2star) / y1star - Pi4, psi * ((Pi7 + Pi10 * y1star) / y2star + Pi3 + Pi8), -Pi1 / x - Pi2, - (1 - x) * (1 + Pi6 * y2star) / (y1star)^2 - Pi4, Pi10 * (psi * x - y2star) / y2star, 0, Pi6 * (1 - x - y1star) / y1star, -psi * x * (Pi7 + Pi10 * y1star) / y2star^2 - Pi8), nrow = 3, ncol = 3)
}


#Numerical check on community matrices for ODE models
#Arguments:
#x, y1star: proportions of space covered by potential competitors and polyps
#y2star: proportion space covered by polyps on potential competitors per unit area: may be greater than 1 (default 0.1)
#psi: area available for settlement on potential competitors per unit area covered by potential competiors (default 2)
#odeoutputname, odefacilitationoutputname, odegrowthfaciloutputname, odeovergrowthoutputname, odeprotectionoutputname: names of .RData files with results from basic, settlement facilitation, growth facilitation, overgrowth, protection models respectively
#Value: prints out numerical and analytical community matrices evaluated at (x, y1star), for mean parameter values over both depths
checkcommunitymatrix <- function(x, y1star, y2star = 0.1, psi = 2, odeoutputname, odefacilitationoutputname, odegrowthfaciloutputname, odeovergrowthoutputname, odeprotectionoutputname){
    state <- c(x, y1star)
    load(file = paste(odeoutputname, ".RData", sep = ""))
    print("Basic model")
    print(paste("Numerical community matrix at mean parameters across depths, x =", x, ", y1star =", y1star, ":"))
    print(jacobian(func = odeproportionalchange, x = state, a0 = mean(a0), a1 = mean(a1), a2 = mean(a2), deltab0 = mean(deltab0), b1 = mean(b1), b2 = mean(b2)))
    print("Analytical community matrix:")
    print(cmbasic(x = x, y1star = y1star, a0 = mean(a0), a1 = mean(a1), a2 = mean(a2), deltab0 = mean(deltab0), b1 = mean(b1), b2 = mean(b2)))

    print("Model with polyps on potential competitors")
    print(paste("Numerical community matrix at mean parameters across depths, x =", x, ", y1star =", y1star, ":"))
    print(jacobian(func = odeproportionalchangethreestate, x = c(state, y2star), a0 = mean(a0), a1 = mean(a1), a2 = mean(a2), deltab0 = mean(deltab0), b1 = mean(b1), b2 = mean(b2), b3 = mean(b1), c0 = mean(deltab0), c1 = mean(b1), c2 = mean(b2), c3 = mean(b1), psi = psi))
    print("Analytical community matrix:")
    print(cmthreestate(x = x, y1star = y1star, y2star = y2star, a0 = mean(a0), a1 = mean(a1), a2 = mean(a2), deltab0 = mean(deltab0), b1 = mean(b1), b2 = mean(b2), b3 = mean(b1), c0 = mean(deltab0), c1 = mean(b1), c2 = mean(b2), c3 = mean(b1), psi = psi)) #we never fitted this model: parameters as in basic model, plus arbitrary choices for others: psi = 2: potential competitors are hemispheres; b3 = mean(b1), c1 = mean(b1), c3 = mean(b1): same budding to and from substrate and potential comp; c0 = mean(deltab0): same settlement on potential comp as substrate; c2 = mean(b2): same death on comp as substrate
        
    load(file = paste(odefacilitationoutputname, ".RData", sep = ""))
    print("------------------------------")
    print("Settlement facilitation model")
    print(paste("Numerical community matrix at mean parameters across depths, x =", x, ", y1star =", y1star, ":"))
    print(jacobian(func = odeproportionalchange, x = state, a0 = mean(a0), a1 = mean(a1), a2 = mean(a2), deltab0 = mean(deltab0), b1 = mean(b1), b2 = mean(b2), m0 = mean(m0)))
    print("Analytical community matrix:")
    print(cmsettlement(x = x, y1star = y1star, a0 = mean(a0), a1 = mean(a1), a2 = mean(a2), deltab0 = mean(deltab0), b1 = mean(b1), b2 = mean(b2), m0 = mean(m0)))

    load(file = paste(odegrowthfaciloutputname, ".RData", sep = ""))
    print("------------------------------")
    print("Growth facilitation model")
    print(paste("Numerical community matrix at mean parameters across depths, x =", x, ", y1star =", y1star, ":"))
    print(jacobian(func = odeproportionalchange, x = state, a0 = mean(a0), a1 = mean(a1), a2 = mean(a2), deltab0 = mean(deltab0), b1 = mean(b1), b2 = mean(b2), m1 = mean(m1)))
    print("Analytical community matrix:")
    print(cmgrowthfacil(x = x, y1star = y1star, a0 = mean(a0), a1 = mean(a1), a2 = mean(a2), deltab0 = mean(deltab0), b1 = mean(b1), b2 = mean(b2), m1 = mean(m1)))

    load(file = paste(odeovergrowthoutputname, ".RData", sep = ""))
    print("------------------------------")
    print("Overgrowth model")
    print(paste("Numerical community matrix at mean parameters across depths, x =", x, ", y1star =", y1star, ":"))
    print(jacobian(func = odeproportionalchange, x = state, a0 = mean(a0), a1 = mean(a1), a2 = mean(a2), deltab0 = mean(deltab0), b1 = mean(b1), b2 = mean(b2), a1y1star = mean(a1y1star)))
    print("Analytical community matrix:")
    print(cmovergrowth(x = x, y1star = y1star, a0 = mean(a0), a1 = mean(a1), a2 = mean(a2), deltab0 = mean(deltab0), b1 = mean(b1), b2 = mean(b2), a1y1star = mean(a1y1star)))

    load(file = paste(odeprotectionoutputname, ".RData", sep = ""))
    print("------------------------------")
    print("Protection model")
    print(paste("Numerical community matrix at mean parameters across depths, x =", x, ", y1star =", y1star, ":"))
    print(jacobian(func = odeproportionalchange, x = state, a0 = mean(a0), a1 = mean(a1), a2 = mean(a2), deltab0 = mean(deltab0), b1 = mean(b1), b2 = mean(b2), m2 = mean(m2)))
    print("Analytical community matrix:")
    print(cmprotection(x = x, y1star = y1star, a0 = mean(a0), a1 = mean(a1), a2 = mean(a2), deltab0 = mean(deltab0), b1 = mean(b1), b2 = mean(b2), m2 = mean(m2)))
    
}

#extract the ODE parameters that are in all models to a printable matrix
#Arguments: fname is name of file containing posterior samples from parameters
#Value: a 14 x 1 matrix containing formatted posterior means and 95% hpd regions for parameters a0, a1, a2, deltab0, b1, b2 (each at 1 m and then at 3 m), rA, rO
basicODEparameters <- function(fname, lambda = 1){
    load(file = paste(fname, ".RData", sep = ""))
    a0summary <- posteriorsummaryvector(a0)
    a1summary <- posteriorsummaryvector(a1)
    a2summary <- posteriorsummaryvector(a2)
    deltab0summary <- posteriorsummaryvector(deltab0)
    b1summary <- posteriorsummaryvector(b1)
    b2summary <- posteriorsummaryvector(b2)
    rA <- 1 - kA
    rO <- 1 - kO
    rAsummary <- posteriorsummaryvector(rA)
    rOsummary <- posteriorsummaryvector(rO)
    mytable <- array(dim = c(14, 1))
    mytable[1, 1] <- extractposteriorsummary(a0summary, 1, format = "%.2g")
    mytable[2, 1] <- extractposteriorsummary(a0summary, 2, format = "%.2g")
    mytable[3, 1] <- extractposteriorsummary(a1summary, 1, format = "%.2g")
    mytable[4, 1] <- extractposteriorsummary(a1summary, 2, format = "%.2g")
    mytable[5, 1] <- extractposteriorsummary(a2summary, 1, format = "%.2g")
    mytable[6, 1] <- extractposteriorsummary(a2summary, 2, format = "%.2g")
    mytable[7, 1] <- extractposteriorsummary(deltab0summary, 1, format = "%.2g")
    mytable[8, 1] <- extractposteriorsummary(deltab0summary, 2, format = "%.2g")
    mytable[9, 1] <- extractposteriorsummary(b1summary, 1, format = "%.2g")
    mytable[10, 1] <- extractposteriorsummary(b1summary, 2, format = "%.2g")
    mytable[11, 1] <- extractposteriorsummary(b2summary, 1, format = "%.2g")
    mytable[12, 1] <- extractposteriorsummary(b2summary, 2, format = "%.2g")
    mytable[13, 1] <- extractposteriorsummary(rAsummary, 1, format = "%.2g")
    mytable[14, 1] <- extractposteriorsummary(rOsummary, 1, format = "%.2g")
    return(mytable)
}

#produce a LaTeX table summarizing posterior distributions of ODE parameters
#Arguments:
#odeoutputname, odefacilitationoutputname, odegrowthfaciloutputname, odeovergrowthoutputname, odeprotectionoutputname: names of .RData files with results from basic, settlement facilitation, growth facilitation, overgrowth, protection models respectively
#fname: name of file to which LaTeX output should be written
#Value:
#LaTeX code for table of posterior means and 95% HPD intervals from each model, written to fname
printODEparameters <- function(odeoutputname, odefacilitationoutputname, odegrowthfaciloutputname, odeovergrowthoutputname, odeprotectionoutputname, fname){
    mytable <- array(dim = c(22, 5))
    rownames(mytable) <- c("$a_0$, \\SI{1}{\\metre}", "$a_0$, \\SI{3}{\\metre}", "$a_1$, \\SI{1}{\\metre}", "$a_1$, \\SI{3}{\\metre}", "$a_2$, \\SI{1}{\\metre}", "$a_2$, \\SI{3}{\\metre}", "$\\delta b_0$, \\SI{1}{\\metre}", "$\\delta b_0$, \\SI{3}{\\metre}", "$b_1$, \\SI{1}{\\metre}", "$b_1$, \\SI{3}{\\metre}", "$b_2$, \\SI{1}{\\metre}", "$b_2$, \\SI{3}{\\metre}", "$r_A$", "$r_O$", "$m_0$, \\SI{1}{\\metre}", "$m_0$, \\SI{3}{\\metre}", "$m_1$, \\SI{1}{\\metre}", "$m_1$, \\SI{3}{\\metre}", "$a_{1, y_1^*}$, \\SI{1}{\\metre}", "$a_{1, y_1^*}$, \\SI{3}{\\metre}", "$m_2$, \\SI{1}{\\metre}", "$m_2$, \\SI{3}{\\metre}")
    colnames(mytable) <- c("basic", "settlement facilitation", "growth facilitation", "overgrowth", "protection")
    mytable[1:14, 1] <- basicODEparameters(fname = odeoutputname)
    mytable[1:14, 2] <- basicODEparameters(fname = odefacilitationoutputname)
    mytable[1:14, 3] <- basicODEparameters(fname = odegrowthfaciloutputname)
    mytable[1:14, 4] <- basicODEparameters(fname = odeovergrowthoutputname)
    mytable[1:14, 5] <- basicODEparameters(fname = odeprotectionoutputname)

    load(file = paste(odefacilitationoutputname, ".RData", sep = ""))
    m0summary <- posteriorsummaryvector(m0)
    mytable[15, 2] <- extractposteriorsummary(m0summary, 1, format = "%.2g")
    mytable[16, 2] <- extractposteriorsummary(m0summary, 2, format = "%.2g")

    load(file = paste(odegrowthfaciloutputname, ".RData", sep = ""))
    m1summary <- posteriorsummaryvector(m1)
    mytable[17, 3] <- extractposteriorsummary(m1summary, 1, format = "%.2g")
    mytable[18, 3] <- extractposteriorsummary(m1summary, 2, format = "%.2g")

    load(file = paste(odeovergrowthoutputname, ".RData", sep = ""))
    a1y1starsummary <- posteriorsummaryvector(a1y1star)
    mytable[19, 4] <- extractposteriorsummary(a1y1starsummary, 1, format = "%.2g")
    mytable[20, 4] <- extractposteriorsummary(a1y1starsummary, 2, format = "%.2g")

    load(file = paste(odeprotectionoutputname, ".RData", sep = ""))
    m2summary <- posteriorsummaryvector(m2)
    mytable[21, 5] <- extractposteriorsummary(m2summary, 1, format = "%.2g")
    mytable[22, 5] <- extractposteriorsummary(m2summary, 2, format = "%.2g")

    xt <- xtable(mytable)
    print(xt, sanitize.text.function = function(x){x}, file = fname)
}

#Bayesian one-sample 95% credible interval for normally-distributed data
#Arguments: vector of observations y, assumed to be independent and identically normally distributed
#Value: list containing sample mean ybar, central 95% credible interval mu95 under noninformative uniform prior on (mu, log sigma)
#Note: this corresponds to the standard one-sample t-interval
#Source: Gelman et al., Bayesian Data Analysis, 2nd ed, pp. 76-78
onesample <- function(y) {
    y <- y[!is.na(y)]
    ybar <- mean(y)
    s <- sd(y)
    n <- length(y)
    tstar <- qt(0.975, df = n - 1)
    mu95 <- c(ybar - tstar * s / sqrt(n), ybar + tstar * s / sqrt(n))
    return(list(ybar = ybar, mu95 = mu95))
}

#plots and credible intervals for environmental data
#Arguments:
#envdata, a data frame containing the variables depthm, salinitypsu, temperaturedegreesC, oxygenmgperl. Note that values are assumed to be stacked in the same date order for each of 1 m and 3 m depth
#envfigname: name of pdf file to which plots will be written
#Value: prints out sample mean and 95% central credible interval for each environmental variable under iid normal model with uninformative uniform priors on (mu, log sigma)
#Also writes pdf envfigname
envanalyses <- function(envdata, envfigname){
    if(any(!(envdata$Date[envdata$depthm == 3] == envdata$Date[envdata$depthm == 1]))) print("Warning: date order does not match between depths")

    oxygen3 <- envdata$oxygenmgperl[envdata$depthm == 3]
    oxygen1 <- envdata$oxygenmgperl[envdata$depthm == 1]
    print("Oxygen 3 m - 1 m:")
    print(onesample(y = oxygen3 - oxygen1))

    temp3 <- envdata$temperaturedegreesC[envdata$depthm == 3]
    temp1 <- envdata$temperaturedegreesC[envdata$depthm == 1]
    print("Temperature 3 m - 1 m:")
    print(onesample(y = temp3 - temp1))

    sal3 <- envdata$salinitypsu[envdata$depthm == 3]
    sal1 <- envdata$salinitypsu[envdata$depthm == 1]
    print("Salinity 3 m - 1 m:")
    print(onesample(y = sal3 - sal1))

    X11()
    par(mfrow = c(3, 1))
    qqnorm(y = oxygen3 - oxygen1, main = "oxygen")
    qqline(y = oxygen3 - oxygen1)
    
    qqnorm(y = temp3 - temp1, main = "temperature")
    qqline(y = temp3 - temp1)

    qqnorm(y = sal3 - sal1, main = "salinity")
    qqline(y = sal3 - sal1)


    pdf(file = envfigname, width = 7, height = 7)
    par(mfrow = c(3, 1))
    par(mar = c(5, 5, 2, 2))
    dates <- as.Date(envdata$Date[envdata$depthm == 3], "%d/%m/%Y")

    plot(dates, oxygen3, type = "l", lwd = 2, lty = "solid", ylim = range(envdata$oxygenmgperl, na.rm = TRUE), xaxt = "n", xlab = "date", ylab = expression(oxygen/mg~l^{-1}))
    axis(side = 1, at = unique(dates), labels = unique(dates))
    lines(dates, oxygen1, lwd = 2, lty = "dashed")
    axlims <- par("usr")
    text(axlims[1] + 0.05 * (axlims[2] - axlims[1]), axlims[4] - 0.2 * (axlims[4] - axlims[3]), "a")
    legend("topright", lwd = c(2, 2), lty = c("dashed", "solid"), pch = c(NA, NA), legend = c("1 m", "3 m"), bty = "n")

    plot(dates, temp3, type = "l", lwd = 2, lty = "solid", ylim = range(envdata$temperaturedegreesC, na.rm = TRUE), xaxt = "n", xlab = "date", ylab = expression(temperature/degree*C))
    axis(side = 1, at = unique(dates), labels = unique(dates))
    lines(dates, temp1, lwd = 2, lty = "dashed")
    axlims <- par("usr")
    text(axlims[1] + 0.05 * (axlims[2] - axlims[1]), axlims[4] - 0.2 * (axlims[4] - axlims[3]), "b")

    plot(dates, sal3, type = "l", lwd = 2, lty = "solid", ylim = range(envdata$salinitypsu, na.rm = TRUE), xaxt = "n", xlab = "date", ylab = "salinity/psu")
    axis(side = 1, at = unique(dates), labels = unique(dates))
    lines(dates, sal1, lwd = 2, lty = "dashed")
    axlims <- par("usr")
    text(axlims[1] + 0.05 * (axlims[2] - axlims[1]), axlims[4] - 0.2 * (axlims[4] - axlims[3]), "c")
    
    dev.off()

}

#simulation-based calibration of ODE model using rstan::sbc
#Arguments:
#odestanfile: name of .stan file specifying model
#dynamicdata: object containing the data needed for the model, generated by datafordynamicmodel()
#M: number of times to condition on draws from prior predictive distribution
#iter: number of iterations (half sampling, half warmup) per simulated data set
#Value: object returned by rstan::sbc
SBCODE <- function(odestanfile, dynamicdata, M, iter){
    odemodel <- stan_model(odestanfile)
    n <- rowSums(dynamicdata$counts)
    sbcodeoutput <- sbc(odemodel, data = list(ntaxa = dynamicdata$ntaxa, npanels = dynamicdata$npanels, sumnt = dynamicdata$sumnt, maxtimeblocks = dynamicdata$tb$maxtimeblocks, maxtimes = dynamicdata$tb$maxtimes, nt = dynamicdata$nt, n = n, ntimeblocks = dynamicdata$tb$ntimeblocks, ntimesinblock = dynamicdata$tb$ntimesinblock, timeblocks = dynamicdata$tb$timeblocks, treatmentapplied = dynamicdata$treatmentapplied, depthcode = dynamicdata$depthcode), M = M, iter = iter, refresh = 0)
    return(sbcodeoutput)
}

#make data for SBC with ODE model
#Arguments:
#SBCODEdatafile: name of file to write data to using stan_rdump
#dymamicdata: object created by datafordynamicmodel()
makeSBCODEdata <- function(SBCODEdatafile, dynamicdata){
    n <- rowSums(dynamicdata$counts)
    ntaxa <- dynamicdata$ntaxa
    npanels <- dynamicdata$npanels
    nt <- dynamicdata$nt
    sumnt <- dynamicdata$sumnt
    times <- dynamicdata$times
    ntimeblocks <- dynamicdata$tb$ntimeblocks
    ntimesinblock <- dynamicdata$tb$ntimesinblock
    timeblocks <- dynamicdata$tb$timeblocks
    maxtimeblocks <- dynamicdata$tb$maxtimeblocks
    maxtimes <- dynamicdata$tb$maxtimes
    treatmentapplied <- dynamicdata$treatmentapplied
    depthcode <- dynamicdata$depthcode
    ddata <- c("n", "ntaxa", "npanels", "nt", "sumnt", "times", "ntimeblocks", "ntimesinblock", "timeblocks", "maxtimeblocks", "maxtimes", "treatmentapplied", "depthcode")
    stan_rdump(list = ddata, file = SBCODEdatafile, quiet = FALSE, append = FALSE)
}


#simulate counts from expected proportions generated from an ODE model by stan
#Arguments:
#dynamicdata: object created by datafordynamicmodel()
#rho: expected proportions in each panel at each time: posterior sample from stan (iterations x stacked panels x taxa)
#niter: number of iterations from which to generate simulated counts by multinomial sampling
#Value:
#simcounts: iterations x stacked panels x taxa array of simulated multinomial counts
simulatefromode <- function(dynamicdata, rho, niter){
    n <- rowSums(dynamicdata$counts)
    simcounts <- array(dim = c(niter, dim(dynamicdata$counts)))
    for(i in 1:niter){
        for(j in 1:dim(dynamicdata$counts)[1]){
            simcounts[i, j, ] <- rmultinom(n = 1, size = n[j], prob = rho[i, j, ])
        }
    }
    return(simcounts)
}

#plot posterior predictive simulations from ODE model, generated by simulatefromode()
#Arguments:
#dynamicdata: object containing the data needed for the model, generated by datafordynamicmodel()
#simdata: array generated by simulatefromode(): iterations x stacked panels x taxa
#alldata: data frame containing treatment variable (assumes that observations sorted by time then panel, so that the first npanels rows give the treatment for each panel in the correct order)
#niter: number of iterations of posterior predictive simulation to plot
#fname: name of pdf file to produce
#Value: pdf file fname: posterior predictive simulation for each panel, multiple iterations overlaid
plotposteriorode <- function(dynamicdata, simdata, alldata, niter, fname){
    n <- rowSums(dynamicdata$counts)
    pdf(file = fname, width = 7, height = 7)
    par(mfrow = c(3, 1))
    par(mar = c(4, 4, 2, 2))
    linestyles <- c("dashed", "solid")
    colors <- brewer.pal(3, "Dark2")[c(2, 1, 3)]
    treatments <- alldata$treatment[1:dynamicdata$npanels]

    plot(range(c(0, dynamicdata$times)), c(0, 1), type = "n", xlab = "week", ylab = expression(italic(A.~aurita)))
    tindex <- 1
    for(j in 1:dynamicdata$npanels){
        timeindex <- tindex:(tindex + dynamicdata$nt[j] - 1)
        for(i in 1:niter){
            lines(c(0, dynamicdata$times[timeindex]), c(0, simdata[i, timeindex, 1] / n[timeindex]), col = adjustcolor(colors[treatments[j]], 0.01), lty = linestyles[dynamicdata$depthcode[j]])
        }
        tindex <- tindex + dynamicdata$nt[j]
    }
    legend("topright", bty = "n", legend = c(expression(paste(1~m, ",", ~italic(C))), expression(paste(3~m, ",", ~italic(C))), expression(paste(1~m, ",", ~italic(A))), expression(paste(3~m, ",", ~italic(A))), expression(paste(1~m, ",", ~italic(O))), expression(paste(3~m, ",", ~italic(O)))), col = colors[c(2, 2, 1, 1, 3, 3)], lty = rep(linestyles, 3), pch = NA)
    
    plot(range(c(0, dynamicdata$times)), c(0, 1), type = "n", xlab = "week", ylab = "bare panel")
    tindex <- 1
    for(j in 1:dynamicdata$npanels){
        timeindex <- tindex:(tindex + dynamicdata$nt[j] - 1)
        for(i in 1:niter){
            lines(c(0, dynamicdata$times[timeindex]), c(1, simdata[i, timeindex, 2] / n[timeindex]), col = adjustcolor(colors[treatments[j]], 0.01), lty = linestyles[dynamicdata$depthcode[j]])
        }
        tindex <- tindex + dynamicdata$nt[j]
    }

    plot(range(c(0, dynamicdata$times)), c(0, 1), type = "n", xlab = "week", ylab = "potential competitors")
    tindex <- 1
    for(j in 1:dynamicdata$npanels){
        timeindex <- tindex:(tindex + dynamicdata$nt[j] - 1)
        for(i in 1:niter){
            lines(c(0, dynamicdata$times[timeindex]), c(0, simdata[i, timeindex, 3] / n[timeindex]), col = adjustcolor(colors[treatments[j]], 0.01), lty = linestyles[dynamicdata$depthcode[j]])
        }
        tindex <- tindex + dynamicdata$nt[j]
    }
    dev.off()
}

#plot efect of A. aurita on proportional population growth rate of potential competitors under each ODE model
#Arguments:
#odeoutputname, odeovergrowthoutputname, odeprotectionoutputname, odefacilitationoutputname, odegrowthfaciloutputname: bases of names for .RData files for base, overgrowth, protection, settlement facilitation and growth facilitation models respectively
#grid: how far out we go along each ilr coordinate when constructing a grid of opoints at which to calculate effects
#gridstep: size of step for grid of points
#rangelim: value at which we truncate colour scale when plotting effects
#fname: pdf file to which figure is written
#Value: pdf file fname, ternary plots of A. aurita effects for each model at each depth, with typical trajectories overlaid
plotallAureliaeffects <- function(odeoutputname, odeovergrowthoutputname, odeprotectionoutputname, odefacilitationoutputname, odegrowthfaciloutputname, grid, gridstep, rangelim, fname){
    pdf(file = fname, width = 7, height = 7)
    par(mfrow = c(5, 2))
    par(mar = c(1, 1, 0.5, 0.5))
    par(xpd = NA)

    print("overgrowth model")
    load(file = paste(odeovergrowthoutputname, ".RData", sep = ""))
    load(file = paste(odeovergrowthoutputname, "_typicalrho.RData", sep = ""))
    plotAureliaeffect(grid = grid, gridstep = gridstep, rangelim = rangelim, typicalrhoset = typicalrhoset, a0 = a0, a1 = a1, a2 = a2, deltab0 = deltab0, a1y1star = a1y1star, m0 = NULL, m1 = NULL, m2 = NULL, model = "overgrowth", labels = c("(a): overgrowth, 1 m", "(b): overgrowth, 3 m"))
    print("protection model")
    load(file = paste(odeprotectionoutputname, ".RData", sep = ""))
    load(file = paste(odeprotectionoutputname, "_typicalrho.RData", sep = ""))
    plotAureliaeffect(grid = grid, gridstep = gridstep, rangelim = rangelim, typicalrhoset = typicalrhoset, a0 = a0, a1 = a1, a2 = a2, deltab0 = deltab0, a1y1star = NULL, m0 = NULL, m1 = NULL, m2 = m2, model = "protection",  labels = c("(c): protection, 1 m", "(d): protection, 3 m"))
    print("basic model")
    load(file = paste(odeoutputname, ".RData", sep = ""))
    load(file = paste(odeoutputname, "_typicalrho.RData", sep = ""))
    plotAureliaeffect(grid = grid, gridstep = gridstep, rangelim = rangelim, typicalrhoset = typicalrhoset, a0 = a0, a1 = a1, a2 = a2, deltab0 = deltab0, a1y1star = NULL, m0 = NULL, m1 = NULL, m2 = NULL, model = "base", labels = c("(e): basic, 1 m", "(f): basic, 3 m"))
    print("settlement facilitation model")
    load(file = paste(odefacilitationoutputname, ".RData", sep = ""))
    load(file = paste(odefacilitationoutputname, "_typicalrho.RData", sep = ""))
    plotAureliaeffect(grid = grid, gridstep = gridstep, rangelim = rangelim, typicalrhoset = typicalrhoset, a0 = a0, a1 = a1, a2 = a2, deltab0 = deltab0, a1y1star = NULL, m0 = m0, m1 = NULL, m2 = NULL, model = "settlement", labels = c("(g): settlement facilitation, 1 m", "(h): settlement facilitation, 3 m"))
    print("growth facilitation model")
    load(file = paste(odegrowthfaciloutputname, ".RData", sep = ""))
    load(file = paste(odegrowthfaciloutputname, "_typicalrho.RData", sep = ""))
    plotAureliaeffect(grid = grid, gridstep = gridstep, rangelim = rangelim, typicalrhoset = typicalrhoset, a0 = a0, a1 = a1, a2 = a2, deltab0 = deltab0, a1y1star = NULL, m0 = NULL, m1 = m1, m2 = NULL, model = "growthfacil", labels = c("(i): growth facilitation, 1 m", "(j): growth facilitation, 3 m"))
    colorBarpoly(colvals = makecolors(x = seq(from = -rangelim, to = rangelim, length.out = 101), rangelim = rangelim), zlim = c(-rangelim, rangelim), parfig = c(0.4, 0.6, 0.4, 0.65))
    dev.off()
}


#for colour bars on ternary plots
#Arguments:
#colvals is a vector of colours,
#zlim is scale limits (vector of two elements)
#parfig and parmar are vectors for par(fig) and par(mar)
#tlabs (optional) specifies tick labels
#tpos (optional) specifies what these labels are
#title (optional) is a title for the colour bar
colorBarpoly <- function(colvals, zlim, parfig = c(0, 1, 0.835, 1), parmar = c(1, 4, 4, 5), tlabs = NULL, tpos = NULL, title = NULL){
    par(fig = parfig, new = TRUE)
    ntick <- 4
    if(is.null(tpos)){
        tpos <- pretty(zlim, n = ntick)
        tlat <- (tpos - min(tpos)) / (diff(zlim))
    } else tlat <- tpos
    if(is.null(tlabs)){
        tlabs <- round(tpos, digits = 0)
    }        
    par(mar = parmar)
    image(t(matrix(1:length(colvals), ncol = 1)), axes = FALSE, col = colvals)
    axis(2, at = tlat, tick = FALSE, labels = tlabs, line = -0.5, las = 1)
    if(!is.null(title)) text(0.5, 5 , title)
    par(fig = c(0, 1, 0, 1), new = FALSE)
}

#turn a composition into x and y coordinates that we can use with ordinary plotting commands, but that match up with plot.acomp
#Argument: z, a 3-part composition
#Value: list of x and y coordinates
comptoxy <- function(z){
    s60 <- sin(pi / 3)
    c60 <- cos(pi / 3)
    x <- z[, 2] + z[, 3] * c60
    y <- z[, 3] * s60
    return(list(x = x, y = y))
}

#build convenient grid for plotting polygons of effect of A. aurita on potential competitors on ternary plot
#Arguments:
#grid: extreme of grid points in ilr coordinates (we go from -grid to grid on each axis)
#gridstep: step size on each axis
#Value:
#a list containing variables midpoint (midpoints of polygons, as compositions), x and y (vertices of polygons, as x and y coordinates for plotting with base R)
getgrid <- function(grid,gridstep){
    xgrid <- seq(from = -grid, to = grid, by = gridstep)
    grd <- expand.grid(o1 = xgrid, o2 =xgrid)
    ngrd <- dim(grd)[1]
    polys <- vector(mode = "list", length = ngrd)
    for(i in 1:ngrd){
        xx <- c(grd[i, 1] - gridstep / 2, grd[i, 1] + gridstep / 2, grd[i, 1] + gridstep / 2, grd[i, 1] - gridstep / 2)
        yy <- c(grd[i, 2] - gridstep / 2, grd[i, 2] - gridstep / 2, grd[i, 2] + gridstep / 2, grd[i, 2] + gridstep / 2)
        corners <- comptoxy(ilrInv(cbind(xx, yy)))
        midpoint <- unclass(ilrInv(grd[i, ]))
        polys[[i]] <- list(grd = grd[i, ], x = corners$x, y = corners$y, midpoint = midpoint)
    }
    return(polys)
}

#ternary plot of effect of A. aurita on proportional population growth rate of potential competitors under a specified model (as colour of polygons), with predicted trajectories overlaid
#Arguments:
#grid: scalar, how far we go from origin in ilr coordinates on each axis (default 3)
#gridstep: size of grid cells on each ilr coordinate (default 0.5)
#rangelim: value of effect at which we truncate when colouring polygons (default 40)
#typicalrhoset: predictions from model, produced by typicalrho()
#a0: potential competitor settlement rate (iterations x depths)
#a1: potential competitor growth rate (iterations x depths)
#deltab0: polyp settlement rate * polyp area (iterations x depths)
#a1y1star: overgrowth rate (iterations x depths), optional
#m0: facilitation effect of polyps on potential comp settlement (iterations x depths), optional
#m1: growth facilitation effect of polyps on potential competitors (iterations x depths), optional
#m2: reduction in potential comp mortality due to polyps (iterations x depths), optional
#model: one of "base", "overgrowth", "settlement", "growthfacil", "protection"
#labels: vector of labels for panels representing effects at 1 m and at 3 m
plotAureliaeffect <- function(grid = 3, gridstep = 0.5, rangelim = 40, typicalrhoset, a0, a1, a2, deltab0, a1y1star = NULL, m0 = NULL, m1 = NULL, m2 = NULL, model = "overgrowth", labels){
    mycolors <- brewer.pal(3, "Dark2")[c(2, 1, 3)]
    mygrid <- getgrid(grid = grid, gridstep = gridstep)
    plot(acomp(c(1, 1, 1)), type = "n", labels = c(expression(italic(A.~aurita)), "bare panel", "potential competitors"))
    p <- lapply(mygrid, function(p){polygon(p$x, p$y, col = gridcolor(p, a0 = a0, a1 = a1, a2 = a2, deltab0 = deltab0, a1y1star = a1y1star, m0 = m0, m1 = m1, m2 = m2, rangelim = rangelim, model = model)$col[1], border = NA)})
    text(0.35, 0.75, labels[1], pos = 2)
    lines.acomp(typicalrhoset$rhoC1summary$metriccentre, col = mycolors[2], lty = "dashed", lwd = 1)
    lines.acomp(typicalrhoset$rhoO1summary$metriccentre, col = mycolors[3], lty = "dashed", lwd = 1)
    lines.acomp(typicalrhoset$rhoA1summary$metriccentre, col = mycolors[1], lty = "dashed", lwd = 1)

    plot(acomp(c(1, 1, 1)), type = "n", labels = c(expression(italic(A.~aurita)), "bare panel", "potential competitors"))
    p <- lapply(mygrid, function(p){polygon(p$x, p$y, col = gridcolor(p, a0 = a0, a1 = a1, a2 = a2, deltab0 = deltab0, a1y1star = a1y1star, m0 = m0, m1 = m1, m2 = m2, rangelim = rangelim, model = model)$col[2], border = NA)})
    text(0.35, 0.75, labels[2], pos = 2)
    lines.acomp(typicalrhoset$rhoC3summary$metriccentre, col = mycolors[2], lty = "solid", lwd = 1)
    lines.acomp(typicalrhoset$rhoO3summary$metriccentre, col = mycolors[3], lty = "solid", lwd = 1)
    lines.acomp(typicalrhoset$rhoA3summary$metriccentre, col = mycolors[1], lty = "solid", lwd = 1)

#    colorBarpoly(colvals = makecolors(x = seq(from = -rangelim, to = rangelim, length.out = 101), rangelim = rangelim), zlim = c(-rangelim, rangelim), parfig = c(0, 0.5, 0.7, 0.85))
}

#get colour required for plotting polygons by effect of A. aurita on potential competitors
#Arguments:
#p: one of the polygons returned by getgrid()
#a0: potential competitor settlement rate (iterations x depths)
#a1: potential competitor growth rate (iterations x depths)
#deltab0: polyp settlement rate * polyp area (iterations x depths)
#a1y1star: overgrowth rate (iterations x depths), optional
#m0: facilitation effect of polyps on potential comp settlement (iterations x depths), optional
#m1: growth facilitation effect of polyps on potential competitors (iterations x depths), optional
#m2: reduction in potential comp mortality due to polyps (iterations x depths), optional
#rangelim: value at which we truncate
#model: one of "base", "overgrowth", "settlement", "growthfacil", "protection"
#Value:
#col: colour for plotting (vector of values at 1 m and 3 m), based on dfdy1star
#dfdy1star: posterior mean effect of A. aurita on potential competitors (vector of values at 1 m an 3 m) under the specified model
gridcolor <- function(p, a0, a1, a2, deltab0, b1, b2, a1y1star= NULL, m0 = NULL, m1 = NULL, m2 = NULL, rangelim, model){
    dfdy1star <- numeric(2)
    for(k in 1:2){#depths
        Pi1 <- a0[, k] / deltab0[, k]
        Pi2  <- a1[, k] / deltab0[, k]
        Pi3 <- a2[, k] / deltab0[, k]
        Pi11 <- m0[, k] / deltab0[, k]
        Pi12 <- m1[, k] / deltab0[, k]
        Pi13 <- a1y1star[, k] / deltab0[, k]
        x <- p$midpoint[3] #potential competitors
        y1star <- p$midpoint[1] #A. aurita
        dfdy1star[k] <- switch(as.character(model),
                               "base" = mean(-Pi1 / x - Pi2),
                               "overgrowth" =  mean(-Pi1 / x - Pi2 + Pi13),
                               "settlement" = mean(-Pi1 / x - Pi2 + Pi11 / x * (1 - x - 2 * y1star)),
                               "growthfacil" = mean(-Pi1 / x - Pi2 + Pi12 * (1 - x) - 2 * Pi12 * y1star),
                               "protection" = mean(-Pi1 / x - Pi2 - m2 * Pi3 * exp(- m2 * y1star)),
                  stop(paste("model", model, "not recognized")))
    }
    col <- c(makecolors(x = dfdy1star[1], rangelim = rangelim), makecolors(x = dfdy1star[2], rangelim = rangelim))
    return(list(col = col, dfdy1star = dfdy1star))
}

#create colors from values of a quantitative variable for plotting on the ColorBrewer RdBu scale, centered on zero
#Arguments:
#vector x of numerical values
#scalar rangelim: values outside [-rangelim, rangelim] truncated to ends of color scale
#Value: values of a colorRampPalette function for the values of x (truncated as appropriate) 
makecolors <- function(x, rangelim){
    x[x < -rangelim] <- -rangelim
    x[x > rangelim] <- rangelim
    colindex <- cut(x, breaks = seq(from = -rangelim, to = rangelim, length.out = 101), include.lowest = TRUE)
    mycol <- colorRampPalette(brewer.pal(11, "RdBu"))(100)[colindex]
    return(mycol)
}

#generate rho from posterior mean parameters under overgrowth model (and simulate multinomial sampling with observed total counts)
#Arguments:
#dynamicdata: object produced by datafordynamicmodel()
#a0: potential competitor settlement rate (iterations x depths)
#a1: potential competitor growth rate (iterations x depths)
#deltab0: polyp settlement rate * polyp area (iterations x depths)
#b1: polyp budding rate (iterations x depths)
#b2: polyp death rate (iterations x depths)
#kA: proportion A. aurita not removed in A treatment (vector: iterations)
#kO: proportion potential competitors not removed in O treatment (vector: iterations)
#a1y1star: overgrowth rate of polyps by potential competitors (iterations x depths)
#niter: number of simulated data sets to write to Stan input files
#fname: base for Stan input file name for each of the simulated data sets
#Value: list containing:
#rhofrommean: expected composition at each time, evaluated at posterior means of parameters (stacked observations x 3, same arrangement as rho from Stan output)
#simcounts: simulated multinomial sample from rhofrommean, with observed total count for each observation
simfrommeanparameters <- function(dynamicdata, a0, a1, a2, deltab0, b1, b2, kA, kO, a1y1star, niter, fname){
    meana0 <- colMeans(a0) #posterior mean parameters
    meana1 <- colMeans(a1)
    meana2 <- colMeans(a2)
    meandeltab0 <- colMeans(deltab0)
    meanb1 <- colMeans(b1)
    meanb2 <- colMeans(b2)
    meankA <- mean(kA)
    meankO <- mean(kO)
    meana1y1star <- colMeans(a1y1star)
    startind <- c(0, cumsum(dynamicdata$nt)) + 1
    endind <- cumsum(dynamicdata$nt)
    rhofrommean <- array(NA, dim = c(dynamicdata$sumnt, 3))

    for(i in 1:dynamicdata$npanels){
        rhofrommean[startind[i]:endind[i], ] <- buildrho(a0 = meana0[dynamicdata$depthcode[i]], a1 = meana1[dynamicdata$depthcode[i]], a2 = meana2[dynamicdata$depthcode[i]], deltab0 = meandeltab0[dynamicdata$depthcode[i]], b1 = meanb1[dynamicdata$depthcode[i]], b2 = meanb2[dynamicdata$depthcode[i]], kA = meankA, kO = meankO, m0 = 0, m2 = 0, psi = 0, a1y1star = meana1y1star[dynamicdata$depthcode[i]], m1 = 0, treatmentapplied = dynamicdata$treatmentapplied[startind[i]:endind[i]], timeblocks = dynamicdata$tb$timeblocks[i, , ], ntimeblocks = dynamicdata$tb$ntimeblocks[i], ntimesinblock = dynamicdata$tb$ntimesinblock[i, ], controltruncate = FALSE)
    }
    totalcount <- rowSums(dynamicdata$counts)
    n <- totalcount
    ntaxa <- dynamicdata$ntaxa
    npanels <- dynamicdata$npanels
    nt <- dynamicdata$nt
    sumnt <- dynamicdata$sumnt
    times <- dynamicdata$times
    ntimeblocks <- dynamicdata$tb$ntimeblocks
    ntimesinblock <- dynamicdata$tb$ntimesinblock
    timeblocks <- dynamicdata$tb$timeblocks
    maxtimeblocks <- dynamicdata$tb$maxtimeblocks
    maxtimes <- dynamicdata$tb$maxtimes
    treatmentapplied <- dynamicdata$treatmentapplied
    depthcode <- dynamicdata$depthcode
    for(j in 1:niter){
        simcounts <- array(NA, dim = c(dynamicdata$sumnt, 3))
        for(i in 1:dynamicdata$sumnt){
            simcounts[i, ] <- rmultinom(n = 1, size = totalcount[i], prob = rhofrommean[i, ])
        }
        counts <- simcounts
        ddata <- c("counts", "ntaxa", "npanels", "nt", "sumnt", "times", "ntimeblocks", "ntimesinblock", "timeblocks", "maxtimeblocks", "maxtimes", "treatmentapplied", "depthcode")
        stan_rdump(list = ddata, file = paste(fname, "_", j, ".txt", sep = ""), quiet = FALSE, append = FALSE)
    }
    return(rhofrommean)
}

#Arguments:
#fname: base of name for simulated ode data files
#outname: name of .RData file to which results will be written
#Samples from which mean is the true value used in simulations:
#a0: potential competitor settlement rate (iterations x depths)
#a1: potential competitor growth rate (iterations x depths)
#deltab0: polyp settlement rate * polyp area (iterations x depths)
#b1: polyp budding rate (iterations x depths)
#b2: polyp death rate (iterations x depths)
#kA: proportion A. aurita not removed in A treatment (vector: iterations)
#kO: proportion potential competitors not removed in O treatment (vector: iterations)
#a1y1star: overgrowth rate of polyps by potential competitors (iterations x depths)
#Value: write to a .RData file in same directory as fname: true values, densities, and whether or not in HPD interval
#Note: only implemented for overgrowth model
readrawsimdata <- function(fname, outname, a0, a1, a2, deltab0, b1, b2, a1y1star, kA, kO){
    meana0 <- colMeans(a0) #posterior mean parameters: used to simulate
    meana1 <- colMeans(a1)
    meana2 <- colMeans(a2)
    meandeltab0 <- colMeans(deltab0)
    meanb1 <- colMeans(b1)
    meanb2 <- colMeans(b2)
    meankA <- mean(kA)
    meankO <- mean(kO)
    meana1y1star <- colMeans(a1y1star)

    simpath <- sub("/[^/]+$", "", simodename) #strip everything after last / to get just the path to the simulated data files
    simfiles <- list.files(path = simpath, pattern = ".txt$")
    densities <- vector("list",length = length(simfiles))
    i <- simfiles[1]
    ipattern <- paste(i, "[[:digit:]].csv", sep = "")
    csvfiles <- list.files(path = simpath, pattern = ipattern)
    truetheta <- c(meana0, meana1, meana2, meandeltab0, meanb1, meanb2, meana1y1star, meankA, meankO)
    inHPDs <- array(dim = c(length(simfiles), length(truetheta)))
    colnames(inHPDs) <- c("a0.1m", "a0.3m", "a1.1m", "a1.3m", "a2.1m", "a2.3m", "deltab0.1m", "deltab0.3m", "b1.1m", "b1.3m", "b2.1m", "b2.3m", "a1y1star.1m", "a1y1star.3m", "kA", "kO")
    for(j in 1:length(simfiles)){
        i <- simfiles[j]
        print(i)
        ipattern <- paste(i, "[[:digit:]].csv", sep = "")
        csvfiles <- list.files(path = simpath, pattern = ipattern)
        print(csvfiles)
        idens <- getsimdens(csvfiles = paste(simpath, csvfiles, sep = "/"), truetheta = truetheta)
        inHPDs[j, ] <- idens$inHPDs
        densities[[j]] <- idens$densities
    }
    save(truetheta, densities, inHPDs, file = outname)
}

#read data from a set of .csv Stan output files generated from simulated data, extract posterior densities of parameters from these, and determine whether true value is in 95% HPD interval for each parameter
#Arguments:
#csvfiles: character vector of file names
#truetheta is vector of true parameter values
#Value:
#returns a list containing:
#inHPDs: 1 x 16 array, with value TRUE if the corresponding true parameter is in the 95% HPD interval for the estimate. Ordered as in truetheta
#densities: list of kernel density estimates for each parameter in order as above: each the output of density()
#Note: the order of elements in truetheta, inHPDs, densities has to be maintained by hand! Only implemented for overgrowth model
getsimdens <- function(csvfiles, truetheta){
    gc()
    standata <- read_stan_csv(csvfiles = csvfiles)
    a0 <- extract(standata, pars = "a0")[[1]]
    a1 <- extract(standata, pars = "a1")[[1]]
    a2 <- extract(standata, pars = "a2")[[1]]
    deltab0 <- extract(standata, pars = "deltab0")[[1]]
    b1 <- extract(standata, pars = "b1")[[1]]
    b2 <- extract(standata, pars = "b2")[[1]]
    a1y1star <- extract(standata, pars = "a1y1star")[[1]]
    kA <- extract(standata, pars = "kA")[[1]]
    kO <- extract(standata, pars = "kO")[[1]]    
    rm(standata)
    gc()
    densities <- vector("list", length = length(truetheta))
    densities[[1]] <- density(a0[, 1], bw = "SJ")
    densities[[2]] <- density(a0[, 2], bw = "SJ")
    densities[[3]] <- density(a1[, 1], bw = "SJ")
    densities[[4]] <- density(a1[, 2], bw = "SJ")
    densities[[5]] <- density(a2[, 1], bw = "SJ")
    densities[[6]] <- density(a2[, 2], bw = "SJ")
    densities[[7]] <- density(deltab0[, 1], bw = "SJ")
    densities[[8]] <- density(deltab0[, 2], bw = "SJ")
    densities[[9]] <- density(b1[, 1], bw = "SJ")
    densities[[10]] <- density(b1[, 2], bw = "SJ")
    densities[[11]] <- density(b2[, 1], bw = "SJ")
    densities[[12]] <- density(b2[, 2], bw = "SJ")
    densities[[13]] <- density(a1y1star[, 1], bw = "SJ")
    densities[[14]] <- density(a1y1star[, 2], bw = "SJ")
    densities[[15]] <- density(kA, bw = "SJ")
    densities[[16]] <- density(kO, bw = "SJ")
    
    inHPDs <- array(dim=c(1, length(truetheta)))
    inHPDs[1] <- inHPD(a0[, 1], truetheta[1])
    inHPDs[2] <- inHPD(a0[, 2], truetheta[2])
    inHPDs[3] <- inHPD(a1[, 1], truetheta[3])
    inHPDs[4] <- inHPD(a1[, 2], truetheta[4])
    inHPDs[5] <- inHPD(a2[, 1], truetheta[5])
    inHPDs[6] <- inHPD(a2[, 2], truetheta[6])
    inHPDs[7] <- inHPD(deltab0[, 1], truetheta[7])
    inHPDs[8] <- inHPD(deltab0[, 2], truetheta[8])
    inHPDs[9] <- inHPD(b1[, 1], truetheta[9])
    inHPDs[10] <- inHPD(b1[, 2], truetheta[10])
    inHPDs[11] <- inHPD(b2[, 1], truetheta[11])
    inHPDs[12] <- inHPD(b2[, 2], truetheta[12])
    inHPDs[13] <- inHPD(a1y1star[, 1], truetheta[13])
    inHPDs[14] <- inHPD(a1y1star[, 2], truetheta[14])
    inHPDs[15] <- inHPD(kA, truetheta[15])
    inHPDs[16] <- inHPD(kO, truetheta[16])
    
    return(list(inHPDs = inHPDs, densities = densities))
}

#is true value in 95% HPD interval?
#Arguments:
#theta is a vector of estimates of a parameter (iterations x 1)
#truetheta is true value
#Value: is truetheta in HPD intervals (logical)?
inHPD <- function(theta,truetheta){
    n <- length(theta)
    hpdi <- HPDinterval(as.mcmc(array(theta, dim = c(n, 1))))
    return(truetheta >= hpdi[1] & truetheta <= hpdi[2])
}

#plot collated output from simulations
#Arguments:
#truetheta is vector of true parameter values
#densities is a list of lists (first dimension replicates, second dimension parameters): each element is a kernel density estimate
#priorhalfnorms: array (one row per parameter) of prior sd and sign for parameters with half-normal priors
#priorbetaparms: array of parameters for beta priors for each of the remaining parameters, in same order as truetheta, one row per parameter
#inHPDs: array, simulations x parameters, logical: was true value of parameter in HPD interval from estimation?
#fname: name of pdf file to produce
#Value: plot posterior and prior densities and true values of each parameter to pdf file fname
#Note: x axis labels, limits, etc must all be maintained by hand. Only currently for overgrowth model
plotsimdens <- function(truetheta, densities, priorhalfnorms, priorbetaparms, inHPDs, fname){
    pdf(file = fname, width = 7, height = 7)
    par(mfrow=c(4, 4))
    par(mar = c(4, 4, 1, 1))
    nparm <- length(densities[[1]])
    ndens <- length(densities)
    xlabs <- c(expression(paste(italic(a)[0]:~1~m)),
               expression(paste(italic(a)[0]:~3~m)),
               expression(paste(italic(a)[1]:~1~m)),
               expression(paste(italic(a)[1]:~3~m)),
               expression(paste(italic(a)[2]:~1~m)),
               expression(paste(italic(a)[2]:~3~m)),
               expression(paste(italic(delta*b)[0]:~1~m)),
               expression(paste(italic(delta*b)[0]:~3~m)),
               expression(paste(italic(b)[1]:~1~m)),
               expression(paste(italic(b)[1]:~3~m)),
               expression(paste(italic(b)[2]:~1~m)),
               expression(paste(italic(b)[2]:~3~m)),
               expression(paste(italic(a)[1*","~italic(y)[1]*"*"]:~1~m)),
               expression(paste(italic(a)[1*","~italic(y)[1]*"*"]:~3~m)),
               expression(paste(italic(k)[A])),
               expression(paste(italic(k)[O]))
               )
    scores <- colSums(inHPDs)
    iter <- dim(inHPDs)[1]
    for(i in 1:(nparm - 2)){
        mylabel <- paste("(", letters[i], "): ", scores[i], "/", iter, sep = "")
        plotparmhalfnorm(densities = densities, i = i, ndens = ndens, truetheta = truetheta, xlabs = xlabs, label = mylabel, priorhalfnorm = priorhalfnorms[i, ])
    }
    for(i in (nparm - 1):nparm){#last two: kA and kO must be between zero and one
        mylabel <- paste("(", letters[i], "): ", scores[i], "/", iter, sep = "")
        plotparmbeta(densities = densities, i = i, ndens = ndens, truetheta = truetheta, xlabs = xlabs, label = mylabel, priorparms = priorbetaparms[(i - nparm + 2), ])
    }
    dev.off()
}

#plot density estimates for a single parameter from multiple simulations, that has to be between zero and one (and has a beta prior)
#Arguments:
#densities is a list of lists (first dimension replicates, second dimension parameters): each element is a kernel density estimate
#i is the element we want to plot (second dimension of densities)
#ndens is number of simulations
#truetheta is vector of true parameter values
#xlabs is vector of x axis labels (parameter names)
#label: if not NULL (default), letter for panel label
#priorparms: vector of parameters for prior beta distribution
#Value: plot kernel density estimates from simulations, line at true value
plotparmbeta <- function(densities, i, ndens, truetheta, xlabs, label = NULL, priorparms){
    mycols <- brewer.pal(3, "Dark2")
    yrange <- 0
    for(j in 1:ndens){
        yrange <- range(c(yrange, densities[[j]][[i]]$y))
    }

    plot(densities[[1]][[i]], xlim = c(0, 1), ylim = yrange, col = adjustcolor("black", 0.1), main="", xlab=xlabs[i])
    for(j in 1:ndens){
      lines(densities[[j]][[i]], col = adjustcolor("black", 0.1))      
    }
    x <- seq(from = 0, to = 1, length.out = 100)
    priordens <- dbeta(x, shape1 = priorparms[1], shape2 = priorparms[2])
    lines(x, priordens, col = mycols[2], lty = "dashed")
    abline(v=truetheta[i], col = mycols[1], lwd = 2)
    if(!is.null(label)){
        axlims <- par("usr")
        text(axlims[1] + 0.3 * (axlims[2] - axlims[1]), axlims[4] - 0.1 * (axlims[4] - axlims[3]), label)
    }
}

#plot density estimates for a single parameter from multiple simulations, that has a half-normal prior
#Arguments:
#densities is a list of lists (first dimension replicates, second dimension parameters): each element is a kernel density estimate
#i is the element we want to plot (second dimension of densities)
#ndens is number of simulations
#truetheta is vector of true parameter values
#xlabs is vector of x axis labels (parameter names)
#label: if not NULL (default), letter for panel label
#priorhalfnorm: sd and sign for half-normal prior
#xlim (default NULL): range for x axis (if null, chosen from data)
#Value: plot kernel density estimates from simulations, line at true value
plotparmhalfnorm <- function(densities, i, ndens, truetheta, xlabs, label = NULL, priorhalfnorm, xlim = NULL){
    mycols <- brewer.pal(3, "Dark2")
    xrange <- 0
    yrange <- 0
    for(j in 1:ndens){
        xrange <- range(c(xrange, truetheta[i], densities[[j]][[i]]$x))
        yrange <- range(c(yrange, densities[[j]][[i]]$y))
    }

    if(!is.null(xlim)){
        xrange <- xlim
    }

    if(priorhalfnorm[2] > 0){
        xrange[1] <- 0
    } else {
        xrange[2] <- 0
    }
    
    plot(densities[[1]][[i]], xlim = xrange, ylim = yrange, col = adjustcolor("black", 0.1), main="", xlab=xlabs[i])
    for(j in 1:ndens){
      lines(densities[[j]][[i]], col = adjustcolor("black", 0.1))      
    }
    x <- seq(from = xrange[1], to = xrange[2], length.out = 100)
    priordens <- 2 * dnorm(x, mean = 0, sd = priorhalfnorm[1]) #half-normal
    lines(x, priordens, col = mycols[2], lty = "dashed")
    abline(v=truetheta[i], col = mycols[1], lwd = 2)
    if(!is.null(label)){
        axlims <- par("usr")
        text(axlims[1] + 0.3 * (axlims[2] - axlims[1]), axlims[4] - 0.1 * (axlims[4] - axlims[3]), label)
    }
}
```
