## Supplementary material for "Negative and positive interspecific interactions involving jellyfish polyps in marine sessile communities": Data and code: aureliamanova.stan

```
functions {
/** inverse ilr transformation of a vector x, using the inverse of the transpose of the V matrix of the ilr (tVinv)
*/
  vector ilrinv(matrix tVinv, vector x, int ntaxa) {
         vector[ntaxa] y;
         y = softmax(tVinv * x);
         return y;
	}
}

data {
  int<lower = 0> ntaxa; //number of taxa
  int<lower = 0> npanels; //number of panels
  int<lower = 0> nblocks; //number of blocks
  matrix[ntaxa, ntaxa - 1] tVinv; //back-transformation matrix for ilr transformation
  int blockindex[npanels]; //block index
  int counts[npanels, ntaxa]; //observed counts
  vector[npanels] d1; //dummy variable for depth contrast
  vector[npanels] d2; //dummy variable for O vs A scrape contrast
  vector[npanels] d3; //dummy variable for O and A vs C scrape contrast
  vector[npanels] d4; //dummy variable for first interaction contrast
  vector[npanels] d5; //dummy variable for second interaction contrast
}

transformed data {
  int<lower = 1> s;
  s = ntaxa - 1;
}

parameters {
  vector[s] mu; //intercept 
  vector[s] alpha; //depth contrast (basis vector d1)
  vector[s] beta1; //scrape contrast: O vs A (basis vector d2)
  vector[s] beta2; //scrape contrast: C vs O and A (basis vector d3)
  vector[s] gamma1; //interaction contrast built from kronecker product of d1, d2
  vector[s] gamma2; //interaction contrast built from kronecker product of d1, d3
  vector[s] zpanel[npanels]; //standardized panel effect
  vector[s] zblock[nblocks]; //standardized block effect
  cholesky_factor_corr[s] LOmegaSigma; //Cholesky factor of prior correlation for panels
  vector<lower=0>[s] tauSigma; //prior scale on covariances for panels
  cholesky_factor_corr[s] LOmegaZeta; //Cholesky factor of prior correlation for blocks
  vector<lower=0>[s] tauZeta; //prior scale on covariances for blocks
}

transformed parameters {
  cholesky_factor_cov[s] LSigma; //Cholesky factor of panel effect covariance matrix
  cholesky_factor_cov[s] LZeta; //Cholesky factor of block effect covariance matrix
  vector[s] x[npanels]; //predicted logratio coordinates
  vector[ntaxa] rho[npanels]; //predicted relative abundances

  LSigma = diag_pre_multiply(tauSigma, LOmegaSigma);
  LZeta = diag_pre_multiply(tauZeta, LOmegaZeta);
  for(i in 1:npanels){
    x[i] = mu + alpha * d1[i] + beta1 * d2[i] + beta2 * d3[i] + gamma1 * d4[i] + gamma2 * d5[i] + LSigma * zpanel[i] + LZeta * zblock[blockindex[i]];
    rho[i] = ilrinv(tVinv, x[i], ntaxa);
  } 

}

model {
  vector[s] mudepth = [2.8, -2.6, -0.1, 0.1]'; //prior mean vector for depth effect
  matrix[s, s] Sigmadepth = [[12.7, 3.2e-4, 5.5e-5, 8.5e-5],
			     [3.2e-4, 7.1, 4.1e-5, 6.4e-5],
			     [5.5e-5, 4.1e-5, 2.1e-1, 1.1e-5],
			     [8.5e-5, 6.4e-5, 1.1e-5, 5.0e-1]]; //prior covariance matrix for depth effect
  for(i in 1:npanels) {
    counts[i] ~ multinomial(rho[i]); // observation model
    zpanel[i] ~ normal(0, 1);
  }
  for(j in 1:nblocks) {
    zblock[j] ~ normal(0, 1);
  }
  tauSigma ~ cauchy(0, 2.5);
  LOmegaSigma ~ lkj_corr_cholesky(2);
  tauZeta ~ cauchy(0, 2.5);
  LOmegaZeta ~ lkj_corr_cholesky(2);
  mu ~ normal(0, 2);
  alpha ~ multi_normal(mudepth, Sigmadepth);
  beta1 ~ normal(0, 1);
  beta2 ~ normal(0, 1);
  gamma1 ~ normal(0, 1);
  gamma2 ~ normal(0, 1);
}

generated quantities{
  vector [npanels] log_lik;
  for (j in 1:npanels)
    log_lik[j] = multinomial_lpmf(counts[j] | rho[j]); //log likelihood for WAIC
}
```
