## Supplementary material for "Negative and positive interspecific interactions involving jellyfish polyps in marine sessile communities": Data and code: aureliaode.stan

```
functions {
  //differential equations: first component is rate of change of potential competitors, second component is rate of change of A. aurita
  vector dz_dt(real t, //time
		vector z, //state {proportions filled by potential competitors and polyps on substrate}
		real a0, //settlement by potential competitors
		real a1, //growth of potential competitors
		real a2, //death of potential competitors
		real deltab0, //area per polyp * settlement of polyps 
		real b1, //growth of polyps
		real b2) //death of polyps
  {
    vector[2] dzdt;
    real x = z[1]; //proportion filled by potential competitors
    real y1star = z[2]; //proportion filled by polyps
    real freespace = 1 - x - y1star;
    dzdt[1] = a0 * freespace + a1 * x * freespace + a2 * x;
    dzdt[2] = deltab0 * freespace + b1 * y1star * freespace +b2 * y1star;
    return dzdt;
  }

  //remove fraction 1 - kA of A. aurita from vector z (will only work with ntaxa = 3, A. aurita second component of z)
  vector Atreatment(vector z, real kA) {
    vector[2] newz;
    newz = z;
    newz[2] = kA * newz[2];
    return newz;
  }

  //remove proportion 1 - kO of potential competitors from vector z (will only work with ntaxa = 3, potential competitors first component of z)
  vector Otreatment(vector z, real kO) {
    vector[2] newz;
    newz = z;
    newz[1] = kO * newz[1];
    return newz;
  }
}

data {
  int <lower = 1> ntaxa; //number of taxa
  int <lower = 1> npanels; //number of panels
  int <lower = 1> sumnt; //total number of observations
  int <lower = 1> maxtimeblocks; //maximum time blocks per panel
  int <lower = 1> maxtimes; //max possible number of observation times in time block
  int nt[npanels]; //number of observations on each panel
  int counts[sumnt, ntaxa]; //stacked multinomial counts: columns are taxa. Sorted by panel, and by time and then pre/post within each panel
  int ntimeblocks[npanels]; //number of time blocks on each panel
  int ntimesinblock[npanels, maxtimeblocks]; //number of times in each time block on each panel
  real timeblocks[npanels, maxtimeblocks, maxtimes]; //observation times in each time block on each panel
  int treatmentapplied[sumnt]; //treatment applied to each observation (0 for no treatment)
  int depthcode[npanels]; //what depth was a panel at? Code 1 for 1 m, 2 for 3 m
}

parameters {
  real <lower = 0> a0[2]; //vector of 2 values: each parameter at 2 different depths
  real <lower = 0> a1[2];
  real <upper = 0> a2[2];
  real <lower = 0> deltab0[2];
  real <lower = 0> b1[2];
  real <upper = 0> b2[2];
  real <lower = 0, upper = 1> kA;
  real <lower = 0, upper = 1> kO;
}

transformed parameters {
  vector[ntaxa] rho[sumnt];
  real t0;
  vector[2] zinit; //this will only work with ntaxa equal to 3
  vector[ntaxa - 1] z[sumnt]; //latent states
  { //new block inside which we can declare integer tindex
      int tindex = 1;
      for(j in 1:npanels) {
	zinit = [0.0, 0.0]';
	t0 = 0.0;
	for(k in 1:ntimeblocks[j]) {
	  z[tindex:(ntimesinblock[j, k] + tindex - 1)] = ode_rk45(dz_dt, zinit, t0, timeblocks[j, k, 1:ntimesinblock[j, k]], a0[depthcode[j]], a1[depthcode[j]], a2[depthcode[j]], deltab0[depthcode[j]], b1[depthcode[j]], b2[depthcode[j]]);
	  t0 = timeblocks[j, k, ntimesinblock[j, k]];
	  zinit = z[ntimesinblock[j, k] + tindex - 1];
	  if(treatmentapplied[tindex + ntimesinblock[j, k]] == 2) {//A treatment
	    zinit = Atreatment(zinit, kA);
	  } else if(treatmentapplied[tindex + ntimesinblock[j, k]] == 3) {//O treatment
	    zinit = Otreatment(zinit, kO);
	  }
	  if(treatmentapplied[tindex + ntimesinblock[j, k]] != 0) {//Post observation: note that if we have a control panel, there is no post observation. Doesn't overflow or cause problems with current data because next observation after this always a pre for next panel, and last panel is not a control
	    z[ntimesinblock[j, k] + tindex] = zinit; 
	    tindex = tindex + 1;
	  }
	  tindex = tindex + ntimesinblock[j, k];
	}
      }
  }
  for(j in 1:sumnt) {
    rho[j, 3] = z[j, 1];
    rho[j, 1] = z[j, 2];
    rho[j, 2] = 1.0 - z[j, 1] - z[j, 2];
  }
}

model {
  for(j in 1:sumnt) {
    counts[j] ~ multinomial(rho[j]); //observation model
  }

  a0 ~ normal(0, 0.05);
  a1 ~ normal(0, 1);
  a2 ~ normal(0, 0.05);
  deltab0 ~ normal(0, 0.02);
  b1 ~ normal(0, 0.2);
  b2 ~ normal(0, 0.05);
  kA ~ beta(2, 2);
  kO ~ beta(2, 2);
}

generated quantities {
  vector[sumnt] log_lik;
  for(j in 1:sumnt) {
    log_lik[j] = multinomial_lpmf(counts[j] | rho[j]); //log likelihood for loo
  }
}
```
