## Supplementary material for "Negative and positive interspecific interactions involving jellyfish polyps in marine sessile communities": Data and code: aureliaode_overgrowth_sbc.stan

```
functions {
  //differential equations: first component is rate of change of potential competitors, second component is rate of change of A. aurita
  vector dz_dt(real t, //time
	       vector z, //state {proportions filled by potential competitors and polyps on substrate}
	       real a0, //settlement by potential competitors
	       real a1, //growth of potential competitors
	       real a1y1star, //overgrowth of polyps by potential competitors
	       real a2, //death of potential competitors
	       real deltab0, //area per polyp * settlement of polyps 
	       real b1, //growth of polyps
	       real b2) //death of polyps
  {
    vector[2] dzdt;
    real x = z[1]; //proportion filled by potential competitors
    real y1star = z[2]; //proportion filled by polyps
    real freespace = 1 - x - y1star;
    dzdt[1] = a0 * freespace + a1 * x * freespace + a1y1star * x * y1star + a2 * x;
    dzdt[2] = deltab0 * freespace + b1 * y1star * freespace - a1y1star * x * y1star + b2 * y1star;
    return dzdt;
  }

  //remove fraction 1 - kA of A. aurita from vector z (will only work with ntaxa = 3, A. aurita second component of z)
  vector Atreatment(vector z, real kA) {
    vector[2] newz;
    newz = z;
    newz[2] = kA * newz[2];
    return newz;
  }

  //remove proportion 1 - kO of potential competitors from vector z (will only work with ntaxa = 3, potential competitors first component of z)
  vector Otreatment(vector z, real kO) {
    vector[2] newz;
    newz = z;
    newz[1] = kO * newz[1];
    return newz;
  }

  //generate normal random numbers, truncated below at zero. mu and sigma are mean and standard deviation of the untruncated normal. Based on section 18.10 in stan user's guide, version 2.27
  real posnormal_rng(real mu, real sigma) {
    real p_lb = normal_cdf(0, mu, sigma);
    real u = uniform_rng(p_lb, 1.0);
    real y = mu + sigma * inv_Phi(u);
    return y;
  }

}

data {
  int <lower = 1> ntaxa; //number of taxa
  int <lower = 1> npanels; //number of panels
  int <lower = 1> sumnt; //total number of observations
  int <lower = 1> maxtimeblocks; //maximum time blocks per panel
  int <lower = 1> maxtimes; //max possible number of observation times in time block
  int nt[npanels]; //number of observations on each panel
  int n[sumnt]; //number of points counted in each stacked multinomial observation
  //  int counts[sumnt, ntaxa]; //stacked multinomial counts: columns are taxa. Sorted by panel, and by time and then pre/post within each panel
  int ntimeblocks[npanels]; //number of time blocks on each panel
  int ntimesinblock[npanels, maxtimeblocks]; //number of times in each time block on each panel
  real timeblocks[npanels, maxtimeblocks, maxtimes]; //observation times in each time block on each panel
  int treatmentapplied[sumnt]; //treatment applied to each observation (0 for no treatment)
  int depthcode[npanels]; //what depth was a panel at? Code 1 for 1 m, 2 for 3 m
}

transformed data {
  real <lower = 0> a0_[2]; //vector of 2 values: each parameter at 2 different depths
  real <lower = 0> a1_[2];
  real <lower = 0> a1y1star_[2];
  real <upper = 0> a2_[2];
  real <lower = 0> deltab0_[2];
  real <lower = 0> b1_[2];
  real <upper = 0> b2_[2];
  real <lower = 0, upper = 1> kA_;
  real <lower = 0, upper = 1> kO_;
  vector[ntaxa] rho_[sumnt];
  real t0_;
  vector[2] zinit_; //this will only work with ntaxa equal to 3
  vector[ntaxa - 1] z_[sumnt]; //latent states
  int counts[sumnt, ntaxa]; //prior predicted stacked multinomial counts
  //sample parameter values from priors
  for(i in 1:2){
    a0_[i] = posnormal_rng(0, 0.05);
    a1_[i] = posnormal_rng(0, 1);
    a1y1star_[i] = posnormal_rng(0, 1);
    a2_[i] = -posnormal_rng(0, 0.05); //a2 must be negative
    deltab0_[i] = posnormal_rng(0, 0.02);
    b1_[i] = posnormal_rng(0, 0.2);
    b2_[i] = -posnormal_rng(0, 0.05); //b2 must be negative
  }
  kA_ = beta_rng(2, 2);
  kO_ = beta_rng(2, 2);

  { //new block inside which we can declare integer tindex
      int tindex = 1;
      for(j in 1:npanels) {
	zinit_ = [0.0, 0.0]';
	t0_ = 0.0;
	for(k in 1:ntimeblocks[j]) {
	  z_[tindex:(ntimesinblock[j, k] + tindex - 1)] = ode_rk45(dz_dt, zinit_, t0_, timeblocks[j, k, 1:ntimesinblock[j, k]], a0_[depthcode[j]], a1_[depthcode[j]], a1y1star_[depthcode[j]], a2_[depthcode[j]], deltab0_[depthcode[j]], b1_[depthcode[j]], b2_[depthcode[j]]);
	  t0_ = timeblocks[j, k, ntimesinblock[j, k]];
	  zinit_ = z_[ntimesinblock[j, k] + tindex - 1];
	  if(treatmentapplied[tindex + ntimesinblock[j, k]] == 2) {//A treatment
	    zinit_ = Atreatment(zinit_, kA_);
	  } else if(treatmentapplied[tindex + ntimesinblock[j, k]] == 3) {//O treatment
	    zinit_ = Otreatment(zinit_, kO_);
	  }
	  if(treatmentapplied[tindex + ntimesinblock[j, k]] != 0) {//Post observation: note that if we have a control panel, there is no post observation. Doesn't overflow or cause problems with current data because next observation after this always a pre for next panel, and last panel is not a control
	    z_[ntimesinblock[j, k] + tindex] = zinit_; 
	    tindex = tindex + 1;
	  }
	  tindex = tindex + ntimesinblock[j, k];
	}
      }
  }
  for(j in 1:sumnt) {
    rho_[j, 3] = z_[j, 1];
    rho_[j, 1] = z_[j, 2];
    rho_[j, 2] = 1.0 - z_[j, 1] - z_[j, 2];
    counts[j] = multinomial_rng(rho_[j], n[j]); // realizations from prior predictive distribution
  }
}

parameters {
  real <lower = 0> a0[2]; //vector of 2 values: each parameter at 2 different depths
  real <lower = 0> a1[2];
  real <lower = 0> a1y1star[2];
  real <upper = 0> a2[2];
  real <lower = 0> deltab0[2];
  real <lower = 0> b1[2];
  real <upper = 0> b2[2];
  real <lower = 0, upper = 1> kA;
  real <lower = 0, upper = 1> kO;
}

transformed parameters {
  vector[ntaxa] rho[sumnt];
  real t0;
  vector[2] zinit; //this will only work with ntaxa equal to 3
  vector[ntaxa - 1] z[sumnt]; //latent states
  { //new block inside which we can declare integer tindex
      int tindex = 1;
      for(j in 1:npanels) {
	zinit = [0.0, 0.0]';
	t0 = 0.0;
	for(k in 1:ntimeblocks[j]) {
	  z[tindex:(ntimesinblock[j, k] + tindex - 1)] = ode_rk45(dz_dt, zinit, t0, timeblocks[j, k, 1:ntimesinblock[j, k]], a0[depthcode[j]], a1[depthcode[j]], a1y1star[depthcode[j]], a2[depthcode[j]], deltab0[depthcode[j]], b1[depthcode[j]], b2[depthcode[j]]);
	  t0 = timeblocks[j, k, ntimesinblock[j, k]];
	  zinit = z[ntimesinblock[j, k] + tindex - 1];
	  if(treatmentapplied[tindex + ntimesinblock[j, k]] == 2) {//A treatment
	    zinit = Atreatment(zinit, kA);
	  } else if(treatmentapplied[tindex + ntimesinblock[j, k]] == 3) {//O treatment
	    zinit = Otreatment(zinit, kO);
	  }
	  if(treatmentapplied[tindex + ntimesinblock[j, k]] != 0) {//Post observation: note that if we have a control panel, there is no post observation. Doesn't overflow or cause problems with current data because next observation after this always a pre for next panel, and last panel is not a control
	    z[ntimesinblock[j, k] + tindex] = zinit; 
	    tindex = tindex + 1;
	  }
	  tindex = tindex + ntimesinblock[j, k];
	}
      }
  }
  for(j in 1:sumnt) {
    rho[j, 3] = z[j, 1];
    rho[j, 1] = z[j, 2];
    rho[j, 2] = 1.0 - z[j, 1] - z[j, 2];

  }
}

model {
  for(j in 1:sumnt) {
    counts[j] ~ multinomial(rho[j]); //observation model
  }

  a0 ~ normal(0, 0.05);
  a1 ~ normal(0, 1);
  a1y1star ~ normal(0, 1);
  a2 ~ normal(0, 0.05);
  deltab0 ~ normal(0, 0.02);
  b1 ~ normal(0, 0.2);
  b2 ~ normal(0, 0.05);
  kA ~ beta(2, 2);
  kO ~ beta(2, 2);
}

generated quantities {
  vector[sumnt] log_lik;
  vector[16] pars_;
  int counts_[sumnt, ntaxa] = counts;
  int ranks_[16];
  pars_[1] = a0_[1];
  pars_[2] = a0_[2];
  pars_[3] = a1_[1];
  pars_[4] = a1_[2];
  pars_[5] = a1y1star_[1];
  pars_[6] = a1y1star_[2];
  pars_[7] = a2_[1];
  pars_[8] = a2_[2];
  pars_[9] = deltab0_[1];
  pars_[10] = deltab0_[2];
  pars_[11] = b1_[1];
  pars_[12] = b1_[2];
  pars_[13] = b2_[1];
  pars_[14] = b2_[2];
  pars_[15] = kA_;
  pars_[16] = kO_;
  ranks_[1] = a0[1] > a0_[1];
  ranks_[2] = a0[2] > a0_[2];
  ranks_[3] = a1[1] > a1_[1];
  ranks_[4] = a1[2] > a1_[2];
  ranks_[5] = a1y1star[1] > a1y1star_[1];
  ranks_[6] = a1y1star[2] > a1y1star_[2];
  ranks_[7] = a2[1] > a2_[1];
  ranks_[8] = a2[2] > a2_[2];
  ranks_[9] = deltab0[1] > deltab0_[1];
  ranks_[10] = deltab0[2] > deltab0_[2];
  ranks_[11] = b1[1] > b1_[1];
  ranks_[12] = b1[2] > b1_[2];
  ranks_[13] = b2[1] > b2_[1];
  ranks_[14] = b2[2] > b2_[2];
  ranks_[5] = kA > kA_;
  ranks_[2] = kO > kO_;
  for(j in 1:sumnt) {
    log_lik[j] = multinomial_lpmf(counts[j] | rho[j]); //log likelihood for loo
  }
}
```
