## Supplementary material for "Negative and positive interspecific interactions involving jellyfish polyps in marine sessile communities": Data and code: envdata.stan

```
data {
  int<lower = 1> n; //number of differences between paired observations
  vector[n] y; //vector of differences between paired observations (e.g. temperature at 3 m - temperature at 1 m on same date
}

parameters {
  real<lower = 0> sigma;
  real mu;
}

model {
  y ~ normal();
  mu ~ normal(0, 2);
  //sigma ~ //need to decide on a suitable prior here: but a model for mu conditional on sigma, as in Bayesian Data Analysis, 2nd ed, chapter 3, is probably better
}

generated quantities{
  vector [n] log_lik;
  log_lik = normal_lpdf(y | mu, sigma); //log likelihood for WAIC
}
```
