## Supplementary material for "Negative and positive interspecific interactions involving jellyfish polyps in marine sessile communities": Data and code: run4chains_nointeract.sh

```
#!/bin/bash
for i in {1..4}
do
    ./aureliamanova_nointeract sample num_samples=2500 num_warmup=2500 save_warmup=1 algorithm algorithm=hmc engine=nuts max_depth=20 random seed=12345 id=$i data file=../data/manovadata.txt output file=../../manovaresults/manovanointeract$i.csv refresh=10 &
done
wait
```
