## Supplementary material for "Negative and positive interspecific interactions involving jellyfish polyps in marine sessile communities": Data and code: run4chainscvnodepth.sh

```
#!/bin/bash
for f in ../../leaveoneoutdata/*.txt
do
    for i in {1..4}
    do
	./aureliamanova_nodepth sample num_samples=2500 num_warmup=2500 save_warmup=1 algorithm algorithm=hmc engine=nuts max_depth=20 random seed=12345 id=$i data file=$f output file=$f$i.nodepth.csv &
    done
done
wait
```
