## Supplementary material for "Negative and positive interspecific interactions involving jellyfish polyps in marine sessile communities": Data and code: readme.txt

Code and data to accompany Boughton et al., Negative and positive interspecific interactions involving jellyfish polyps in marine sessile communities.

Matthew Spencer
School of Environmental Sciences
University of Liverpool
Liverpool
L69 3GP
UK


The following instructions were testd on a 64-bit Ubuntu 20.04.4 system with an Intel Core i7-4600M CPU, 15.3 GiB RAM, R 4.1.3, cmdstan-2.27.0. You will need the R packages compositions, RColorBrewer, coda, rstan, mvtnorm, rethinking, hdrcde, scatterplot3d, mvQuad, matrixStats, loo, xtable, deSolve and numDeriv.

Estimation is done using cmdstan, with data manipulation and plotting in R. R code is in aurelia/aureliaproject/R: all the necessary functions are in aureliafuncs.R, while the script that uses them to perform analysis tasks is aurelia.R (with logical variables at the start of the script determining which tasks are carried out when the script is run). Stan code and bash scripts are in aurelia/aureliaproject/stan. The Stan code will need to be compiled before use.

Compositional manova on final composition
-----------------------------------------

1. Open the script aurelia/aureliaproject/R/aurelia.R, and change line 24 to read
setupmanova <- TRUE

2. Open R and type
source("aurelia.R")
to create the data files needed by stan. Then in aurelia.R, change line 24 back to
setupmanova <- FALSE

3. In another terminal, cd to the location of your cmdstan installation and type make [path to aurelia/aureliaproject/stan]/aureliamanova
to compile the full model (with an interaction between depth and treatment), followed by
make [path to aurelia/aureliaproject/stan]/aureliamanova_nointeract
to compile the model without an interaction between depth and treatment,
make [path to aurelia/aureliaproject/stan]/aureliamanova_noscrape
to compile the model without an interaction between depth and treatment or a treatment effect, and
make [path to aurelia/aureliaproject/stan]/aureliamanova_nodepth
to compile the model without an interaction between depth and treatment or a depth effect.

3. cd to aurelia/aureliaproject/stan and type
./run4chains.sh
to fit the full model (with an interaction between depth and treatment).
Then type
./run4chains_nointeract.sh
to fit a model without an interaction between depth and treatment.
Each of these should only take a few minutes.

4. In aurelia.R, change line 24 to read
setupmanova <- FALSE
and change line 32 to read
makeleaveoneoutdata <- TRUE

5. In R, type
source("aurelia.R")
to make leave-one-out cross-validation data sets. Then in aurelia.R, change line 32 back to
makeleaveoneoutdata <- FALSE

6. To carry out leave-one-out cross-validation, cd to aurelia/aureliaproject/stan and type each of the following in turn:
./run4chainscvinteraction.sh
./run4chainscvnointeraction.sh
./run4chainscvnoscrape.sh
./run4chainscvnodepth.sh

NOTE: In each of these shell scripts, if you just want to test the code, you may want to change the values of num_samples and num_warmup (set to 2500 for the manuscript) to smaller numbers such as 250. You should expect each of these to take about 10 times as long as run4chains.sh with num_samples and num_warmup set to 2500.

7. To process the leave-one-out cross-validation output, in aurelia.R, change lines 33 to 38 to read
processleaveoneoutdata <- TRUE
 processleaveoneoutdata <- TRUE #process stan leave-one-out output
processfull <- TRUE #which manova models to process within processleaveoneoutdata block
processnointeract <- TRUE
processnoscrape <- TRUE
processnodepth <- TRUE
comparecv <- TRUE #compare manova models using cross-validation

NOTE: if you just want to test the code, you may want to change the number of iterations and Monte Carlo iterations on lines 228 and 229 to smaller values (e.g. 2 rather than 1e3):
    niter <- 1e3 #number of stan iterations to use
    mciter <- 1e3 #number of Monte Carlo iterations to use to approximate integrals over block and panel effects

With each of these set to 1e3, you might expect the next step to take 48 h or more.

8. Now in R type
source("aurelia.R")
to process the leave-one-out cross-validation output. Once this is done, set the logical variables on lines 33 to 38 back to FALSE.

9. In aurelia.R, change lines 25-27 to
readmanovaresults <- TRUE #extract results for manova from stan output files
loadmanovaresults <- TRUE #load already-extraced manova results from .RData
posteriorpredictive <- TRUE #posterior predictive checks
to look at trace plots, effective sample size and Rhat statistics, and to generate posterior predictive plots (which will be written to aurelia/aureliaproject/diagnosticplots).

In R, type

source("aurelia.R")

to run these. Once this is done, change the logical variables on lines 25 and 27 back to FALSE, but leave loadmanovaresults <- TRUE on line 26.

10. In aurelia.R, change lines 30 and 31 to
plotmanovaresults <- TRUE #plot results from manova (and parameter tables)
hierarchicaldiagnostics <- TRUE #diagnostics for hierarchical parts of model

and then in R type

source("aurelia.R")

to produce figures, summary statistics and tables that appear in the manuscript. The figures will be written as pdfs to aurelia/aureliaproject/latex and aurelia/aureliaproject/diagnosticplots. The table of parameter estimates will be written to aurelia/aureliaproject/latex/manovaparameters.tex.
Once this is complete, set the logical variables on lines 30 and 31 back to FALSE.

11. If you want to carry out simulation-based calibration, in aurelia.R, change lines 28-29 to
doSBC <- TRUE #simulation-based calibration of manova
summarizeSBC  <- TRUE #summarize results of simulation-based calibration of manova

NOTE: this will take a long time (maybe about 32 h) with the parameters used in the manuscript. To test the code, change the parameters M and iter on line 191:

sbcoutput <- SBC(manovastanfile = manovastanfile, manovainputs = manovainputs, ntaxa = ntaxa, V = V, M = 500, iter = 5000)

to smaller values, e.g. M = 10, iter = 10.

Then in R type

source("aurelia.R")

Simulation-based calibration plots will be written to aurelia/manovaresults as pdf files. Once complete, set the logical variables on lines 28-29 back to FALSE.

Analysis of environmental data
------------------------------

1. In aurelia.R, change line 68 to read
analyzeenvironmentaldata  <- TRUE #salinity, temperature, oxygen analyses

and then in R type

source("aurelia.R")

Summary statistics will be printed out, and the figure ../latex/envdata.pdf will be produced. When this is complete, change the logical variable on line 68 back to FALSE.

Models for community dynamics
-----------------------------
1. In aurelia.R, change line 39 to read
setupdynamicmodel <- TRUE #write data for dynamic models in stan

and then in R type

source("aurelia.R")

to create the data files needed for dynamic models. Then change the logical variable on line 39 back to FALSE.

2. In another terminal, cd to the location of your cmdstan installation and type the following:

make [path to aurelia/aureliaproject/stan]/aureliaode
make [path to aurelia/aureliaproject/stan]/aureliaode_overgrowth
make [path to aurelia/aureliaproject/stan]/aureliaode_protection
make [path to aurelia/aureliaproject/stan]/aureliaode_facilitation
make [path to aurelia/aureliaproject/stan]/aureliaode_growthfacil

to compile the basic ODE model and models with overgrowth, protection from predators, facilitation of settlement and facilitation of growth respectively.

3. In a terminal, cd to aurelia/aureliaproject/stan and run the following commands:

./run4chainsode.sh
./run4chainsode_overgrowth.sh
./run4chainsode_protection.sh
./run4chainsode_facilitation.sh
./run4chainsode_growthfacil.sh

to run the dynamic models.
NOTE: with 2500 warmup and 2500 sampling iterations, as used in the manuscript, these may take 1-3 hours each. To test the code, you may want to set num_samples and num_warmup to smaller values (e.g. 250) in each case.

4. In aurelia.R, change line 40 to read

readoderesults <- TRUE #extract results for ODE model from stan output files

then in R, type

source("aurelia.R")

to read the Stan output from the basic ODE model, produce trace plots and print out effective sample sizes and Rhat statistics. When this is done, set the logical variable on line 40 back to FALSE, and repeat setting the logical variables on lines 43 (overgrowth), 46 (protection), 45 (facilitation) and 47 (growth facilitation) in turn.

5. In aurelia.R, change line 65 to read

compareodemodels <- TRUE #compare fitted ODE models

then in R, type

source("aurelia.R")

to compare ODE models based on expected log predictive density. When this is done, set the logical variable on line 65 back to FALSE.

6. In aurelia.R, change line 62 to read

buildtypicalrho <- TRUE #ODE solution for typical panels from each combination

Also change line 51 to read

loadoderesults <- TRUE #load already-extraded ODE results

and make sure that the logical variables on lines 83 to 89 (usefacil to usegrowthfacil) are all set to FALSE.

Then in R, type

source("aurelia.R")

to generate predicted trajectories for typical panels from each treatment. Set the logical variable on line 51 back to FALSE afterwards.

NOTE: generating predicted trajectories may take about 15 minutes per model with niter set to 1e4, as in the manuscript. To test the code, set niter on lines 620, 623, 626, 629, 632, 635, 638, 641 to a smaller value, e.g. 1e3.

7. Repeat step 6, in turn setting the logical variables on the following pairs of lines to TRUE (and then back to FALSE afterwards):

54, 85: for overgrowth model
57, 88: for protection model
56, 87: for settlement facilitation model
58, 89: for growth facilitation model

When done, set line 62 back to

buildtypicalrho <- FALSE #ODE solution for typical panels from each combination

8. The main results in the manuscript are based on the overgrowth model. To use results from this model, change line 54 in aurelia. R to read

loadodeovergrowthresults <- TRUE #load already-extraded ODE results with overgrowth

and change line 85 to read

useovergrowth <- TRUE #use overgrowth in ODE model?

(If you want to work with output from other models, set the corresponding logical variables as in step 6 or 7.)

9. To plot output from the ODE models, change line 63 in aurelia.R to

plotoderesults <- TRUE #plot results from ODE model (requires buildtypicalrho to have been run first). Then in R type

source("aurelia.R")

Plots of the raw time series data (../latex/timeseries.pdf) and time series with fitted ODE models (../latex/timeseries_odeovergrowth.pdf) will be written to pdf files. When done, change the logical variable on line 63 back to FALSE.

10. To check whether the A treatment was also removing potential competitors, change line 64 in aurelia.R to read

checkAremoval  <- TRUE #check whether A treatment was also removing potential competitors

Then in R type

source("aurelia.R")

Plots of potential competitor sample proportions before and after A. aurita removal treatment (../latex/checkAremovaleffects.pdf) and sample proportions of A. aurita before after A. aurita removal treatment (../latex/checkAremovalproportion.pdf) will be written to pdf files. When done, change the logical variable on line 64 back to FALSE.

11. To produce a table of parameter estimates for the ODE model, set line 67 in aurelia.R to read

printODEtable <- TRUE #print ODE parameters to LaTeX

Then in R type

source("aurelia.R")

Parameter estimates will be written to ../latex/ODEparameters.tex. When done, set the logical variable on line 67 back to FALSE.

12. To carry out a graphical posterior predictive check for the ODE model, set line 71 to read

posteriorpredictiveODE <- TRUE #posterior predictive checks for ODE model

Then in R type

source("aurelia.R")

Plots of simulated samples generated from the ODE model will be written to ../latex/posteriorpredictiveovergrowth.pdf. Whe done, set the logical variable on line 71 back to FALSE.


13. To plot the estimated effect of A. aurita on the proportional population growth rate of potential competitors under each ODE model, in aurelia.R change line 72 to read

plotODEeffects <- TRUE #ternary plots of effect of A. aurita on proportional population growth rate of potential competitors

Ternary plots of this effect under each model will be written to ../latex/Aureliaeffects.pdf. This may take a few minutes. When done, change the logical variable on line 72 back to FALSE.

14. To generate simulated data from the ODE model to check estimation, set line 73 in aurelia.R to

setupsimODEdata <- TRUE #generate simulated data sets for checking performance of estimation

Then in R type

source("aurelia.R")

This will generate 10 simulated data sets. When done, set the logical variable on line 73 back to FALSE.

15. To fit ODE models to simulated data, cd to aurelia/aureliaproject/stan and type

./run4chainsode_overgrowth_manysim.sh

NOTE: with 2500 warmup and 2500 sampling iterations, as in the manuscript, this may take approximately 10 h. To test the code, set num_samples and num_warmup to smaller values, e.g. 25. This may still take a while (up to about 50 minutes).

16. To process the fitted ODE results from simulated data, in aurelia.R change lines 74-75 to read

checksimODEdata <- TRUE #process Stan output from simulated data sets
readrawsimoutput <- TRUE #read the raw Stan output from simulated data sets (if FALSE, load processed output from .RData file)

and in R, type

source("aurelia.R")

A figure showing true parameter values, priors and estimated distributions will be written to ../latex/simODEperformance.pdf. When done, set the logical variables on lines 74 and 75 back to FALSE.
